# Transcriptomic responses of the human kidney to acute injury at single cell resolution

**DOI:** 10.1101/2021.12.15.472619

**Authors:** Christian Hinze, Christine Kocks, Janna Leiz, Nikos Karaiskos, Anastasiya Boltengagen, Christopher Mark Skopnik, Jan Klocke, Jan-Hendrik Hardenberg, Helena Stockmann, Inka Gotthardt, Benedikt Obermayer, Laleh Haghverdi, Emanuel Wyler, Markus Landthaler, Sebastian Bachmann, Andreas C. Hocke, Victor Corman, Jonas Busch, Wolfgang Schneider, Nina Himmerkus, Markus Bleich, Kai-Uwe Eckardt, Philipp Enghard, Nikolaus Rajewsky, Kai M. Schmidt-Ott

**Affiliations:** Department of Nephrology and Medical Intensive Care, Charité-Universitaetsmedizin Berlin, Corporate Member of Freie Universitaet Berlin and Humboldt-Universitaet zu Berlin, Berlin, Germany; Max Delbrueck Center for Molecular Medicine in the Helmholtz Association, Berlin, Germany; Berlin Institute for Medical Systems Biology, Max Delbrueck Center in the Helmholtz Association, Berlin, Germany; Deutsches Rheumaforschungszentrum, an Institute of the Leibniz Foundation, Berlin, Germany; Berlin Institute of Health, Berlin, Germany; Core Unit Bioinformatics, BIH/Charité/MDC, Berlin, Germany; Institute for Functional Anatomy, Charité-Universitaetsmedizin Berlin, Corporate Member of Freie Universitaet Berlin and Humboldt-Universitaet zu Berlin, Berlin, Germany; Department of Infectious Diseases and Respiratory Medicine, Charité-Universitaetsmedizin Berlin, Corporate Member of Freie Universitaet Berlin and Humboldt-Universitaet zu Berlin, Berlin, Germany; Institute of Virology, Charité-Universitaetsmedizin, Corporate Member of Freie Universitaet Berlin and Humboldt-Universitaet zu Berlin, Berlin, Germany; Department of Urology, Charité-Universitaetsmedizin Berlin, Corporate Member of Freie Universitaet Berlin and Humboldt-Universitaet zu Berlin, Berlin, Germany; Department of Pathology, Charité-Universitaetsmedizin Berlin, Corporate Member of Freie Universitaet Berlin and Humboldt-Universitaet zu Berlin, Berlin, Germany; Institute of Physiology, Christian-Albrechts-Universitaet, Kiel, Germany

## Abstract

**Background:** Acute kidney injury (AKI) occurs frequently in critically ill patients and is associated with adverse outcomes. Cellular mechanisms underlying AKI and kidney cell responses to injury remain incompletely understood.

**Methods:** We performed single-nuclei transcriptomics, bulk transcriptomics, molecular imaging studies, and conventional histology on kidney tissues from 8 individuals with severe AKI (stage 2 or 3 according to Kidney Disease: Improving Global Outcomes (KDIGO) criteria). Specimens were obtained within 1-2 hours after individuals had succumbed to critical illness associated with respiratory infections, with 4 of 8 individuals diagnosed with COVID-19. Control kidney tissues were obtained post-mortem or after nephrectomy from individuals without AKI.

**Results:** High-depth single cell-resolved gene expression data of human kidneys affected by AKI revealed enrichment of novel injury-associated cell states within the major cell types of the tubular epithelium, in particular in proximal tubules, thick ascending limbs and distal convoluted tubules. Four distinct, hierarchically interconnected injured cell states were distinguishable and characterized by transcriptome patterns associated with oxidative stress, hypoxia, interferon response, and epithelial-to-mesenchymal transition, respectively. Transcriptome differences between individuals with AKI were driven primarily by the cell type-specific abundance of these four injury subtypes rather than by private molecular responses. AKI-associated changes in gene expression between individuals with and without COVID-19 were similar.

**Conclusion:** The study provides an extensive resource of the cell type-specific transcriptomic responses associated with critical illness-associated AKI in humans, highlighting recurrent disease-associated signatures and inter-individual heterogeneity. Personalized molecular disease assessment in human AKI may foster the development of tailored therapies.

## Introduction

Acute kidney injury (AKI) is a frequently observed clinical syndrome, which associates with high morbidity and mortality^1-6^. More than 10% of all hospitalized individuals and more than 50% of critically ill individuals admitted to intensive care units develop AKI^2,3,7^. Despite its extensive clinical and economic impact, AKI therapy is largely limited to best supportive care and kidney replacement therapies (hemodialysis or hemofiltration) in patients with advanced kidney failure^8-10^. Targeted therapies preventing AKI or fostering recovery from AKI are still missing.

Numerous attempts have been made using animal models and human samples to uncover underlying mechanisms of AKI, to identify therapeutic targets and to identify disease biomarkers^11-17^. However, studies in a controlled clinical setting with cell type-specific gene expression resolution of human AKI are lacking.

Although AKI is uniformly defined by changes in serum creatinine levels and/or urinary output, previous studies suggest a vast underlying heterogeneity and complexity of AKI with an unknown number of AKI subtypes, suggesting that personalized approaches in the treatment of AKI may be needed^15,18-20^. Most recently, the question of AKI subtypes was intensively debated when high incidence rates of AKI were observed in individuals with COVID-19^21-24^. In particular the question was raised whether COVID-19 entails a specific molecular subtype of AKI, either through renal viral tropism or other systemic effects^25-29^.

Single-cell gene expression approaches provide powerful tools to investigate cell type-specific changes and cellular interactions and thus may help to delineate potential molecular subtypes of AKI. Recent mouse studies underlined the potential of single cell resolution for our understanding of AKI and revealed new molecular cell states associated with AKI^11,12,30^. Here, we present the first comparative single cell census of the human kidney in individuals with AKI compared to controls without AKI.

## Results

### Single-nuclei RNA sequencing from human kidney samples enables the investigation of cell type-specific gene expression changes in acute kidney injury

To access cellular responses in AKI, we conducted a single cell transcriptome census of human AKI utilizing single-nuclei RNA sequencing (snRNA-seq) of kidney samples from individuals with AKI and control individuals without AKI (Fig. 1A). Kidney samples from individuals with AKI who succumbed to critical illness were obtained within 1-2 hours post-mortem with consent of next of kin. All AKI individuals (n=8, AKI_1-8_) had developed clinical criteria of severe AKI (as defined by KDIGO criteria for AKI stage 2 or stage 3) within 5 days prior to sampling. All individuals had developed AKI in a clinical setting of critical illness, severe respiratory infections, and systemic inflammation, including four cases of COVID-19-associated AKI (Suppl. Table S1). To control for baseline characteristics inherent to human kidneys obtained under clinical conditions and to quantitate the extent of post-mortem effects on gene expression, we used control kidney samples. They included normal kidney tissue collected during tumor nephrectomies (n=3; Control_TN1-TN3_; for clinical parameters of all individuals see Suppl. Table S1). In addition, we obtained post-mortem kidney tissue at three different time points (15 min, 60 min, 120 min) after the cessation of circulation (Control_15 min; 60 min; 120 min_) from a brain-dead individual without clinical evidence of AKI.

**Figure 1:**
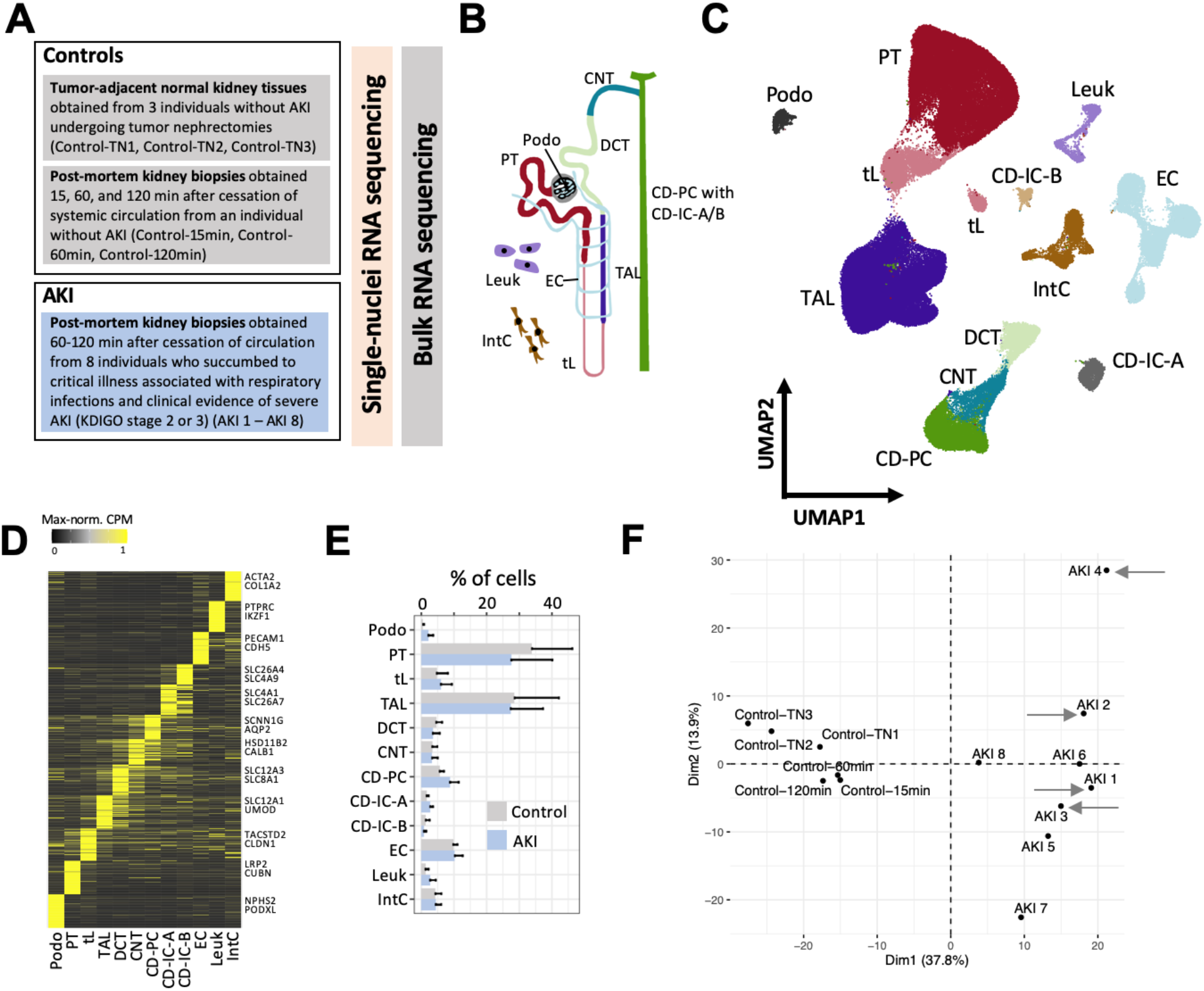
A single cell census of human AKI. **A**. Overview of the study and samples subjected to snRNA-seq and bulk RNA-seq. **B**. Major cell types of the human kidney. (Podo – podocytes, PT – proximal tubule, tL – thin limb, TAL – thick ascending limb, DCT – distal convoluted tubule, CNT – connecting tubule, CD-PC/IC-A/IC-B – collecting duct principal/intercalated cells type A and B, Leuk – leukocytes, IntC – interstitial cells). **C**. Uniform manifold approximation and projection (UMAP) of all kidney cells from snRNA-seq from individuals with AKI and controls. **D**. Heatmap of marker genes of each major cell type. Examples of known cell type marker genes are indicated. Expression values are shown as per-gene maximum-normalized counts per million (CPM). **E**. Relative abundances of major cell types in individuals with AKI and controls (mean and standard deviation). **F**. Principal component analysis of all study individuals using pseudobulk data per individual from all proximal tubule (PT) cells and PT-specific highly variable genes (see Suppl. Fig. S2 for other cell types and whole tissue). COVID-associated AKI cases are highlighted by grey arrows.

Single-nuclei RNA-seq of all samples resulted in 106,971 sequenced cells with a median of 2139 detected genes and 4008 unique transcripts per cell (Suppl. Fig. S1). Joint unbiased clustering and cell type identification with known marker genes allowed the identification of the expected major kidney cell types (Fig. 1B-D). There were no overt differences in major cell type abundances between AKI and controls (Fig. 1E). Principal component analysis (PCA) indicated that the presence of AKI (versus absence of clinical AKI) was the main driver of cell type-specific and global gene expression differences between the samples (Fig. 1F, Suppl. Fig. S2). In contrast, PCA did not identify a major impact of the sampling method (tumor versus post-mortem biopsy), the sampling time after cessation of circulation (15 min, 60 min or 120 min) or the presence of COVID-19-associated AKI (versus AKI associated with other respiratory infections) on global or cell-type-specific gene expression (Fig. 1F, Suppl. Fig S2). Interestingly, we observed heterogeneity of kidney cell gene expression between different individuals with AKI, suggesting different molecular subtypes of AKI (Fig. 1F).

### Kidney tubular epithelial cells from different parts of the nephron show strong gene expression responses to AKI

Kidney ischemia-reperfusion injury in mice, the most frequently applied experimental model of human AKI, results in a predominant injury of cells of the proximal tubule (PT), the most abundant cell type of the kidney. Therefore, many previous studies focused on this cell type^13,14^. However, in humans there is considerable uncertainty regarding the impact of AKI on molecular states of different kidney cell types^31^. To assess the cell type-specific response to AKI systematically, we performed differential gene expression analysis within the major kidney cell types comparing AKI to control kidneys using DESeq2^32^ (see Suppl. Table S2 for a full list of differentially expressed genes per cell type). Profound transcriptomic responses to AKI were observed in kidney tubule cells of the PT, the thick ascending limb of the loop of Henle (TAL), the distal convoluted tubule (DCT), and connecting tubules (CNT), cell types that reside predominantly in the cortex and outer medulla of the kidney, regions that are known to be particular susceptible to ischemic or hypoxic injury^13-15,33^ (Fig. 2A). In contrast, less pronounced transcriptomic responses to AKI were observed in thin limbs (tL), collecting duct principal cells (CD-PCs) and collecting duct intercalated cells (CD-ICs), consistent with the predominant localization of these cell types in the inner medulla of the kidney, which is adapted to a low oxygen environment, has lower energy expenditure and is less susceptible to hypoxia or ischemia when compared to more cortical regions^33^. Podocytes, endothelial cells and interstitial cells also displayed less pronounced transcriptomic responses in AKI.

**Figure 2:**
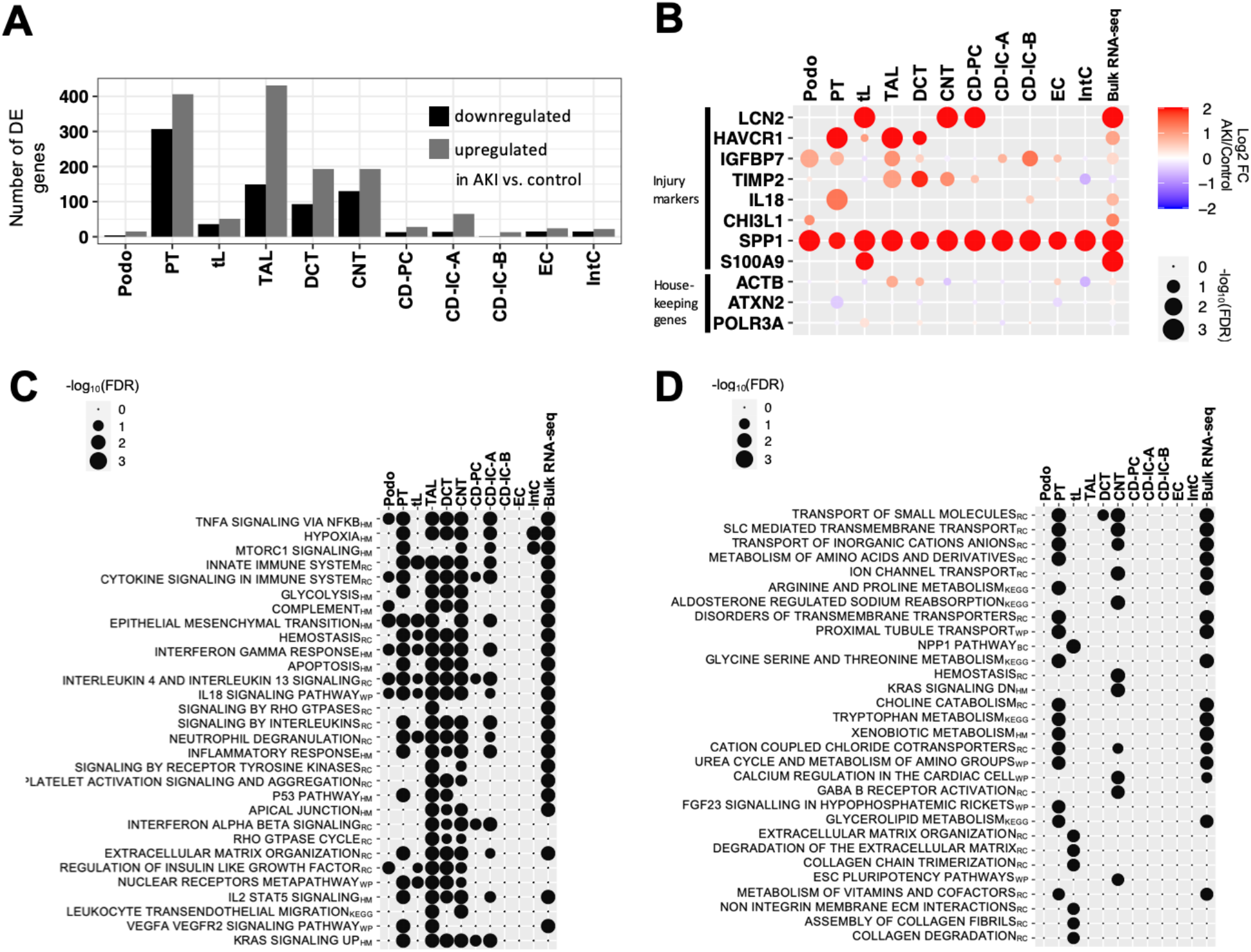
Cell type-specific responses of kidney cells to acute injury. **A**. Absolute numbers of differentially expressed (DE) genes upregulated and downregulated in AKI versus controls within major kidney cell types. **B**. Dot plot displaying the degree of differential expression for known injury marker genes and housekeeping control genes (actin beta (ACTB), ataxin 2 (ATXN2) and RNA polymerase III subunit A (POLR3A)). **C**,**D**. Dot plot for top enriched pathways (defined by FDR) in genes upregulated (C) and downregulated (D) in AKI versus controls. HM – Molecular Signatures Database (MSigDB) hallmark gene sets, MSigDB canonical pathway gene sets derived from RC – Reactome, WP – WikiPathways and KEGG -Kyoto Encyclopedia of Genes and Genomes.

Among differentially expressed genes were known markers of renal cell stress, which encode proteins that have been proposed as kidney injury markers, such as neutrophil gelatinase-associated lipocalin/lipocalin 2 (LCN2), kidney injury molecule 1 (HAVCR1), and insulin-like growth factor binding protein 7 (IGFBP7)^34^. Importantly, our data provided the opportunity to identify the major cellular sources where these transcripts were synthesized in response to injury. For instance, consistent with previous reports on mouse and human AKI, LCN2 was primarily upregulated in CNT and CD-PC^12,35^, while HAVCR1 was primarily upregulated in PT^12,36^ (Fig. 2B). Secreted phosphoprotein 1 (SPP1), encoding for the secreted glycoprotein osteopontin, was found to be upregulated in virtually all non-leukocyte kidney cell types. This was strongly reminiscent of the situation in mouse AKI, where osteopontin upregulation was similarly observed in multiple kidney cell types^12^, and where osteopontin inhibition attenuated renal injury^37^, suggesting a conserved, targetable AKI pathway. IGFBP7 protein was previously found to be primarily of PT origin in diseased human kidneys^38^. Consistently, we found an upregulation of IGFBP7 mRNA in PT cells (Fig. 2B). However, we also found IGFBP7 to be upregulated in podocytes and TALs (Fig. 2B), findings which we were able to validate by IGFBP7 in situ hybridization (Suppl. Fig. S3). This indicates that our single cell transcriptome database is consistent with prior knowledge and provides an opportunity to uncover novel information regarding the cellular origin of AKI-associated transcripts.

Pathway analyses of differentially expressed genes indicated that a proportion of genes upregulated in AKI were associated with inflammatory response-associated pathways (tumor necrosis factor alpha, interferon gamma and interleukin signaling), hypoxia response, and epithelial to mesenchymal transition (EMT, Fig. 2C, Suppl. Table S3). Importantly, our analyses indicated that most functional pathways were upregulated simultaneously in multiple kidney tubule cell types suggesting common AKI response patterns across the nephron. Several studies have indicated an AKI-associated metabolic shift in tubular epithelia and a downregulation of genes associated with tubular transport processes^33,39^. Consistently, we observed that genes downregulated in AKI were mostly related to molecule transport and metabolism (Fig. 2D).

Since our cohort included four individuals with COVID-19-associated AKI, we compared their kidney cell type-specific gene expression with that of individuals with non-COVID-19 respiratory infection-associated AKI. Only few differentially expressed genes were identified, suggesting that the major transcriptomic responses of kidney cells in COVID-19 were not substantially different from those in other forms of AKI (see Suppl. Table S4 for the respective gene lists).

Importantly, the genes and pathways that were differentially regulated in AKI versus control according to single nuclei sequencing data displayed concordant regulation in bulk mRNA sequencing from separate kidney samples of the same patients, providing additional validation (Fig. 2B, C, D).

### Profound effects of AKI on kidney cell state abundance

To achieve a more fine-grained analysis of cellular subclasses, we performed subclusterings of the major kidney cell types. Thereby, we were able to derive 74 kidney cell populations based on their transcriptomes, which included known cellular subtypes of kidney cells (e. g. S1, S2, and S3 segments of the PT) and additional novel cell populations (designated as “New” cell populations, Fig. 3, Suppl. Fig. S4-5). These “New” cell populations were still attributable to major kidney cell types based on their transcriptomes (Fig. 1C), but they were not characteristic of the known anatomic sub cell types, suggesting that they represent injury-related cell states.

**Figure 3:**
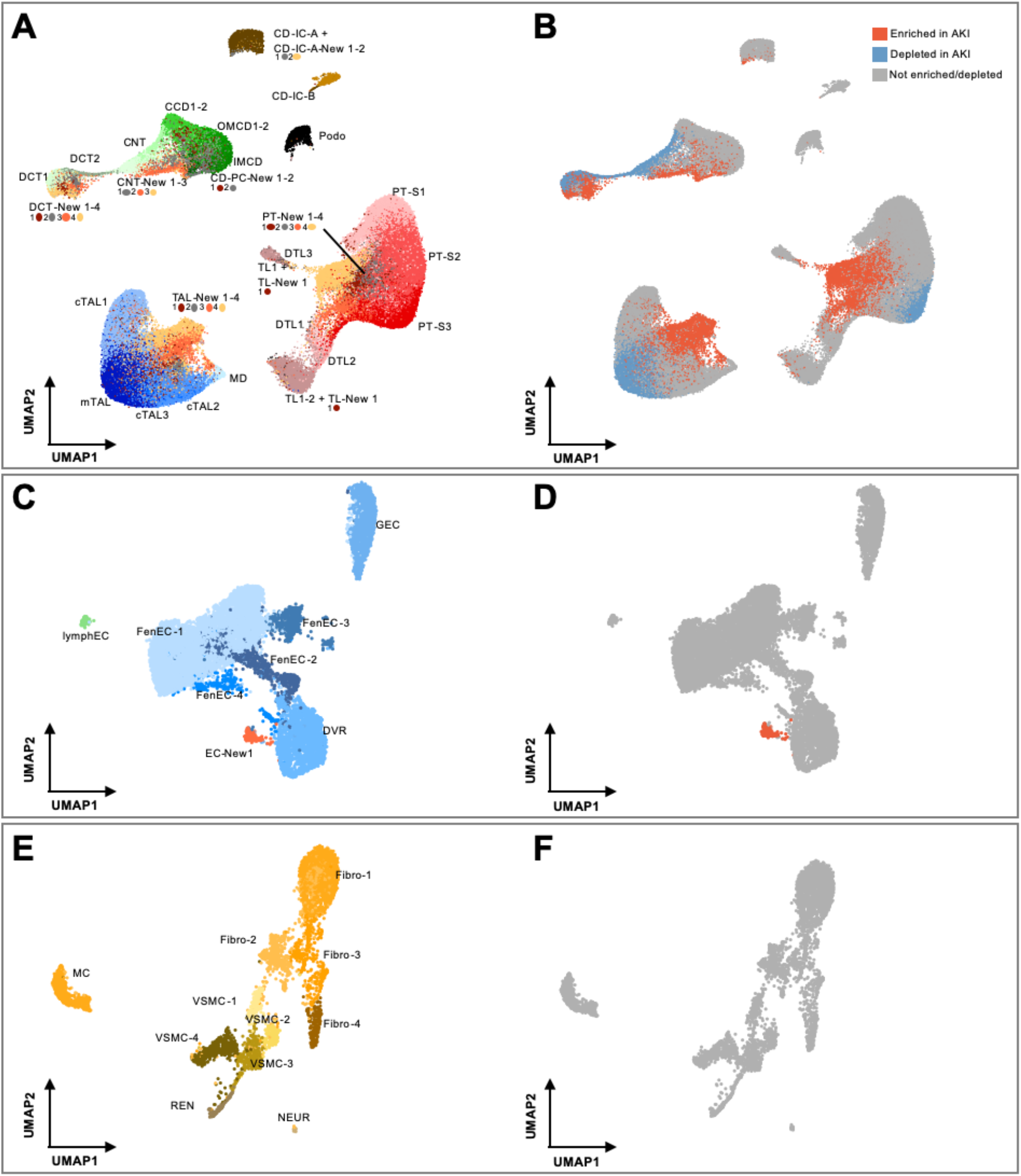
AKI leads to depletion of differentiated cell states and enrichment of “New” cell states within the kidney epithelium. **A, B**. UMAP plot of subclustered kidney tubular epithelial cells (A) and their enrichment or depletion in AKI based on statistical testing of relative abundances within the respective major cell type (B) (see methods for details). In A, cellular subtypes of the kidney tubule are annotated as indicated. To enhance visibility, color code is indicated below the respective labeling. In B, the same UMAP plot as in A is color-coded based on enrichment (red) or depletion (blue) in AKI individuals. **C, D**. Analogous plots for subclustering of endothelial cells (ECs). Please note the emergence of one AKI-associated subcluster, EC-New 1. **E, F**. Analogous plots for subclusterings of interstitial cells. PT-S1-3 – PT S1-3 segments, c/mTAL – cortical/medullary TAL, TL, DTL – thin limb and descending thin limb, CCD, OMCD, IMCD – cortical/outer and inner medullary collecting duct principal cell; lymphEC – lymphatic EC, GEC – glomerular EC, FenEC 1-4 – fenestrated endothelial cell types, DVR – descending vasa recta; MC – mesangial cells, VSMC – vascular smooth muscle cells, REN – renin-transcribing cells, Fibro – fibroblasts, NEUR – neuronal cells.

We analyzed, which of the identified cell subpopulations differed in abundance in individuals with AKI, yielding depleted and enriched subpopulations (Fig. 3B). Profound depletion in AKI was observed within cells of the PT (in particular those representing the S3 segment), consistent with its known susceptibility to injury and its tendency to undergo dedifferentiation in AKI^11,13,14,40^. Unexpectedly, in addition to PT, differentiated medullary TAL, DCT, CNT cells were also substantially depleted in AKI. Inversely, profound enrichment in AKI was observed of the “New” cell subpopulations associated with these same cell types, indicating that PT, TAL, DCT and CNT displayed the most profound responses to AKI and confirming the notion that “New” subpopulations represent injury-associated cell states (Fig. 3B). “New” subpopulations within cells of collecting duct (CD-PC, CD-IC-A, CD-IC-B), ascending and descending thin limbs (ATL, DTL) were also enriched in AKI, although they represented only small subpopulations (Fig. 3B), suggesting that these cell types are less susceptible or reside in less susceptible regions of the kidney. Non-epithelial cell types of the kidney, such as endothelial cells, interstitial cells and leukocytes displayed no enrichment or depletion in AKI, with the exception of one subtype of endothelial cells (EC-New 1), which was enriched in AKI and showed a transcriptional profile resembling endothelia of descending vasa recta (Fig. 3C-F; Suppl. Fig. S4).

### Analyses of AKI-induced cell states suggest four distinct injury response patterns

We next conducted further analyses to characterize the “New” cell subpopulations detected within the tubular epithelial compartment of the kidney. Quantification of the four “New” cell clusters associated with the PT (PT-New 1-4) indicated that almost one third (31.8%) of PT cells in AKI samples belonged to these clusters (Fig. 4A). We next identified marker genes for PT-New 1-4 (Fig. 4B) and performed pathway analysis. Enriched gene sets included oxidative stress signaling and the nuclear transcription factor erythroid 2-related factor 2 (NRF2) pathway (PT-New 1), the hypoxia response pathway (PT-New 2), the interferon gamma response and genes encoding for ribosomal proteins (PT-New 3) as well as genes associated with epithelial-mesenchymal transition (EMT) (PT-New 4) (Fig. 4B, see https://www.gsea-msigdb.org, Hallmark and canonical pathways, for pathway definitions). Nevertheless, PT-New 1-4 showed some overlap and trajectory analysis using partition-based graph abstraction (PAGA)^41^ suggested hierarchical relationships between healthy PT cells and cells representing PT-New 1-4, with PT-New 4 displaying the most distant gene expression signature from healthy PT.

**Figure 4:**
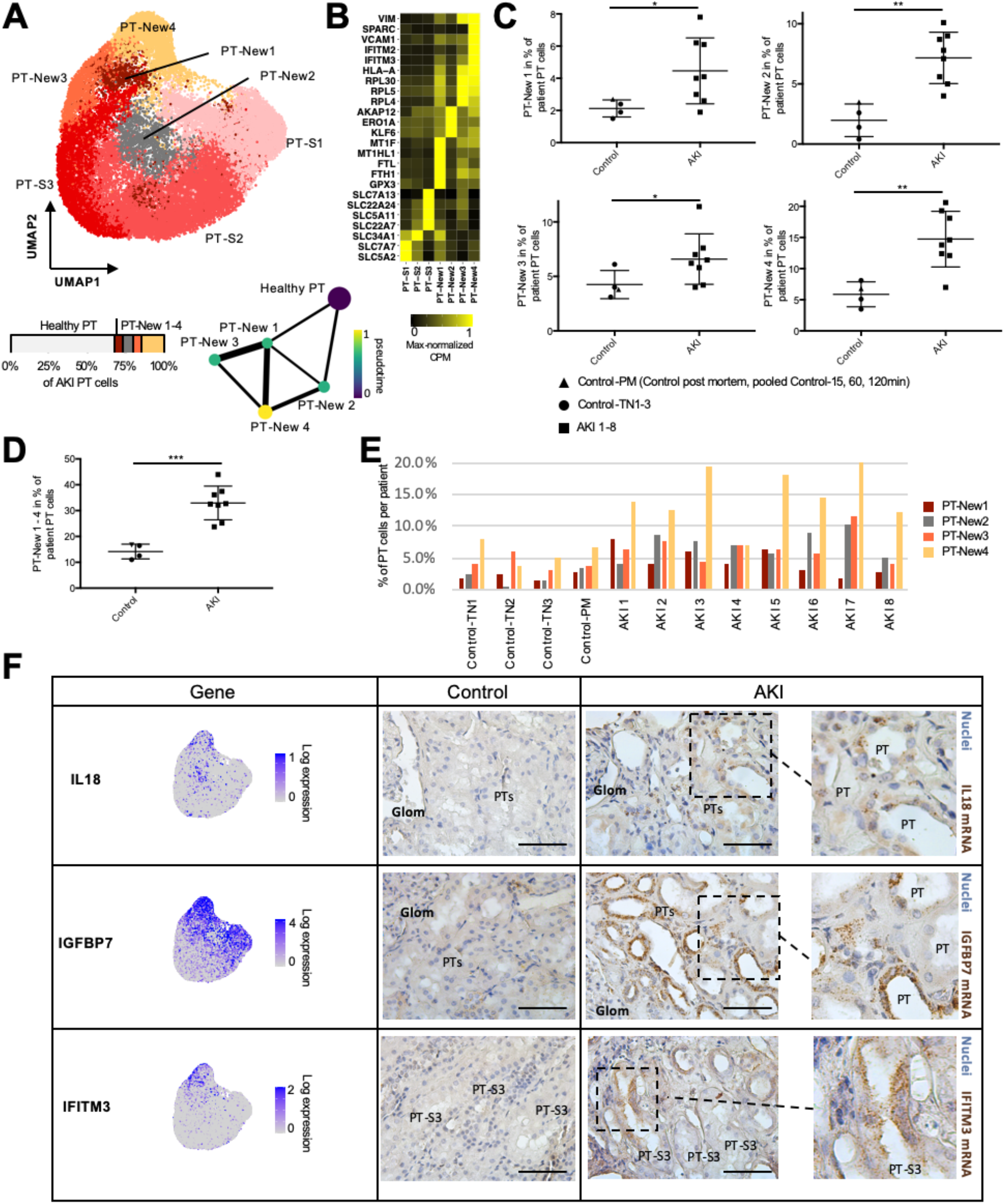
PT AKI-enriched cell states reveal four distinct injury response patterns. **A**. UMAP plot of subclustering of the PT with the anatomical PT segments S1-3 (PT-S1-3) and the AKI-associated cell states PT-New 1-4. Below the UMAP are a bar plot displaying the relative abundances of PT-New 1-4 with respect to all AKI PT cells and a trajectory analysis using partition-based graph abstraction (PAGA) highlighting diffusion pseudotime. Line widths of the connecting edges represent statistical connectivity between the nodes^41^. Healthy PT-S1-3 were summarized to healthy PT for this analysis. **B**. Heatmap of selected marker genes for the identified PT cell subpopulations. **C**. Plots display relative abundance of PT-New 1-4 as percentage of all PT cells. **D**. Plot displays relative abundance of combined PT-New 1-4 as percentage of all PT cells. **E**. Individual abundances of PT-New 1-4 for control and AKI individuals. **F**. Feature plots on the UMAP presented in A. and RNAscope in situ hybridizations of injury markers interleukin 18 (IL18), insulin-like growth factor binding protein 7 (IGFBP7) and interferon-induced transmembrane protein 3 (IFITM3). Scale bar: 50 µm. P-value: *<0.05, **<0.01, ***<0.001, n.s. – not significant. Control-PM – pooled samples (Control_15 min_, Control_60 min_, Control_120 min_) of post mortem non-AKI control individual.

We compared PT-New 1-4 to previously identified PT-derived injured cell states in mouse renal ischemia-reperfusion injury, designated as “injured PT S1/S2 cells”, “injured PT S3 cells”, “severe injured PT cells” and “failed repair” cells^11^. We trained a multinomial model using marker genes of these clusters using a cross-species mouse/human comparative approach (see methods), which indicated that PT-New 1 (oxidative stress) showed similarity to injured mouse S1/2 cells, whereas PT-New 2 (hypoxia) resembled injured S3 cells (Suppl. Fig. S6). PT-New 3 and PT-New 4 most closely resembled “failed repair” PT cells in mice (Suppl. Fig. S6). Cells from PT-New 3 and PT-New 4 expressed the EMT marker VIM. PT-New 4 cells were additionally marked by VCAM1, a marker that has previously been associated with the “failed repair” state of injured PT cells^42^. PT-New 1, PT-New 3 and PT-New 4 also showed high expression of interferon target genes (e. g. IFITM2, IFITM3) and human leukocyte antigen HLA-A, which is consistent with the association of injured PT cells with immune responses and inflammation^42^. Together these observations suggest that “New” cell populations represent four distinct but hierarchically connected injured PT cell states.

The individual abundances of PT-New 1-4 and the combined abundances of PT-New 1-4 were significantly increased in the AKI samples (Fig. 4C, D). Importantly, the distribution of PT-New 1-4 among individuals with AKI displayed marked heterogeneity (Fig. 4C, E). For instance, the relative abundance of PT-New 4 (EMT/ “failed repair”) varied by a factor of three among samples from different individuals with AKI (compare PT-New 4 between AKI 4 and AKI 7 in Fig. 4E). The combined injury-associated PT clusters (PT-New 1-4) made up between 20% and 45% of all proximal tubule cells in individuals with AKI (Fig. 4C). Together these observations highlight the presence of recurrent AKI-associated cell states, but they also indicate substantial inter-individual heterogeneity.

To localize “New” PT-associated clusters in injured kidney tissues, we performed RNAscope *in situ* hybridizations for transcripts overexpressed in PT-New 1-4 clusters (Fig. 4F). Interleukin-18 (IL18) mRNA was specifically expressed in occasional cells within clusters PT-New 2 and PT-New 4. Consistently, in situ hybridizations indicated that IL18 mRNA was expressed in a small subpopulation of PT cells of AKI kidneys compared with little or no expression in control kidneys (Fig. 4F). In contrast, insulin-like growth factor binding protein 7 (IGFBP7) was widely expressed in PT-New 1-4 cells. Consistently, in situ hybridizations of AKI kidneys demonstrated IGFBP7 mRNA expression in a substantial subset of PT cells (Fig. 4F), while only few PT cells of control kidneys expressed IGFBP7. Interferon-induced transmembrane protein 3 (IFITM3) was expressed in a subset of cells within PT-New 1, 3, and 4. In situ hybridization confirmed IFITM3 expression within severely injured PT cells of AKI kidneys, while IFITM3 was undetectable in control kidneys (Fig. 4F).

### The abundance of injury response patterns varies among cell types of the kidney tubule

We next turned to other kidney epithelial cell types and their response to injury. We compared AKI-enriched “New” cell states in tL, TAL, DCT, CNT, CD-PC and CD-IC to those in PT (Fig. 5, Suppl. Fig. S7-9). Remarkably, the transcriptomic responses of the different tubular epithelial cell types to AKI displayed a marked overlap. For instance, “New” cell populations residing in TAL (TAL-New 1-4) and DCT (DCT-New 1-4) displayed 4 injured cell states with marker genes and functional pathways similar to PT-New 1-4 (Suppl. Table S5). This suggests conserved injury responses across different kidney cell types. The percentage of cells displaying AKI-associated “New” cell states varied markedly among major cell types of the kidney: 31.8% of PT cells; 36.4% of TAL cells; 59.6% of DCT cells; 43.5% of CNT cells; 2.1% of tL cells; 19.1% of CD-PCs and 5.7% of CD-ICs (Fig. 4-5, Suppl. Fig. S7-9).

**Figure 5:**
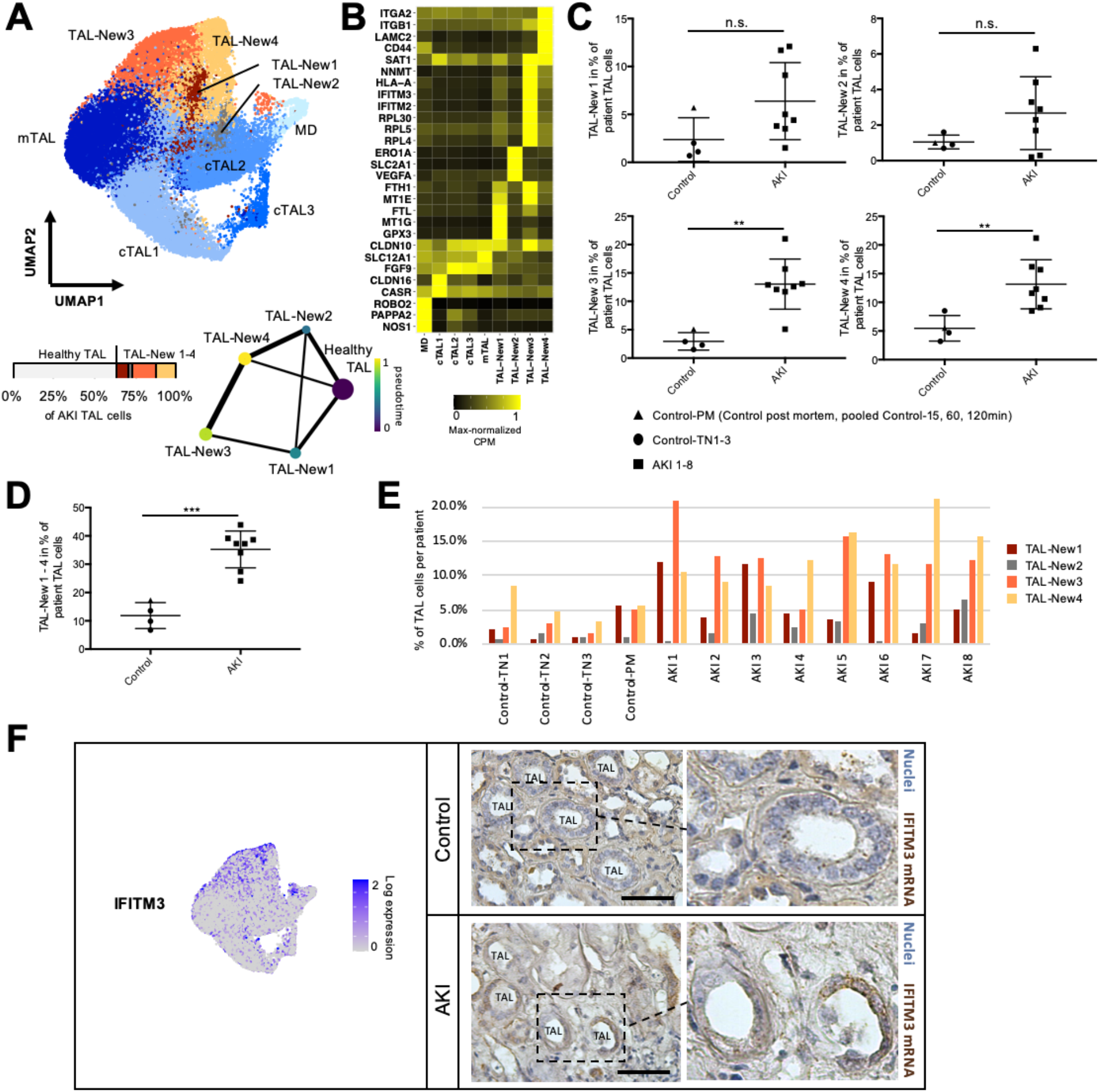
AKI-associated cell states within the thick ascending limb (TAL). **A**. UMAP plot of TAL subclustering with the anatomical segments cTAL 1-3 and mTAL and the AKI-associated cell states TAL-New 1-4. Below the UMAP are a bar plot displaying the relative abundances of TAL-New 1-4 with respect to all AKI TAL cells and a trajectory analysis using partition-based graph abstraction (PAGA) highlighting diffusion pseudotime. Line widths of the connecting edges represent statistical connectivity between the nodes^41^. Healthy cTAL 1-3 and mTAL were summarized to healthy TAL for this analysis. **B**. Heatmap of selected marker genes for the identified TAL cell subpopulations. **C**. Plot displaying relative abundances of TAL-New 1-4 with respect to individual’s TAL cells. **D**. Relative abundances of combined TAL-New 1-4 cells with respect to individual’s TAL cells **E**. Individual abundances of TAL-New 1-4 for control and AKI individuals. **F**. Feature plot on the UMAP presented in A. and RNAscope in situ hybridization of interferon-induced transmembrane protein 3 (IFITM3). Scale bar: 50 µm. P-value: *<0.05, **<0.01, ***<0.001, n.s. – not significant. Control-PM – pooled samples (Control_15 min_, Control_60 min_, Control_120 min_) of post mortem non-AKI control individual.

Similar to the PT, we also observed inter-individual heterogeneity for the AKI-associated cell states in other cell types of the kidney tubule (Fig. 5E, Suppl. Fig. S7-9). Although, similar clusters as PT-New 1-4 were present in the other kidney tubule cell types, their relative abundance was cell type-dependent. In PTs and DCTs, the most abundant AKI-associated cell state was that associated with EMT (PT-New 4 and DCT-New 4). In contrast, the most abundant TAL cell state was TAL-New 3 (interferon gamma signaling-associated). IFITM3, an interferon-associated gene and marker gene of TAL-New 3, was expressed in injured TALs of AKI kidneys when analyzed by in situ hybridization (Fig. 5F). We conclude that injury responses in different epithelial cell types of the kidney associate with common molecular pathways and marker genes, although they display cell type-specific and inter-individual heterogeneity.

## Discussion

This study identifies a strong impact of human AKI associated with critical illness on the kidney transcriptome at single cell resolution. We provide an atlas of single cell transcriptomes and uncover novel AKI-induced transcriptomic responses at unprecedented cellular resolution and transcriptional depth. We find that the dominant AKI-associated transcriptomic alterations reside within the different cell types of the kidney’s tubular epithelium with surprisingly few transcriptomic alterations in other cell types. We uncover four AKI-associated “New” transcriptomic cell states, which emerge abundantly in PT, TAL and DCT and display a remarkable overlap of marker genes and enriched molecular pathways between these different tubular epithelial cell types, suggesting common injury mechanisms. Finally, we report a strong transcriptional heterogeneity among individuals with AKI, which is explained by the inter-individual differences in cell type-specific abundances of injury-associated cell states.

Our study provides insights into the pathophysiology of human AKI and indicates injury-associated responses in several different types of tubular epithelial cells. Previous studies of AKI in animal models (mostly ischemia-reperfusion injury in mice) have focused on the PT because cells of the PT are abundant and because PTs display marked susceptibility and responsiveness to ischemic injury^13,14,31,33,43-45^. Fewer studies have examined effects of AKI on other cell types. However, some previous rodent studies also showed evidence of injury in distal parts of the kidney tubule including TALs^46^ and collecting ducts^47^, similar to what we demonstrate here in human AKI. Presumed mechanisms of tubular injury and tubular transcriptome responses are related to the high metabolic demands of tubular cells, in particular proximal tubules and TALs, which become overwhelmed in the setting of ischemia or hypoxia ^31,46,48,49^. Tubular stress in this setting is documented by novel induction of injury-associated transcripts and down-regulation of differentiation markers (e. g. solute transporters). Most recently, transcriptome studies at single cell resolution in mouse AKI models confirmed these findings and indicated predominant cellular responses in the proximal tubule compartment of the kidney^11,12,30^. Our human AKI data are consistent with the proximal tubular responses described in mice, but they suggest an unexpectedly widespread molecular response in other types of kidney tubular epithelia, including TAL, DCT, CNT and collecting duct. It is tempting to assume that this difference reflects the more complex, multifactorial pathogenesis of AKI in critically ill patients compared to ischemia reperfusion models, but additional interspecies differences can so far not be excluded.

Given the observed induction of AKI-associated cell states in different segments of the kidney tubule, questions arise regarding their functional significance. It is noteworthy that we found a particularly pronounced activation of NRF2 target genes in the “New 1” clusters in several cell types (PT-New 1, TAL-New 1, DCT-New 1, CNT-New 1, CD-PC-New 1, TL-New 1). NRF2 signaling is induced by oxidative stress, plays a role in the induction of antioxidative genes and has been associated with protection from kidney injury^50-52^. Trajectory analyses and comparative genomics indicated that these cells were likely derived from S1/2 segments of the PT and represent an early stage of injury. Cells of the “New 2” cluster exhibited a hypoxia signature. Hypoxia has also previously been recognized as an important mechanism of AKI, due to low baseline tissue oxygen concentrations in the kidney, which further decline under conditions leading to AKI. Induction of hypoxia-inducible genes through stabilization of hypoxia-inducible transcription factors in different kidney cell populations or selectively in TAL or PT cells was found to ameliorate ischemic or toxic AKI^33,53^. In contrast, cells of the “New 3” and “New 4” clusters expressed an EMT signature and pro-inflammatory genes and resembled cells previously described as a “failed repair” state, which is associated with progression to kidney fibrosis^11,42^. In trajectory analyses these cells were transcriptionally most distant from healthy tubular epithelium in PT and TAL.

Our study included patients with COVID-19-associated AKI. There is an ongoing debate on the mechanisms of AKI in COVID-19, particularly with regard to whether there is a COVID-19-specific kidney pathophysiology that is different from other critical illness-associated forms of AKI^26,28,54,55^. Our study did not uncover a specific transcriptomic signature associated with COVID-19-associated AKI and suggests that COVID-19 AKI is on a common molecular spectrum with AKI associated with other types of respiratory failure and critical illness.

In summary, we observed that AKI in humans with critical illness and systemic inflammation is associated with widespread transcriptomic responses within a spectrum of kidney cell types, uncovering novel cell states and potential targets for AKI therapies. These findings suggest that precision approaches like single cell transcriptomics maybe suitable tools to overcome the current limitations in diagnosing and treating subtypes of AKI.

## Methods

### Specimen collection

After consent of next of kin, post mortem biopsies were collected using 18G biopsy needles within 2 hours from death from individuals who had died in a clinical setting of critical illness on intensive care units of Charité-Universitaetsmedizin Berlin (ethics approval EA2/045/18) (see Suppl. Table S1 for detailed patient characteristics). Control tissue from tumor-adjacent normal tissue of tumor nephrectomies was collected during tumor nephrectomies (ethics approval EA4/026/18). Kidney specimen were either stored in pre-cooled RNAlater at 4°C for 24 hours and then stored at −80°C (for snRNA-seq) or in 4% formaldehyde (for histopathological studies and in situ hybridizations).

### Single-nuclei sequencing

Kidney specimen subjected to snRNA-seq were kept at 4°C at all times. All specimens were treated as described in detail in Leiz, Hinze et al. 2021^56^.

Main steps included: Specimens were thoroughly minced in nuclear lysis buffer 1 (nuclear lysis buffer (Sigma) + Ribolock (1U/µl) + VRC (10mM)) and homogenated using a dounce homogenizer with pastel A (Sigma D8938-1SET), filtered (100 µm), homogenized again (douncer with pastel B), filtered through a 35 µm strainer and centrifuged (5 min, 500g). The pellet was then resuspended in nuclear lysis buffer 2 (nuclear lysis buffer + Ribolock (1U/µl)). To remove debris from the suspension, we underlayed the suspension with lysis buffer containing 10% sucrose and 1U/µl of Ribolock. After centrifugation (5 min, 500g), the supernatant and debris were carefully removed. Pelleted nuclei were resuspended in PBS/0.04%BSA + Ribolock (1U/µl), filtered through a 20 µm strainer and stained with DAPI.

All samples were subjected to single-nuclei sequencing following the 10x genomics protocol for Chromium Next GEM Single Cell 3’ v3.1 chemistry targeting 10000 nuclei. Obtained libraries were sequenced on Illumina HiSeq 4000 sequencers (paired-end). Digital expression matrices were generated using the Cellranger software version 3.0.2 with –force-cells 10000 against a genome composed of the human HG38 genome (GRCh38 3.0.0).

### Single-nuclei sequencing data analysis

Initial filtering was performed by excluding nuclei with more than 5% mitochondrial reads and less than 500 detected genes. Nuclei passing this filter of all samples were then analyzed using Seurat’s best practice work flow for data integration using the reciprocal PCA approach (https://satijalab.org/seurat/articles/integration_rpca.html) with default parameters. Emerging clusters were then analyzed for marker gene expression using Seurat’s FindAllMarkers function and subsequently assigned to the major renal cell types. Each major cell type was then subclustered using Seurat’s best practice standard work flow for data integration:(https://satijalab.org/seurat/articles/integration_introduction.html). Followed by RunUMAP(seu, dims = 1:30), FindNeighbors(seu, dims = 1:30) and FindClusters(seu, resolution = 0.5) using the integrated assay. Marker genes for all emerging clusters were calculated.

During the next steps, the aim was to identify destroyed nuclei and doublets. For the destroyed nuclei, in the initial clustering, they clearly clustered away from major kidney cell types and showed a significantly reduced complexity of gene expression (nUMI, nGene) when compared to major cell types. Doublets could be detected by clusters in the subclusterings of the major cell types showing canonical marker genes from another major cell type (e.g. TAL marker gene expression in a PT subcluster). Destroyed and doublet cells were removed and the whole clustering process was repeated to avoid any influence of the excluded cells on the clusterings.

Enrichment testing for all major cell types was performed by calculating relative abundances of each generated subcluster per patient as percent of the respective major cell type. A p-value was computed by using a Mann-Whitney-U test comparing relative abundances of AKI samples versus control samples for each cluster and each major cell type. Enrichment scores were calculated by −log10 transforming the p-values and considered significant if p-value<0.05. The three replicates from Control-PM were averaged to one value per comparison.

### RNA extraction and alignment for bulk RNA sequencing

The RNeasy Micro Kit (#74004, Qiagen, Hilden, Germany) was used to extract total RNA from kidney biopsies stored at −80°C. For tissue disruption frozen biopsy samples were transferred to ceramic bead-filled tubes (#KT03961-1-102.BK, Bertin Technologies, Montigny-le-Bretonneux, France) containing 700 µl QIAazol Lysis reagent (#79306, Qiagen) and homogenized for 2x 20 sec at 5000 rpm using a Precellys 24 tissue homogenizer (Bertin Technologies).

The lysate was incubated for 5 min at room temperature, mixed with 140 μl chloroform, and centrifuged (4 °C, 12,000 x g, 15 min). The supernatant was transferred to a fresh tube. Subsequent RNA purification was performed according to the “Purification of Total RNA from Animal and Human Tissues” protocol starting at step 4 (https://www.qiagen.com/de/shop/pcr/rneasy-micro-kit/#resources).

RNA concentration and integrity were evaluated with a NanoDrop Spectrophotometer (Thermo Scientific, Waltham, MA) and 2100 Bioanalyzer Instrument (Agilent Technologies, Santa Clara, CA). Paired-end RNA sequencing (Truseq stranded mRNA, 2×100bp) was performed on a Novaseq 6000 SP flow cell. Provided FASTQ files were aligned using STAR and the same genome as for the snRNA-seq data (GRCh38 3.0.0 with SARS-CoV1/2), reads were then counted using featureCounts^57^ with -p -t exon -O -g gene_id -s 0.

### Differential gene expression analysis

Differential gene expression analysis was performed using the DESeq2 pacakge (version 1.28.1). Input to DESeq2 were count matrices generated by adding the counts from all cells per sample and major cell type (for snRNA-seq) or the raw count matrices (for bulk RNA-seq). For snRNA-seq, only genes expressed in 10% or 500 cells were considered. DESeq dataset was generated by:

dds <- DESeqDataSetFromMatrix(countData = my.count.data, colData = col.data, design = ∼ condition + perc.mt),

followed by:

dds <- estimateSizeFactors(dds, type=‘poscounts’)

dds <- DESeq(dds)

“condition” was either AKI or control, my.count.data was generated for each major cell type, separately (snRNA-seq). Desired comparisons were derived by:

my.result = results(dds, contrast=c(“condition”,”Control”,”AKI”))

Results were then filtered by demanding adjusted p-value<0.001 and absolute log2 fold change larger than 2.

### Pathway enrichment analysis

Differentially expressed genes were analyzed, separately, for genes up- and downregulated in AKI versus controls using the MSigDB web interface:

http://www.gsea-msigdb.org/gsea/msigdb/annotate.jsp with the hallmark gene sets and curated gene sets (C1) from Biocarta, Kegg, Reactome and WikiPathways.

### Partition-based graph abstraction

The full integrated assay from the Seurat object of the respective cell type were imported into scanpy version 1.5.0. PCA and neighborhood graph were computed:

~~~
sc.tl.pca(data, use_highly_variable=False)
sc.pp.neighbors(data, n_pcs=30)
~~~

Healthy subclusters (e.g. PT-S1-3) were summarized into one category (e.g. Healthy PT) to exclude the anatomical axis from this analysis. PAGA graph and diffusion pseudotime (dpt) were calculated. The root of diffusion pseudotime was defined to be in the first element of the healthy cells (e.g. first element in healthy PT). To further exclude the anatomic axis from pseudotime analysis, all dpt values for the healthy cells were set to 0. PAGA graph was then plotted with threshold 0.15 highlighting dpt. Dpt was min-max normalized for plotting.

### Cross-species approach

Mouse ischemia reperfusion data was downloaded from GEO (GSE139107)^11^, meta data of PT subclustering from the original publication was kindly provided by Ben Humphreys. Multinomial classification was performed using the glmnet package version 2.0-16. The training set included randomly selected cells (2/3 of cells) from injured mouse PT subclusterings (Suppl. Fig. S6). Genes used in the training were highly variable features of the mouse AKI data. Human orthologous genes were generated using Biomart. Test data were the remaining 1/3 of mouse PT injured subclusters. Glmnet produces different models for different values of lambda which determines how hard overcomplexity of the respective model gets punished. Each so-generated model was tested on the test data and the model with the highest accuracy on the test data was determined. The so selected model was then applied to our human PT subclusters PT-New 1-4.

### RNA *in situ* hybridization - RNAscope

The RNAscope 2.5 HD reagent kit-brown (#322300, Advanced Cell Diagnostics (ACD), Newark, CA) was used to perform chromogenic *in situ* hybridization on formalin-fixed paraffin embedded mouse kidney sections with probes directed against *IFITM3* (#1062531-C1, ACD), *IGFBP7* (#316681, ACD) and *IL18* (#400301, ACD).

Kidney slices were fixed in 4% formaldehyde embedded in paraffin by the Department of Pathology of Charité-Universitaetsmedizin Berlin. Paraffin-embedded kidney slices were cut into 5 µm sections, plated on Superfrost Plus slides, air-dried overnight, baked for 1 hour at 60°C, cooled for 30 minutes, dewaxed, and air-dried again. Subsequent pretreatment and RNAscope assay procedures for all probes were performed according to the “Formalin-Fixed Paraffin-Embedded (FFPE) Sample Preparation and Pretreatment” and “RNAscope 2.5 HD Detection Reagent BROWN” protocols (ACD documents #322452 and #322310) as recommended by the manufacturer. Sections were counterstained with hematoxylin before dehydrating and applying coverslips using a xylene-based mounting medium. Images of the hybridized sections were captured on a Leica DM2000 LED bright field microscope.

## Acknowledgements

This work was supported by grants to KMSO from the Deutsche Forschungsgemeinschaft (DFG; SFB 1365, GRK 2318 and FOR 2841), to C. H. from the DFG (HI 2238/2-1) and the Berlin Institue of Health (BIH) and MDC (BIH and MDC Focus Area Translational Vascular Biomedicine), and by the Urological Research Foundation (Berlin). ACH was supported by the Berlin University Alliance GC2 Global Health (Corona Virus Pre-Exploration Project), the BMBF (RAPID and Organo-Strat 01KX2021) the DFG (SFB-TR 84, B6 / Z1a). PE was supported by DFG grant EN 924/5-1, the BIH and the Jackstaedt Foundation. NK was supported by DFG grants RA 838/5-1 and KA 5006/1-1. AB was supported by funding from the Gottfried Wilhelm Leibniz Prize of the DFG of NR.

We thank Tatjana Luganskaja for excellent technical support. We thank Benjamin D. Humphreys and his lab for providing supplemental information to their original publication^11^.

## Supplemental Tables

External file SupplTableS1.xlsx

**Supplemental Table S1:** Clinical data and results from histopathological analyses.

External file SupplTableS2.xlsx

**Supplemental Table S2:** Full results from differential gene expression analyses AKI versus control.

External file SupplTableS3.xlsx

**Supplemental Table S3:** Results from pathway enrichment analyses.

External file SupplTableS4.xlsx

**Supplemental Table S4:** Full results from differential gene expression analyses COVID AKI versus non-COVID AKI.

External file SupplTableS5.xlsx

**Supplemental Table S5:** Marker gene comparison between AKI-induced cell states of PT, TAL and DCT.

## Notes

### Competing Interest Statement

The authors have declared no competing interest.

