## Supplementary material for "Transcriptomic responses of the human kidney to acute injury at single cell resolution"

**External file SupplTableS1.xlsx**

**Supplemental Table S1: Clinical data and results from histopathological analyses.**

**External file SupplTableS2.xlsx**

**Supplemental Table S2: Full results from differential gene expression analyses AKI versus control.**

**External file SupplTableS3.xlsx**

**Supplemental Table S3: Results from pathway enrichment analyses.**

**External file SupplTableS4.xlsx**

**Supplemental Table S4: Full results from differential gene expression analyses COVID AKI versus non-COVID AKI.**

**External file SupplTableS5.xlsx**

**Supplemental Table S5: Marker gene comparison between AKI-induced cell states of PT, TAL and DCT.**

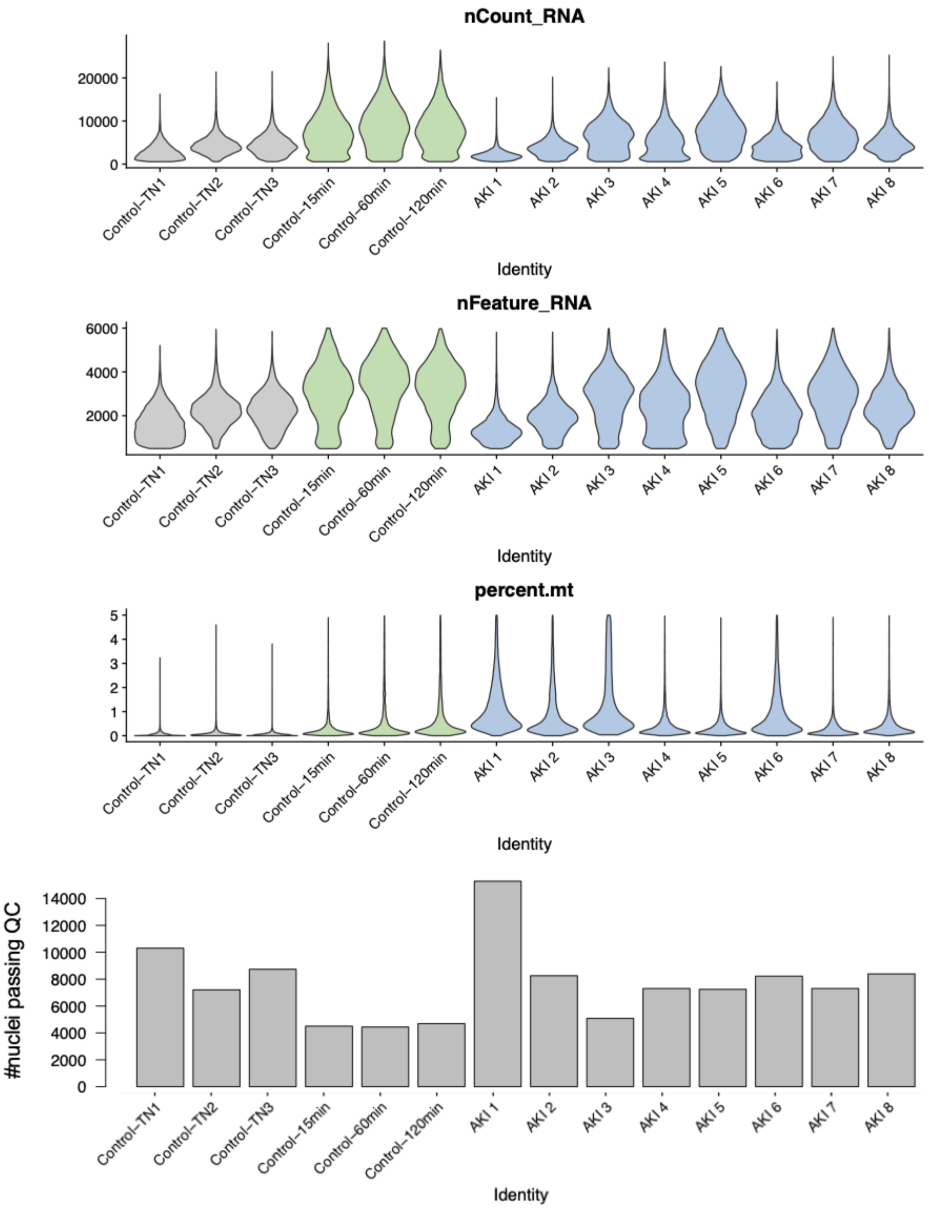


***Supplemental Figure S1: Basic statistics of snRNA-seq libraries.*** *Displayed are plots for number of detected transcripts (nCount_RNA), genes (nFeature_RNA), percent mitochondrial reads (percent.mt) and the number of nuclei which passed quality control per sample.*

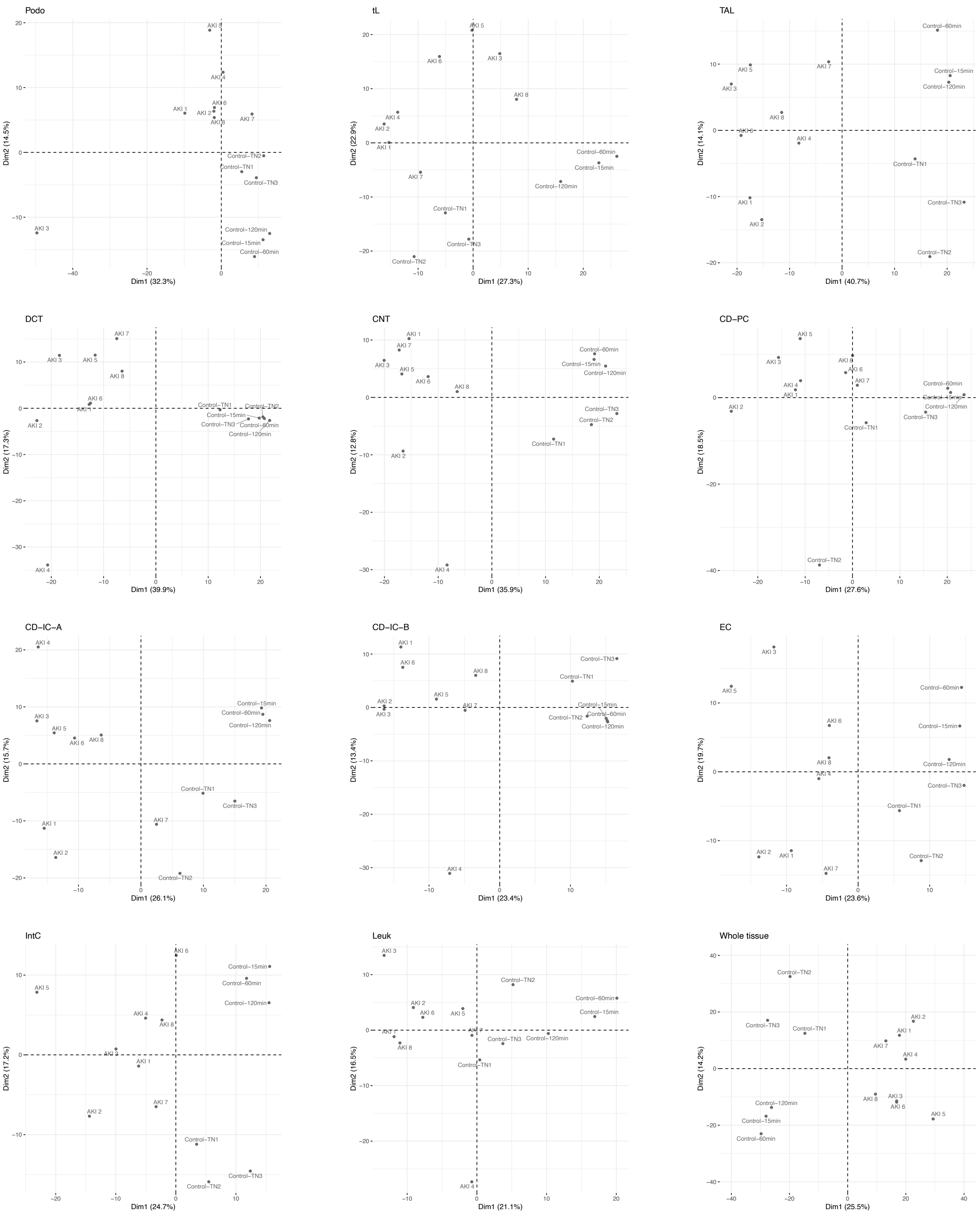


***Supplemental Figure S2:*** *PCA analyses analogous to Fig. 1F using cell type-specific highly variable genes for the remaining cell types and whole tissue.*

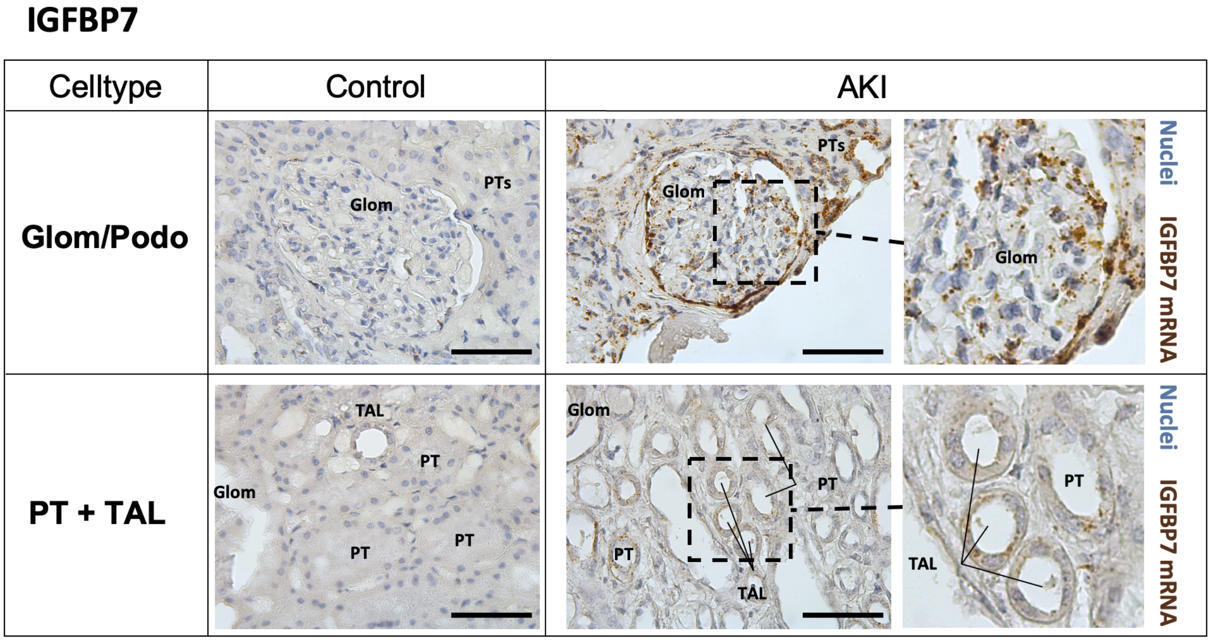


***Supplemental Figure S3:*** *RNAscope in situ hybridizations of injury insulin-like growth factor binding protein 7 (IGFBP7) on post mortem control and AKI sample. Scale bar: 50µm.*

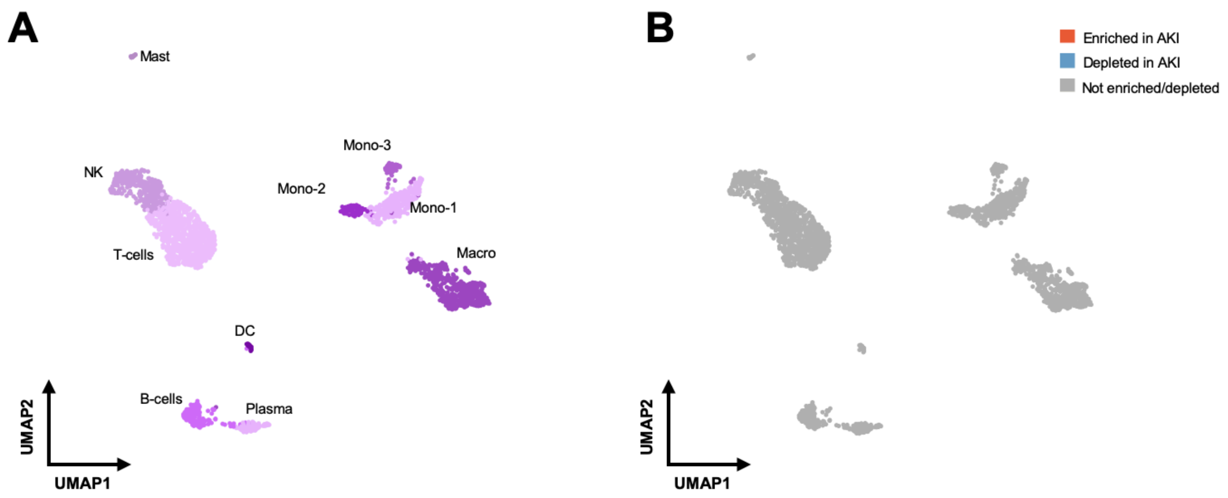


***Supplemental Figure S4: Subclustering of leukocytes. A, B.*** *UMAP plot of subclustered leukocytes (A) and their enrichment or depletion in AKI based on statistical testing of relative abundances within the leukocytes (B) (see methods for details). In B, the same UMAP plot as in A is color-coded based on enrichment (red) or depletion (blue) in AKI individuals. Mast – mast cells, DC – dendritic cells, NK – natural killer cells, Plasma – plasma cells, Mono – monocytes, Macro – macrophages*

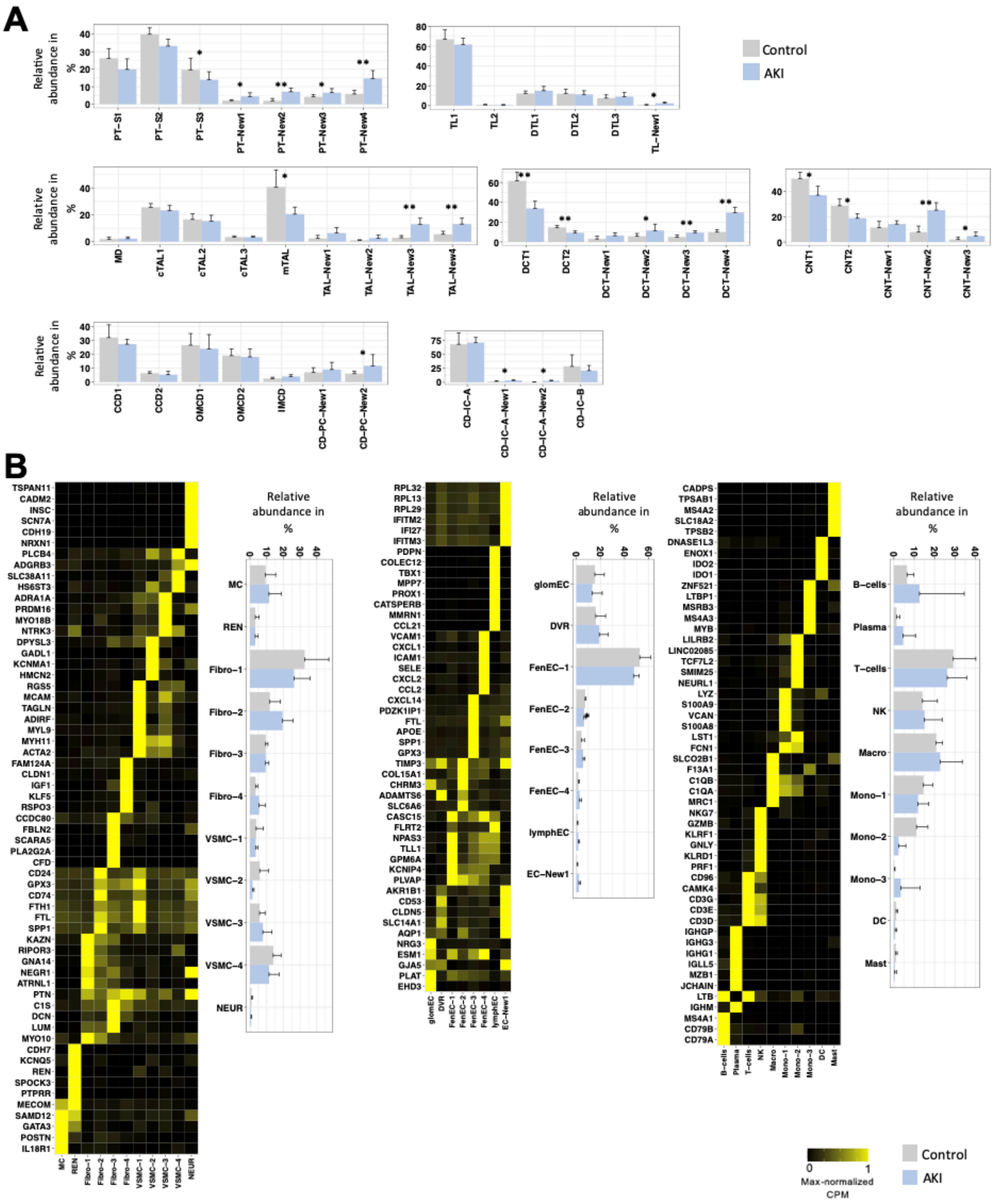


***Supplemental Figure S5: Abundances and marker gene expression of cell populations from subclusterings. A.*** *Relative abundances per major cell type (CD-IC-A and CD-IC-B were analyzed together as CD-IC) of renal cell subpopulations from subclustering analyses of kidney epithelial cells.* ***B.*** *Heatmaps with marker genes and*

*abundance plots as in (A) for ECs, IntCs and leukocytes. Note that marker gene heatmaps for kidney epithelial cell subclusters are shown in figures 4, 5 and Suppl. Fig. S7-9. PT-S1-3 – PT S1-3 segments, c/mTAL – cortical/medullary TAL, TL, DTL – thin limb, descending thin limb, CCD, OMCD, IMCD – cortical/outer and inner medullary collecting duct principal cell; lymphEC – lymphatic EC, GEC – glomerular EC, FenEC 1-4 – fenestrated endothelial cell types, DVR – descending vasa recta; MC – mesangial cells, VSMC – vascular smooth muscle cells, REN – renin-transcribing cells, Fibro – fibroblasts, NEUR – neuronal cells; Mast – mast cells, DC – dendritic cells, NK – natural killer cells, Plasma – plasma cells, Mono – monocytes, Macro – macrophages.*

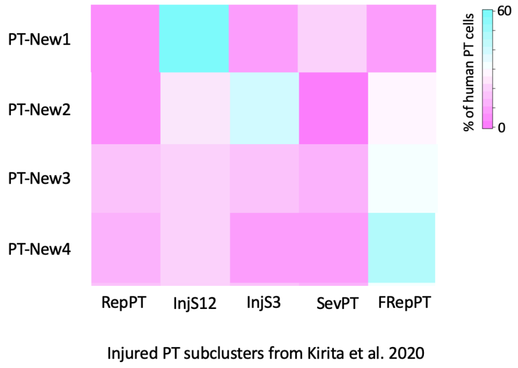


**Supplemental Figure S6:** Results from cross-species approach using published mouse ischemia reperfusion-induced AKI data[1]. The color code indicates the percentage of PT cells from PT-New1-4 in our snRNA-seq data which were assigned to the indicated mouse PT clusters. Abbreviations from the original publication: RepPT – repairing PT, InjS12 – injured S1/2 PT segments, InjS3 – injured S3 segment, SevPT – severely injured PT, FRepPT – failed repair PT.

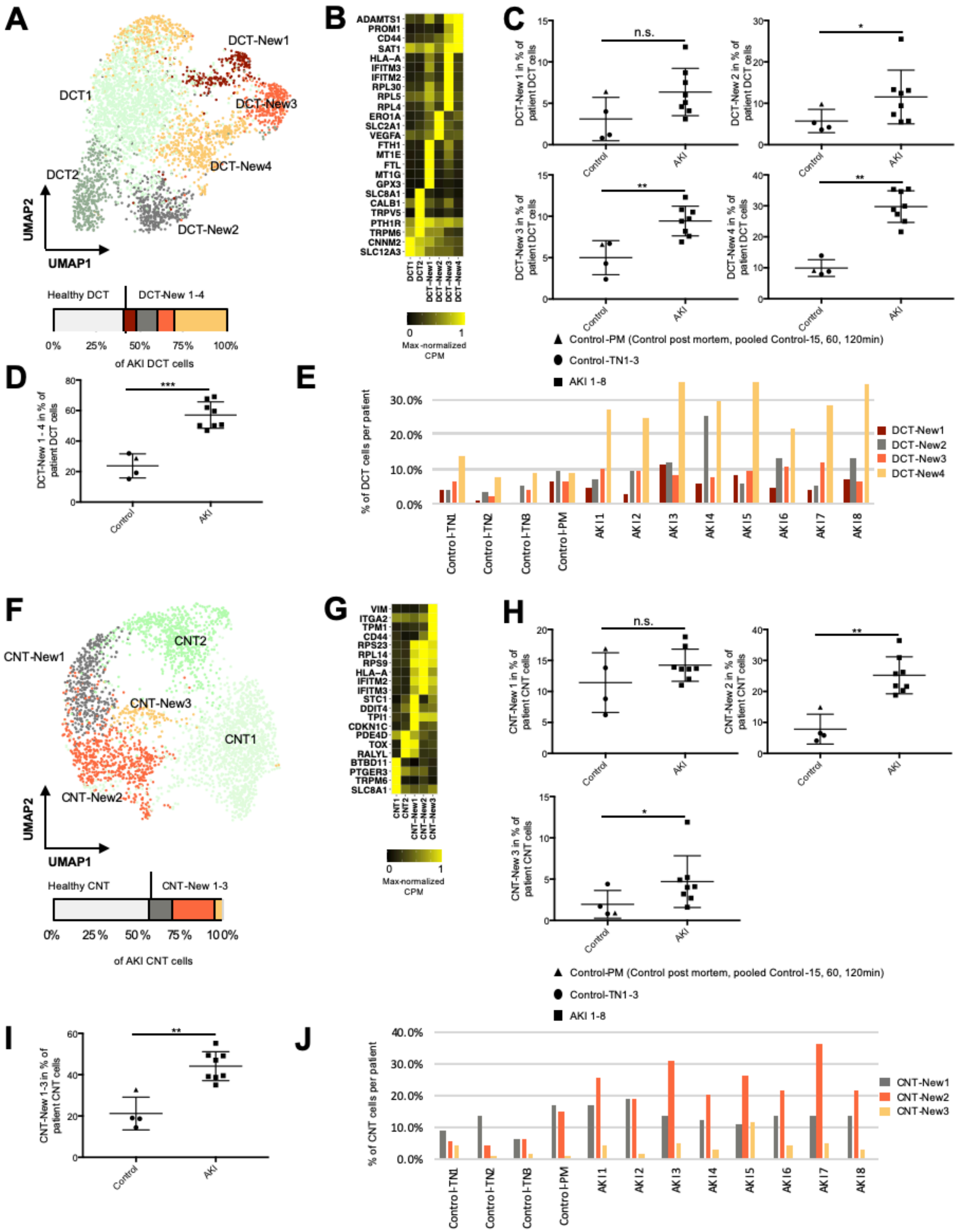


**Supplemental Figure S7:** ***A.*** *UMAP plot of subclustering of the DCT. Below the UMAP is a bar plot displaying the relative abundances of DCT-New 1-4 with respect to all AKI*

*DCT cells.* ***B.*** *Heatmap of selected marker genes for the identified DCT cell subpopulations.* ***C.*** *Plots displaying relative abundances of DCT-New 1-4 with respect to the individual's DCT cells, separately.* ***D.*** *Relative abundances of combined DCT-New 1-4 with respect to the individual's DCT cells.* ***E.*** *Individual abundances of DCT-New 1-4 for control and AKI individuals. P-value: *<0.05, **<0.01, ***<0.001, n.s. – not significant. Control-PM – pooled samples (*Control$_{15min}$, Control$_{60min}$, Control$_{120min}$*) of post mortem non-AKI control individual.*

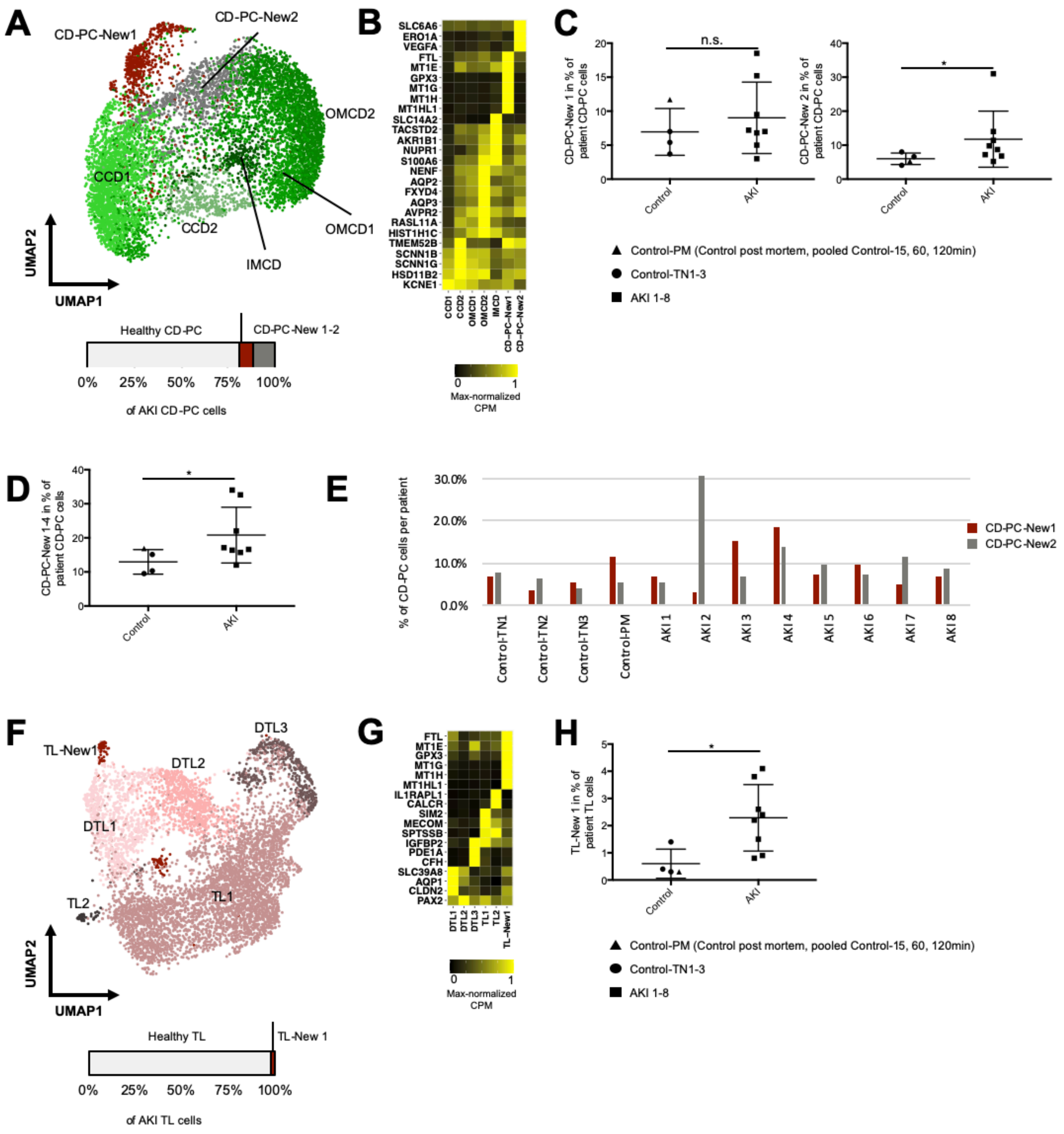


**Supplemental Figure S8:** Analogous figure as shown in Suppl. Fig. S6 for CD-PC (A-E) and thin limbs (F-J). Note that for the thin limbs in F, DTL clusters 1-3, no ascending TL clusters and clusters only named TL1, 2 are depicted. The reason for this is (compare to CD-PC) that we expect very little ascending thin limb cells in our data. The TL clusters TL1 and 2 express SPTSSB, which is an ascending thin limb marker gene, but only little CLCNKA. TL1 and TL2 are very abundant compared to the DTL clusters and are therefore unlikely to be clean ascending thin limb clusters. Cells with characteristics as TL1 and 2 (expression of SPTSSB, little expression of AQP1 and CLCNKA) were described in mouse kidneys before[2].

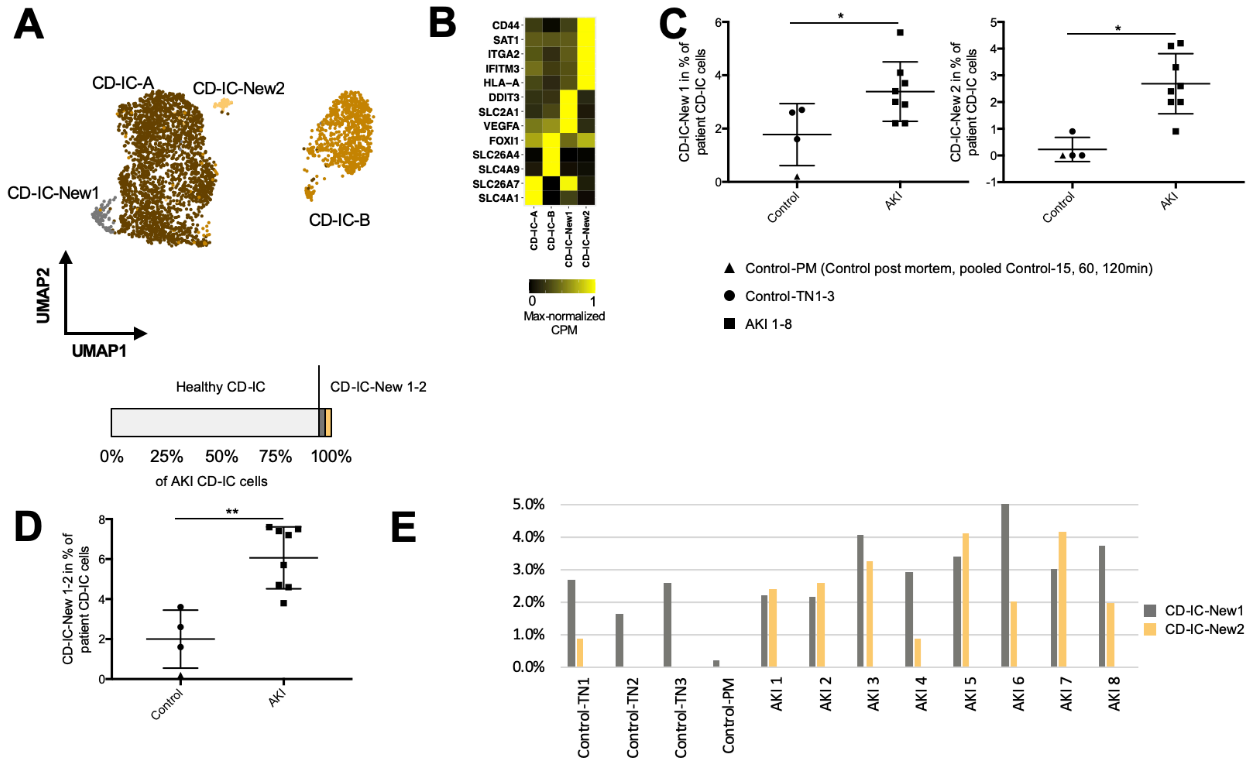


**Supplemental Figure S9:** Analogous figure as shown in Suppl. Fig. S6 for CD-IC.
