## Supplemental Table S1 for "Transcriptomic responses of the human kidney to acute injury at single cell resolution"

| Suppl. Table S1 | Clinical and histopathological data of included individuals | | | | | | | | | | | | |
|---|---|---|---|---|---|---|---|---|---|---|---|---|---|
| | Sample | Control-TN1 | Control-TN2 | Control-TN3 | Control-PM | AKI 1 | AKI 2 | AKI 3 | AKI 4 | AKI 5 | AKI 6 | AKI 7 | AKI 8 |
| | Group | Control | Control | Control | Control | AKI | AKI | AKI | AKI | AKI | AKI | AKI | AKI |
| | Type of kidney sample | Tumor-adjacent normal kidney tissue | Tumor-adjacent normal kidney tissue | Tumor-adjacent normal kidney tissue | Post mortem kidney biopsy | Post mortem kidney biopsy | Post mortem kidney biopsy | Post mortem kidney biopsy | Post mortem kidney biopsy | Post mortem kidney biopsy | Post mortem kidney biopsy | Post mortem kidney biopsy | Post mortem kidney biopsy |
| | Gender (0=m, 1=f) | 1 | 0 | 0 | 0 | 0 | 1 | 1 | 0 | 0 | 0 | 0 | 1 |
| | Age, years | 51 | 61 | 54 | 21 | 57 | 68 | 57 | 53 | 61 | 67 | 72 | 69 |
| | Ethnicity (0=caucasian, 1=african, 2=other) | 0 | 0 | 0 | 0 | 0 | 0 | 0 | 0 | 0 | 0 | 0 | 0 |
| | **Preexisting comorbidities** | | | | | | | | | | | | |
| | Hypertension (0=no, 1=yes) | 1 | 0 | 1 | 0 | 0 | 1 | 1 | 0 | 1 | 1 | 1 | 1 |
| | Diabetes (0=no, 1=yes) | 0 | 0 | 0 | 0 | 0 | 0 | 1 | 0 | 1 | 0 | 0 | 0 |
| | Smoking (0=no, 1=yes) | 0 | 0 | 0 | 0 | 1 | 0 | 0 | 0 | 1 | 0 | 1 | 0 |
| | Coronary heart disease (0=no, 1=yes) | 0 | 0 | 0 | 0 | 0 | 0 | 0 | 0 | 0 | 0 | 1 | 1 |
| | Peripheral vascular disease (0=no, 1=yes) | 0 | 0 | 0 | 0 | 0 | 0 | 0 | 0 | 0 | 1 | 1 | 0 |
| | Stroke/transient ischemic attack (0=no, 1=yes) | 0 | 0 | 0 | 0 | 0 | 0 | 0 | 0 | 1 | 0 | 0 | 0 |
| | Chronic heart failure (0=no, 1=yes) | 0 | 0 | 0 | 0 | 0 | 0 | 1 | 0 | 0 | 0 | 1 | 0 |
| | Atrial fibrillation (0=no, 1=yes) | 0 | 0 | 0 | 0 | 0 | 0 | 0 | 0 | 0 | 1 | 0 | 0 |
| | Chronic obstructive pulmonary disease (0=no, 1=yes) | 0 | 0 | 0 | 0 | 0 | 1 | 0 | 0 | 0 | 0 | 0 | 0 |
| | Asthma (0=no, 1=yes) | 0 | 0 | 0 | 0 | 0 | 0 | 0 | 0 | 0 | 0 | 0 | 0 |
| | Obstructive sleep apnea (0=no, 1=yes) | 0 | 0 | 0 | 0 | 0 | 0 | 0 | 0 | 0 | 0 | 0 | 0 |
| | Active malignancy (0=no, 1=yes) | 1 | 1 | 1 | 0 | 0 | 0 | 0 | 1 | 0 | 0 | 0 | 1 |
| | Obesity (body mass index>30) (0=no, 1=yes) | 0 | 0 | 0 | 1 | 0 | 1 | 0 | 0 | 0 | 0 | 0 | 0 |
| | Days after onset COVID symptoms (applies to COVID-19 patients) | . | . | . | . | 17 | 34 | 27 | 21 | . | . | . | . |
| | Primary cause of death | . | . | . | Cerebral edema | Pneumonia, COVID-19 associated | Pneumonia, COVID-19 associated | Pneumonia, COVID-19 associated | Pneumonia, COVID-19 associated | Pneumonia | Pneumonia | Pneumonia | Pneumonia |
| | AKI stage (KDIGO) at time of tissue sampling/sampling (0 = no AKI, 1 = Stage1 ; 2 =Stage2; 3 = Stage3) | 0 | 0 | 0 | 0 | 3 | 3 | 3 | 3 | 2 | 2 | 3 | 3 |
| | Start Stage 1 (indicate number of days before tissue sampling) | . | . | . | . | 1 | 2 | 2 | 1 | 1 | 2 | 3 | 3 |
| | Start Stage 2 (indicate number of days before tissue sampling) | . | . | . | . | 1 | 2 | 2 | 1 | 1 | 1 | 3 | 3 |
| | Start Stage 3 (indicate number of days before tissue sampling) | . | . | . | . | 1 | 2 | 2 | 1 | | | 3 | 3 |
| | Start of RRT (# days before tissue sampling, 0 if no RRT within 72 hours before tissue sampling) | 0 | 0 | 0 | 0 | 1 | 2 | 1 | 1 | 0 | 0 | 3 | 3 |
| | Need for vasopressor therapy before tissue sampling (# days before tissue sampling, 0= no need for vasopressors within 72 hour | 0 | 0 | 0 | 1 | 1 | 2 | 3 | 2 | 1 | 0 | 2 | 2 |
| | Need for mechanical ventilation (# days before tissue sampling, 0= no need for mechanical ventilation within 72 hours before t | 0 | 0 | 0 | 1 | 1 | 25 | 19 | 2 | 1 | 0 | 3 | 2 |
| | Start of extracorporeal membrane oxygenation (# days before tissue sampling, 0 if no ECMO within 72 hours before tissue samp | 0 | 0 | 0 | 0 | 0 | 2 | 2 | 2 | 0 | 0 | 0 | 0 |
| | Last serum procalcitonin (µg/l) (if available within 96 hours before tissue sampling, otherwise ".") | . | . | . | 0.70 | 0.95 | 67.9 | 1.31 | 76 | 18.7 | 4.88 | 0.32 | 1.45 |
| | Last serum C-reactive protein (mg/dl)(if available within 96 hours before tissue sampling, otherwise ".") | . | . | . | 45.90 | 436 | 299 | 128.1 | 21 | 276 | 102 | 73.8 | 282.6 |
| | Last blood leukocyte count (units)(if available within 96 hours before tissue sampling, otherwise ".") | 6.69 | 8.5 | 7.9 | 18.51 | 15.63 | 31.43 | 9.69 | 8.56 | 1.66 | 12.63 | 10.89 | 10.52 |
| | last serum interleukin-6 level (ng/l)(if available within 96 hours before tissue sampling, otherwise ".") | . | . | . | . | 2684 | 49873 | 2176 | 14655 | 30955 | 142 | 226.1 | 1697 |
| | last serum ferritin (µg/l)(if available within 96 hours before tissue sampling, otherwise ".") | . | . | . | . | 3048 | 5073 | 8936 | 8496 | 612 | 2724 | 433.5 | 46299 |
| | last hematocrit (units)(if available within 96 hours before tissue sampling, otherwise ".") | 0.414 | 0.431 | 0.496 | 0.46 | 0.289 | 0.279 | 0.283 | 0.292 | 0.32 | 0.237 | 0.263 | 0.364 |
| | last hemoglobin (g/dl)(if available within 96 hours before tissue sampling, otherwise ".") | 13.7 | 14.1 | 15.9 | 16.1 | 10 | 7 | 9.2 | 9.9 | 10.2 | 7.8 | 9 | 11.8 |
| | last thrombocyte count (count/nl)(if available within 96 hours before tissue sampling, otherwise ".") | 233 | 329 | 307 | 358.0 | 95 | 375 | 143 | 83 | 66 | 156 | 10.89 | 37 |
| | last neutrophil count (count/nl)(if available within 96 hours before tissue sampling, otherwise ".") | 3.71 | 5.89 | 5.07 | 14.7 | 17.505 | 31.43 | 13.776 | . | . | 9.88 | . | . |
| | last lymphocyte count (count/nl)(if available within 96 hours before tissue sampling, otherwise ".") | 2.11 | 1.47 | 1.74 | 1.7 | 0.91 | 3.1 | 0.68 | . | . | 0.62 | . | 0.147 |
| | Last serum total bilirubin level (mg/dl) (if available within 96 hours before tissue sampling, otherwise ".") | . | . | . | 0.4 | 8 | 1.29 | 2.52 | 0.71 | 1.53 | 17.17 | . | 1.19 |
| | Last serum asparate amino transferase level (U/l) (if available within 96 hours before tissue sampling, otherwise ".") | . | . | . | 55 | 67 | 135 | 10270 | 285 | 61 | 146 | 29 | 141 |
| | Last serum alanin amino transferase level (U/l) (if available within 96 hours before tissue sampling, otherwise ".") | 30 | 26 | 23 | 105 | 39 | 46 | 1566 | 21 | 21 | 85 | 11 | 101 |
| | Last serum lactate dehydrogenase level (U/l) (if available within 96 hours before tissue sampling, otherwise ".") | 183 | 283 | 218 | 294 | 1087 | 702 | 14446 | 1135 | 192 | 357 | 180 | 333 |
| | Last serum D-dimer level (mg/l) (if available within 96 hours before sampling, otherwise ".") | . | . | . | . | >20 | 6.39 | 11.75 | 20 | 1.25 | 10.46 | . | . |
| | **Kidney histology result** | | | | | | | | | | | | |
| | Thrombotic microangiopathy (0=no, 1=yes) | 0 | 0 | 0 | 0 | 1 | 0 | 0 | 0 | 0 | 0 | 0 | 0 |
| | Globally sclerosed glomeruli/total number glomeruli | 0/>30 | 0/>30 | 0/>30 | 0/29 | 0/18 | 0/12 | 2/7 | 0/10 | 1/22 | 2/32 | 23/48 | 3/32 |
| | Acute tubular injury (ATI; tubular dilation, epithelial flattening, cell sloughing, coagulative necrosis; 0 - no, 1 - mild (dilatation, flattening, sloughing), 2 - moderate (focal necrosis), 3 - severe (infarction)) | 0 | 0 | 0 | 0 | 0 | 1 | 1 | 1 | 0 | 2 | 0 | 2 |
| | Tubular vacuolization (0 - absent, 1 - <50%, 2 - >50%) | 0 | 0 | 0 | 0 | 2 | 2 | 0 | 2 | 2 | 0 | 0 | 2 |
