## Supplemental Table S3 for "Transcriptomic responses of the human kidney to acute injury at single cell resolution"

Enrichment for genes upregulated in AKI vs. control

**Note that cell types with no significant enrichment results are not listed.**

Overlap Results

| | |
|---|---|
| Collection(s): | CP:BIOCARTA, CP:KEGG, CP:REACTOME, CP:WIKIPATHWAYS, H |
| # overlaps shown: | 28 |
| # genesets in collections: | 2747 |
| # genes in comparison (n): | 15 |
| # genes in universe (N): | 40312 |

| Gene Set Name | # Genes in Gene Set (K) | Description | # Genes in Overlap (k) | k/K | p-value | FDR q-value |
|---|---|---|---|---|---|---|
| WP_IL17_SIGNALING_PATHWAY | 32 | IL17 signaling pathway | 3 | 0.0938 | 2.05E-07 | 5.61E-04 |
| WP_THE_INFLUENCE_OF_LAMINOPATHIES_ON_WNT_SIGNALING | 40 | The influence of laminopathies on Wnt signaling | 3 | 0.075 | 4.08E-07 | 5.61E-04 |
| HALLMARK_EPITHELIAL_MESENCHYMAL_TRANSITION | 200 | Genes defining epithelial-mesenchymal transition, as in wound healing, fibrosis and metastasis. | 4 | 0.02 | 7.69E-07 | 7.04E-04 |
| WP_THE_OVERLAP_BETWEEN_SIGNAL_TRANSDUCTION_PATHWAYS_THAT_CONTRIBUTE_TO_A_RANGE_OF_LMNA_LAMINOPATHIES | 60 | The Overlap Between Signal Transduction Pathways that Contribute to a Range of LMNA Laminopathies | 3 | 0.05 | 1.41E-06 | 8.97E-04 |
| WP_LUNG_FIBROSIS | 63 | Lung fibrosis | 3 | 0.0476 | 1.63E-06 | 8.97E-04 |
| REACTOME_INTERLEUKIN_4_AND_INTERLEUKIN_13_SIGNALING | 111 | Interleukin-4 and Interleukin-13 signaling | 3 | 0.027 | 9.02E-06 | 3.95E-03 |
| WP_TRANSCRIPTIONAL_CASCADE_REGULATING_ADIPOGENESIS | 13 | Transcriptional cascade regulating adipogenesis | 2 | 0.1538 | 1.01E-05 | 3.95E-03 |
| REACTOME_REGULATION_OF_INSULIN_LIKE_GROWTH_FACTOR_IGF_TRANSPORT_AND_UPTAKE_BY_INSULIN_LIKE_GROWTH_FACTOR_BINDING_PROTEINS_IGFBPS | 124 | Regulation of Insulin-like Growth Factor (IGF) transport and uptake by Insulin-like Growth Factor Binding Proteins (IGFBPs) | 3 | 0.0242 | 1.26E-05 | 4.32E-03 |
| WP_ADIPOGENESIS | 131 | Adipogenesis | 3 | 0.0229 | 1.48E-05 | 4.53E-03 |
| BIOCARTA_IL6_PATHWAY | 21 | IL 6 signaling pathway | 2 | 0.0952 | 2.70E-05 | 7.42E-03 |
| WP_TRANSCRIPTION_FACTOR_REGULATION_IN_ADIPOGENESIS | 22 | Transcription factor regulation in adipogenesis | 2 | 0.0909 | 2.97E-05 | 7.42E-03 |
| WP_EXTRACELLULAR_VESICLES_IN_THE_CROSSTALK_OF_CARDIAC_CELLS | 28 | Extracellular vesicles in the crosstalk of cardiac cells | 2 | 0.0714 | 4.86E-05 | 9.59E-03 |
| HALLMARK_COMPLEMENT | 200 | Genes encoding components of the complement system, which is part of the innate immune system. | 3 | 0.015 | 5.24E-05 | 9.59E-03 |
| HALLMARK_INTERFERON_GAMMA_RESPONSE | 200 | Genes up-regulated in response to IFNG [GeneID=3458]. | 3 | 0.015 | 5.24E-05 | 9.59E-03 |
| HALLMARK_TNFA_SIGNALING_VIA_NFKB | 200 | Genes regulated by NF-kB in response to TNF [GeneID=7124]. | 3 | 0.015 | 5.24E-05 | 9.59E-03 |
| WP_WHITE_FAT_CELL_DIFFERENTIATION | 33 | White fat cell differentiation | 2 | 0.0606 | 6.78E-05 | 1.16E-02 |
| HALLMARK_ANGIOGENESIS | 36 | Genes up-regulated during formation of blood vessels (angiogenesis). | 2 | 0.0556 | 8.08E-05 | 1.31E-02 |
| WP_IL6_SIGNALING_PATHWAY | 43 | IL-6 signaling pathway | 2 | 0.0465 | 1.16E-04 | 1.69E-02 |
| REACTOME_CYTOKINE_SIGNALING_IN_IMMUNE_SYSTEM | 719 | Cytokine Signaling in Immune system | 4 | 0.0056 | 1.17E-04 | 1.69E-02 |
| REACTOME_INTERLEUKIN_10_SIGNALING | 46 | Interleukin-10 signaling | 2 | 0.0435 | 1.32E-04 | 1.82E-02 |
| WP_IL18_SIGNALING_PATHWAY | 279 | IL-18 signaling pathway | 3 | 0.0108 | 1.40E-04 | 1.84E-02 |
| WP_IL4_SIGNALING_PATHWAY | 55 | IL-4 Signaling Pathway | 2 | 0.0364 | 1.90E-04 | 2.37E-02 |
| REACTOME_SIGNALING_BY_PDGF | 58 | Signaling by PDGF | 2 | 0.0345 | 2.11E-04 | 2.42E-02 |
| WP_TGFBETA_RECEPTOR_SIGNALING | 58 | TGF-beta Receptor Signaling | 2 | 0.0345 | 2.11E-04 | 2.42E-02 |
| WP_TGFBETA_RECEPTOR_SIGNALLING_IN_SKELETAL_DYSPLASIAS | 62 | TGF-beta Receptor Signalling in Skeletal Dysplasias | 2 | 0.0323 | 2.41E-04 | 2.65E-02 |
| WP_ONCOSTATIN_M_SIGNALING_PATHWAY | 66 | Oncostatin M Signaling Pathway | 2 | 0.0303 | 2.73E-04 | 2.89E-02 |
| REACTOME_TRANSCRIPTIONAL_REGULATION_OF_WHITE_ADIPOCYTE_DIFFERENTIATION | 84 | Transcriptional regulation of white adipocyte differentiation | 2 | 0.0238 | 4.43E-04 | 4.50E-02 |
| REACTOME_TRANSCRIPTIONAL_REGULATION_OF_GRANULOPOIESIS | 90 | Transcriptional regulation of granulopoiesis | 2 | 0.0222 | 5.08E-04 | 4.98E-02 |

Enrichment for genes upregulated in AKI vs. control

Overlap Results

| | |
|---|---|
| Collection(s): | CP:BIOCARTA, CP:KEGG, CP:REACTOME, CP:WIKIPATHWAYS, H |
| # overlaps shown: | 100 |
| # genesets in collections: | 2747 |
| # genes in comparison (n): | 389 |
| # genes in universe (N): | 40312 |

| Gene Set Name | # Genes in Gene Set (K) | Description | # Genes in Overlap (k) | k/K | p-value | FDR q-value |
|---|---|---|---|---|---|---|
| HALLMARK_TNFA_SIGNALING_VIA_NFKB | 200 | Genes regulated by NF-kB in response to TNF [GeneID=7124]. | 35 | 0.175 | 2.05E-33 | 5.64E-30 |
| HALLMARK_HYPOXIA | 200 | Genes up-regulated in response to low oxygen levels (hypoxia). | 34 | 0.17 | 4.85E-32 | 6.66E-29 |
| HALLMARK_MTORC1_SIGNALING | 200 | Genes up-regulated through activation of mTORC1 complex. | 28 | 0.14 | 3.73E-24 | 3.42E-21 |
| HALLMARK_GLYCOLYSIS | 200 | Genes encoding proteins involved in glycolysis and gluconeogenesis. | 23 | 0.115 | 4.23E-18 | 2.90E-15 |
| WP_PATHWAYS_IN_CLEAR_CELL_RENAL_CELL_CARCINOMA | 87 | Pathways in clear cell renal cell carcinoma | 16 | 0.1839 | 2.64E-16 | 1.45E-13 |
| HALLMARK_INFLAMMATORY_RESPONSE | 200 | Genes defining inflammatory response. | 19 | 0.095 | 1.19E-13 | 5.45E-11 |
| WP_GLYCOLYSIS_AND_GLUCONEOGENESIS | 45 | Glycolysis and Gluconeogenesis | 11 | 0.2444 | 4.44E-13 | 1.74E-10 |
| WP_CORI_CYCLE | 17 | Cori Cycle | 8 | 0.4706 | 1.58E-12 | 5.41E-10 |
| WP_COMPUTATIONAL_MODEL_OF_AEROBIC_GLYCOLYSIS | 12 | Computational Model of Aerobic Glycolysis | 7 | 0.5833 | 5.61E-12 | 1.71E-09 |
| HALLMARK_P53_PATHWAY | 200 | Genes involved in p53 pathways and networks. | 17 | 0.085 | 1.42E-11 | 3.89E-09 |
| REACTOME_CYTOKINE_SIGNALING_IN_IMMUNE_SYSTEM | 719 | Cytokine Signaling in Immune system | 30 | 0.0417 | 2.81E-11 | 7.03E-09 |
| WP_IL18_SIGNALING_PATHWAY | 279 | IL-18 signaling pathway | 19 | 0.0681 | 4.30E-11 | 9.84E-09 |
| REACTOME_GLYCOLYSIS | 72 | Glycolysis | 11 | 0.1528 | 1.05E-10 | 2.21E-08 |
| HALLMARK_EPITHELIAL_MESENCHYMAL_TRANSITION | 200 | Genes defining epithelial-mesenchymal transition, as in wound healing, fibrosis and metastasis. | 16 | 0.08 | 1.40E-10 | 2.57E-08 |
| HALLMARK_INTERFERON_GAMMA_RESPONSE | 200 | Genes up-regulated in response to IFNG [GeneID=3458]. | 16 | 0.08 | 1.40E-10 | 2.57E-08 |
| REACTOME_SIGNALING_BY_INTERLEUKINS | 463 | Signaling by Interleukins | 23 | 0.0497 | 2.19E-10 | 3.75E-08 |
| KEGG_GLYCOLYSIS_GLUCONEOGENESIS | 62 | Glycolysis / Gluconeogenesis | 10 | 0.1613 | 4.29E-10 | 6.94E-08 |
| REACTOME_METABOLISM_OF_CARBOHYDRATES | 293 | Metabolism of carbohydrates | 18 | 0.0614 | 7.40E-10 | 1.13E-07 |
| REACTOME_GLUCOSE_METABOLISM | 92 | Glucose metabolism | 11 | 0.1196 | 1.57E-09 | 2.27E-07 |
| WP_APOPTOSISRELATED_NETWORK_DUE_TO_ALTERED_NOTCH3_IN_OVARIAN_CANCER | 54 | Apoptosis-related network due to altered Notch3 in ovarian cancer | 9 | 0.1667 | 2.40E-09 | 3.29E-07 |
| KEGG_PATHWAYS_IN_CANCER | 325 | Pathways in cancer | 18 | 0.0554 | 3.81E-09 | 4.99E-07 |
| REACTOME_INNATE_IMMUNE_SYSTEM | 1117 | Innate Immune System | 34 | 0.0304 | 5.00E-09 | 6.24E-07 |
| HALLMARK_APOPTOSIS | 161 | Genes mediating programmed cell death (apoptosis) by activation of caspases. | 13 | 0.0807 | 6.79E-09 | 8.10E-07 |
| WP_METABOLIC_REPROGRAMMING_IN_COLON_CANCER | 44 | Metabolic reprogramming in colon cancer | 8 | 0.1818 | 9.16E-09 | 1.05E-06 |
| REACTOME_INTERLEUKIN_4_AND_INTERLEUKIN_13_SIGNALING | 111 | Interleukin-4 and Interleukin-13 signaling | 11 | 0.0991 | 1.17E-08 | 1.29E-06 |
| WP_NUCLEAR_RECEPTORS_METAPATHWAY | 321 | Nuclear Receptors Meta-Pathway | 17 | 0.053 | 2.00E-08 | 2.12E-06 |
| REACTOME_TRANSCRIPTIONAL_REGULATION_BY_TP53 | 363 | Transcriptional Regulation by TP53 | 18 | 0.0496 | 2.11E-08 | 2.14E-06 |
| WP_GLYCOLYSIS_IN_SENESCENCE | 11 | Glycolysis in senescence | 5 | 0.4545 | 3.59E-08 | 3.52E-06 |
| WP_NRF2_PATHWAY | 145 | NRF2 pathway | 11 | 0.0759 | 1.87E-07 | 1.77E-05 |
| WP_PI3KAKT_SIGNALING_PATHWAY | 345 | PI3K-Akt Signaling Pathway | 16 | 0.0464 | 3.17E-07 | 2.90E-05 |
| WP_VEGFAVEGFR2_SIGNALING_PATHWAY | 438 | VEGFA-VEGFR2 Signaling Pathway | 18 | 0.0411 | 3.46E-07 | 3.07E-05 |
| WP_FOCAL_ADHESIONPI3KAKTMTORSIGNALING_PATHWAY | 309 | Focal Adhesion-PI3K-Akt-mTOR-signaling pathway | 15 | 0.0485 | 4.17E-07 | 3.58E-05 |
| WP_SMALL_CELL_LUNG_CANCER | 98 | Small cell lung cancer | 9 | 0.0918 | 4.89E-07 | 4.07E-05 |
| WP_HEPATITIS_C_AND_HEPATOCELLULAR_CARCINOMA | 50 | Hepatitis C and Hepatocellular Carcinoma | 7 | 0.14 | 5.16E-07 | 4.17E-05 |
| HALLMARK_IL2_STAT5_SIGNALING | 199 | Genes up-regulated by STAT5 in response to IL2 stimulation. | 12 | 0.0603 | 6.27E-07 | 4.92E-05 |
| REACTOME_HEMOSTASIS | 678 | Hemostasis | 22 | 0.0324 | 9.76E-07 | 7.45E-05 |
| REACTOME_METABOLISM_OF_LIPIDS | 741 | Metabolism of lipids | 23 | 0.031 | 1.18E-06 | 8.73E-05 |
| KEGG_SMALL_CELL_LUNG_CANCER | 84 | Small cell lung cancer | 8 | 0.0952 | 1.61E-06 | 1.16E-04 |
| REACTOME_EXTRACELLULAR_MATRIX_ORGANIZATION | 301 | Extracellular matrix organization | 14 | 0.0465 | 1.67E-06 | 1.18E-04 |
| WP_SELENIUM_MICRONUTRIENT_NETWORK | 91 | Selenium Micronutrient Network | 8 | 0.0879 | 2.96E-06 | 2.03E-04 |
| KEGG_FOCAL_ADHESION | 199 | Focal adhesion | 11 | 0.0553 | 4.27E-06 | 2.86E-04 |
| HALLMARK_KRAS_SIGNALING_UP | 200 | Genes up-regulated by KRAS activation. | 11 | 0.055 | 4.48E-06 | 2.86E-04 |
| HALLMARK_MYOGENESIS | 200 | Genes involved in development of skeletal muscle (myogenesis). | 11 | 0.055 | 4.48E-06 | 2.86E-04 |
| WP_FOCAL_ADHESION | 202 | Focal Adhesion | 11 | 0.0545 | 4.92E-06 | 3.07E-04 |
| REACTOME_RNA_POLYMERASE_II_TRANSCRIPTION | 1374 | RNA Polymerase II Transcription | 32 | 0.0233 | 5.02E-06 | 3.07E-04 |
| WP_7Q1123_COPY_NUMBER_VARIATION_SYNDROME | 98 | 7q11.23 copy number variation syndrome | 8 | 0.0816 | 5.17E-06 | 3.07E-04 |
| REACTOME_NEUTROPHIL_DEGRANULATION | 479 | Neutrophil degranulation | 17 | 0.0355 | 5.25E-06 | 3.07E-04 |
| REACTOME_DISSOLUTION_OF_FIBRIN_CLOT | 13 | Dissolution of Fibrin Clot | 4 | 0.3077 | 5.70E-06 | 3.19E-04 |
| WP_MAMMARY_GLAND_DEVELOPMENT_PATHWAY_PUBERTY_STAGE_2_OF_4 | 13 | Mammary gland development pathway - Puberty (Stage 2 of 4) | 4 | 0.3077 | 5.70E-06 | 3.19E-04 |
| HALLMARK_ANDROGEN_RESPONSE | 100 | Genes defining response to androgens. | 8 | 0.08 | 6.01E-06 | 3.30E-04 |
| WP_P53_TRANSCRIPTIONAL_GENE_NETWORK | 72 | p53 transcriptional gene network | 7 | 0.0972 | 6.34E-06 | 3.42E-04 |
| HALLMARK_CHOLESTEROL_HOMEOSTASIS | 74 | Genes involved in cholesterol homeostasis. | 7 | 0.0946 | 7.62E-06 | 3.97E-04 |
| HALLMARK_REACTIVE_OXYGEN_SPECIES_PATHWAY | 49 | Genes up-regulated by reactive oxigen species (ROS). | 6 | 0.1224 | 7.65E-06 | 3.97E-04 |
| REACTOME_REPRESSION_OF_WNT_TARGET_GENES | 14 | Repression of WNT target genes | 4 | 0.2857 | 7.92E-06 | 4.03E-04 |
| REACTOME_MET_ACTIVATES_PTK2_SIGNALING | 30 | MET activates PTK2 signaling | 5 | 0.1667 | 9.53E-06 | 4.71E-04 |
| REACTOME_VESICLE_MEDIATED_TRANSPORT | 724 | Vesicle-mediated transport | 21 | 0.029 | 9.61E-06 | 4.71E-04 |
| WP_PHYTOCHEMICAL_ACTIVITY_ON_NRF2_TRANSCRIPTIONAL_ACTIVATION | 15 | Phytochemical activity on NRF2 transcriptional activation | 4 | 0.2667 | 1.07E-05 | 5.15E-04 |
| KEGG_STARCH_AND_SUCROSE_METABOLISM | 52 | Starch and sucrose metabolism | 6 | 0.1154 | 1.09E-05 | 5.15E-04 |
| REACTOME_SIGNALING_BY_MET | 79 | Signaling by MET | 7 | 0.0886 | 1.18E-05 | 5.48E-04 |
| REACTOME_POST_TRANSLATIONAL_PROTEIN_MODIFICATION | 1435 | Post-translational protein modification | 32 | 0.0223 | 1.20E-05 | 5.49E-04 |
| REACTOME_GLYCOGEN_SYNTHESIS | 16 | Glycogen synthesis | 4 | 0.25 | 1.42E-05 | 6.38E-04 |
| REACTOME_MEMBRANE_TRAFFICKING | 629 | Membrane Trafficking | 19 | 0.0302 | 1.46E-05 | 6.47E-04 |
| WP_IL4_SIGNALING_PATHWAY | 55 | IL-4 Signaling Pathway | 6 | 0.1091 | 1.51E-05 | 6.49E-04 |
| WP_RANKLRANK_RECEPTOR_ACTIVATOR_OF_NFKB_LIGAND_SIGNALING_PATHWAY | 55 | RANKL/RANK (Receptor activator of NFKB (ligand)) Signaling Pathway | 6 | 0.1091 | 1.51E-05 | 6.49E-04 |
| WP_DNA_DAMAGE_RESPONSE_ONLY_ATM_DEPENDENT | 114 | DNA Damage Response (only ATM dependent) | 8 | 0.0702 | 1.58E-05 | 6.68E-04 |
| KEGG_FRUCTOSE_AND_MANNOSE_METABOLISM | 34 | Fructose and mannose metabolism | 5 | 0.1471 | 1.80E-05 | 7.39E-04 |
| REACTOME_GLUCONEOGENESIS | 34 | Gluconeogenesis | 5 | 0.1471 | 1.80E-05 | 7.39E-04 |
| BIOCARTA_ETS_PATHWAY | 17 | METS affect on Macrophage Differentiation | 4 | 0.2353 | 1.84E-05 | 7.43E-04 |
| HALLMARK_IL6_JAK_STAT3_SIGNALING | 87 | Genes up-regulated by IL6 [GeneID=3569] via STAT3 [GeneID=6774], e.g., during acute phase response. | 7 | 0.0805 | 2.22E-05 | 8.85E-04 |
| BIOCARTA_CARDIACEGF_PATHWAY | 18 | Role of EGF Receptor Transactivation by GPCRs in Cardiac Hypertrophy | 4 | 0.2222 | 2.35E-05 | 9.21E-04 |
| HALLMARK_HEDGEHOG_SIGNALING | 36 | Genes up-regulated by activation of hedgehog signaling. | 5 | 0.1389 | 2.40E-05 | 9.30E-04 |
| WP_MYOMETRIAL_RELAXATION_AND_CONTRACTION_PATHWAYS | 158 | Myometrial Relaxation and Contraction Pathways | 9 | 0.057 | 2.51E-05 | 9.57E-04 |
| WP_PHOTODYNAMIC_THERAPYINDUCED_HIF1_SURVIVAL_SIGNALING | 37 | Photodynamic therapy-induced HIF-1 survival signaling | 5 | 0.1351 | 2.76E-05 | 1.03E-03 |
| HALLMARK_ESTROGEN_RESPONSE_EARLY | 200 | Genes defining early response to estrogen. | 10 | 0.05 | 2.76E-05 | 1.03E-03 |
| WP_GENOTOXICITY_PATHWAY | 63 | Genotoxicity pathway | 6 | 0.0952 | 3.32E-05 | 1.21E-03 |
| REACTOME_TRANSPORT_OF_SMALL_MOLECULES | 728 | Transport of small molecules | 20 | 0.0275 | 3.36E-05 | 1.21E-03 |
| BIOCARTA_ATM_PATHWAY | 20 | ATM Signaling Pathway | 4 | 0.2 | 3.66E-05 | 1.31E-03 |
| WP_GLYCOGEN_SYNTHESIS_AND_DEGRADATION | 40 | Glycogen Synthesis and Degradation | 5 | 0.125 | 4.06E-05 | 1.43E-03 |
| REACTOME_MET_PROMOTES_CELL_MOTILITY | 41 | MET promotes cell motility | 5 | 0.122 | 4.59E-05 | 1.60E-03 |
| WP_TGFBETA_SIGNALING_PATHWAY | 133 | TGF-beta Signaling Pathway | 8 | 0.0602 | 4.80E-05 | 1.63E-03 |
| WP_HIF1A_AND_PPARG_REGULATION_OF_GLYCOLYSIS | 8 | HIF1A and PPARG regulation of glycolysis | 3 | 0.375 | 4.82E-05 | 1.63E-03 |
| WP_TNF_RELATED_WEAK_INDUCER_OF_APOPTOSIS_TWEAK_SIGNALING_PATHWAY | 42 | TNF related weak inducer of apoptosis (TWEAK) Signaling Pathway | 5 | 0.119 | 5.17E-05 | 1.73E-03 |
| WP_PI3KAKTMTOR_VITD3_SIGNALLING | 22 | PI3K/AKT/mTOR - VitD3 Signalling | 4 | 0.1818 | 5.44E-05 | 1.80E-03 |
| WP_DNA_DAMAGE_RESPONSE | 69 | DNA Damage Response | 6 | 0.087 | 5.58E-05 | 1.82E-03 |
| WP_CILIARY_LANDSCAPE | 219 | Ciliary landscape | 10 | 0.0457 | 5.95E-05 | 1.92E-03 |
| KEGG_RENAL_CELL_CARCINOMA | 70 | Renal cell carcinoma | 6 | 0.0857 | 6.05E-05 | 1.93E-03 |
| REACTOME_NUCLEOTIDE_SALVAGE | 23 | Nucleotide salvage | 4 | 0.1739 | 6.54E-05 | 2.02E-03 |
| WP_NRF2ARE_REGULATION | 23 | NRF2-ARE regulation | 4 | 0.1739 | 6.54E-05 | 2.02E-03 |
| WP_PHOTODYNAMIC_THERAPYINDUCED_NFE2L2_NRF2_SURVIVAL_SIGNALING | 23 | Photodynamic therapy-induced NFE2L2 (NRF2) survival signaling | 4 | 0.1739 | 6.54E-05 | 2.02E-03 |
| REACTOME_CELLULAR_RESPONSES_TO_EXTERNAL_STIMULI | 706 | Cellular responses to external stimuli | 19 | 0.0269 | 6.90E-05 | 2.08E-03 |
| REACTOME_INTERLEUKIN_1_FAMILY_SIGNALING | 140 | Interleukin-1 family signaling | 8 | 0.0571 | 6.90E-05 | 2.08E-03 |
| WP_PROSTAGLANDIN_SYNTHESIS_AND_REGULATION | 45 | Prostaglandin Synthesis and Regulation | 5 | 0.1111 | 7.25E-05 | 2.17E-03 |
| WP_FOLATE_METABOLISM | 73 | Folate Metabolism | 6 | 0.0822 | 7.67E-05 | 2.27E-03 |
| KEGG_DORSO_VENTRAL_AXIS_FORMATION | 24 | Dorso-ventral axis formation | 4 | 0.1667 | 7.79E-05 | 2.28E-03 |
| REACTOME_INTERLEUKIN_10_SIGNALING | 46 | Interleukin-10 signaling | 5 | 0.1087 | 8.07E-05 | 2.33E-03 |
| HALLMARK_UV_RESPONSE_DN | 144 | Genes down-regulated in response to ultraviolet (UV) radiation. | 8 | 0.0556 | 8.42E-05 | 2.41E-03 |
| WP_RAS_SIGNALING | 185 | Ras Signaling | 9 | 0.0486 | 8.55E-05 | 2.42E-03 |

PT UP

| | | | | | | |
|---|---|---|---|---|---|---|
| WP_UNFOLDED_PROTEIN_RESPONSE | 25 | Unfolded protein response | 4 | 0.16 | 9.20E-05 | 2.57E-03 |
| REACTOME_MAPK_FAMILY_SIGNALING_CASCADES | 327 | MAPK family signaling cascades | 12 | 0.0367 | 9.25E-05 | 2.57E-03 |
| WP_ARYL_HYDROCARBON_RECEPTOR_NETPATH | 48 | Aryl Hydrocarbon Receptor Netpath | 5 | 0.1042 | 9.93E-05 | 2.73E-03 |

Enrichment for genes downregulated in AKI vs. control

Overlap Results

| | |
|---|---|
| Collection(s): | CP:BIOCARTA, CP:KEGG, CP:REACTOME, CP:WIKIPATHWAYS, H |
| # overlaps shown: | 33 |
| # genesets in collections: | 2747 |
| # genes in comparison (n): | 271 |
| # genes in universe (N): | 40312 |

| Gene Set Name | # Genes in Gene Set (K) | Description | # Genes in Overlap (k) | k/K | p-value | FDR q-value |
|---|---|---|---|---|---|---|
| REACTOME_SLC_MEDIATED_TRANSMEMBRANE_TRANSPORT | 250 | SLC-mediated transmembrane transport | 18 | 0.072 | 1.35E-13 | 3.70E-10 |
| REACTOME_TRANSPORT_OF_INORGANIC_CATIONS_ANIONS_AND_AMINO_ACIDS_OLIGOPEPTIDES | 106 | Transport of inorganic cations/anions and amino acids/oligopeptides | 12 | 0.1132 | 8.39E-12 | 1.15E-08 |
| REACTOME_METABOLISM_OF_AMINO_ACIDS_AND_DERIVATIVES | 374 | Metabolism of amino acids and derivatives | 19 | 0.0508 | 1.32E-11 | 1.21E-08 |
| REACTOME_TRANSPORT_OF_SMALL_MOLECULES | 728 | Transport of small molecules | 23 | 0.0316 | 1.16E-09 | 7.95E-07 |
| REACTOME_SLC_TRANSPORTER_DISORDERS | 99 | SLC transporter disorders | 10 | 0.101 | 1.47E-09 | 8.08E-07 |
| KEGG_ARGININE_AND_PROLINE_METABOLISM | 54 | Arginine and proline metabolism | 8 | 0.1481 | 3.00E-09 | 1.37E-06 |
| REACTOME_DISORDERS_OF_TRANSMEMBRANE_TRANSPORTERS | 176 | Disorders of transmembrane transporters | 11 | 0.0625 | 3.56E-08 | 1.40E-05 |
| WP_PROXIMAL_TUBULE_TRANSPORT | 58 | Proximal tubule transport | 7 | 0.1207 | 1.29E-07 | 4.42E-05 |
| KEGG_GLYCINE_SERINE_AND_THREONINE_METABOLISM | 31 | Glycine, serine and threonine metabolism | 5 | 0.1613 | 1.95E-06 | 5.95E-04 |
| REACTOME_CHOLINE_CATABOLISM | 6 | Choline catabolism | 3 | 0.5 | 5.92E-06 | 1.63E-03 |
| KEGG_TRYPTOPHAN_METABOLISM | 40 | Tryptophan metabolism | 5 | 0.125 | 7.18E-06 | 1.79E-03 |
| HALLMARK_XENOBIOTIC_METABOLISM | 200 | Genes encoding proteins involved in processing of drugs and other xenobiotics. | 9 | 0.045 | 9.45E-06 | 2.14E-03 |
| REACTOME_CATION_COUPLED_CHLORIDE_COTRANSPORTERS | 7 | Cation-coupled Chloride cotransporters | 3 | 0.4286 | 1.03E-05 | 2.14E-03 |
| WP_UREA_CYCLE_AND_METABOLISM_OF_AMINO_GROUPS | 21 | Urea cycle and metabolism of amino groups | 4 | 0.1905 | 1.09E-05 | 2.14E-03 |
| WP_FGF23_SIGNALLING_IN_HYPOPHOSPHATEMIC_RICKETS_AND_RELATED_DISORDERS | 22 | FGF23 signalling in Hypophosphatemic rickets and related disorders | 4 | 0.1818 | 1.33E-05 | 2.43E-03 |
| KEGG_GLYCEROLIPID_METABOLISM | 49 | Glycerolipid metabolism | 5 | 0.102 | 1.98E-05 | 3.40E-03 |
| REACTOME_METABOLISM_OF_VITAMINS_AND_COFACTORS | 189 | Metabolism of vitamins and cofactors | 8 | 0.0423 | 4.58E-05 | 7.41E-03 |
| REACTOME_GLYOXYLATE_METABOLISM_AND_GLYCINE_DEGRADATION | 31 | Glyoxylate metabolism and glycine degradation | 4 | 0.129 | 5.45E-05 | 8.31E-03 |
| REACTOME_AMINO_ACID_TRANSPORT_ACROSS_THE_PLASMA_MEMBRANE | 33 | Amino acid transport across the plasma membrane | 4 | 0.1212 | 7.01E-05 | 1.01E-02 |
| WP_GENES_INVOLVED_IN_MALE_INFERTILITY | 155 | Genes involved in male infertility | 7 | 0.0452 | 9.16E-05 | 1.26E-02 |
| REACTOME_TRYPTOPHAN_CATABOLISM | 14 | Tryptophan catabolism | 3 | 0.2143 | 1.04E-04 | 1.35E-02 |
| REACTOME_METABOLISM_OF_LIPIDS | 741 | Metabolism of lipids | 15 | 0.0202 | 1.70E-04 | 2.08E-02 |
| REACTOME_SENSORY_PROCESSING_OF_SOUND | 77 | Sensory processing of sound | 5 | 0.0649 | 1.76E-04 | 2.08E-02 |
| REACTOME_METABOLISM_OF_CARBOHYDRATES | 293 | Metabolism of carbohydrates | 9 | 0.0307 | 1.81E-04 | 2.08E-02 |
| REACTOME_AFLATOXIN_ACTIVATION_AND_DETOXIFICATION | 19 | Aflatoxin activation and detoxification | 3 | 0.1579 | 2.69E-04 | 2.84E-02 |
| REACTOME_METABOLISM_OF_COFACTORS | 19 | Metabolism of cofactors | 3 | 0.1579 | 2.69E-04 | 2.84E-02 |
| REACTOME_CELLULAR_HEXOSE_TRANSPORT | 21 | Cellular hexose transport | 3 | 0.1429 | 3.65E-04 | 3.72E-02 |
| WP_AMINO_ACID_METABOLISM | 91 | Amino Acid metabolism | 5 | 0.0549 | 3.84E-04 | 3.77E-02 |
| REACTOME_FRUCTOSE_CATABOLISM | 5 | Fructose catabolism | 2 | 0.4 | 4.44E-04 | 4.07E-02 |
| REACTOME_SODIUM_COUPLED_PHOSPHATE_COTRANSPORTERS | 5 | Sodium-coupled phosphate cotransporters | 2 | 0.4 | 4.44E-04 | 4.07E-02 |
| REACTOME_THE_CANONICAL_RETINOID_CYCLE_IN_RODS_TWILIGHT_VISION | 23 | The canonical retinoid cycle in rods (twilight vision) | 3 | 0.1304 | 4.82E-04 | 4.13E-02 |
| WP_KYNURENINE_PATHWAY_AND_LINKS_TO_CELLULAR_SENESCENCE | 23 | Kynurenine Pathway and links to Cellular Senescence | 3 | 0.1304 | 4.82E-04 | 4.13E-02 |
| REACTOME_SENSORY_PROCESSING_OF_SOUND_BY_OUTER_HAIR_CELLS_OF_THE_COCHLEA | 55 | Sensory processing of sound by outer hair cells of the cochlea | 4 | 0.0727 | 5.20E-04 | 4.33E-02 |

Enrichment for genes upregulated in AKI vs. control

Overlap Results

| | |
|---|---|
| Collection(s): | CP:BIOCARTA, CP:KEGG, CP:REACTOME, CP:WIKIPATHWAYS, H |
| # overlaps shown: | 43 |
| # genesets in collections: | 2747 |
| # genes in comparison (n): | 51 |
| # genes in universe (N): | 40312 |

| Gene Set Name | # Genes in Gene Set (K) | Description | # Genes in Overlap (k) | k/K | p-value | FDR q-value |
|---|---|---|---|---|---|---|
| WP_COMPLEMENT_AND_COAGULATION_CASCADES | 59 | Complement and Coagulation Cascades | 5 | 0.0847 | 1.26E-08 | 3.46E-05 |
| KEGG_COMPLEMENT_AND_COAGULATION_CASCADES | 69 | Complement and coagulation cascades | 5 | 0.0725 | 2.80E-08 | 3.85E-05 |
| REACTOME_INNATE_IMMUNE_SYSTEM | 1117 | Innate Immune System | 11 | 0.0098 | 1.22E-07 | 1.11E-04 |
| REACTOME_FORMATION_OF_FIBRIN_CLOT_CLOTTING_CASCADE | 39 | Formation of Fibrin Clot (Clotting Cascade) | 4 | 0.1026 | 1.81E-07 | 1.24E-04 |
| WP_PHYTOCHEMICAL_ACTIVITY_ON_NRF2_TRANSCRIPTIONAL_ACTIVATION | 15 | Phytochemical activity on NRF2 transcriptional activation | 3 | 0.2 | 8.59E-07 | 4.72E-04 |
| WP_NRF2ARE_REGULATION | 23 | NRF2-ARE regulation | 3 | 0.1304 | 3.32E-06 | 1.30E-03 |
| WP_PHOTODYNAMIC_THERAPYINDUCED_NFE2L2_NRF2_SURVIVAL_SIGNALING | 23 | Photodynamic therapy-induced NFE2L2 (NRF2) survival signaling | 3 | 0.1304 | 3.32E-06 | 1.30E-03 |
| KEGG_NICOTINATE_AND_NICOTINAMIDE_METABOLISM | 24 | Nicotinate and nicotinamide metabolism | 3 | 0.125 | 3.79E-06 | 1.30E-03 |
| HALLMARK_EPITHELIAL_MESENCHYMAL_TRANSITION | 200 | Genes defining epithelial-mesenchymal transition, as in wound healing, fibrosis and metastasis. | 5 | 0.025 | 5.58E-06 | 1.70E-03 |
| WP_HUMAN_COMPLEMENT_SYSTEM | 99 | Human Complement System | 4 | 0.0404 | 7.83E-06 | 2.15E-03 |
| REACTOME_METABOLIC_DISORDERS_OF_BIOLOGICAL_OXIDATION_ENZYMES | 34 | Metabolic disorders of biological oxidation enzymes | 3 | 0.0882 | 1.11E-05 | 2.77E-03 |
| REACTOME_DISEASES_OF_METABOLISM | 246 | Diseases of metabolism | 5 | 0.0203 | 1.52E-05 | 3.47E-03 |
| WP_FERROPTOSIS | 40 | Ferroptosis | 3 | 0.075 | 1.82E-05 | 3.85E-03 |
| HALLMARK_COAGULATION | 138 | Genes encoding components of blood coagulation system; also up-regulated in platelets. | 4 | 0.029 | 2.90E-05 | 5.69E-03 |
| REACTOME_NEUTROPHIL_DEGRANULATION | 479 | Neutrophil degranulation | 6 | 0.0125 | 3.12E-05 | 5.69E-03 |
| HALLMARK_REACTIVE_OXYGEN_SPECIES_PATHWAY | 49 | Genes up-regulated by reactive oxigen species (ROS). | 3 | 0.0612 | 3.37E-05 | 5.69E-03 |
| WP_NRF2_PATHWAY | 145 | NRF2 pathway | 4 | 0.0276 | 3.52E-05 | 5.69E-03 |
| WP_APOPTOSISRELATED_NETWORK_DUE_TO_ALTERED_NOTCH3_IN_OVARIAN_CANCER | 54 | Apoptosis-related network due to altered Notch3 in ovarian cancer | 3 | 0.0556 | 4.52E-05 | 6.90E-03 |
| WP_NUCLEAR_RECEPTORS_METAPATHWAY | 321 | Nuclear Receptors Meta-Pathway | 5 | 0.0156 | 5.40E-05 | 7.81E-03 |
| WP_LUNG_FIBROSIS | 63 | Lung fibrosis | 3 | 0.0476 | 7.18E-05 | 9.86E-03 |
| WP_METAPATHWAY_BIOTRANSFORMATION_PHASE_I_AND_II | 185 | Metapathway biotransformation Phase I and II | 4 | 0.0216 | 9.07E-05 | 1.19E-02 |
| REACTOME_GLUTATHIONE_SYNTHESIS_AND_RECYCLING | 12 | Glutathione synthesis and recycling | 2 | 0.1667 | 1.03E-04 | 1.28E-02 |
| BIOCARTA_EXTRINSIC_PATHWAY | 13 | Extrinsic Prothrombin Activation Pathway | 2 | 0.1538 | 1.21E-04 | 1.35E-02 |
| HALLMARK_ESTROGEN_RESPONSE_EARLY | 200 | Genes defining early response to estrogen. | 4 | 0.02 | 1.22E-04 | 1.35E-02 |
| HALLMARK_INTERFERON_GAMMA_RESPONSE | 200 | Genes up-regulated in response to IFNG [GeneID=3458]. | 4 | 0.02 | 1.22E-04 | 1.35E-02 |
| REACTOME_BIOLOGICAL_OXIDATIONS | 222 | Biological oxidations | 4 | 0.018 | 1.83E-04 | 1.93E-02 |
| REACTOME_HEMOSTASIS | 678 | Hemostasis | 6 | 0.0088 | 2.10E-04 | 2.13E-02 |
| REACTOME_NICOTINAMIDE_SALVAGING | 19 | Nicotinamide salvaging | 2 | 0.1053 | 2.65E-04 | 2.60E-02 |
| BIOCARTA_AMI_PATHWAY | 20 | Acute Myocardial Infarction | 2 | 0.1 | 2.94E-04 | 2.78E-02 |
| REACTOME_COMMON_PATHWAY_OF_FIBRIN_CLOT_FORMATION | 22 | Common Pathway of Fibrin Clot Formation | 2 | 0.0909 | 3.57E-04 | 2.98E-02 |
| WP_FGF23_SIGNALLING_IN_HYPOPHOSPHATEMIC_RICKETS_AND_RELATED_DISORDERS | 22 | FGF23 signalling in Hypophosphatemic rickets and related disorders | 2 | 0.0909 | 3.57E-04 | 2.98E-02 |
| WP_GLUTATHIONE_METABOLISM | 22 | Glutathione metabolism | 2 | 0.0909 | 3.57E-04 | 2.98E-02 |
| REACTOME_PHASE_II_CONJUGATION_OF_COMPOUNDS | 109 | Phase II - Conjugation of compounds | 3 | 0.0275 | 3.64E-04 | 2.98E-02 |
| REACTOME_INTERLEUKIN_4_AND_INTERLEUKIN_13_SIGNALING | 111 | Interleukin-4 and Interleukin-13 signaling | 3 | 0.027 | 3.84E-04 | 2.98E-02 |
| BIOCARTA_INTRINSIC_PATHWAY | 23 | Intrinsic Prothrombin Activation Pathway | 2 | 0.087 | 3.90E-04 | 2.98E-02 |
| REACTOME_INTRINSIC_PATHWAY_OF_FIBRIN_CLOT_FORMATION | 23 | Intrinsic Pathway of Fibrin Clot Formation | 2 | 0.087 | 3.90E-04 | 2.98E-02 |
| WP_IL18_SIGNALING_PATHWAY | 279 | IL-18 signaling pathway | 4 | 0.0143 | 4.34E-04 | 3.23E-02 |
| REACTOME_PURINERGIC_SIGNALING_IN_LEISHMANIASIS_INFECTION | 26 | Purinergic signaling in leishmaniasis infection | 2 | 0.0769 | 5.00E-04 | 3.62E-02 |
| REACTOME_REGULATION_OF_INSULIN_LIKE_GROWTH_FACTOR_IGF_TRANSPORT_AND_UPTAKE_BY_INSULIN_LIKE_GROWTH_FACTOR_BINDING_PROTEINS_IGFBPS | 124 | Regulation of Insulin-like Growth Factor (IGF) transport and uptake by Insulin-like Growth Factor Binding Proteins (IGFBPs) | 3 | 0.0242 | 5.31E-04 | 3.74E-02 |
| WP_RAS_AND_BRADYKININ_PATHWAYS_IN_COVID19 | 29 | RAS and Bradykinin pathways in COVID-19 | 2 | 0.069 | 6.23E-04 | 4.27E-02 |
| REACTOME_RESPONSE_TO_ELEVATED_PLATELET_CYTOSOLIC_CA2 | 132 | Response to elevated platelet cytosolic Ca2+ | 3 | 0.0227 | 6.37E-04 | 4.27E-02 |
| REACTOME_NICOTINATE_METABOLISM | 31 | Nicotinate metabolism | 2 | 0.0645 | 7.13E-04 | 4.66E-02 |
| WP_TRANSSULFURATION_AND_ONE_CARBON_METABOLISM | 32 | Trans-sulfuration and one carbon metabolism | 2 | 0.0625 | 7.60E-04 | 4.85E-02 |

Enrichment for genes downregulated in AKI vs. control

Overlap Results

| | |
|---|---|
| Collection(s): | CP:BIOCARTA, CP:KEGG, CP:REACTOME, CP:WIKIPATHWAYS, H |
| # overlaps shown: | 20 |
| # genesets in collections: | 2747 |
| # genes in comparison (n): | 34 |
| # genes in universe (N): | 40312 |

| Gene Set Name | # Genes in Gene Set (K) | Description | # Genes in Overlap (k) | k/K | p-value | FDR q-value |
|---|---|---|---|---|---|---|
| BIOCARTA_NPP1_PATHWAY | 10 | Regulators of Bone Mineralization | 3 | 0.3 | 6.55E-08 | 1.80E-04 |
| REACTOME_EXTRACELLULAR_MATRIX_ORGANIZATION | 301 | Extracellular matrix organization | 5 | 0.0166 | 5.23E-06 | 4.87E-03 |
| REACTOME_DEGRADATION_OF_THE_EXTRACELLULAR_MATRIX | 140 | Degradation of the extracellular matrix | 4 | 0.0286 | 5.96E-06 | 4.87E-03 |
| REACTOME_COLLAGEN_CHAIN_TRIMERIZATION | 44 | Collagen chain trimerization | 3 | 0.0682 | 7.09E-06 | 4.87E-03 |
| REACTOME_NON_INTEGRIN_MEMBRANE_ECM_INTERACTIONS | 59 | Non-integrin membrane-ECM interactions | 3 | 0.0508 | 1.73E-05 | 7.72E-03 |
| REACTOME_ASSEMBLY_OF_COLLAGEN_FIBRILS_AND_OTHER_MULTIMERIC_STRUCTURES | 61 | Assembly of collagen fibrils and other multimeric structures | 3 | 0.0492 | 1.91E-05 | 7.72E-03 |
| REACTOME_COLLAGEN_DEGRADATION | 64 | Collagen degradation | 3 | 0.0469 | 2.20E-05 | 7.72E-03 |
| HALLMARK_EPITHELIAL_MESENCHYMAL_TRANSITION | 200 | Genes defining epithelial-mesenchymal transition, as in wound healing, fibrosis and metastasis. | 4 | 0.02 | 2.43E-05 | 7.72E-03 |
| REACTOME_COLLAGEN_BIOSYNTHESIS_AND_MODIFYING_ENZYMES | 67 | Collagen biosynthesis and modifying enzymes | 3 | 0.0448 | 2.53E-05 | 7.72E-03 |
| BIOCARTA_VITCB_PATHWAY | 11 | Vitamin C in the Brain | 2 | 0.1818 | 3.78E-05 | 1.04E-02 |
| BIOCARTA_ACE2_PATHWAY | 13 | Angiotensin-converting enzyme 2 regulates heart function | 2 | 0.1538 | 5.35E-05 | 1.32E-02 |
| REACTOME_COLLAGEN_FORMATION | 90 | Collagen formation | 3 | 0.0333 | 6.12E-05 | 1.32E-02 |
| BIOCARTA_PLATELETAPP_PATHWAY | 14 | Platelet Amyloid Precursor Protein Pathway | 2 | 0.1429 | 6.24E-05 | 1.32E-02 |
| REACTOME_ANCHORING_FIBRIL_FORMATION | 15 | Anchoring fibril formation | 2 | 0.1333 | 7.20E-05 | 1.41E-02 |
| WP_SMALL_CELL_LUNG_CANCER | 98 | Small cell lung cancer | 3 | 0.0306 | 7.89E-05 | 1.45E-02 |
| REACTOME_CROSSLINKING_OF_COLLAGEN_FIBRILS | 18 | Crosslinking of collagen fibrils | 2 | 0.1111 | 1.05E-04 | 1.80E-02 |
| WP_OVERVIEW_OF_NANOPARTICLE_EFFECTS | 19 | Overview of nanoparticle effects | 2 | 0.1053 | 1.17E-04 | 1.89E-02 |
| BIOCARTA_AMI_PATHWAY | 20 | Acute Myocardial Infarction | 2 | 0.1 | 1.30E-04 | 1.98E-02 |
| BIOCARTA_INTRINSIC_PATHWAY | 23 | Intrinsic Prothrombin Activation Pathway | 2 | 0.087 | 1.73E-04 | 2.50E-02 |
| REACTOME_LAMININ_INTERACTIONS | 30 | Laminin interactions | 2 | 0.0667 | 2.96E-04 | 4.06E-02 |

TAL UP

Enrichment for genes upregulated in AKI vs. control

Overlap Results

| | |
|---|---|
| Collection(s): | CP:BIOCARTA, CP:KEGG, CP:REACTOME, CP:WIKIPATHWAYS, H |
| # overlaps shown: | 100 |
| # genesets in collections: | 2747 |
| # genes in comparison (n): | 420 |
| # genes in universe (N): | 40312 |

| Gene Set Name | # Genes in Gene Set (K) | Description | # Genes in Overlap (k) | k/K | p-value | FDR q-value |
|---|---|---|---|---|---|---|
| HALLMARK_TNFA_SIGNALING_VIA_NFKB | 200 | Genes regulated by NF-kB in response to TNF [GeneID=7124]. | 32 | 0.16 | 2.74E-28 | 7.53E-25 |
| REACTOME_INNATE_IMMUNE_SYSTEM | 1117 | Innate Immune System | 56 | 0.0501 | 2.62E-22 | 3.60E-19 |
| REACTOME_CYTOKINE_SIGNALING_IN_IMMUNE_SYSTEM | 719 | Cytokine Signaling in Immune system | 41 | 0.057 | 1.68E-18 | 1.54E-15 |
| HALLMARK_COMPLEMENT | 200 | Genes encoding components of the complement system, which is part of the innate immune system. | 23 | 0.115 | 2.27E-17 | 1.25E-14 |
| HALLMARK_EPITHELIAL_MESENCHYMAL_TRANSITION | 200 | Genes defining epithelial-mesenchymal transition, as in wound healing, fibrosis and metastasis. | 23 | 0.115 | 2.27E-17 | 1.25E-14 |
| REACTOME_HEMOSTASIS | 678 | Hemostasis | 38 | 0.056 | 5.61E-17 | 2.57E-14 |
| HALLMARK_INTERFERON_GAMMA_RESPONSE | 200 | Genes up-regulated in response to IFNG [GeneID=3458]. | 22 | 0.11 | 2.94E-16 | 1.15E-13 |
| HALLMARK_APOPTOSIS | 161 | Genes mediating programmed cell death (apoptosis) by activation of caspases. | 20 | 0.1242 | 6.21E-16 | 2.13E-13 |
| REACTOME_INTERLEUKIN_4_AND_INTERLEUKIN_13_SIGNALING | 111 | Interleukin-4 and Interleukin-13 signaling | 17 | 0.1532 | 2.71E-15 | 8.28E-13 |
| WP_IL18_SIGNALING_PATHWAY | 279 | IL-18 signaling pathway | 24 | 0.086 | 3.53E-15 | 9.71E-13 |
| REACTOME_SIGNALING_BY_RHO_GTPASES_MIRO_GTPASES_AND_RHOBTB3 | 717 | Signaling by Rho GTPases, Miro GTPases and RHOBTB3 | 36 | 0.0502 | 1.11E-14 | 2.77E-12 |
| REACTOME_SIGNALING_BY_INTERLEUKINS | 463 | Signaling by Interleukins | 29 | 0.0626 | 1.89E-14 | 4.32E-12 |
| REACTOME_NEUTROPHIL_DEGRANULATION | 479 | Neutrophil degranulation | 29 | 0.0605 | 4.48E-14 | 9.47E-12 |
| HALLMARK_HYPOXIA | 200 | Genes up-regulated in response to low oxygen levels (hypoxia). | 18 | 0.09 | 4.80E-12 | 9.42E-10 |
| REACTOME_SIGNALING_BY_RECEPTOR_TYROSINE_KINASES | 504 | Signaling by Receptor Tyrosine Kinases | 27 | 0.0536 | 5.91E-12 | 1.05E-09 |
| REACTOME_PLATELET_ACTIVATION_SIGNALING_AND_AGGREGATION | 261 | Platelet activation, signaling and aggregation | 20 | 0.0766 | 6.12E-12 | 1.05E-09 |
| HALLMARK_APICAL_JUNCTION | 200 | Genes encoding components of apical junction complex. | 17 | 0.085 | 4.69E-11 | 7.15E-09 |
| HALLMARK_INFLAMMATORY_RESPONSE | 200 | Genes defining inflammatory response. | 17 | 0.085 | 4.69E-11 | 7.15E-09 |
| REACTOME_RHO_GTPASE_CYCLE | 444 | RHO GTPase cycle | 23 | 0.0518 | 4.31E-10 | 6.24E-08 |
| REACTOME_EXTRACELLULAR_MATRIX_ORGANIZATION | 301 | Extracellular matrix organization | 19 | 0.0631 | 5.67E-10 | 7.79E-08 |
| REACTOME_REGULATION_OF_INSULIN_LIKE_GROWTH_FACTOR_IGF_TRANSPORT_AND_UPTAKE_BY_INSULIN_LIKE_GROWTH_FACTOR_BINDING_PROTEINS_IGFBPS | 124 | Regulation of Insulin-like Growth Factor (IGF) transport and uptake by Insulin-like Growth Factor Binding Proteins (IGFBPs) | 13 | 0.1048 | 6.84E-10 | 8.94E-08 |
| WP_NUCLEAR_RECEPTORS_METAPATHWAY | 321 | Nuclear Receptors Meta-Pathway | 19 | 0.0592 | 1.65E-09 | 2.06E-07 |
| HALLMARK_IL2_STAT5_SIGNALING | 199 | Genes up-regulated by STAT5 in response to IL2 stimulation. | 15 | 0.0754 | 3.42E-09 | 3.87E-07 |
| KEGG_LEUKOCYTE_TRANSENDOTHELIAL_MIGRATION | 116 | Leukocyte transendothelial migration | 12 | 0.1034 | 3.63E-09 | 3.87E-07 |
| HALLMARK_ESTROGEN_RESPONSE_EARLY | 200 | Genes defining early response to estrogen. | 15 | 0.075 | 3.67E-09 | 3.87E-07 |
| HALLMARK_P53_PATHWAY | 200 | Genes involved in p53 pathways and networks. | 15 | 0.075 | 3.67E-09 | 3.87E-07 |
| REACTOME_PROGRAMMED_CELL_DEATH | 208 | Programmed Cell Death | 15 | 0.0721 | 6.25E-09 | 6.36E-07 |
| REACTOME_REGULATED_NECROSIS | 56 | Regulated Necrosis | 9 | 0.1607 | 6.53E-09 | 6.40E-07 |
| REACTOME_POST_TRANSLATIONAL_PROTEIN_MODIFICATION | 1435 | Post-translational protein modification | 41 | 0.0286 | 7.27E-09 | 6.89E-07 |
| WP_VEGFAVEGFR2_SIGNALING_PATHWAY | 438 | VEGFA-VEGFR2 Signaling Pathway | 21 | 0.0479 | 9.69E-09 | 8.87E-07 |
| REACTOME_SIGNALING_BY_SCF_KIT | 43 | Signaling by SCF-KIT | 8 | 0.186 | 1.37E-08 | 1.21E-06 |
| WP_ADIPOGENESIS | 131 | Adipogenesis | 12 | 0.0916 | 1.45E-08 | 1.25E-06 |
| REACTOME_DEVELOPMENTAL_BIOLOGY | 1143 | Developmental Biology | 35 | 0.0306 | 1.80E-08 | 1.50E-06 |
| HALLMARK_COAGULATION | 138 | Genes encoding components of blood coagulation system; also up-regulated in platelets. | 12 | 0.087 | 2.62E-08 | 2.11E-06 |
| REACTOME_DEGRADATION_OF_THE_EXTRACELLULAR_MATRIX | 140 | Degradation of the extracellular matrix | 12 | 0.0857 | 3.07E-08 | 2.41E-06 |
| REACTOME_CELL_JUNCTION_ORGANIZATION | 92 | Cell junction organization | 10 | 0.1087 | 4.57E-08 | 3.48E-06 |
| WP_HEPATITIS_C_AND_HEPATOCELLULAR_CARCINOMA | 50 | Hepatitis C and Hepatocellular Carcinoma | 8 | 0.16 | 4.76E-08 | 3.53E-06 |
| WP_APOPTOSIS_MODULATION_AND_SIGNALING | 93 | Apoptosis Modulation and Signaling | 10 | 0.1075 | 5.07E-08 | 3.67E-06 |
| KEGG_PATHWAYS_IN_CANCER | 325 | Pathways in cancer | 17 | 0.0523 | 7.25E-08 | 5.11E-06 |
| WP_HUMAN_COMPLEMENT_SYSTEM | 99 | Human Complement System | 10 | 0.101 | 9.25E-08 | 6.35E-06 |
| REACTOME_TOLL_LIKE_RECEPTOR_CASCADES | 155 | Toll-like Receptor Cascades | 12 | 0.0774 | 9.51E-08 | 6.37E-06 |
| WP_IL4_SIGNALING_PATHWAY | 55 | IL-4 Signaling Pathway | 8 | 0.1455 | 1.03E-07 | 6.74E-06 |
| REACTOME_NUCLEOTIDE_BINDING_DOMAIN_LEUCINE_RICH_REPEAT_CONTAINING_RECEPTOR_NLR_SIGNALING_PATHWAYS | 56 | Nucleotide-binding domain, leucine rich repeat containing receptor (NLR) signaling pathways | 8 | 0.1429 | 1.19E-07 | 7.61E-06 |
| WP_EBOLA_VIRUS_PATHWAY_ON_HOST | 131 | Ebola Virus Pathway on Host | 11 | 0.084 | 1.43E-07 | 8.92E-06 |
| KEGG_TIGHT_JUNCTION | 132 | Tight junction | 11 | 0.0833 | 1.54E-07 | 9.22E-06 |
| REACTOME_RESPONSE_TO_ELEVATED_PLATELET_CYTOSOLIC_CA2 | 132 | Response to elevated platelet cytosolic Ca2+ | 11 | 0.0833 | 1.54E-07 | 9.22E-06 |
| WP_COMPLEMENT_AND_COAGULATION_CASCADES | 59 | Complement and Coagulation Cascades | 8 | 0.1356 | 1.81E-07 | 1.06E-05 |
| HALLMARK_ESTROGEN_RESPONSE_LATE | 200 | Genes defining late response to estrogen. | 13 | 0.065 | 2.16E-07 | 1.23E-05 |
| HALLMARK_UV_RESPONSE_DN | 144 | Genes down-regulated in response to ultraviolet (UV) radiation. | 11 | 0.0764 | 3.73E-07 | 2.09E-05 |
| REACTOME_RHO_GTPASE_EFFECTORS | 324 | RHO GTPase Effectors | 16 | 0.0494 | 3.80E-07 | 2.09E-05 |
| REACTOME_RIPK1_MEDIATED_REGULATED_NECROSIS | 29 | RIPK1-mediated regulated necrosis | 6 | 0.2069 | 4.79E-07 | 2.58E-05 |
| REACTOME_SIGNALING_BY_CSF3_G_CSF | 30 | Signaling by CSF3 (G-CSF) | 6 | 0.2 | 5.93E-07 | 3.13E-05 |
| REACTOME_TRAIL_SIGNALING | 8 | TRAIL signaling | 4 | 0.5 | 7.87E-07 | 4.08E-05 |
| REACTOME_VESICLE_MEDIATED_TRANSPORT | 724 | Vesicle-mediated transport | 24 | 0.0331 | 8.43E-07 | 4.29E-05 |
| WP_SMALL_CELL_LUNG_CANCER | 98 | Small cell lung cancer | 9 | 0.0918 | 9.22E-07 | 4.55E-05 |
| REACTOME_TRANSPORT_OF_SMALL_MOLECULES | 728 | Transport of small molecules | 24 | 0.033 | 9.28E-07 | 4.55E-05 |
| REACTOME_NERVOUS_SYSTEM_DEVELOPMENT | 580 | Nervous system development | 21 | 0.0362 | 1.03E-06 | 4.98E-05 |
| REACTOME_CELL_CELL_COMMUNICATION | 130 | Cell-Cell communication | 10 | 0.0769 | 1.19E-06 | 5.63E-05 |
| HALLMARK_ALLOGRAFT_REJECTION | 200 | Genes up-regulated during transplant rejection. | 12 | 0.06 | 1.47E-06 | 6.52E-05 |
| HALLMARK_GLYCOLYSIS | 200 | Genes encoding proteins involved in glycolysis and gluconeogenesis. | 12 | 0.06 | 1.47E-06 | 6.52E-05 |
| HALLMARK_KRAS_SIGNALING_UP | 200 | Genes up-regulated by KRAS activation. | 12 | 0.06 | 1.47E-06 | 6.52E-05 |
| WP_APOPTOSISRELATED_NETWORK_DUE_TO_ALTERED_NOTCH3_IN_OVARIAN_CANCER | 54 | Apoptosis-related network due to altered Notch3 in ovarian cancer | 7 | 0.1296 | 1.47E-06 | 6.52E-05 |
| REACTOME_INTERFERON_SIGNALING | 203 | Interferon Signaling | 12 | 0.0591 | 1.72E-06 | 7.50E-05 |
| REACTOME_SIGNALING_BY_MET | 79 | Signaling by MET | 8 | 0.1013 | 1.78E-06 | 7.63E-05 |
| REACTOME_NOD1_2_SIGNALING_PATHWAY | 36 | NOD1/2 Signaling Pathway | 6 | 0.1667 | 1.85E-06 | 7.80E-05 |
| WP_FGF23_SIGNALLING_IN_HYPOPHOSPHATEMIC_RICKETS_AND_RELATED_DISORDERS | 22 | FGF23 signalling in Hypophosphatemic rickets and related disorders | 5 | 0.2273 | 2.73E-06 | 1.14E-04 |
| WP_NRF2_PATHWAY | 145 | NRF2 pathway | 10 | 0.069 | 3.20E-06 | 1.31E-04 |
| WP_GASTRIN_SIGNALING_PATHWAY | 114 | Gastrin Signaling Pathway | 9 | 0.0789 | 3.28E-06 | 1.33E-04 |
| REACTOME_MEMBRANE_TRAFFICKING | 629 | Membrane Trafficking | 21 | 0.0334 | 3.67E-06 | 1.45E-04 |
| HALLMARK_IL6_JAK_STAT3_SIGNALING | 87 | Genes up-regulated by IL6 [GeneID=3569] via STAT3 [GeneID=6774], e.g., during acute phase response. | 8 | 0.092 | 3.70E-06 | 1.45E-04 |
| REACTOME_RHOA_GTPASE_CYCLE | 149 | RHOA GTPase cycle | 10 | 0.0671 | 4.09E-06 | 1.58E-04 |
| WP_PI3KAKT_SIGNALING_PATHWAY | 345 | PI3K-Akt Signaling Pathway | 15 | 0.0435 | 4.19E-06 | 1.60E-04 |
| REACTOME_INTERLEUKIN_6_FAMILY_SIGNALING | 24 | Interleukin-6 family signaling | 5 | 0.2083 | 4.33E-06 | 1.63E-04 |
| WP_VITAMIN_D_RECEPTOR_PATHWAY | 185 | Vitamin D Receptor Pathway | 11 | 0.0595 | 4.39E-06 | 1.63E-04 |
| REACTOME_EPHB_MEDIATED_FORWARD_SIGNALING | 42 | EPHB-mediated forward signaling | 6 | 0.1429 | 4.71E-06 | 1.73E-04 |
| REACTOME_INACTIVATION_OF_CSF3_G_CSF_SIGNALING | 25 | Inactivation of CSF3 (G-CSF) signaling | 5 | 0.2 | 5.36E-06 | 1.94E-04 |
| WP_FOCAL_ADHESIONPI3KAKTMTORSIGNALING_PATHWAY | 309 | Focal Adhesion-PI3K-Akt-mTOR-signaling pathway | 14 | 0.0453 | 5.48E-06 | 1.95E-04 |
| REACTOME_INTERFERON_GAMMA_SIGNALING | 93 | Interferon gamma signaling | 8 | 0.086 | 6.11E-06 | 2.15E-04 |
| KEGG_COMPLEMENT_AND_COAGULATION_CASCADES | 69 | Complement and coagulation cascades | 7 | 0.1014 | 7.84E-06 | 2.73E-04 |
| REACTOME_MYD88_INDEPENDENT_TLR4_CASCADE | 97 | MyD88-independent TLR4 cascade | 8 | 0.0825 | 8.36E-06 | 2.87E-04 |
| WP_THYMIC_STROMAL_LYMPHOPOIETIN_TSLP_SIGNALING_PATHWAY | 47 | Thymic Stromal LymphoPoietin (TSLP) Signaling Pathway | 6 | 0.1277 | 9.24E-06 | 3.13E-04 |
| WP_EPITHELIAL_TO_MESENCHYMAL_TRANSITION_IN_COLORECTAL_CANCER | 164 | Epithelial to mesenchymal transition in colorectal cancer | 10 | 0.061 | 9.56E-06 | 3.20E-04 |
| WP_P53_TRANSCRIPTIONAL_GENE_NETWORK | 72 | p53 transcriptional gene network | 7 | 0.0972 | 1.04E-05 | 3.45E-04 |
| REACTOME_INTERFERON_ALPHA_BETA_SIGNALING | 73 | Interferon alpha/beta signaling | 7 | 0.0959 | 1.14E-05 | 3.74E-04 |
| HALLMARK_CHOLESTEROL_HOMEOSTASIS | 74 | Genes involved in cholesterol homeostasis. | 7 | 0.0946 | 1.25E-05 | 4.04E-04 |
| REACTOME_TOLL_LIKE_RECEPTOR_TLR1_TLR2_CASCADE | 103 | Toll Like Receptor TLR1:TLR2 Cascade | 8 | 0.0777 | 1.30E-05 | 4.16E-04 |
| REACTOME_MET_ACTIVATES_PTK2_SIGNALING | 30 | MET activates PTK2 signaling | 5 | 0.1667 | 1.38E-05 | 4.35E-04 |
| WP_SENESCENCE_AND_AUTOPHAGY_IN_CANCER | 108 | Senescence and Autophagy in Cancer | 8 | 0.0741 | 1.84E-05 | 5.71E-04 |
| REACTOME_DEATH_RECEPTOR_SIGNALLING | 141 | Death Receptor Signalling | 9 | 0.0638 | 1.85E-05 | 5.71E-04 |
| REACTOME_REGULATION_OF_KIT_SIGNALING | 16 | Regulation of KIT signaling | 4 | 0.25 | 1.91E-05 | 5.84E-04 |
| REACTOME_APOPTOSIS | 179 | Apoptosis | 10 | 0.0559 | 2.05E-05 | 6.18E-04 |
| REACTOME_SIGNALING_BY_PTK6 | 54 | Signaling by PTK6 | 6 | 0.1111 | 2.09E-05 | 6.24E-04 |
| REACTOME_METAL_SEQUESTRATION_BY_ANTIMICROBIAL_PROTEINS | 6 | Metal sequestration by antimicrobial proteins | 3 | 0.5 | 2.19E-05 | 6.48E-04 |
| WP_PATHOGENIC_ESCHERICHIA_COLI_INFECTION | 55 | Pathogenic Escherichia coli infection | 6 | 0.1091 | 2.32E-05 | 6.69E-04 |
| WP_RANKLRANK_RECEPTOR_ACTIVATOR_OF_NFKB_LIGAND_SIGNALING_PATHWAY | 55 | RANKL/RANK (Receptor activator of NFKB (ligand)) Signaling Pathway | 6 | 0.1091 | 2.32E-05 | 6.69E-04 |
| REACTOME_ADAPTIVE_IMMUNE_SYSTEM | 825 | Adaptive Immune System | 23 | 0.0279 | 2.34E-05 | 6.69E-04 |
| KEGG_PATHOGENIC_ESCHERICHIA_COLI_INFECTION | 56 | Pathogenic Escherichia coli infection | 6 | 0.1071 | 2.58E-05 | 7.31E-04 |

TAL UP

| | | | | | | |
|---|---|---|---|---|---|---|
| KEGG_ECM_RECEPTOR_INTERACTION | 84 | ECM-receptor interaction | 7 | 0.0833 | 2.88E-05 | 7.99E-04 |
| KEGG_SMALL_CELL_LUNG_CANCER | 84 | Small cell lung cancer | 7 | 0.0833 | 2.88E-05 | 7.99E-04 |
| REACTOME_SIGNALING_BY_PDGF | 58 | Signaling by PDGF | 6 | 0.1034 | 3.16E-05 | 8.68E-04 |

Enrichment for genes upregulated in AKI vs. control

Overlap Results

| | |
|---|---|
| Collection(s): | CP:BIOCARTA, CP:KEGG, CP:REACTOME, CP:WIKIPATHWAYS, H |
| # overlaps shown: | 94 |
| # genesets in collections: | 2747 |
| # genes in comparison (n): | 185 |
| # genes in universe (N): | 40312 |

| Gene Set Name | # Genes in Gene Set (K) | Description | # Genes in Overlap (k) | k/K | p-value | FDR q-value |
|---|---|---|---|---|---|---|
| HALLMARK_APOPTOSIS | 161 | Genes mediating programmed cell death (apoptosis) by activation of caspases. | 15 | 0.0932 | 1.31E-15 | 3.59E-12 |
| HALLMARK_TNFA_SIGNALING_VIA_NFKB | 200 | Genes regulated by NF-kB in response to TNF [GeneID=7124]. | 15 | 0.075 | 3.32E-14 | 4.55E-11 |
| HALLMARK_HYPOXIA | 200 | Genes up-regulated in response to low oxygen levels (hypoxia). | 12 | 0.06 | 1.76E-10 | 1.61E-07 |
| REACTOME_CYTOKINE_SIGNALING_IN_IMMUNE_SYSTEM | 719 | Cytokine Signaling in Immune system | 19 | 0.0264 | 1.12E-09 | 7.71E-07 |
| HALLMARK_GLYCOLYSIS | 200 | Genes encoding proteins involved in glycolysis and gluconeogenesis. | 11 | 0.055 | 2.58E-09 | 1.15E-06 |
| HALLMARK_KRAS_SIGNALING_UP | 200 | Genes up-regulated by KRAS activation. | 11 | 0.055 | 2.58E-09 | 1.15E-06 |
| REACTOME_HEMOSTASIS | 678 | Hemostasis | 18 | 0.0265 | 2.93E-09 | 1.15E-06 |
| REACTOME_INNATE_IMMUNE_SYSTEM | 1117 | Innate Immune System | 22 | 0.0197 | 1.13E-08 | 3.88E-06 |
| HALLMARK_INTERFERON_GAMMA_RESPONSE | 200 | Genes up-regulated in response to IFNG [GeneID=3458]. | 10 | 0.05 | 3.43E-08 | 1.05E-05 |
| WP_NUCLEAR_RECEPTORS_METAPATHWAY | 321 | Nuclear Receptors Meta-Pathway | 11 | 0.0343 | 3.24E-07 | 8.89E-05 |
| WP_SMALL_CELL_LUNG_CANCER | 98 | Small cell lung cancer | 7 | 0.0714 | 3.72E-07 | 9.29E-05 |
| REACTOME_PLATELET_ACTIVATION_SIGNALING_AND_AGGREGATION | 261 | Platelet activation, signaling and aggregation | 10 | 0.0383 | 4.08E-07 | 9.34E-05 |
| REACTOME_INTERLEUKIN_4_AND_INTERLEUKIN_13_SIGNALING | 111 | Interleukin-4 and Interleukin-13 signaling | 7 | 0.0631 | 8.69E-07 | 1.84E-04 |
| REACTOME_EXTRACELLULAR_MATRIX_ORGANIZATION | 301 | Extracellular matrix organization | 10 | 0.0332 | 1.48E-06 | 2.91E-04 |
| REACTOME_SIGNALING_BY_INTERLEUKINS | 463 | Signaling by Interleukins | 12 | 0.0259 | 1.78E-06 | 3.15E-04 |
| REACTOME_REGULATION_OF_INSULIN_LIKE_GROWTH_FACTOR_IGF_TRANSPORT_AND_UPTAKE_BY_INSULIN_LIKE_GROWTH_FACTOR_BINDING_PROTEINS_IGFBPS | 124 | Regulation of Insulin-like Growth Factor (IGF) transport and uptake by Insulin-like Growth Factor Binding Proteins (IGFBPs) | 7 | 0.0565 | 1.83E-06 | 3.15E-04 |
| KEGG_ECM_RECEPTOR_INTERACTION | 84 | ECM-receptor interaction | 6 | 0.0714 | 2.60E-06 | 4.20E-04 |
| HALLMARK_IL6_JAK_STAT3_SIGNALING | 87 | Genes up-regulated by IL6 [GeneID=3569] via STAT3 [GeneID=6774], e.g., during acute phase response. | 6 | 0.069 | 3.19E-06 | 4.88E-04 |
| HALLMARK_ESTROGEN_RESPONSE_EARLY | 200 | Genes defining early response to estrogen. | 8 | 0.04 | 4.40E-06 | 6.36E-04 |
| WP_APOPTOSIS_MODULATION_AND_SIGNALING | 93 | Apoptosis Modulation and Signaling | 6 | 0.0645 | 4.71E-06 | 6.47E-04 |
| WP_APOPTOSISRELATED_NETWORK_DUE_TO_ALTERED_NOTCH3_IN_OVARIAN_CANCER | 54 | Apoptosis-related network due to altered Notch3 in ovarian cancer | 5 | 0.0926 | 5.08E-06 | 6.47E-04 |
| WP_NRF2_PATHWAY | 145 | NRF2 pathway | 7 | 0.0483 | 5.18E-06 | 6.47E-04 |
| REACTOME_NUCLEOTIDE_BINDING_DOMAIN_LEUCINE_RICH_REPEAT_CONTAINING_RECEPTOR_NLR_SIGNALING_PATHWAYS | 56 | Nucleotide-binding domain, leucine rich repeat containing receptor (NLR) signaling pathways | 5 | 0.0893 | 6.09E-06 | 7.28E-04 |
| WP_IL18_SIGNALING_PATHWAY | 279 | IL-18 signaling pathway | 9 | 0.0323 | 6.36E-06 | 7.28E-04 |
| BIOCARTA_SODD_PATHWAY | 10 | SODD/TNFR1 Signaling Pathway | 3 | 0.3 | 1.11E-05 | 1.22E-03 |
| WP_FOCAL_ADHESIONPI3KAKTMTORSIGNALING_PATHWAY | 309 | Focal Adhesion-PI3K-Akt-mTOR-signaling pathway | 9 | 0.0291 | 1.44E-05 | 1.52E-03 |
| REACTOME_NEUTROPHIL_DEGRANULATION | 479 | Neutrophil degranulation | 11 | 0.023 | 1.50E-05 | 1.53E-03 |
| WP_PI3KAKT_SIGNALING_PATHWAY | 345 | PI3K-Akt Signaling Pathway | 9 | 0.0261 | 3.41E-05 | 3.31E-03 |
| REACTOME_RESPONSE_TO_ELEVATED_PLATELET_CYTOSOLIC_CA2 | 132 | Response to elevated platelet cytosolic Ca2+ | 6 | 0.0455 | 3.49E-05 | 3.31E-03 |
| HALLMARK_IL2_STAT5_SIGNALING | 199 | Genes up-regulated by STAT5 in response to IL2 stimulation. | 7 | 0.0352 | 4.02E-05 | 3.56E-03 |
| HALLMARK_COMPLEMENT | 200 | Genes encoding components of the complement system, which is part of the innate immune system. | 7 | 0.035 | 4.15E-05 | 3.56E-03 |
| HALLMARK_P53_PATHWAY | 200 | Genes involved in p53 pathways and networks. | 7 | 0.035 | 4.15E-05 | 3.56E-03 |
| REACTOME_DEGRADATION_OF_THE_EXTRACELLULAR_MATRIX | 140 | Degradation of the extracellular matrix | 6 | 0.0429 | 4.86E-05 | 3.96E-03 |
| WP_APOPTOSIS | 86 | Apoptosis | 5 | 0.0581 | 4.97E-05 | 3.96E-03 |
| HALLMARK_APICAL_SURFACE | 44 | Genes encoding proteins over-represented on the apical surface of epithelial cells, e.g., important for cell polarity (apical area). | 4 | 0.0909 | 5.05E-05 | 3.96E-03 |
| REACTOME_FOXO_MEDIATED_TRANSCRIPTION_OF_CELL_CYCLE_GENES | 17 | FOXO-mediated transcription of cell cycle genes | 3 | 0.1765 | 6.17E-05 | 4.71E-03 |
| WP_HEPATITIS_C_AND_HEPATOCELLULAR_CARCINOMA | 50 | Hepatitis C and Hepatocellular Carcinoma | 4 | 0.08 | 8.38E-05 | 6.22E-03 |
| HALLMARK_INTERFERON_ALPHA_RESPONSE | 97 | Genes up-regulated in response to alpha interferon proteins. | 5 | 0.0515 | 8.83E-05 | 6.38E-03 |
| WP_PHOTODYNAMIC_THERAPYINDUCED_AP1_SURVIVAL_SIGNALING | 51 | Photodynamic therapy-induced AP-1 survival signaling. | 4 | 0.0784 | 9.07E-05 | 6.39E-03 |
| HALLMARK_ANDROGEN_RESPONSE | 100 | Genes defining response to androgens. | 5 | 0.05 | 1.02E-04 | 7.01E-03 |
| REACTOME_REGULATED_NECROSIS | 56 | Regulated Necrosis | 4 | 0.0714 | 1.31E-04 | 8.77E-03 |
| BIOCARTA_CASPASE_PATHWAY | 22 | Caspase Cascade in Apoptosis | 3 | 0.1364 | 1.37E-04 | 8.89E-03 |
| KEGG_PATHWAYS_IN_CANCER | 325 | Pathways in cancer | 8 | 0.0246 | 1.39E-04 | 8.89E-03 |
| REACTOME_DISEASES_OF_METABOLISM | 246 | Diseases of metabolism | 7 | 0.0285 | 1.51E-04 | 9.44E-03 |
| WP_PHOTODYNAMIC_THERAPYINDUCED_NFE2L2_NRF2_SURVIVAL_SIGNALING | 23 | Photodynamic therapy-induced NFE2L2 (NRF2) survival signaling | 3 | 0.1304 | 1.57E-04 | 9.61E-03 |
| WP_GASTRIN_SIGNALING_PATHWAY | 114 | Gastrin Signaling Pathway | 5 | 0.0439 | 1.89E-04 | 1.13E-02 |
| REACTOME_FOXO_MEDIATED_TRANSCRIPTION | 66 | FOXO-mediated transcription | 4 | 0.0606 | 2.48E-04 | 1.45E-02 |
| REACTOME_EFFECTS_OF_PIP2_HYDROLYSIS | 27 | Effects of PIP2 hydrolysis | 3 | 0.1111 | 2.56E-04 | 1.47E-02 |
| KEGG_P53_SIGNALING_PATHWAY | 68 | p53 signaling pathway | 4 | 0.0588 | 2.78E-04 | 1.53E-02 |
| REACTOME_GLYCOSAMINOGLYCAN_METABOLISM | 124 | Glycosaminoglycan metabolism | 5 | 0.0403 | 2.79E-04 | 1.53E-02 |
| REACTOME_CELL_SURFACE_INTERACTIONS_AT_THE_VASCULAR_WALL | 194 | Cell surface interactions at the vascular wall | 6 | 0.0309 | 2.89E-04 | 1.56E-02 |
| REACTOME_RIPK1_MEDIATED_REGULATED_NECROSIS | 29 | RIPK1-mediated regulated necrosis | 3 | 0.1034 | 3.18E-04 | 1.67E-02 |
| HALLMARK_ALLOGRAFT_REJECTION | 200 | Genes up-regulated during transplant rejection. | 6 | 0.03 | 3.40E-04 | 1.67E-02 |
| HALLMARK_APICAL_JUNCTION | 200 | Genes encoding components of apical junction complex. | 6 | 0.03 | 3.40E-04 | 1.67E-02 |
| HALLMARK_INFLAMMATORY_RESPONSE | 200 | Genes defining inflammatory response. | 6 | 0.03 | 3.40E-04 | 1.67E-02 |
| HALLMARK_XENOBIOTIC_METABOLISM | 200 | Genes encoding proteins involved in processing of drugs and other xenobiotics. | 6 | 0.03 | 3.40E-04 | 1.67E-02 |
| REACTOME_LAMININ_INTERACTIONS | 30 | Laminin interactions | 3 | 0.1 | 3.52E-04 | 1.67E-02 |
| REACTOME_MET_ACTIVATES_PTK2_SIGNALING | 30 | MET activates PTK2 signaling | 3 | 0.1 | 3.52E-04 | 1.67E-02 |
| REACTOME_INTERFERON_ALPHA_BETA_SIGNALING | 73 | Interferon alpha/beta signaling | 4 | 0.0548 | 3.65E-04 | 1.69E-02 |
| REACTOME_INTERFERON_SIGNALING | 203 | Interferon Signaling | 6 | 0.0296 | 3.68E-04 | 1.69E-02 |
| KEGG_ARRHYTHMOGENIC_RIGHT_VENTRICULAR_CARDIOMYOPATHY_ARVC | 74 | Arrhythmogenic right ventricular cardiomyopathy (ARVC) | 4 | 0.0541 | 3.84E-04 | 1.70E-02 |
| KEGG_CELL_ADHESION_MOLECULES_CAMS | 133 | Cell adhesion molecules (CAMs) | 5 | 0.0376 | 3.86E-04 | 1.70E-02 |
| REACTOME_HS_GAG_BIOSYNTHESIS | 31 | HS-GAG biosynthesis | 3 | 0.0968 | 3.89E-04 | 1.70E-02 |
| WP_ARRHYTHMOGENIC_RIGHT_VENTRICULAR_CARDIOMYOPATHY | 76 | Arrhythmogenic Right Ventricular Cardiomyopathy | 4 | 0.0526 | 4.26E-04 | 1.81E-02 |
| KEGG_ALANINE_ASPARTATE_AND_GLUTAMATE_METABOLISM | 32 | Alanine, aspartate and glutamate metabolism | 3 | 0.0938 | 4.28E-04 | 1.81E-02 |
| REACTOME_METABOLISM_OF_CARBOHYDRATES | 293 | Metabolism of carbohydrates | 7 | 0.0239 | 4.35E-04 | 1.81E-02 |
| WP_INFLAMMATORY_RESPONSE_PATHWAY | 33 | Inflammatory Response Pathway | 3 | 0.0909 | 4.69E-04 | 1.92E-02 |
| REACTOME_DISEASES_OF_GLYCOSYLATION | 143 | Diseases of glycosylation | 5 | 0.035 | 5.37E-04 | 2.17E-02 |
| REACTOME_REGULATION_OF_TNFR1_SIGNALING | 35 | Regulation of TNFR1 signaling | 3 | 0.0857 | 5.59E-04 | 2.22E-02 |
| REACTOME_SYNTHESIS_OF_UDP_N_ACETYL_GLUCOSAMINE | 8 | Synthesis of UDP-N-acetyl-glucosamine | 2 | 0.25 | 5.76E-04 | 2.23E-02 |
| REACTOME_TRAIL_SIGNALING | 8 | TRAIL signaling | 2 | 0.25 | 5.76E-04 | 2.23E-02 |
| REACTOME_NOD1_2_SIGNALING_PATHWAY | 36 | NOD1/2 Signaling Pathway | 3 | 0.0833 | 6.07E-04 | 2.32E-02 |
| KEGG_SMALL_CELL_LUNG_CANCER | 84 | Small cell lung cancer | 4 | 0.0476 | 6.22E-04 | 2.34E-02 |
| REACTOME_METABOLISM_OF_LIPIDS | 741 | Metabolism of lipids | 11 | 0.0148 | 6.76E-04 | 2.51E-02 |
| KEGG_APOPTOSIS | 87 | Apoptosis | 4 | 0.046 | 7.10E-04 | 2.60E-02 |
| WP_FERROPTOSIS | 40 | Ferroptosis | 3 | 0.075 | 8.29E-04 | 3.00E-02 |
| REACTOME_CELL_JUNCTION_ORGANIZATION | 92 | Cell junction organization | 4 | 0.0435 | 8.76E-04 | 3.08E-02 |
| REACTOME_EPH_EPHRIN_SIGNALING | 92 | EPH-Ephrin signaling | 4 | 0.0435 | 8.76E-04 | 3.08E-02 |
| REACTOME_MET_PROMOTES_CELL_MOTILITY | 41 | MET promotes cell motility | 3 | 0.0732 | 8.92E-04 | 3.10E-02 |
| REACTOME_INTERFERON_GAMMA_SIGNALING | 93 | Interferon gamma signaling | 4 | 0.043 | 9.12E-04 | 3.13E-02 |
| WP_TNF_ALPHA_SIGNALING_PATHWAY | 94 | TNF alpha Signaling Pathway | 4 | 0.0426 | 9.49E-04 | 3.21E-02 |
| REACTOME_EPHB_MEDIATED_FORWARD_SIGNALING | 42 | EPHB-mediated forward signaling | 3 | 0.0714 | 9.57E-04 | 3.21E-02 |
| REACTOME_POST_TRANSLATIONAL_PROTEIN_MODIFICATION | 1435 | Post-translational protein modification | 16 | 0.0111 | 1.01E-03 | 3.34E-02 |
| REACTOME_RHO_GTPASE_CYCLE | 444 | RHO GTPase cycle | 8 | 0.018 | 1.09E-03 | 3.50E-02 |
| KEGG_AMINO_SUGAR_AND_NUCLEOTIDE_SUGAR_METABOLISM | 44 | Amino sugar and nucleotide sugar metabolism | 3 | 0.0682 | 1.10E-03 | 3.50E-02 |
| REACTOME_TNF_SIGNALING | 44 | TNF signaling | 3 | 0.0682 | 1.10E-03 | 3.50E-02 |
| REACTOME_TYPE_I_HEMIDESMOSOME_ASSEMBLY | 11 | Type I hemidesmosome assembly | 2 | 0.1818 | 1.12E-03 | 3.54E-02 |
| WP_PROSTAGLANDIN_SYNTHESIS_AND_REGULATION | 45 | Prostaglandin Synthesis and Regulation | 3 | 0.0667 | 1.17E-03 | 3.66E-02 |
| WP_NANOPARTICLE_TRIGGERED_REGULATED_NECROSIS | 12 | Nanoparticle triggered regulated necrosis | 2 | 0.1667 | 1.34E-03 | 4.14E-02 |
| KEGG_CYTOKINE_CYTOKINE_RECEPTOR_INTERACTION | 265 | Cytokine-cytokine receptor interaction | 6 | 0.0226 | 1.47E-03 | 4.48E-02 |
| HALLMARK_REACTIVE_OXYGEN_SPECIES_PATHWAY | 49 | Genes up-regulated by reactive oxigen species (ROS). | 3 | 0.0612 | 1.50E-03 | 4.48E-02 |
| KEGG_GLYCEROLIPID_METABOLISM | 49 | Glycerolipid metabolism | 3 | 0.0612 | 1.50E-03 | 4.48E-02 |
| REACTOME_ADVANCED_GLYCOSYLATION_ENDPRODUCT_RECEPTOR_SIGNALING | 13 | Advanced glycosylation endproduct receptor signaling | 2 | 0.1538 | 1.58E-03 | 4.62E-02 |
| REACTOME_TNFR1_INDUCED_PROAPOPTOTIC_SIGNALING | 13 | TNFR1-induced proapoptotic signaling | 2 | 0.1538 | 1.58E-03 | 4.62E-02 |

Enrichment for genes downregulated in AKI vs. control

Overlap Results

| | |
|---|---|
| Collection(s): | CP:BIOCARTA, CP:KEGG, CP:REACTOME, CP:WIKIPATHWAYS, H |
| # overlaps shown: | 1 |
| # genesets in collections: | 2747 |
| # genes in comparison (n): | 91 |
| # genes in universe (N): | 40312 |

| Gene Set Name | # Genes in Gene Set (K) | Description | # Genes in Overlap (k) | k/K | p-value | FDR q-value |
|---|---|---|---|---|---|---|
| REACTOME_TRANSPORT_OF_SMALL_MOLECULES | 728 | Transport of small molecules | 10 | 0.0137 | 5.98E-06 | 1.64E-02 |

Enrichment for genes upregulated in AKI vs. control

Overlap Results

| | |
|---|---|
| Collection(s): | CP:BIOCARTA, CP:KEGG, CP:REACTOME, CP:WIKIPATHWAYS, H |
| # overlaps shown: | 89 |
| # genesets in collections: | 2747 |
| # genes in comparison (n): | 186 |
| # genes in universe (N): | 40312 |

| Gene Set Name | # Genes in Gene Set (K) | Description | # Genes in Overlap (k) | k/K | p-value | FDR q-value |
|---|---|---|---|---|---|---|
| HALLMARK_TNFA_SIGNALING_VIA_NFKB | 200 | Genes regulated by NF-kB in response to TNF [GeneID=7124]. | 14 | 0.07 | 6.75E-13 | 1.86E-09 |
| REACTOME_INNATE_IMMUNE_SYSTEM | 1117 | Innate Immune System | 27 | 0.0242 | 2.35E-12 | 3.03E-09 |
| REACTOME_CYTOKINE_SIGNALING_IN_IMMUNE_SYSTEM | 719 | Cytokine Signaling in Immune system | 22 | 0.0306 | 3.31E-12 | 3.03E-09 |
| HALLMARK_GLYCOLYSIS | 200 | Genes encoding proteins involved in glycolysis and gluconeogenesis. | 13 | 0.065 | 1.17E-11 | 5.64E-09 |
| HALLMARK_INTERFERON_GAMMA_RESPONSE | 200 | Genes up-regulated in response to IFNG [GeneID=3458]. | 13 | 0.065 | 1.17E-11 | 5.64E-09 |
| REACTOME_NEUTROPHIL_DEGRANULATION | 479 | Neutrophil degranulation | 18 | 0.0376 | 1.23E-11 | 5.64E-09 |
| HALLMARK_HYPOXIA | 200 | Genes up-regulated in response to low oxygen levels (hypoxia). | 12 | 0.06 | 1.87E-10 | 7.34E-08 |
| REACTOME_INTERFERON_ALPHA_BETA_SIGNALING | 73 | Interferon alpha/beta signaling | 8 | 0.1096 | 1.84E-09 | 6.31E-07 |
| HALLMARK_IL2_STAT5_SIGNALING | 199 | Genes up-regulated by STAT5 in response to IL2 stimulation. | 11 | 0.0553 | 2.59E-09 | 7.23E-07 |
| REACTOME_INTERLEUKIN_4_AND_INTERLEUKIN_13_SIGNALING | 111 | Interleukin-4 and Interleukin-13 signaling | 9 | 0.0811 | 2.63E-09 | 7.23E-07 |
| REACTOME_INTERFERON_GAMMA_SIGNALING | 93 | Interferon gamma signaling | 8 | 0.086 | 1.29E-08 | 3.21E-06 |
| REACTOME_TRAIL_SIGNALING | 8 | TRAIL signaling | 4 | 0.5 | 3.03E-08 | 6.93E-06 |
| HALLMARK_COMPLEMENT | 200 | Genes encoding components of the complement system, which is part of the innate immune system. | 10 | 0.05 | 3.61E-08 | 7.63E-06 |
| REACTOME_INTERFERON_SIGNALING | 203 | Interferon Signaling | 10 | 0.0493 | 4.16E-08 | 8.16E-06 |
| REACTOME_HEMOSTASIS | 678 | Hemostasis | 16 | 0.0236 | 1.24E-07 | 2.27E-05 |
| HALLMARK_IL6_JAK_STAT3_SIGNALING | 87 | Genes up-regulated by IL6 [GeneID=3569] via STAT3 [GeneID=6774], e.g., during acute phase response. | 7 | 0.0805 | 1.70E-07 | 2.92E-05 |
| KEGG_CELL_ADHESION_MOLECULES_CAMS | 133 | Cell adhesion molecules (CAMs) | 8 | 0.0602 | 2.12E-07 | 3.42E-05 |
| REACTOME_SIGNALING_BY_INTERLEUKINS | 463 | Signaling by Interleukins | 13 | 0.0281 | 2.83E-07 | 4.31E-05 |
| HALLMARK_ESTROGEN_RESPONSE_EARLY | 200 | Genes defining early response to estrogen. | 9 | 0.045 | 4.31E-07 | 6.22E-05 |
| WP_IL18_SIGNALING_PATHWAY | 279 | IL-18 signaling pathway | 10 | 0.0358 | 7.87E-07 | 1.08E-04 |
| KEGG_LEUKOCYTE_TRANSENDOTHELIAL_MIGRATION | 116 | Leukocyte transendothelial migration | 7 | 0.0603 | 1.21E-06 | 1.59E-04 |
| REACTOME_EXTRACELLULAR_MATRIX_ORGANIZATION | 301 | Extracellular matrix organization | 10 | 0.0332 | 1.56E-06 | 1.95E-04 |
| KEGG_NATURAL_KILLER_CELL_MEDIATED_CYTOTOXICITY | 137 | Natural killer cell mediated cytotoxicity | 7 | 0.0511 | 3.69E-06 | 4.41E-04 |
| HALLMARK_ALLOGRAFT_REJECTION | 200 | Genes up-regulated during transplant rejection. | 8 | 0.04 | 4.58E-06 | 4.66E-04 |
| HALLMARK_EPITHELIAL_MESENCHYMAL_TRANSITION | 200 | Genes defining epithelial-mesenchymal transition, as in wound healing, fibrosis and metastasis. | 8 | 0.04 | 4.58E-06 | 4.66E-04 |
| HALLMARK_INFLAMMATORY_RESPONSE | 200 | Genes defining inflammatory response. | 8 | 0.04 | 4.58E-06 | 4.66E-04 |
| HALLMARK_KRAS_SIGNALING_UP | 200 | Genes up-regulated by KRAS activation. | 8 | 0.04 | 4.58E-06 | 4.66E-04 |
| WP_APOPTOSIS_MODULATION_AND_SIGNALING | 93 | Apoptosis Modulation and Signaling | 6 | 0.0645 | 4.86E-06 | 4.77E-04 |
| WP_APOPTOSISRELATED_NETWORK_DUE_TO_ALTERED_NOTCH3_IN_OVARIAN_CANCER | 54 | Apoptosis-related network due to altered Notch3 in ovarian cancer | 5 | 0.0926 | 5.22E-06 | 4.94E-04 |
| HALLMARK_INTERFERON_ALPHA_RESPONSE | 97 | Genes up-regulated in response to alpha interferon proteins. | 6 | 0.0619 | 6.21E-06 | 5.68E-04 |
| HALLMARK_APOPTOSIS | 161 | Genes mediating programmed cell death (apoptosis) by activation of caspases. | 7 | 0.0435 | 1.07E-05 | 9.44E-04 |
| REACTOME_ENDOSOMAL_VACUOLAR_PATHWAY | 11 | Endosomal/Vacuolar pathway | 3 | 0.2727 | 1.55E-05 | 1.33E-03 |
| WP_TYPE_II_INTERFERON_SIGNALING_IFNG | 37 | Type II interferon signaling (IFNG) | 4 | 0.1081 | 2.57E-05 | 2.14E-03 |
| WP_TRANSCRIPTIONAL_CASCADE_REGULATING_ADIPOGENESIS | 13 | Transcriptional cascade regulating adipogenesis | 3 | 0.2308 | 2.67E-05 | 2.16E-03 |
| REACTOME_CELL_CELL_COMMUNICATION | 130 | Cell-Cell communication | 6 | 0.0462 | 3.30E-05 | 2.59E-03 |
| WP_ADIPOGENESIS | 131 | Adipogenesis | 6 | 0.0458 | 3.45E-05 | 2.63E-03 |
| REACTOME_CELL_SURFACE_INTERACTIONS_AT_THE_VASCULAR_WALL | 194 | Cell surface interactions at the vascular wall | 7 | 0.0361 | 3.54E-05 | 2.63E-03 |
| HALLMARK_APICAL_JUNCTION | 200 | Genes encoding components of apical junction complex. | 7 | 0.035 | 4.29E-05 | 3.10E-03 |
| KEGG_ECM_RECEPTOR_INTERACTION | 84 | ECM-receptor interaction | 5 | 0.0595 | 4.55E-05 | 3.17E-03 |
| HALLMARK_COAGULATION | 138 | Genes encoding components of blood coagulation system; also up-regulated in platelets. | 6 | 0.0435 | 4.62E-05 | 3.17E-03 |
| REACTOME_DEGRADATION_OF_THE_EXTRACELLULAR_MATRIX | 140 | Degradation of the extracellular matrix | 6 | 0.0429 | 5.00E-05 | 3.33E-03 |
| REACTOME_TP53_REGULATES_TRANSCRIPTION_OF_CELL_DEATH_GENES | 44 | TP53 Regulates Transcription of Cell Death Genes | 4 | 0.0909 | 5.16E-05 | 3.33E-03 |
| REACTOME_DEATH_RECEPTOR_SIGNALLING | 141 | Death Receptor Signalling | 6 | 0.0426 | 5.21E-05 | 3.33E-03 |
| HALLMARK_UV_RESPONSE_DN | 144 | Genes down-regulated in response to ultraviolet (UV) radiation. | 6 | 0.0417 | 5.85E-05 | 3.65E-03 |
| KEGG_JAK_STAT_SIGNALING_PATHWAY | 155 | Jak-STAT signaling pathway | 6 | 0.0387 | 8.80E-05 | 5.37E-03 |
| HALLMARK_ANDROGEN_RESPONSE | 100 | Genes defining response to androgens. | 5 | 0.05 | 1.05E-04 | 6.25E-03 |
| WP_IL4_SIGNALING_PATHWAY | 55 | IL-4 Signaling Pathway | 4 | 0.0727 | 1.25E-04 | 7.28E-03 |
| REACTOME_REGULATED_NECROSIS | 56 | Regulated Necrosis | 4 | 0.0714 | 1.34E-04 | 7.65E-03 |
| REACTOME_CHAPERONE_MEDIATED_AUTOPHAGY | 22 | Chaperone Mediated Autophagy | 3 | 0.1364 | 1.40E-04 | 7.82E-03 |
| WP_THE_OVERLAP_BETWEEN_SIGNAL_TRANSDUCTION_PATHWAYS_THAT_CONTRIBUTE_TO_A_RANGE_OF_LMNA_LAMINOPATHIES | 60 | The Overlap Between Signal Transduction Pathways that Contribute to a Range of LMNA Laminopathies | 4 | 0.0667 | 1.75E-04 | 9.61E-03 |
| REACTOME_ANTIGEN_PRESENTATION_FOLDING_ASSEMBLY_AND_PEPTIDE_LOADING_OF_CLASS_I_MHC | 25 | Antigen Presentation: Folding, assembly and peptide loading of class I MHC | 3 | 0.12 | 2.06E-04 | 1.11E-02 |
| WP_LUNG_FIBROSIS | 63 | Lung fibrosis | 4 | 0.0635 | 2.11E-04 | 1.12E-02 |
| REACTOME_PLATELET_ACTIVATION_SIGNALING_AND_AGGREGATION | 261 | Platelet activation, signaling and aggregation | 7 | 0.0268 | 2.24E-04 | 1.16E-02 |
| WP_VITAMIN_D_RECEPTOR_PATHWAY | 185 | Vitamin D Receptor Pathway | 6 | 0.0324 | 2.31E-04 | 1.17E-02 |
| WP_ENDOCHONDRAL_OSSIFICATION | 65 | Endochondral Ossification | 4 | 0.0615 | 2.38E-04 | 1.17E-02 |
| WP_ENDOCHONDRAL_OSSIFICATION_WITH_SKELETAL_DYSPLASIAS | 65 | Endochondral Ossification with Skeletal Dysplasias | 4 | 0.0615 | 2.38E-04 | 1.17E-02 |
| KEGG_CYTOKINE_CYTOKINE_RECEPTOR_INTERACTION | 265 | Cytokine-cytokine receptor interaction | 7 | 0.0264 | 2.46E-04 | 1.19E-02 |
| WP_ONCOSTATIN_M_SIGNALING_PATHWAY | 66 | Oncostatin M Signaling Pathway | 4 | 0.0606 | 2.53E-04 | 1.20E-02 |
| REACTOME_REGULATION_OF_INSULIN_LIKE_GROWTH_FACTOR_IGF_TRANSPORT_AND_UPTAKE_BY_INSULIN_LIKE_GROWTH_FACTOR_BINDING_PROTEINS_IGFBPS | 124 | Regulation of Insulin-like Growth Factor (IGF) transport and uptake by Insulin-like Growth Factor Binding Proteins (IGFBPs) | 5 | 0.0403 | 2.86E-04 | 1.33E-02 |
| REACTOME_METAL_SEQUESTRATION_BY_ANTIMICROBIAL_PROTEINS | 6 | Metal sequestration by antimicrobial proteins | 2 | 0.3333 | 3.14E-04 | 1.44E-02 |
| WP_GLUCOCORTICOID_RECEPTOR_PATHWAY | 71 | Glucocorticoid Receptor Pathway | 4 | 0.0563 | 3.35E-04 | 1.51E-02 |
| HALLMARK_ESTROGEN_RESPONSE_LATE | 200 | Genes defining late response to estrogen. | 6 | 0.03 | 3.50E-04 | 1.53E-02 |
| HALLMARK_MTORC1_SIGNALING | 200 | Genes up-regulated through activation of mTORC1 complex. | 6 | 0.03 | 3.50E-04 | 1.53E-02 |
| REACTOME_TIGHT_JUNCTION_INTERACTIONS | 30 | Tight junction interactions | 3 | 0.1 | 3.58E-04 | 1.54E-02 |
| REACTOME_RESPONSE_TO_ELEVATED_PLATELET_CYTOSOLIC_CA2 | 132 | Response to elevated platelet cytosolic Ca2+ | 5 | 0.0379 | 3.82E-04 | 1.61E-02 |
| REACTOME_CLASS_I_MHC_MEDIATED_ANTIGEN_PROCESSING_PRESENTATION | 377 | Class I MHC mediated antigen processing & presentation | 8 | 0.0212 | 3.91E-04 | 1.63E-02 |
| REACTOME_PROGRAMMED_CELL_DEATH | 208 | Programmed Cell Death | 6 | 0.0288 | 4.31E-04 | 1.75E-02 |
| WP_PROLACTIN_SIGNALING_PATHWAY | 76 | Prolactin Signaling Pathway | 4 | 0.0526 | 4.34E-04 | 1.75E-02 |
| REACTOME_METABOLISM_OF_CARBOHYDRATES | 293 | Metabolism of carbohydrates | 7 | 0.0239 | 4.50E-04 | 1.79E-02 |
| WP_WHITE_FAT_CELL_DIFFERENTIATION | 33 | White fat cell differentiation | 3 | 0.0909 | 4.76E-04 | 1.87E-02 |
| REACTOME_ADAPTIVE_IMMUNE_SYSTEM | 825 | Adaptive Immune System | 12 | 0.0145 | 4.87E-04 | 1.88E-02 |
| REACTOME_LATE_ENDOSOMAL_MICROAUTOPHAGY | 34 | Late endosomal microautophagy | 3 | 0.0882 | 5.21E-04 | 1.99E-02 |
| REACTOME_SIGNALING_BY_RECEPTOR_TYROSINE_KINASES | 504 | Signaling by Receptor Tyrosine Kinases | 9 | 0.0179 | 5.97E-04 | 2.25E-02 |
| WP_APOPTOSIS | 86 | Apoptosis | 4 | 0.0465 | 6.94E-04 | 2.58E-02 |
| KEGG_APOPTOSIS | 87 | Apoptosis | 4 | 0.046 | 7.25E-04 | 2.65E-02 |
| WP_NUCLEAR_RECEPTORS_METAPATHWAY | 321 | Nuclear Receptors Meta-Pathway | 7 | 0.0218 | 7.70E-04 | 2.78E-02 |
| WP_THE_INFLUENCE_OF_LAMINOPATHIES_ON_WNT_SIGNALING | 40 | The influence of laminopathies on Wnt signaling | 3 | 0.075 | 8.42E-04 | 3.00E-02 |
| REACTOME_CELL_JUNCTION_ORGANIZATION | 92 | Cell junction organization | 4 | 0.0435 | 8.93E-04 | 3.15E-02 |
| REACTOME_MET_PROMOTES_CELL_MOTILITY | 41 | MET promotes cell motility | 3 | 0.0732 | 9.06E-04 | 3.15E-02 |
| WP_EPITHELIAL_TO_MESENCHYMAL_TRANSITION_IN_COLORECTAL_CANCER | 164 | Epithelial to mesenchymal transition in colorectal cancer | 5 | 0.0305 | 1.02E-03 | 3.50E-02 |
| WP_IL6_SIGNALING_PATHWAY | 43 | IL-6 signaling pathway | 3 | 0.0698 | 1.04E-03 | 3.53E-02 |
| REACTOME_RHO_GTPASE_CYCLE | 444 | RHO GTPase cycle | 8 | 0.018 | 1.13E-03 | 3.71E-02 |
| REACTOME_REGULATION_BY_C_FLIP | 11 | Regulation by c-FLIP | 2 | 0.1818 | 1.13E-03 | 3.71E-02 |
| WP_GLYCOLYSIS_IN_SENESCENCE | 11 | Glycolysis in senescence | 2 | 0.1818 | 1.13E-03 | 3.71E-02 |
| REACTOME_DAP12_INTERACTIONS | 45 | DAP12 interactions | 3 | 0.0667 | 1.19E-03 | 3.84E-02 |
| KEGG_TYPE_II_DIABETES_MELLITUS | 47 | Type II diabetes mellitus | 3 | 0.0638 | 1.35E-03 | 4.23E-02 |
| REACTOME_TP53_REGULATES_TRANSCRIPTION_OF_DEATH_RECEPTORS_AND_LIGANDS | 12 | TP53 Regulates Transcription of Death Receptors and Ligands | 2 | 0.1667 | 1.36E-03 | 4.23E-02 |
| WP_COMPUTATIONAL_MODEL_OF_AEROBIC_GLYCOLYSIS | 12 | Computational Model of Aerobic Glycolysis | 2 | 0.1667 | 1.36E-03 | 4.23E-02 |
| WP_MAMMARY_GLAND_DEVELOPMENT_PATHWAY_PUBERTY_STAGE_2_OF_4 | 13 | Mammary gland development pathway - Puberty (Stage 2 of 4) | 2 | 0.1538 | 1.60E-03 | 4.93E-02 |

CNT DOWN

Enrichment for genes downregulated in AKI vs. control

Overlap Results

| | |
|---|---|
| Collection(s): | CP:BIOCARTA, CP:KEGG, CP:REACTOME, CP:WIKIPATHWAYS, H |
| # overlaps shown: | 21 |
| # genesets in collections: | 2747 |
| # genes in comparison (n): | 125 |
| # genes in universe (N): | 40312 |

| Gene Set Name | # Genes in Gene Set (K) | Description | # Genes in Overlap (k) | k/K | p-value | FDR q-value |
|---|---|---|---|---|---|---|
| REACTOME_TRANSPORT_OF_SMALL_MOLECULES | 728 | Transport of small molecules | 21 | 0.0288 | 1.14E-14 | 3.14E-11 |
| REACTOME_ION_CHANNEL_TRANSPORT | 183 | Ion channel transport | 11 | 0.0601 | 1.48E-11 | 2.03E-08 |
| REACTOME_STIMULI_SENSING_CHANNELS | 106 | Stimuli-sensing channels | 9 | 0.0849 | 5.04E-11 | 4.62E-08 |
| KEGG_ALDOSTERONE_REGULATED_SODIUM_REABSORPTION | 42 | Aldosterone-regulated sodium reabsorption | 6 | 0.1429 | 3.77E-09 | 2.59E-06 |
| REACTOME_SLC_MEDIATED_TRANSMEMBRANE_TRANSPORT | 250 | SLC-mediated transmembrane transport | 8 | 0.032 | 1.23E-06 | 6.66E-04 |
| REACTOME_HEMOSTASIS | 678 | Hemostasis | 12 | 0.0177 | 1.45E-06 | 6.66E-04 |
| HALLMARK_KRAS_SIGNALING_DN | 200 | Genes down-regulated by KRAS activation. | 7 | 0.035 | 3.24E-06 | 1.27E-03 |
| WP_VITAMIN_DSENSITIVE_CALCIUM_SIGNALING_IN_DEPRESSION | 40 | Vitamin D-sensitive calcium signaling in depression | 4 | 0.1 | 7.38E-06 | 2.41E-03 |
| WP_CALCIUM_REGULATION_IN_THE_CARDIAC_CELL | 152 | Calcium Regulation in the Cardiac Cell | 6 | 0.0395 | 8.44E-06 | 2.41E-03 |
| WP_SPLICING_FACTOR_NOVA_REGULATED_SYNAPTIC_PROTEINS | 42 | Splicing factor NOVA regulated synaptic proteins | 4 | 0.0952 | 9.00E-06 | 2.41E-03 |
| REACTOME_GABA_B_RECEPTOR_ACTIVATION | 43 | GABA B receptor activation | 4 | 0.093 | 9.90E-06 | 2.41E-03 |
| WP_MYOMETRIAL_RELAXATION_AND_CONTRACTION_PATHWAYS | 158 | Myometrial Relaxation and Contraction Pathways | 6 | 0.038 | 1.05E-05 | 2.41E-03 |
| REACTOME_TRANSPORT_OF_INORGANIC_CATIONS_ANIONS_AND_AMINO_ACIDS_OLIGOPEPTIDES | 106 | Transport of inorganic cations/anions and amino acids/oligopeptides | 5 | 0.0472 | 2.09E-05 | 4.41E-03 |
| WP_ESC_PLURIPOTENCY_PATHWAYS | 118 | ESC Pluripotency Pathways | 5 | 0.0424 | 3.50E-05 | 6.86E-03 |
| REACTOME_GABA_RECEPTOR_ACTIVATION | 60 | GABA receptor activation | 4 | 0.0667 | 3.76E-05 | 6.88E-03 |
| WP_RAS_AND_BRADYKININ_PATHWAYS_IN_COVID19 | 29 | RAS and Bradykinin pathways in COVID-19 | 3 | 0.1034 | 1.00E-04 | 1.72E-02 |
| REACTOME_INTERACTION_BETWEEN_L1_AND_ANKYRINS | 31 | Interaction between L1 and Ankyrins | 3 | 0.0968 | 1.23E-04 | 1.98E-02 |
| REACTOME_PLATELET_HOMEOSTASIS | 86 | Platelet homeostasis | 4 | 0.0465 | 1.54E-04 | 2.35E-02 |
| REACTOME_INWARDLY_RECTIFYING_K_CHANNELS | 35 | Inwardly rectifying K+ channels | 3 | 0.0857 | 1.77E-04 | 2.56E-02 |
| REACTOME_CATION_COUPLED_CHLORIDE_COTRANSPORTERS | 7 | Cation-coupled Chloride cotransporters | 2 | 0.2857 | 1.98E-04 | 2.72E-02 |
| WP_G_PROTEIN_SIGNALING_PATHWAYS | 96 | G Protein Signaling Pathways | 4 | 0.0417 | 2.35E-04 | 3.07E-02 |

Enrichment for genes upregulated in AKI vs. control

Overlap Results

| | |
|---|---|
| Collection(s): | CP:BIOCARTA, CP:KEGG, CP:REACTOME, CP:WIKIPATHWAYS, H |
| # overlaps shown: | 9 |
| # genesets in collections: | 2747 |
| # genes in comparison (n): | 28 |
| # genes in universe (N): | 40312 |

| Gene Set Name | # Genes in Gene Set (K) | Description | # Genes in Overlap (k) | k/K | p-value | FDR q-value |
|---|---|---|---|---|---|---|
| HALLMARK_KRAS_SIGNALING_UP | 200 | Genes up-regulated by KRAS activation. | 4 | 0.02 | 1.10E-05 | 1.17E-02 |
| WP_ENDOCHONDRAL_OSSIFICATION | 65 | Endochondral Ossification | 3 | 0.0462 | 1.27E-05 | 1.17E-02 |
| WP_ENDOCHONDRAL_OSSIFICATION_WITH_SKELETAL_DYSPLASIAS | 65 | Endochondral Ossification with Skeletal Dysplasias | 3 | 0.0462 | 1.27E-05 | 1.17E-02 |
| REACTOME_INTERFERON_ALPHA_BETA_SIGNALING | 73 | Interferon alpha/beta signaling | 3 | 0.0411 | 1.81E-05 | 1.24E-02 |
| HALLMARK_PEROXISOME | 104 | Genes encoding components of peroxisome. | 3 | 0.0288 | 5.21E-05 | 2.48E-02 |
| WP_OSTEOCLAST_SIGNALING | 16 | Osteoclast Signaling | 2 | 0.125 | 5.55E-05 | 2.48E-02 |
| REACTOME_INTERLEUKIN_4_AND_INTERLEUKIN_13_SIGNALING | 111 | Interleukin-4 and Interleukin-13 signaling | 3 | 0.027 | 6.33E-05 | 2.48E-02 |
| REACTOME_CYTOKINE_SIGNALING_IN_IMMUNE_SYSTEM | 719 | Cytokine Signaling in Immune system | 5 | 0.007 | 1.24E-04 | 3.85E-02 |
| REACTOME_DEGRADATION_OF_THE_EXTRACELLULAR_MATRIX | 140 | Degradation of the extracellular matrix | 3 | 0.0214 | 1.26E-04 | 3.85E-02 |

Enrichment for genes upregulated in AKI vs. control

Overlap Results

| | |
|---|---|
| Collection(s): | CP:BIOCARTA, CP:KEGG, CP:REACTOME, CP:WIKIPATHWAYS, H |
| # overlaps shown: | 100 |
| # genesets in collections: | 2747 |
| # genes in comparison (n): | 61 |
| # genes in universe (N): | 40312 |

| Gene Set Name | # Genes in Gene Set (K) | Description | # Genes in Overlap (k) | k/K | p-value | FDR q-value |
|---|---|---|---|---|---|---|
| HALLMARK_INTERFERON_GAMMA_RESPONSE | 200 | Genes up-regulated in response to IFNG [GeneID=3458]. | 11 | 0.055 | 1.14E-14 | 3.14E-11 |
| REACTOME_INTERFERON_ALPHA_BETA_SIGNALING | 73 | Interferon alpha/beta signaling | 7 | 0.0959 | 1.92E-11 | 2.63E-08 |
| REACTOME_CYTOKINE_SIGNALING_IN_IMMUNE_SYSTEM | 719 | Cytokine Signaling in Immune system | 13 | 0.0181 | 4.97E-11 | 4.55E-08 |
| REACTOME_INTERFERON_SIGNALING | 203 | Interferon Signaling | 7 | 0.0345 | 2.57E-08 | 1.76E-05 |
| WP_ADIPOGENESIS | 131 | Adipogenesis | 6 | 0.0458 | 5.03E-08 | 2.76E-05 |
| REACTOME_INTERFERON_GAMMA_SIGNALING | 93 | Interferon gamma signaling | 5 | 0.0538 | 3.15E-07 | 1.44E-04 |
| REACTOME_SIGNALING_BY_INTERLEUKINS | 463 | Signaling by Interleukins | 8 | 0.0173 | 4.93E-07 | 1.53E-04 |
| REACTOME_ENDOSOMAL_VACUOLAR_PATHWAY | 11 | Endosomal/Vacuolar pathway | 3 | 0.2727 | 5.39E-07 | 1.53E-04 |
| HALLMARK_ALLOGRAFT_REJECTION | 200 | Genes up-regulated during transplant rejection. | 6 | 0.03 | 6.12E-07 | 1.53E-04 |
| HALLMARK_KRAS_SIGNALING_UP | 200 | Genes up-regulated by KRAS activation. | 6 | 0.03 | 6.12E-07 | 1.53E-04 |
| HALLMARK_TNFA_SIGNALING_VIA_NFKB | 200 | Genes regulated by NF-kB in response to TNF [GeneID=7124]. | 6 | 0.03 | 6.12E-07 | 1.53E-04 |
| REACTOME_INTERLEUKIN_4_AND_INTERLEUKIN_13_SIGNALING | 111 | Interleukin-4 and Interleukin-13 signaling | 5 | 0.045 | 7.60E-07 | 1.74E-04 |
| WP_HEPATITIS_C_AND_HEPATOCELLULAR_CARCINOMA | 50 | Hepatitis C and Hepatocellular Carcinoma | 4 | 0.08 | 1.04E-06 | 2.19E-04 |
| WP_THE_OVERLAP_BETWEEN_SIGNAL_TRANSDUCTION_PATHWAYS_THAT_CONTRIBUTE_TO_A_RANGE_OF_LMNA_LAMINOPATHIES | 60 | The Overlap Between Signal Transduction Pathways that Contribute to a Range of LMNA Laminopathies | 4 | 0.0667 | 2.17E-06 | 4.26E-04 |
| WP_ONCOSTATIN_M_SIGNALING_PATHWAY | 66 | Oncostatin M Signaling Pathway | 4 | 0.0606 | 3.19E-06 | 5.84E-04 |
| REACTOME_ANTIGEN_PRESENTATION_FOLDING_ASSEMBLY_AND_PEPTIDE_LOADING_OF_CLASS_I_MHC | 25 | Antigen Presentation: Folding, assembly and peptide loading of class I MHC | 3 | 0.12 | 7.40E-06 | 1.27E-03 |
| KEGG_GALACTOSE_METABOLISM | 26 | Galactose metabolism | 3 | 0.1154 | 8.36E-06 | 1.35E-03 |
| WP_PATHWAYS_IN_CLEAR_CELL_RENAL_CELL_CARCINOMA | 87 | Pathways in clear cell renal cell carcinoma | 4 | 0.046 | 9.61E-06 | 1.47E-03 |
| WP_PANCREATIC_ADENOCARCINOMA_PATHWAY | 89 | Pancreatic adenocarcinoma pathway | 4 | 0.0449 | 1.05E-05 | 1.52E-03 |
| HALLMARK_EPITHELIAL_MESENCHYMAL_TRANSITION | 200 | Genes defining epithelial-mesenchymal transition, as in wound healing, fibrosis and metastasis. | 5 | 0.025 | 1.36E-05 | 1.69E-03 |
| HALLMARK_HYPOXIA | 200 | Genes up-regulated in response to low oxygen levels (hypoxia). | 5 | 0.025 | 1.36E-05 | 1.69E-03 |
| HALLMARK_INFLAMMATORY_RESPONSE | 200 | Genes defining inflammatory response. | 5 | 0.025 | 1.36E-05 | 1.69E-03 |
| KEGG_ALLOGRAFT_REJECTION | 37 | Allograft rejection | 3 | 0.0811 | 2.47E-05 | 2.83E-03 |
| WP_TYPE_II_INTERFERON_SIGNALING_IFNG | 37 | Type II interferon signaling (IFNG) | 3 | 0.0811 | 2.47E-05 | 2.83E-03 |
| KEGG_GRAFT_VERSUS_HOST_DISEASE | 41 | Graft-versus-host disease | 3 | 0.0732 | 3.37E-05 | 3.71E-03 |
| KEGG_TYPE_I_DIABETES_MELLITUS | 43 | Type I diabetes mellitus | 3 | 0.0698 | 3.90E-05 | 4.12E-03 |
| REACTOME_INNATE_IMMUNE_SYSTEM | 1117 | Innate Immune System | 9 | 0.0081 | 4.43E-05 | 4.39E-03 |
| WP_INTEGRATED_CANCER_PATHWAY | 45 | Integrated Cancer Pathway | 3 | 0.0667 | 4.47E-05 | 4.39E-03 |
| REACTOME_RUNX3_REGULATES_CDKN1A_TRANSCRIPTION | 7 | RUNX3 regulates CDKN1A transcription | 2 | 0.2857 | 4.71E-05 | 4.46E-03 |
| REACTOME_POST_TRANSLATIONAL_PROTEIN_MODIFICATION | 1435 | Post-translational protein modification | 10 | 0.007 | 5.46E-05 | 5.00E-03 |
| REACTOME_DEGRADATION_OF_THE_EXTRACELLULAR_MATRIX | 140 | Degradation of the extracellular matrix | 4 | 0.0286 | 6.24E-05 | 5.53E-03 |
| KEGG_AUTOIMMUNE_THYROID_DISEASE | 52 | Autoimmune thyroid disease | 3 | 0.0577 | 6.91E-05 | 5.93E-03 |
| WP_IL4_SIGNALING_PATHWAY | 55 | IL-4 Signaling Pathway | 3 | 0.0545 | 8.18E-05 | 6.81E-03 |
| WP_TGFBETA_RECEPTOR_SIGNALING | 58 | TGF-beta Receptor Signaling | 3 | 0.0517 | 9.59E-05 | 7.75E-03 |
| WP_FOCAL_ADHESIONPI3KAKTMTORSIGNALING_PATHWAY | 309 | Focal Adhesion-PI3K-Akt-mTOR-signaling pathway | 5 | 0.0162 | 1.07E-04 | 8.42E-03 |
| WP_TGFBETA_RECEPTOR_SIGNALLING_IN_SKELETAL_DYSPLASIAS | 62 | TGF-beta Receptor Signalling in Skeletal Dysplasias | 3 | 0.0484 | 1.17E-04 | 8.43E-03 |
| WP_LUNG_FIBROSIS | 63 | Lung fibrosis | 3 | 0.0476 | 1.23E-04 | 8.43E-03 |
| WP_NOTCH_SIGNALING_PATHWAY_NETPATH | 63 | Notch Signaling Pathway Netpath | 3 | 0.0476 | 1.23E-04 | 8.43E-03 |
| BIOCARTA_EPONFKB_PATHWAY | 11 | Erythropoietin mediated neuroprotection through NF-kB | 2 | 0.1818 | 1.23E-04 | 8.43E-03 |
| WP_NAD_METABOLISM_SIRTUINS_AND_AGING | 11 | NAD metabolism, sirtuins and aging | 2 | 0.1818 | 1.23E-04 | 8.43E-03 |
| WP_PROTEASOME_DEGRADATION | 64 | Proteasome Degradation | 3 | 0.0469 | 1.29E-04 | 8.62E-03 |
| KEGG_PATHWAYS_IN_CANCER | 325 | Pathways in cancer | 5 | 0.0154 | 1.36E-04 | 8.88E-03 |
| WP_AGERAGE_PATHWAY | 66 | AGE/RAGE pathway | 3 | 0.0455 | 1.41E-04 | 9.00E-03 |
| WP_LNCRNAMEDIATED_MECHANISMS_OF_THERAPEUTIC_RESISTANCE | 12 | LncRNA-mediated mechanisms of therapeutic resistance | 2 | 0.1667 | 1.47E-04 | 9.19E-03 |
| KEGG_VIRAL_MYOCARDITIS | 70 | Viral myocarditis | 3 | 0.0429 | 1.68E-04 | 1.02E-02 |
| WP_CHROMOSOMAL_AND_MICROSATELLITE_INSTABILITY_IN_COLORECTAL_CANCER | 74 | Chromosomal and microsatellite instability in colorectal cancer | 3 | 0.0405 | 1.98E-04 | 1.18E-02 |
| REACTOME_REGULATION_OF_IFNG_SIGNALING | 14 | Regulation of IFNG signaling | 2 | 0.1429 | 2.03E-04 | 1.18E-02 |
| REACTOME_SYNTHESIS_OF_PROSTAGLANDINS_PG_AND_THROMBOXANES_TX | 15 | Synthesis of Prostaglandins (PG) and Thromboxanes (TX) | 2 | 0.1333 | 2.34E-04 | 1.31E-02 |
| WP_PHYTOCHEMICAL_ACTIVITY_ON_NRF2_TRANSCRIPTIONAL_ACTIVATION | 15 | Phytochemical activity on NRF2 transcriptional activation | 2 | 0.1333 | 2.34E-04 | 1.31E-02 |
| HALLMARK_IL2_STAT5_SIGNALING | 199 | Genes up-regulated by STAT5 in response to IL2 stimulation. | 4 | 0.0201 | 2.41E-04 | 1.32E-02 |
| HALLMARK_MTORC1_SIGNALING | 200 | Genes up-regulated through activation of mTORC1 complex. | 4 | 0.02 | 2.46E-04 | 1.32E-02 |
| REACTOME_CLASS_I_MHC_MEDIATED_ANTIGEN_PROCESSING_PRESENTATION | 377 | Class I MHC mediated antigen processing & presentation | 5 | 0.0133 | 2.70E-04 | 1.42E-02 |
| REACTOME_FOXO_MEDIATED_TRANSCRIPTION_OF_CELL_CYCLE_GENES | 17 | FOXO-mediated transcription of cell cycle genes | 2 | 0.1176 | 3.02E-04 | 1.54E-02 |
| REACTOME_IL_6_TYPE_CYTOKINE_RECEPTOR_LIGAND_INTERACTIONS | 17 | IL-6-type cytokine receptor ligand interactions | 2 | 0.1176 | 3.02E-04 | 1.54E-02 |
| KEGG_ANTIGEN_PROCESSING_AND_PRESENTATION | 88 | Antigen processing and presentation | 3 | 0.0341 | 3.30E-04 | 1.65E-02 |
| WP_ALLOGRAFT_REJECTION | 89 | Allograft Rejection | 3 | 0.0337 | 3.41E-04 | 1.67E-02 |
| REACTOME_NICOTINAMIDE_SALVAGING | 19 | Nicotinamide salvaging | 2 | 0.1053 | 3.79E-04 | 1.83E-02 |
| REACTOME_TRANSCRIPTIONAL_REGULATION_BY_RUNX3 | 96 | Transcriptional regulation by RUNX3 | 3 | 0.0312 | 4.26E-04 | 2.02E-02 |
| HALLMARK_INTERFERON_ALPHA_RESPONSE | 97 | Genes up-regulated in response to alpha interferon proteins. | 3 | 0.0309 | 4.39E-04 | 2.05E-02 |
| BIOCARTA_IL6_PATHWAY | 21 | IL 6 signaling pathway | 2 | 0.0952 | 4.64E-04 | 2.06E-02 |
| BIOCARTA_P53HYPOXIA_PATHWAY | 21 | Hypoxia and p53 in the Cardiovascular system | 2 | 0.0952 | 4.64E-04 | 2.06E-02 |
| WP_GLOBO_SPHINGOLIPID_METABOLISM | 21 | Globo Sphingolipid Metabolism | 2 | 0.0952 | 4.64E-04 | 2.06E-02 |
| WP_FGF23_SIGNALLING_IN_HYPOPHOSPHATEMIC_RICKETS_AND_RELATED_DISORDERS | 22 | FGF23 signalling in Hypophosphatemic rickets and related disorders | 2 | 0.0909 | 5.10E-04 | 2.22E-02 |
| REACTOME_DISEASES_OF_METABOLISM | 246 | Diseases of metabolism | 4 | 0.0163 | 5.38E-04 | 2.29E-02 |
| BIOCARTA_PTDINS_PATHWAY | 23 | Phosphoinositides and their downstream targets. | 2 | 0.087 | 5.58E-04 | 2.29E-02 |
| WP_NRF2ARE_REGULATION | 23 | NRF2-ARE regulation | 2 | 0.087 | 5.58E-04 | 2.29E-02 |
| WP_THE_HUMAN_IMMUNE_RESPONSE_TO_TUBERCULOSIS | 23 | The human immune response to tuberculosis | 2 | 0.087 | 5.58E-04 | 2.29E-02 |
| REACTOME_ANTIGEN_PROCESSING_CROSS_PRESENTATION | 106 | Antigen processing-Cross presentation | 3 | 0.0283 | 5.69E-04 | 2.30E-02 |
| WP_SENESCENCE_AND_AUTOPHAGY_IN_CANCER | 108 | Senescence and Autophagy in Cancer | 3 | 0.0278 | 6.01E-04 | 2.32E-02 |
| KEGG_NICOTINATE_AND_NICOTINAMIDE_METABOLISM | 24 | Nicotinate and nicotinamide metabolism | 2 | 0.0833 | 6.08E-04 | 2.32E-02 |
| REACTOME_INTERLEUKIN_6_FAMILY_SIGNALING | 24 | Interleukin-6 family signaling | 2 | 0.0833 | 6.08E-04 | 2.32E-02 |
| REACTOME_OTHER_INTERLEUKIN_SIGNALING | 24 | Other interleukin signaling | 2 | 0.0833 | 6.08E-04 | 2.32E-02 |
| REACTOME_INACTIVATION_OF_CSF3_G_CSF_SIGNALING | 25 | Inactivation of CSF3 (G-CSF) signaling | 2 | 0.08 | 6.61E-04 | 2.49E-02 |
| WP_GASTRIN_SIGNALING_PATHWAY | 114 | Gastrin Signaling Pathway | 3 | 0.0263 | 7.03E-04 | 2.61E-02 |
| REACTOME_REGULATION_OF_IFNA_SIGNALING | 26 | Regulation of IFNA signaling | 2 | 0.0769 | 7.15E-04 | 2.62E-02 |
| KEGG_PENTOSE_PHOSPHATE_PATHWAY | 27 | Pentose phosphate pathway | 2 | 0.0741 | 7.72E-04 | 2.79E-02 |
| REACTOME_NEUTROPHIL_DEGRANULATION | 479 | Neutrophil degranulation | 5 | 0.0104 | 7.98E-04 | 2.85E-02 |
| BIOCARTA_G1_PATHWAY | 28 | Cell Cycle: G1/S Check Point | 2 | 0.0714 | 8.30E-04 | 2.89E-02 |
| WP_EXTRACELLULAR_VESICLES_IN_THE_CROSSTALK_OF_CARDIAC_CELLS | 28 | Extracellular vesicles in the crosstalk of cardiac cells | 2 | 0.0714 | 8.30E-04 | 2.89E-02 |
| WP_CELL_CYCLE | 122 | Cell Cycle | 3 | 0.0246 | 8.56E-04 | 2.91E-02 |
| WP_IL18_SIGNALING_PATHWAY | 279 | IL-18 signaling pathway | 4 | 0.0143 | 8.59E-04 | 2.91E-02 |
| KEGG_CELL_CYCLE | 125 | Cell cycle | 3 | 0.024 | 9.19E-04 | 3.08E-02 |
| REACTOME_SIGNALING_BY_CSF3_G_CSF | 30 | Signaling by CSF3 (G-CSF) | 2 | 0.0667 | 9.53E-04 | 3.12E-02 |
| WP_MATRIX_METALLOPROTEINASES | 30 | Matrix Metalloproteinases | 2 | 0.0667 | 9.53E-04 | 3.12E-02 |
| REACTOME_NICOTINATE_METABOLISM | 31 | Nicotinate metabolism | 2 | 0.0645 | 1.02E-03 | 3.29E-02 |
| WP_EBOLA_VIRUS_PATHWAY_ON_HOST | 131 | Ebola Virus Pathway on Host | 3 | 0.0229 | 1.05E-03 | 3.36E-02 |
| WP_IL17_SIGNALING_PATHWAY | 32 | IL17 signaling pathway | 2 | 0.0625 | 1.08E-03 | 3.39E-02 |
| WP_OVARIAN_INFERTILITY_GENES | 32 | Ovarian Infertility Genes | 2 | 0.0625 | 1.08E-03 | 3.39E-02 |
| KEGG_CELL_ADHESION_MOLECULES_CAMS | 133 | Cell adhesion molecules (CAMs) | 3 | 0.0226 | 1.10E-03 | 3.39E-02 |
| REACTOME_EXTRACELLULAR_MATRIX_ORGANIZATION | 301 | Extracellular matrix organization | 4 | 0.0133 | 1.14E-03 | 3.47E-02 |
| REACTOME_ACTIVATION_OF_MATRIX_METALLOPROTEINASES | 33 | Activation of Matrix Metalloproteinases | 2 | 0.0606 | 1.15E-03 | 3.48E-02 |
| KEGG_NATURAL_KILLER_CELL_MEDIATED_CYTOTOXICITY | 137 | Natural killer cell mediated cytotoxicity | 3 | 0.0219 | 1.20E-03 | 3.54E-02 |
| HALLMARK_COAGULATION | 138 | Genes encoding components of blood coagulation system; also up-regulated in platelets. | 3 | 0.0217 | 1.22E-03 | 3.54E-02 |
| KEGG_FRUCTOSE_AND_MANNOSE_METABOLISM | 34 | Fructose and mannose metabolism | 2 | 0.0588 | 1.22E-03 | 3.54E-02 |
| REACTOME_METABOLIC_DISORDERS_OF_BIOLOGICAL_OXIDATION_ENZYMES | 34 | Metabolic disorders of biological oxidation enzymes | 2 | 0.0588 | 1.22E-03 | 3.54E-02 |
| HALLMARK_ANGIOGENESIS | 36 | Genes up-regulated during formation of blood vessels (angiogenesis). | 2 | 0.0556 | 1.37E-03 | 3.81E-02 |
| REACTOME_INTERLEUKIN_7_SIGNALING | 36 | Interleukin-7 signaling | 2 | 0.0556 | 1.37E-03 | 3.81E-02 |

| | | | | | | |
|---|---|---|---|---|---|---|
| WP_HOSTPATHOGEN_INTERACTION_OF_HUMAN_CORONA_VIRUSES_MAPK_SIGNALING | 36 | Host-pathogen interaction of human corona viruses - MAPK signaling | 2 | 0.0556 | 1.37E-03 | 3.81E-02 |
| WP_TYPE_2_PAPILLARY_RENAL_CELL_CARCINOMA | 36 | Type 2 papillary renal cell carcinoma | 2 | 0.0556 | 1.37E-03 | 3.81E-02 |
| WP_NRF2_PATHWAY | 145 | NRF2 pathway | 3 | 0.0207 | 1.41E-03 | 3.87E-02 |

Enrichment for genes upregulated in AKI vs. control

Overlap Results

| | |
|---|---|
| Collection(s): | CP:BIOCARTA, CP:KEGG, CP:REACTOME, CP:WIKIPATHWAYS, H |
| # overlaps shown: | 2 |
| # genesets in collections: | 2747 |
| # genes in comparison (n): | 13 |
| # genes in universe (N): | 40312 |

| Gene Set Name | # Genes in Gene Set (K) | Description | # Genes in Overlap (k) | k/K | p-value | FDR q-value |
|---|---|---|---|---|---|---|
| REACTOME_DEGRADATION_OF_THE_EXTRACELLULAR_MATRIX | 140 | Degradation of the extracellular matrix | 3 | 0.0214 | 1.14E-05 | 3.14E-02 |
| WP_EXTRACELLULAR_VESICLES_IN_THE_CROSSTALK_OF_CARDIAC_CELLS | 28 | Extracellular vesicles in the crosstalk of cardiac cells | 2 | 0.0714 | 3.61E-05 | 4.96E-02 |

Enrichment for genes upregulated in AKI vs. control

Overlap Results

| | |
|---|---|
| Collection(s): | CP:BIOCARTA, CP:KEGG, CP:REACTOME, CP:WIKIPATHWAYS, H |
| # overlaps shown: | 5 |
| # genesets in collections: | 2747 |
| # genes in comparison (n): | 21 |
| # genes in universe (N): | 40312 |

| Gene Set Name | # Genes in Gene Set (K) | Description | # Genes in Overlap (k) | k/K | p-value | FDR q-value |
|---|---|---|---|---|---|---|
| HALLMARK_HYPOXIA | 200 | Genes up-regulated in response to low oxygen levels (hypoxia). | 5 | 0.025 | 5.45E-08 | 1.50E-04 |
| WP_VITAMIN_D_RECEPTOR_PATHWAY | 185 | Vitamin D Receptor Pathway | 4 | 0.0216 | 2.42E-06 | 3.02E-03 |
| HALLMARK_MTORC1_SIGNALING | 200 | Genes up-regulated through activation of mTORC1 complex. | 4 | 0.02 | 3.29E-06 | 3.02E-03 |
| WP_ADIPOGENESIS | 131 | Adipogenesis | 3 | 0.0229 | 4.27E-05 | 2.93E-02 |
| WP_FGF23_SIGNALLING_IN_HYPOPHOSPHATEMIC_RICKETS_AND_RELATED_DISORDERS | 22 | FGF23 signalling in Hypophosphatemic rickets and related disorders | 2 | 0.0909 | 5.93E-05 | 3.26E-02 |

Enrichment for genes upregulated in AKI vs. control

Overlap Results

| | |
|---|---|
| Collection(s): | CP:BIOCARTA, CP:KEGG, CP:REACTOME, CP:WIKIPATHWAYS, H |
| # overlaps shown: | 1 |
| # genesets in collections: | 2747 |
| # genes in comparison (n): | 24 |
| # genes in universe (N): | 40312 |

| Gene Set Name | # Genes in Gene Set (K) | Description | # Genes in Overlap (k) | k/K | p-value | FDR q-value |
|---|---|---|---|---|---|---|
| WP_GLUCOCORTICOID_RECEPTOR_PATHWAY | 71 | Glucocorticoid Receptor Pathway | 3 | 0.0423 | 1.03E-05 | 2.83E-02 |

Gene/Gene Set Overlap Matrix

Enrichment for genes upregulated in AKI vs. control

Overlap Results

| | |
|---|---|
| Collection(s): | CP:BIOCARTA, CP:KEGG, CP:REACTOME, CP:WIKIPATHWAYS, H |
| # overlaps shown: | 100 |
| # genesets in collections: | 2747 |
| # genes in comparison (n): | 872 |
| # genes in universe (N): | 40312 |

| Gene Set Name | # Genes in Gene Set (K) | Description | # Genes in Overlap (k) | k/K | p-value | FDR q-value |
|---|---|---|---|---|---|---|
| REACTOME_INNATE_IMMUNE_SYSTEM | 1117 | Innate Immune System | 105 | 0.094 | 5.54E-37 | 1.52E-33 |
| HALLMARK_TNFA_SIGNALING_VIA_NFKB | 200 | Genes regulated by NF-kB in response to TNF [GeneID=7124]. | 43 | 0.215 | 4.17E-30 | 5.73E-27 |
| REACTOME_NEUTROPHIL_DEGRANULATION | 479 | Neutrophil degranulation | 59 | 0.1232 | 4.19E-27 | 3.84E-24 |
| REACTOME_CYTOKINE_SIGNALING_IN_IMMUNE_SYSTEM | 719 | Cytokine Signaling in Immune system | 69 | 0.096 | 5.22E-25 | 3.59E-22 |
| HALLMARK_HYPOXIA | 200 | Genes up-regulated in response to low oxygen levels (hypoxia). | 33 | 0.165 | 1.25E-19 | 5.71E-17 |
| HALLMARK_P53_PATHWAY | 200 | Genes involved in p53 pathways and networks. | 33 | 0.165 | 1.25E-19 | 5.71E-17 |
| WP_IL18_SIGNALING_PATHWAY | 279 | IL-18 signaling pathway | 37 | 0.1326 | 1.54E-18 | 6.03E-16 |
| REACTOME_SIGNALING_BY_INTERLEUKINS | 463 | Signaling by Interleukins | 47 | 0.1015 | 2.29E-18 | 7.86E-16 |
| HALLMARK_INFLAMMATORY_RESPONSE | 200 | Genes defining inflammatory response. | 30 | 0.15 | 8.74E-17 | 2.40E-14 |
| HALLMARK_INTERFERON_GAMMA_RESPONSE | 200 | Genes up-regulated in response to IFNG [GeneID=3458]. | 30 | 0.15 | 8.74E-17 | 2.40E-14 |
| HALLMARK_COMPLEMENT | 200 | Genes encoding components of the complement system, which is part of the innate immune system. | 28 | 0.14 | 5.67E-15 | 1.42E-12 |
| REACTOME_HEMOSTASIS | 678 | Hemostasis | 51 | 0.0752 | 1.96E-14 | 4.48E-12 |
| REACTOME_DEVELOPMENTAL_BIOLOGY | 1143 | Developmental Biology | 69 | 0.0604 | 2.78E-14 | 5.88E-12 |
| REACTOME_CELLULAR_RESPONSES_TO_EXTERNAL_STIMULI | 706 | Cellular responses to external stimuli | 51 | 0.0722 | 9.42E-14 | 1.85E-11 |
| HALLMARK_GLYCOLYSIS | 200 | Genes encoding proteins involved in glycolysis and gluconeogenesis. | 26 | 0.13 | 3.11E-13 | 5.69E-11 |
| HALLMARK_IL6_JAK_STAT3_SIGNALING | 87 | Genes up-regulated by IL6 [GeneID=3569] via STAT3 [GeneID=6774], e.g., during acute phase response. | 18 | 0.2069 | 4.30E-13 | 7.38E-11 |
| REACTOME_NERVOUS_SYSTEM_DEVELOPMENT | 580 | Nervous system development | 44 | 0.0759 | 9.35E-13 | 1.51E-10 |
| WP_HUMAN_COMPLEMENT_SYSTEM | 99 | Human Complement System | 18 | 0.1818 | 4.41E-12 | 6.73E-10 |
| WP_VEGFAVEGFR2_SIGNALING_PATHWAY | 438 | VEGFA-VEGFR2 Signaling Pathway | 36 | 0.0822 | 1.16E-11 | 1.68E-09 |
| HALLMARK_COAGULATION | 138 | Genes encoding components of blood coagulation system; also up-regulated in platelets. | 20 | 0.1449 | 2.24E-11 | 3.08E-09 |
| REACTOME_TOLL_LIKE_RECEPTOR_CASCADES | 155 | Toll-like Receptor Cascades | 21 | 0.1355 | 2.65E-11 | 3.46E-09 |
| HALLMARK_EPITHELIAL_MESENCHYMAL_TRANSITION | 200 | Genes defining epithelial-mesenchymal transition, as in wound healing, fibrosis and metastasis. | 23 | 0.115 | 9.05E-11 | 1.13E-08 |
| REACTOME_EXTRACELLULAR_MATRIX_ORGANIZATION | 301 | Extracellular matrix organization | 28 | 0.093 | 1.34E-10 | 1.60E-08 |
| REACTOME_ADAPTIVE_IMMUNE_SYSTEM | 825 | Adaptive Immune System | 49 | 0.0594 | 2.94E-10 | 3.37E-08 |
| REACTOME_METABOLISM_OF_CARBOHYDRATES | 293 | Metabolism of carbohydrates | 27 | 0.0922 | 3.53E-10 | 3.88E-08 |
| HALLMARK_APOPTOSIS | 161 | Genes mediating programmed cell death (apoptosis) by activation of caspases. | 20 | 0.1242 | 3.81E-10 | 4.02E-08 |
| HALLMARK_MTORC1_SIGNALING | 200 | Genes up-regulated through activation of mTORC1 complex. | 22 | 0.11 | 5.45E-10 | 5.54E-08 |
| REACTOME_INTERLEUKIN_1_FAMILY_SIGNALING | 140 | Interleukin-1 family signaling | 18 | 0.1286 | 1.63E-09 | 1.60E-07 |
| REACTOME_POST_TRANSLATIONAL_PROTEIN_MODIFICATION | 1435 | Post-translational protein modification | 68 | 0.0474 | 1.81E-09 | 1.71E-07 |
| HALLMARK_IL2_STAT5_SIGNALING | 199 | Genes up-regulated by STAT5 in response to IL2 stimulation. | 21 | 0.1055 | 2.85E-09 | 2.61E-07 |
| HALLMARK_ALLOGRAFT_REJECTION | 200 | Genes up-regulated during transplant rejection. | 21 | 0.105 | 3.12E-09 | 2.77E-07 |
| REACTOME_INFECTIOUS_DISEASE | 924 | Infectious disease | 50 | 0.0541 | 4.34E-09 | 3.72E-07 |
| REACTOME_RESPONSE_OF_EIF2AK4_GCN2_TO_AMINO_ACID_DEFICIENCY | 102 | Response of EIF2AK4 (GCN2) to amino acid deficiency | 15 | 0.1471 | 5.73E-09 | 4.77E-07 |
| REACTOME_TOLL_LIKE_RECEPTOR_TLR1_TLR2_CASCADE | 103 | Toll Like Receptor TLR1:TLR2 Cascade | 15 | 0.1456 | 6.58E-09 | 5.32E-07 |
| REACTOME_DISEASES_OF_IMMUNE_SYSTEM | 31 | Diseases of Immune System | 9 | 0.2903 | 1.31E-08 | 1.03E-06 |
| HALLMARK_ESTROGEN_RESPONSE_EARLY | 200 | Genes defining early response to estrogen. | 20 | 0.1 | 1.70E-08 | 1.30E-06 |
| REACTOME_CLASS_I_MHC_MEDIATED_ANTIGEN_PROCESSING_PRESENTATION | 377 | Class I MHC mediated antigen processing & presentation | 28 | 0.0743 | 2.03E-08 | 1.51E-06 |
| KEGG_CYTOKINE_CYTOKINE_RECEPTOR_INTERACTION | 265 | Cytokine-cytokine receptor interaction | 23 | 0.0868 | 2.18E-08 | 1.57E-06 |
| REACTOME_RNA_POLYMERASE_II_TRANSCRIPTION | 1374 | RNA Polymerase II Transcription | 63 | 0.0459 | 2.47E-08 | 1.72E-06 |
| WP_VITAMIN_D_RECEPTOR_PATHWAY | 185 | Vitamin D Receptor Pathway | 19 | 0.1027 | 2.50E-08 | 1.72E-06 |
| WP_INTEGRATED_CANCER_PATHWAY | 45 | Integrated Cancer Pathway | 10 | 0.2222 | 3.43E-08 | 2.30E-06 |
| REACTOME_TRANSCRIPTIONAL_REGULATION_BY_TP53 | 363 | Transcriptional Regulation by TP53 | 27 | 0.0744 | 3.52E-08 | 2.30E-06 |
| WP_PATHWAYS_IN_CLEAR_CELL_RENAL_CELL_CARCINOMA | 87 | Pathways in clear cell renal cell carcinoma | 13 | 0.1494 | 4.87E-08 | 3.11E-06 |
| WP_FOLATE_METABOLISM | 73 | Folate Metabolism | 12 | 0.1644 | 5.34E-08 | 3.34E-06 |
| REACTOME_INTEGRIN_SIGNALING | 27 | Integrin signaling | 8 | 0.2963 | 7.16E-08 | 4.37E-06 |
| REACTOME_PLATELET_ACTIVATION_SIGNALING_AND_AGGREGATION | 261 | Platelet activation, signaling and aggregation | 22 | 0.0843 | 7.37E-08 | 4.40E-06 |
| WP_SELENIUM_MICRONUTRIENT_NETWORK | 91 | Selenium Micronutrient Network | 13 | 0.1429 | 8.43E-08 | 4.92E-06 |
| WP_FOCAL_ADHESIONPI3KAKTMTORSIGNALING_PATHWAY | 309 | Focal Adhesion-PI3K-Akt-mTOR-signaling pathway | 24 | 0.0777 | 8.98E-08 | 5.14E-06 |
| REACTOME_EPH_EPHRIN_SIGNALING | 92 | EPH-Ephrin signaling | 13 | 0.1413 | 9.62E-08 | 5.39E-06 |
| REACTOME_TRANSPORT_OF_SMALL_MOLECULES | 728 | Transport of small molecules | 40 | 0.0549 | 1.04E-07 | 5.69E-06 |
| WP_ONCOSTATIN_M_SIGNALING_PATHWAY | 66 | Oncostatin M Signaling Pathway | 11 | 0.1667 | 1.66E-07 | 8.92E-06 |
| WP_ADIPOGENESIS | 131 | Adipogenesis | 15 | 0.1145 | 1.76E-07 | 9.28E-06 |
| REACTOME_MYD88_INDEPENDENT_TLR4_CASCADE | 97 | MyD88-independent TLR4 cascade | 13 | 0.134 | 1.82E-07 | 9.43E-06 |
| REACTOME_REGULATION_OF_TLR_BY_ENDOGENOUS_LIGAND | 21 | Regulation of TLR by endogenous ligand | 7 | 0.3333 | 1.93E-07 | 9.82E-06 |
| KEGG_P53_SIGNALING_PATHWAY | 68 | p53 signaling pathway | 11 | 0.1618 | 2.27E-07 | 1.14E-05 |
| REACTOME_REGULATION_OF_IFNG_SIGNALING | 14 | Regulation of IFNG signaling | 6 | 0.4286 | 2.61E-07 | 1.28E-05 |
| KEGG_COMPLEMENT_AND_COAGULATION_CASCADES | 69 | Complement and coagulation cascades | 11 | 0.1594 | 2.65E-07 | 1.28E-05 |
| WP_APOPTOSIS | 86 | Apoptosis | 12 | 0.1395 | 3.44E-07 | 1.63E-05 |
| REACTOME_EUKARYOTIC_TRANSLATION_INITIATION | 120 | Eukaryotic Translation Initiation | 14 | 0.1167 | 3.56E-07 | 1.66E-05 |
| WP_P53_TRANSCRIPTIONAL_GENE_NETWORK | 72 | p53 transcriptional gene network | 11 | 0.1528 | 4.14E-07 | 1.89E-05 |
| REACTOME_DEGRADATION_OF_THE_EXTRACELLULAR_MATRIX | 140 | Degradation of the extracellular matrix | 15 | 0.1071 | 4.20E-07 | 1.89E-05 |
| REACTOME_SIGNALING_BY_RHO_GTPASES_MIRO_GTPASES_AND_RHOBTB3 | 717 | Signaling by Rho GTPases, Miro GTPases and RHOBTB3 | 38 | 0.053 | 5.24E-07 | 2.32E-05 |
| BIOCARTA_G2_PATHWAY | 24 | Cell Cycle: G2/M Checkpoint | 7 | 0.2917 | 5.43E-07 | 2.37E-05 |
| WP_PROLACTIN_SIGNALING_PATHWAY | 76 | Prolactin Signaling Pathway | 11 | 0.1447 | 7.26E-07 | 3.12E-05 |
| REACTOME_METABOLISM_OF_RNA | 672 | Metabolism of RNA | 36 | 0.0536 | 8.10E-07 | 3.39E-05 |
| WP_APOPTOSIS_MODULATION_AND_SIGNALING | 93 | Apoptosis Modulation and Signaling | 12 | 0.129 | 8.16E-07 | 3.39E-05 |
| KEGG_AXON_GUIDANCE | 129 | Axon guidance | 14 | 0.1085 | 8.68E-07 | 3.56E-05 |
| REACTOME_INTERLEUKIN_4_AND_INTERLEUKIN_13_SIGNALING | 111 | Interleukin-4 and Interleukin-13 signaling | 13 | 0.1171 | 8.88E-07 | 3.59E-05 |
| REACTOME_EUKARYOTIC_TRANSLATION_ELONGATION | 94 | Eukaryotic Translation Elongation | 12 | 0.1277 | 9.17E-07 | 3.60E-05 |
| WP_TNF_ALPHA_SIGNALING_PATHWAY | 94 | TNF alpha Signaling Pathway | 12 | 0.1277 | 9.17E-07 | 3.60E-05 |
| REACTOME_SIGNALING_BY_MET | 79 | Signaling by MET | 11 | 0.1392 | 1.08E-06 | 4.12E-05 |
| WP_REGULATORY_CIRCUITS_OF_THE_STAT3_SIGNALING_PATHWAY | 79 | Regulatory circuits of the STAT3 signaling pathway | 11 | 0.1392 | 1.08E-06 | 4.12E-05 |
| WP_GASTRIN_SIGNALING_PATHWAY | 114 | Gastrin Signaling Pathway | 13 | 0.114 | 1.21E-06 | 4.54E-05 |
| REACTOME_ANTIMICROBIAL_PEPTIDES | 97 | Antimicrobial peptides | 12 | 0.1237 | 1.29E-06 | 4.79E-05 |
| REACTOME_IRAK4_DEFICIENCY_TLR2_4 | 18 | IRAK4 deficiency (TLR2/4) | 6 | 0.3333 | 1.50E-06 | 5.48E-05 |
| REACTOME_FORMATION_OF_FIBRIN_CLOT_CLOTTING_CASCADE | 39 | Formation of Fibrin Clot (Clotting Cascade) | 8 | 0.2051 | 1.58E-06 | 5.63E-05 |
| REACTOME_PLATELET_AGGREGATION_PLUG_FORMATION | 39 | Platelet Aggregation (Plug Formation) | 8 | 0.2051 | 1.58E-06 | 5.63E-05 |
| REACTOME_SELENOAMINO_ACID_METABOLISM | 118 | Selenoamino acid metabolism | 13 | 0.1102 | 1.79E-06 | 6.31E-05 |
| REACTOME_CELLULAR_RESPONSE_TO_STARVATION | 157 | Cellular response to starvation | 15 | 0.0955 | 1.82E-06 | 6.34E-05 |
| WP_VITAMIN_B12_METABOLISM | 53 | Vitamin B12 Metabolism | 9 | 0.1698 | 1.88E-06 | 6.46E-05 |
| HALLMARK_APICAL_JUNCTION | 200 | Genes encoding components of apical junction complex. | 17 | 0.085 | 1.95E-06 | 6.54E-05 |
| HALLMARK_ESTROGEN_RESPONSE_LATE | 200 | Genes defining late response to estrogen. | 17 | 0.085 | 1.95E-06 | 6.54E-05 |
| REACTOME_INTERLEUKIN_1_SIGNALING | 102 | Interleukin-1 signaling | 12 | 0.1176 | 2.22E-06 | 7.33E-05 |
| WP_DNA_DAMAGE_RESPONSE | 69 | DNA Damage Response | 10 | 0.1449 | 2.29E-06 | 7.41E-05 |
| REACTOME_TRANSLATION | 295 | Translation | 21 | 0.0712 | 2.29E-06 | 7.41E-05 |
| WP_PI3KAKT_SIGNALING_PATHWAY | 345 | PI3K-Akt Signaling Pathway | 23 | 0.0667 | 2.35E-06 | 7.47E-05 |
| REACTOME_SIGNALING_BY_RECEPTOR_TYROSINE_KINASES | 504 | Signaling by Receptor Tyrosine Kinases | 29 | 0.0575 | 2.37E-06 | 7.47E-05 |
| WP_INSULIN_SIGNALING | 161 | Insulin Signaling | 15 | 0.0932 | 2.50E-06 | 7.80E-05 |
| WP_IL4_SIGNALING_PATHWAY | 55 | IL-4 Signaling Pathway | 9 | 0.1636 | 2.60E-06 | 8.01E-05 |
| KEGG_APOPTOSIS | 87 | Apoptosis | 11 | 0.1264 | 2.86E-06 | 8.73E-05 |
| REACTOME_METAL_SEQUESTRATION_BY_ANTIMICROBIAL_PROTEINS | 6 | Metal sequestration by antimicrobial proteins | 4 | 0.6667 | 3.15E-06 | 9.51E-05 |
| KEGG_RIBOSOME | 88 | Ribosome | 11 | 0.125 | 3.21E-06 | 9.57E-05 |
| REACTOME_PROGRAMMED_CELL_DEATH | 208 | Programmed Cell Death | 17 | 0.0817 | 3.33E-06 | 9.84E-05 |
| WP_REGULATION_OF_TOLLLIKE_RECEPTOR_SIGNALING_PATHWAY | 145 | Regulation of toll-like receptor signaling pathway | 14 | 0.0966 | 3.53E-06 | 1.02E-04 |
| REACTOME_INTERLEUKIN_12_FAMILY_SIGNALING | 57 | Interleukin-12 family signaling | 9 | 0.1579 | 3.53E-06 | 1.02E-04 |
| WP_SENESCENCE_AND_AUTOPHAGY_IN_CANCER | 108 | Senescence and Autophagy in Cancer | 12 | 0.1111 | 4.06E-06 | 1.16E-04 |
| WP_MONOAMINE_TRANSPORT | 32 | Monoamine Transport | 7 | 0.2188 | 4.54E-06 | 1.29E-04 |

Bulk RNA-seq UP

| | | | | | | |
|---|---|---|---|---|---|---|
| REACTOME_COMMON_PATHWAY_OF_FIBRIN_CLOT_FORMATION | 22 | Common Pathway of Fibrin Clot Formation | 6 | 0.2727 | 5.59E-06 | 1.57E-04 |
| REACTOME_CELL_SURFACE_INTERACTIONS_AT_THE_VASCULAR_WALL | 194 | Cell surface interactions at the vascular wall | 16 | 0.0825 | 5.72E-06 | 1.59E-04 |
| REACTOME_SIGNALING_BY_ROBO_RECEPTORS | 218 | Signaling by ROBO receptors | 17 | 0.078 | 6.25E-06 | 1.72E-04 |

Enrichment for genes downregulated in AKI vs. control

Overlap Results

| | |
|---|---|
| Collection(s): | CP:BIOCARTA, CP:KEGG, CP:REACTOME, CP:WIKIPATHWAYS, H |
| # overlaps shown: | 95 |
| # genesets in collections: | 2747 |
| # genes in comparison (n): | 767 |
| # genes in universe (N): | 40312 |

| Gene Set Name | # Genes in Gene Set (K) | Description | # Genes in Overlap (k) | k/K | p-value |
|---|---|---|---|---|---|
| REACTOME_METABOLISM_OF_AMINO_ACIDS_AND_DERIVATIVES | 374 | Metabolism of amino acids and derivatives | 38 | 0.1016 | 6.65E-17 |
| REACTOME_TRANSPORT_OF_SMALL_MOLECULES | 728 | Transport of small molecules | 52 | 0.0714 | 4.97E-16 |
| KEGG_VALINE_LEUCINE_AND_ISOLEUCINE_DEGRADATION | 44 | Valine, leucine and isoleucine degradation | 14 | 0.3182 | 4.92E-14 |
| REACTOME_SLC_MEDIATED_TRANSMEMBRANE_TRANSPORT | 250 | SLC-mediated transmembrane transport | 26 | 0.104 | 3.15E-12 |
| REACTOME_TRANSPORT_OF_INORGANIC_CATIONS_ANIONS_AND_AMINO_ACIDS_OLIGOPEPTIDES | 106 | Transport of inorganic cations/anions and amino acids/oligopeptides | 17 | 0.1604 | 1.90E-11 |
| KEGG_GLYCINE_SERINE_AND_THREONINE_METABOLISM | 31 | Glycine, serine and threonine metabolism | 10 | 0.3226 | 1.81E-10 |
| REACTOME_GLYOXYLATE_METABOLISM_AND_GLYCINE_DEGRADATION | 31 | Glyoxylate metabolism and glycine degradation | 10 | 0.3226 | 1.81E-10 |
| HALLMARK_XENOBIOTIC_METABOLISM | 200 | Genes encoding proteins involved in processing of drugs and other xenobiotics. | 21 | 0.105 | 3.20E-10 |
| KEGG_ARGININE_AND_PROLINE_METABOLISM | 54 | Arginine and proline metabolism | 12 | 0.2222 | 3.41E-10 |
| WP_AMINO_ACID_METABOLISM | 91 | Amino Acid metabolism | 14 | 0.1538 | 2.00E-09 |
| KEGG_PEROXISOME | 78 | Peroxisome | 13 | 0.1667 | 2.74E-09 |
| KEGG_BETA_ALANINE_METABOLISM | 22 | beta-Alanine metabolism | 8 | 0.3636 | 4.18E-09 |
| REACTOME_METABOLISM_OF_LIPIDS | 741 | Metabolism of lipids | 40 | 0.054 | 5.23E-09 |
| KEGG_PROPANOATE_METABOLISM | 33 | Propanoate metabolism | 9 | 0.2727 | 7.99E-09 |
| REACTOME_FATTY_ACID_METABOLISM | 177 | Fatty acid metabolism | 18 | 0.1017 | 9.72E-09 |
| HALLMARK_FATTY_ACID_METABOLISM | 158 | Genes encoding proteins involved in metabolism of fatty acids. | 17 | 0.1076 | 1.07E-08 |
| KEGG_TRYPTOPHAN_METABOLISM | 40 | Tryptophan metabolism | 9 | 0.225 | 5.03E-08 |
| REACTOME_BRANCHED_CHAIN_AMINO_ACID_CATABOLISM | 21 | Branched-chain amino acid catabolism | 7 | 0.3333 | 8.10E-08 |
| REACTOME_PROTEIN_LOCALIZATION | 164 | Protein localization | 16 | 0.0976 | 1.14E-07 |
| WP_TRANSSULFURATION_AND_ONE_CARBON_METABOLISM | 32 | Trans-sulfuration and one carbon metabolism | 8 | 0.25 | 1.16E-07 |
| REACTOME_PEROXISOMAL_PROTEIN_IMPORT | 63 | Peroxisomal protein import | 10 | 0.1587 | 3.02E-07 |
| REACTOME_BIOLOGICAL_OXIDATIONS | 222 | Biological oxidations | 18 | 0.0811 | 3.12E-07 |
| WP_PROXIMAL_TUBULE_TRANSPORT | 58 | Proximal tubule transport | 9 | 0.1552 | 1.44E-06 |
| WP_ONE_CARBON_METABOLISM | 31 | One Carbon Metabolism | 7 | 0.2258 | 1.55E-06 |
| REACTOME_CHOLINE_CATABOLISM | 6 | Choline catabolism | 4 | 0.6667 | 1.89E-06 |
| REACTOME_SLC_TRANSPORTER_DISORDERS | 99 | SLC transporter disorders | 11 | 0.1111 | 3.04E-06 |
| REACTOME_METABOLISM_OF_WATER_SOLUBLE_VITAMINS_AND_COFACTORS | 123 | Metabolism of water-soluble vitamins and cofactors | 12 | 0.0976 | 4.33E-06 |
| REACTOME_DISORDERS_OF_TRANSMEMBRANE_TRANSPORTERS | 176 | Disorders of transmembrane transporters | 14 | 0.0795 | 7.86E-06 |
| KEGG_DRUG_METABOLISM_CYTOCHROME_P450 | 72 | Drug metabolism - cytochrome P450 | 9 | 0.125 | 9.11E-06 |
| WP_TRYPTOPHAN_METABOLISM | 42 | Tryptophan metabolism | 7 | 0.1667 | 1.33E-05 |
| WP_METAPATHWAY_BIOTRANSFORMATION_PHASE_I_AND_II | 185 | Metapathway biotransformation Phase I and II | 14 | 0.0757 | 1.39E-05 |
| KEGG_HISTIDINE_METABOLISM | 29 | Histidine metabolism | 6 | 0.2069 | 1.52E-05 |
| REACTOME_NEURONAL_SYSTEM | 410 | Neuronal System | 22 | 0.0537 | 1.59E-05 |
| REACTOME_TRANSMISSION_ACROSS_CHEMICAL_SYNAPSES | 269 | Transmission across Chemical Synapses | 17 | 0.0632 | 1.87E-05 |
| KEGG_GLYCOLYSIS_GLUCONEOGENESIS | 62 | Glycolysis / Gluconeogenesis | 8 | 0.129 | 2.26E-05 |
| KEGG_CITRATE_CYCLE_TCA_CYCLE | 31 | Citrate cycle (TCA cycle) | 6 | 0.1935 | 2.28E-05 |
| KEGG_ALANINE_ASPARTATE_AND_GLUTAMATE_METABOLISM | 32 | Alanine, aspartate and glutamate metabolism | 6 | 0.1875 | 2.76E-05 |
| HALLMARK_ADIPOGENESIS | 200 | Genes up-regulated during adipocyte differentiation (adipogenesis). | 14 | 0.07 | 3.31E-05 |
| REACTOME_CARDIAC_CONDUCTION | 127 | Cardiac conduction | 11 | 0.0866 | 3.32E-05 |
| KEGG_GLYCEROLIPID_METABOLISM | 49 | Glycerolipid metabolism | 7 | 0.1429 | 3.77E-05 |
| WP_UREA_CYCLE_AND_METABOLISM_OF_AMINO_GROUPS | 21 | Urea cycle and metabolism of amino groups | 5 | 0.2381 | 3.89E-05 |
| KEGG_BUTANOATE_METABOLISM | 34 | Butanoate metabolism | 6 | 0.1765 | 3.97E-05 |
| REACTOME_ION_CHANNEL_TRANSPORT | 183 | Ion channel transport | 13 | 0.071 | 5.38E-05 |
| WP_ALANINE_AND_ASPARTATE_METABOLISM | 12 | Alanine and aspartate metabolism | 4 | 0.3333 | 5.70E-05 |
| WP_BIOMARKERS_FOR_UREA_CYCLE_DISORDERS | 12 | Biomarkers for urea cycle disorders | 4 | 0.3333 | 5.70E-05 |
| HALLMARK_BILE_ACID_METABOLISM | 112 | Genes involve in metabolism of bile acids and salts. | 10 | 0.0893 | 5.80E-05 |
| KEGG_PROXIMAL_TUBULE_BICARBONATE_RECLAMATION | 23 | Proximal tubule bicarbonate reclamation | 5 | 0.2174 | 6.23E-05 |
| REACTOME_INTRINSIC_PATHWAY_OF_FIBRIN_CLOT_FORMATION | 23 | Intrinsic Pathway of Fibrin Clot Formation | 5 | 0.2174 | 6.23E-05 |
| REACTOME_MITOCHONDRIAL_FATTY_ACID_BETA_OXIDATION | 37 | Mitochondrial Fatty Acid Beta-Oxidation | 6 | 0.1622 | 6.54E-05 |
| REACTOME_FRUCTOSE_CATABOLISM | 5 | Fructose catabolism | 3 | 0.6 | 6.67E-05 |
| WP_CATALYTIC_CYCLE_OF_MAMMALIAN_FLAVINCONTAINING_MONOOXYGENASES_FMOS | 5 | Catalytic cycle of mammalian Flavin-containing MonoOxygenases (FMOs) | 3 | 0.6 | 6.67E-05 |
| REACTOME_ION_HOMEOSTASIS | 54 | Ion homeostasis | 7 | 0.1296 | 7.15E-05 |
| WP_ONE_CARBON_METABOLISM_AND_RELATED_PATHWAYS | 54 | One carbon metabolism and related pathways | 7 | 0.1296 | 7.15E-05 |
| REACTOME_METABOLISM_OF_VITAMINS_AND_COFACTORS | 189 | Metabolism of vitamins and cofactors | 13 | 0.0688 | 7.49E-05 |
| REACTOME_FORMATION_OF_FIBRIN_CLOT_CLOTTING_CASCADE | 39 | Formation of Fibrin Clot (Clotting Cascade) | 6 | 0.1538 | 8.89E-05 |
| KEGG_PYRUVATE_METABOLISM | 40 | Pyruvate metabolism | 6 | 0.15 | 1.03E-04 |
| KEGG_FATTY_ACID_METABOLISM | 42 | Fatty acid metabolism | 6 | 0.1429 | 1.36E-04 |
| WP_NUCLEAR_RECEPTORS_METAPATHWAY | 321 | Nuclear Receptors Meta-Pathway | 17 | 0.053 | 1.66E-04 |
| KEGG_CALCIUM_SIGNALING_PATHWAY | 178 | Calcium signaling pathway | 12 | 0.0674 | 1.69E-04 |
| KEGG_LYSINE_DEGRADATION | 44 | Lysine degradation | 6 | 0.1364 | 1.77E-04 |
| WP_BIOGENIC_AMINE_SYNTHESIS | 16 | Biogenic Amine Synthesis | 4 | 0.25 | 1.97E-04 |
| REACTOME_PHASE_I_FUNCTIONALIZATION_OF_COMPOUNDS | 106 | Phase I - Functionalization of compounds | 9 | 0.0849 | 1.98E-04 |
| REACTOME_PEROXISOMAL_LIPID_METABOLISM | 29 | Peroxisomal lipid metabolism | 5 | 0.1724 | 2.00E-04 |
| KEGG_RETINOL_METABOLISM | 64 | Retinol metabolism | 7 | 0.1094 | 2.13E-04 |
| REACTOME_CATION_COUPLED_CHLORIDE_COTRANSPORTERS | 7 | Cation-coupled Chloride cotransporters | 3 | 0.4286 | 2.27E-04 |
| REACTOME_FRUCTOSE_METABOLISM | 7 | Fructose metabolism | 3 | 0.4286 | 2.27E-04 |
| WP_DISORDERS_OF_THE_KREBS_CYCLE | 7 | Disorders of the Krebs cycle | 3 | 0.4286 | 2.27E-04 |
| WP_OXYSTEROLS_DERIVED_FROM_CHOLESTEROL | 46 | Oxysterols derived from cholesterol | 6 | 0.1304 | 2.28E-04 |
| WP_TCA_CYCLE_AKA_KREBS_OR_CITRIC_ACID_CYCLE | 18 | TCA Cycle (aka Krebs or citric acid cycle) | 4 | 0.2222 | 3.22E-04 |
| KEGG_METABOLISM_OF_XENOBIOTICS_BY_CYTOCHROME_P450 | 70 | Metabolism of xenobiotics by cytochrome P450 | 7 | 0.1 | 3.72E-04 |
| REACTOME_AFLATOXIN_ACTIVATION_AND_DETOXIFICATION | 19 | Aflatoxin activation and detoxification | 4 | 0.2105 | 4.01E-04 |
| KEGG_CYSTEINE_AND_METHIONINE_METABOLISM | 34 | Cysteine and methionine metabolism | 5 | 0.1471 | 4.33E-04 |
| WP_NRF2_PATHWAY | 145 | NRF2 pathway | 10 | 0.069 | 4.83E-04 |
| HALLMARK_HEME_METABOLISM | 200 | Genes involved in metabolism of heme (a cofactor consisting of iron and porphyrin) and erythroblast differentiation. | 12 | 0.06 | 4.90E-04 |
| REACTOME_CLASS_I_PEROXISOMAL_MEMBRANE_PROTEIN_IMPORT | 20 | Class I peroxisomal membrane protein import | 4 | 0.2 | 4.94E-04 |
| REACTOME_RAS_ACTIVATION_UPON_CA2_INFLUX_THROUGH_NMDA_RECEPTOR | 20 | Ras activation upon Ca2+ influx through NMDA receptor | 4 | 0.2 | 4.94E-04 |
| REACTOME_MULTIFUNCTIONAL_ANION_EXCHANGERS | 9 | Multifunctional anion exchangers | 3 | 0.3333 | 5.29E-04 |
| REACTOME_NEGATIVE_REGULATION_OF_NMDA_RECEPTOR_MEDIATED_NEURONAL_TRANSMISSION | 21 | Negative regulation of NMDA receptor-mediated neuronal transmission | 4 | 0.1905 | 6.01E-04 |
| REACTOME_UNBLOCKING_OF_NMDA_RECEPTORS_GLUTAMATE_BINDING_AND_ACTIVATION | 21 | Unblocking of NMDA receptors, glutamate binding and activation | 4 | 0.1905 | 6.01E-04 |
| REACTOME_NEUROTRANSMITTER_RECEPTORS_AND_POSTSYNAPTIC_SIGNAL_TRANSMISSION | 205 | Neurotransmitter receptors and postsynaptic signal transmission | 12 | 0.0585 | 6.10E-04 |
| REACTOME_ION_TRANSPORT_BY_P_TYPE_ATPASES | 55 | Ion transport by P-type ATPases | 6 | 0.1091 | 6.11E-04 |
| REACTOME_SENSORY_PROCESSING_OF_SOUND_BY_OUTER_HAIR_CELLS_OF_THE_COCHLEA | 55 | Sensory processing of sound by outer hair cells of the cochlea | 6 | 0.1091 | 6.11E-04 |
| REACTOME_SENSORY_PROCESSING_OF_SOUND | 77 | Sensory processing of sound | 7 | 0.0909 | 6.66E-04 |
| WP_CALCIUM_REGULATION_IN_THE_CARDIAC_CELL | 152 | Calcium Regulation in the Cardiac Cell | 10 | 0.0658 | 6.99E-04 |
| REACTOME_CITRIC_ACID_CYCLE_TCA_CYCLE | 22 | Citric acid cycle (TCA cycle) | 4 | 0.1818 | 7.24E-04 |
| REACTOME_DEFECTS_IN_VITAMIN_AND_COFACTOR_METABOLISM | 22 | Defects in vitamin and cofactor metabolism | 4 | 0.1818 | 7.24E-04 |
| REACTOME_CAMK_IV_MEDIATED_PHOSPHORYLATION_OF_CREB | 10 | CaMK IV-mediated phosphorylation of CREB | 3 | 0.3 | 7.45E-04 |
| BIOCARTA_INTRINSIC_PATHWAY | 23 | Intrinsic Prothrombin Activation Pathway | 4 | 0.1739 | 8.63E-04 |
| REACTOME_LONG_TERM_POTENTIATION | 23 | Long-term potentiation | 4 | 0.1739 | 8.63E-04 |
| WP_VITAMIN_DSENSITIVE_CALCIUM_SIGNALING_IN_DEPRESSION | 40 | Vitamin D-sensitive calcium signaling in depression | 5 | 0.125 | 9.33E-04 |
| REACTOME_DAG_AND_IP3_SIGNALING | 41 | DAG and IP3 signaling | 5 | 0.122 | 1.05E-03 |
| REACTOME_LYSINE_CATABOLISM | 12 | Lysine catabolism | 3 | 0.25 | 1.33E-03 |
| REACTOME_MUSCLE_CONTRACTION | 195 | Muscle contraction | 11 | 0.0564 | 1.37E-03 |
| REACTOME_ASSEMBLY_AND_CELL_SURFACE_PRESENTATION_OF_NMDA_RECEPTORS | 44 | Assembly and cell surface presentation of NMDA receptors | 5 | 0.1136 | 1.45E-03 |
| WP_RENIN_ANGIOTENSIN_ALDOSTERONE_SYSTEM_RAAS | 44 | Renin Angiotensin Aldosterone System (RAAS) | 5 | 0.1136 | 1.45E-03 |

Gene/Gene Set Overlap Matrix

| Entrez Gene Id | Gene Symbol | Gene Description | REACTOME_METABOLISM_OF_AMINO_ACIDS_AND_DERIVATIVES | REACTOME_TRANSPORT_OF_SMALL_MOLECULES | KEGG_VALINE_LEUCINE_AND_ISOLEUCINE_DEGRADATION |
|---|---|---|---|---|---|
| 594 | BCKDHB | branched chain keto acid dehydrogenase E1 subunit beta [Source:HGNC Symbol;Acc:HGNC:987] | REACTOME_METABOLISM_OF_AMINO_ACIDS_AND_DERIVATIVES | | KEGG_VALINE_LEUCINE_AND_ISOLEUCINE_DEGRADATION |
| 1629 | DBT | dihydrolipoamide branched chain transacylase E2 [Source:HGNC Symbol;Acc:HGNC:2698] | REACTOME_METABOLISM_OF_AMINO_ACIDS_AND_DERIVATIVES | | KEGG_VALINE_LEUCINE_AND_ISOLEUCINE_DEGRADATION |
| 64087 | MCCC2 | methylcrotonoyl-CoA carboxylase 2 [Source:HGNC Symbol;Acc:HGNC:6937] | REACTOME_METABOLISM_OF_AMINO_ACIDS_AND_DERIVATIVES | | KEGG_VALINE_LEUCINE_AND_ISOLEUCINE_DEGRADATION |
| 501 | ALDH7A1 | aldehyde dehydrogenase 7 family member A1 [Source:HGNC Symbol;Acc:HGNC:877] | REACTOME_METABOLISM_OF_AMINO_ACIDS_AND_DERIVATIVES | | KEGG_VALINE_LEUCINE_AND_ISOLEUCINE_DEGRADATION |
| 26275 | HIBCH | 3-hydroxyisobutyryl-CoA hydrolase [Source:HGNC Symbol;Acc:HGNC:4908] | REACTOME_METABOLISM_OF_AMINO_ACIDS_AND_DERIVATIVES | | KEGG_VALINE_LEUCINE_AND_ISOLEUCINE_DEGRADATION |
| 4329 | ALDH6A1 | aldehyde dehydrogenase 6 family member A1 [Source:HGNC Symbol;Acc:HGNC:7179] | REACTOME_METABOLISM_OF_AMINO_ACIDS_AND_DERIVATIVES | | KEGG_VALINE_LEUCINE_AND_ISOLEUCINE_DEGRADATION |
| 3712 | IVD | isovaleryl-CoA dehydrogenase [Source:HGNC Symbol;Acc:HGNC:6186] | REACTOME_METABOLISM_OF_AMINO_ACIDS_AND_DERIVATIVES | | KEGG_VALINE_LEUCINE_AND_ISOLEUCINE_DEGRADATION |
| 27034 | ACAD8 | acyl-CoA dehydrogenase family member 8 [Source:HGNC Symbol;Acc:HGNC:87] | REACTOME_METABOLISM_OF_AMINO_ACIDS_AND_DERIVATIVES | | KEGG_VALINE_LEUCINE_AND_ISOLEUCINE_DEGRADATION |
| 1610 | DAO | D-amino acid oxidase [Source:HGNC Symbol;Acc:HGNC:2671] | REACTOME_METABOLISM_OF_AMINO_ACIDS_AND_DERIVATIVES | | |
| 275 | AMT | aminomethyltransferase [Source:HGNC Symbol;Acc:HGNC:473] | REACTOME_METABOLISM_OF_AMINO_ACIDS_AND_DERIVATIVES | | |
| 2628 | GATM | glycine amidinotransferase [Source:HGNC Symbol;Acc:HGNC:4175] | REACTOME_METABOLISM_OF_AMINO_ACIDS_AND_DERIVATIVES | | |
| 635 | BHMT | betaine--homocysteine S-methyltransferase [Source:HGNC Symbol;Acc:HGNC:1047] | REACTOME_METABOLISM_OF_AMINO_ACIDS_AND_DERIVATIVES | | |
| 51268 | PIPOX | pipecolic acid and sarcosine oxidase [Source:HGNC Symbol;Acc:HGNC:17804] | REACTOME_METABOLISM_OF_AMINO_ACIDS_AND_DERIVATIVES | | |
| 6470 | SHMT1 | serine hydroxymethyltransferase 1 [Source:HGNC Symbol;Acc:HGNC:10850] | REACTOME_METABOLISM_OF_AMINO_ACIDS_AND_DERIVATIVES | | |
| 26227 | PHGDH | phosphoglycerate dehydrogenase [Source:HGNC Symbol;Acc:HGNC:8923] | REACTOME_METABOLISM_OF_AMINO_ACIDS_AND_DERIVATIVES | | |
| 1757 | SARDH | sarcosine dehydrogenase [Source:HGNC Symbol;Acc:HGNC:10536] | REACTOME_METABOLISM_OF_AMINO_ACIDS_AND_DERIVATIVES | | |
| 29958 | DMGDH | dimethylglycine dehydrogenase [Source:HGNC Symbol;Acc:HGNC:24475] | REACTOME_METABOLISM_OF_AMINO_ACIDS_AND_DERIVATIVES | | |
| 8659 | ALDH4A1 | aldehyde dehydrogenase 4 family member A1 [Source:HGNC Symbol;Acc:HGNC:406] | REACTOME_METABOLISM_OF_AMINO_ACIDS_AND_DERIVATIVES | | |
| 58510 | PRODH2 | proline dehydrogenase 2 [Source:HGNC Symbol;Acc:HGNC:17325] | REACTOME_METABOLISM_OF_AMINO_ACIDS_AND_DERIVATIVES | | |
| 4967 | OGDH | oxoglutarate dehydrogenase [Source:HGNC Symbol;Acc:HGNC:8124] | REACTOME_METABOLISM_OF_AMINO_ACIDS_AND_DERIVATIVES | | |
| 5827 | PXMP2 | peroxisomal membrane protein 2 [Source:HGNC Symbol;Acc:HGNC:9716] | REACTOME_METABOLISM_OF_AMINO_ACIDS_AND_DERIVATIVES | | |
| 55526 | DHTKD1 | dehydrogenase E1 and transketolase domain containing 1 [Source:HGNC Symbol;Acc:HGNC:23537] | REACTOME_METABOLISM_OF_AMINO_ACIDS_AND_DERIVATIVES | | |
| 112817 | HOGA1 | 4-hydroxy-2-oxoglutarate aldolase 1 [Source:HGNC Symbol;Acc:HGNC:25155] | REACTOME_METABOLISM_OF_AMINO_ACIDS_AND_DERIVATIVES | | |
| 1644 | DDC | dopa decarboxylase [Source:HGNC Symbol;Acc:HGNC:2719] | REACTOME_METABOLISM_OF_AMINO_ACIDS_AND_DERIVATIVES | | |
| 191 | AHCY | adenosylhomocysteinase [Source:HGNC Symbol;Acc:HGNC:343] | REACTOME_METABOLISM_OF_AMINO_ACIDS_AND_DERIVATIVES | | |
| 445 | ASS1 | argininosuccinate synthase 1 [Source:HGNC Symbol;Acc:HGNC:758] | REACTOME_METABOLISM_OF_AMINO_ACIDS_AND_DERIVATIVES | | |
| 2747 | GLUD2 | glutamate dehydrogenase 2 [Source:HGNC Symbol;Acc:HGNC:4336] | REACTOME_METABOLISM_OF_AMINO_ACIDS_AND_DERIVATIVES | | |
| 1158 | CKM | creatine kinase, M-type [Source:HGNC Symbol;Acc:HGNC:1994] | REACTOME_METABOLISM_OF_AMINO_ACIDS_AND_DERIVATIVES | | |
| 79814 | AGMAT | agmatinase [Source:HGNC Symbol;Acc:HGNC:18407] | REACTOME_METABOLISM_OF_AMINO_ACIDS_AND_DERIVATIVES | | |
| 10841 | FTCD | formimidoyltransferase cyclodeaminase [Source:HGNC Symbol;Acc:HGNC:3974] | REACTOME_METABOLISM_OF_AMINO_ACIDS_AND_DERIVATIVES | | |
| 1621 | DBH | dopamine beta-hydroxylase [Source:HGNC Symbol;Acc:HGNC:2689] | REACTOME_METABOLISM_OF_AMINO_ACIDS_AND_DERIVATIVES | | |
| 23498 | HAAO | 3-hydroxyanthranilate 3,4-dioxygenase [Source:HGNC Symbol;Acc:HGNC:4796] | REACTOME_METABOLISM_OF_AMINO_ACIDS_AND_DERIVATIVES | | |
| 169355 | IDO2 | indoleamine 2,3-dioxygenase 2 [Source:HGNC Symbol;Acc:HGNC:27269] | REACTOME_METABOLISM_OF_AMINO_ACIDS_AND_DERIVATIVES | | |
| 2346 | FOLH1 | folate hydrolase 1 [Source:HGNC Symbol;Acc:HGNC:3788] | REACTOME_METABOLISM_OF_AMINO_ACIDS_AND_DERIVATIVES | | |
| 2875 | GPT | glutamic--pyruvic transaminase [Source:HGNC Symbol;Acc:HGNC:4552] | REACTOME_METABOLISM_OF_AMINO_ACIDS_AND_DERIVATIVES | | |
| 5860 | QDPR | quinoid dihydropteridine reductase [Source:HGNC Symbol;Acc:HGNC:9752] | REACTOME_METABOLISM_OF_AMINO_ACIDS_AND_DERIVATIVES | | |
| 5053 | PAH | phenylalanine hydroxylase [Source:HGNC Symbol;Acc:HGNC:8582] | REACTOME_METABOLISM_OF_AMINO_ACIDS_AND_DERIVATIVES | | |
| 10247 | RIDA | reactive intermediate imine deaminase A homolog [Source:HGNC Symbol;Acc:HGNC:16897] | REACTOME_METABOLISM_OF_AMINO_ACIDS_AND_DERIVATIVES | | |
| 23382 | AHCYL2 | adenosylhomocysteinase like 2 [Source:HGNC Symbol;Acc:HGNC:22204] | | REACTOME_TRANSPORT_OF_SMALL_MOLECULES | |
| 340024 | SLC6A19 | solute carrier family 6 member 19 [Source:HGNC Symbol;Acc:HGNC:27960] | | REACTOME_TRANSPORT_OF_SMALL_MOLECULES | |
| 348932 | SLC6A18 | solute carrier family 6 member 18 [Source:HGNC Symbol;Acc:HGNC:26441] | | REACTOME_TRANSPORT_OF_SMALL_MOLECULES | |
| 153201 | SLC36A2 | solute carrier family 36 member 2 [Source:HGNC Symbol;Acc:HGNC:18762] | | REACTOME_TRANSPORT_OF_SMALL_MOLECULES | |
| 142680 | SLC34A3 | solute carrier family 34 member 3 [Source:HGNC Symbol;Acc:HGNC:20305] | | REACTOME_TRANSPORT_OF_SMALL_MOLECULES | |
| 6569 | SLC34A1 | solute carrier family 34 member 1 [Source:HGNC Symbol;Acc:HGNC:11019] | | REACTOME_TRANSPORT_OF_SMALL_MOLECULES | |
| 6550 | SLC9A3 | solute carrier family 9 member A3 [Source:HGNC Symbol;Acc:HGNC:11073] | | REACTOME_TRANSPORT_OF_SMALL_MOLECULES | |
| 6559 | SLC12A3 | solute carrier family 12 member 3 [Source:HGNC Symbol;Acc:HGNC:10912] | | REACTOME_TRANSPORT_OF_SMALL_MOLECULES | |
| 6557 | SLC12A1 | solute carrier family 12 member 1 [Source:HGNC Symbol;Acc:HGNC:10910] | | REACTOME_TRANSPORT_OF_SMALL_MOLECULES | |
| 9990 | SLC12A6 | solute carrier family 12 member 6 [Source:HGNC Symbol;Acc:HGNC:10914] | | REACTOME_TRANSPORT_OF_SMALL_MOLECULES | |
| 6521 | SLC4A1 | solute carrier family 4 member 1 (Diego blood group) [Source:HGNC Symbol;Acc:HGNC:11027] | | REACTOME_TRANSPORT_OF_SMALL_MOLECULES | |
| 5172 | SLC26A4 | solute carrier family 26 member 4 [Source:HGNC Symbol;Acc:HGNC:8818] | | REACTOME_TRANSPORT_OF_SMALL_MOLECULES | |
| 6546 | SLC8A1 | solute carrier family 8 member A1 [Source:HGNC Symbol;Acc:HGNC:11068] | | REACTOME_TRANSPORT_OF_SMALL_MOLECULES | |
| 159963 | SLC5A12 | solute carrier family 5 member 12 [Source:HGNC Symbol;Acc:HGNC:28750] | | REACTOME_TRANSPORT_OF_SMALL_MOLECULES | |
| 115019 | SLC26A9 | solute carrier family 26 member 9 [Source:HGNC Symbol;Acc:HGNC:14469] | | REACTOME_TRANSPORT_OF_SMALL_MOLECULES | |
| 8501 | SLC43A1 | solute carrier family 43 member 1 [Source:HGNC Symbol;Acc:HGNC:9225] | | REACTOME_TRANSPORT_OF_SMALL_MOLECULES | |
| 389015 | SLC9A4 | solute carrier family 9 member A4 [Source:HGNC Symbol;Acc:HGNC:11077] | | REACTOME_TRANSPORT_OF_SMALL_MOLECULES | |
| 6524 | SLC5A2 | solute carrier family 5 member 2 [Source:HGNC Symbol;Acc:HGNC:11037] | | REACTOME_TRANSPORT_OF_SMALL_MOLECULES | |
| 6540 | SLC6A13 | solute carrier family 6 member 13 [Source:HGNC Symbol;Acc:HGNC:11046] | | REACTOME_TRANSPORT_OF_SMALL_MOLECULES | |
| 283375 | SLC39A5 | solute carrier family 39 member 5 [Source:HGNC Symbol;Acc:HGNC:20502] | | REACTOME_TRANSPORT_OF_SMALL_MOLECULES | |
| 154091 | SLC2A12 | solute carrier family 2 member 12 [Source:HGNC Symbol;Acc:HGNC:18067] | | REACTOME_TRANSPORT_OF_SMALL_MOLECULES | |
| 66035 | SLC2A11 | solute carrier family 2 member 11 [Source:HGNC Symbol;Acc:HGNC:14239] | | REACTOME_TRANSPORT_OF_SMALL_MOLECULES | |
| 169026 | SLC30A8 | solute carrier family 30 member 8 [Source:HGNC Symbol;Acc:HGNC:20303] | | REACTOME_TRANSPORT_OF_SMALL_MOLECULES | |
| 9058 | SLC13A2 | solute carrier family 13 member 2 [Source:HGNC Symbol;Acc:HGNC:10917] | | REACTOME_TRANSPORT_OF_SMALL_MOLECULES | |
| 353189 | SLCO4C1 | solute carrier organic anion transporter family member 4C1 [Source:HGNC Symbol;Acc:HGNC:23612] | | REACTOME_TRANSPORT_OF_SMALL_MOLECULES | |
| 9153 | SLC28A2 | solute carrier family 28 member 2 [Source:HGNC Symbol;Acc:HGNC:11002] | | REACTOME_TRANSPORT_OF_SMALL_MOLECULES | |
| 760 | CA2 | carbonic anhydrase 2 [Source:HGNC Symbol;Acc:HGNC:1373] | | REACTOME_TRANSPORT_OF_SMALL_MOLECULES | |
| 27329 | ANGPTL3 | angiopoietin like 3 [Source:HGNC Symbol;Acc:HGNC:491] | | REACTOME_TRANSPORT_OF_SMALL_MOLECULES | |
| 8029 | CUBN | cubilin [Source:HGNC Symbol;Acc:HGNC:2548] | | REACTOME_TRANSPORT_OF_SMALL_MOLECULES | |
| 481 | ATP1B1 | ATPase Na+/K+ transporting subunit beta 1 [Source:HGNC Symbol;Acc:HGNC:804] | | REACTOME_TRANSPORT_OF_SMALL_MOLECULES | |
| 81693 | AMN | amnion associated transmembrane protein [Source:HGNC Symbol;Acc:HGNC:14604] | | REACTOME_TRANSPORT_OF_SMALL_MOLECULES | |
| 64241 | ABCG8 | ATP binding cassette subfamily G member 8 [Source:HGNC Symbol;Acc:HGNC:13887] | | REACTOME_TRANSPORT_OF_SMALL_MOLECULES | |
| 11160 | ERLIN2 | ER lipid raft associated 2 [Source:HGNC Symbol;Acc:HGNC:1356] | | REACTOME_TRANSPORT_OF_SMALL_MOLECULES | |
| 368 | ABCC6 | ATP binding cassette subfamily C member 6 [Source:HGNC Symbol;Acc:HGNC:57] | | REACTOME_TRANSPORT_OF_SMALL_MOLECULES | |
| 816 | CAMK2B | calcium/calmodulin dependent protein kinase II beta [Source:HGNC Symbol;Acc:HGNC:1461] | | REACTOME_TRANSPORT_OF_SMALL_MOLECULES | |
| 815 | CAMK2A | calcium/calmodulin dependent protein kinase II alpha [Source:HGNC Symbol;Acc:HGNC:1460] | | REACTOME_TRANSPORT_OF_SMALL_MOLECULES | |
| 818 | CAMK2G | calcium/calmodulin dependent protein kinase II gamma [Source:HGNC Symbol;Acc:HGNC:1463] | | REACTOME_TRANSPORT_OF_SMALL_MOLECULES | |
| 2785 | GNG3 | G protein subunit gamma 3 [Source:HGNC Symbol;Acc:HGNC:4405] | | REACTOME_TRANSPORT_OF_SMALL_MOLECULES | |
| 11194 | ABCB8 | ATP binding cassette subfamily B member 8 [Source:HGNC Symbol;Acc:HGNC:49] | | REACTOME_TRANSPORT_OF_SMALL_MOLECULES | |
| 491 | ATP2B2 | ATPase plasma membrane Ca2+ transporting 2 [Source:HGNC Symbol;Acc:HGNC:815] | | REACTOME_TRANSPORT_OF_SMALL_MOLECULES | |
| 479 | ATP12A | ATPase H+/K+ transporting non-gastric alpha2 subunit [Source:HGNC Symbol;Acc:HGNC:13816] | | REACTOME_TRANSPORT_OF_SMALL_MOLECULES | |
| 7442 | TRPV1 | transient receptor potential cation channel subfamily V member 1 [Source:HGNC Symbol;Acc:HGNC:12716] | | REACTOME_TRANSPORT_OF_SMALL_MOLECULES | |
| 59341 | TRPV4 | transient receptor potential cation channel subfamily V member 4 [Source:HGNC Symbol;Acc:HGNC:18083] | | REACTOME_TRANSPORT_OF_SMALL_MOLECULES | |
| 140803 | TRPM6 | transient receptor potential cation channel subfamily M member 6 [Source:HGNC Symbol;Acc:HGNC:17995] | | REACTOME_TRANSPORT_OF_SMALL_MOLECULES | |
| 80036 | TRPM3 | transient receptor potential cation channel subfamily M member 3 [Source:HGNC Symbol;Acc:HGNC:17992] | | REACTOME_TRANSPORT_OF_SMALL_MOLECULES | |
| 65266 | WNK4 | WNK lysine deficient protein kinase 4 [Source:HGNC Symbol;Acc:HGNC:14544] | | REACTOME_TRANSPORT_OF_SMALL_MOLECULES | |
| 6338 | SCNN1B | sodium channel epithelial 1 subunit beta [Source:HGNC Symbol;Acc:HGNC:10600] | | REACTOME_TRANSPORT_OF_SMALL_MOLECULES | |
| 1184 | CLCN5 | chloride voltage-gated channel 5 [Source:HGNC Symbol;Acc:HGNC:2023] | | REACTOME_TRANSPORT_OF_SMALL_MOLECULES | |
| 345 | APOC3 | apolipoprotein C3 [Source:HGNC Symbol;Acc:HGNC:610] | | REACTOME_TRANSPORT_OF_SMALL_MOLECULES | |
| 120 | ADD3 | adducin 3 [Source:HGNC Symbol;Acc:HGNC:245] | | REACTOME_TRANSPORT_OF_SMALL_MOLECULES | |
| 26234 | FBXL5 | F-box and leucine rich repeat protein 5 [Source:HGNC Symbol;Acc:HGNC:13602] | | REACTOME_TRANSPORT_OF_SMALL_MOLECULES | |
| 10367 | MICU1 | mitochondrial calcium uptake 1 [Source:HGNC Symbol;Acc:HGNC:1530] | | REACTOME_TRANSPORT_OF_SMALL_MOLECULES | |
| 224 | ALDH3A2 | aldehyde dehydrogenase 3 family member A2 [Source:HGNC Symbol;Acc:HGNC:403] | | | KEGG_VALINE_LEUCINE_AND_ISOLEUCINE_DEGRADATION |
| 34 | ACADM | acyl-CoA dehydrogenase medium chain [Source:HGNC Symbol;Acc:HGNC:89] | | | KEGG_VALINE_LEUCINE_AND_ISOLEUCINE_DEGRADATION |
| 3033 | HADH | hydroxyacyl-CoA dehydrogenase [Source:HGNC Symbol;Acc:HGNC:4799] | | | KEGG_VALINE_LEUCINE_AND_ISOLEUCINE_DEGRADATION |
| 18 | ABAT | 4-aminobutyrate aminotransferase [Source:HGNC Symbol;Acc:HGNC:23] | | | KEGG_VALINE_LEUCINE_AND_ISOLEUCINE_DEGRADATION |
| 84693 | MCEE | methylmalonyl-CoA epimerase [Source:HGNC Symbol;Acc:HGNC:16732] | | | KEGG_VALINE_LEUCINE_AND_ISOLEUCINE_DEGRADATION |
| 5019 | OXCT1 | 3-oxoacid CoA-transferase 1 [Source:HGNC Symbol;Acc:HGNC:8527] | | | KEGG_VALINE_LEUCINE_AND_ISOLEUCINE_DEGRADATION |
| 132158 | GLYCTK | glycerate kinase [Source:HGNC Symbol;Acc:HGNC:24247] | | | |
| 50 | ACO2 | aconitase 2 [Source:HGNC Symbol;Acc:HGNC:118] | | | |
| 128 | ADH5 | alcohol dehydrogenase 5 (class III), chi polypeptide [Source:HGNC Symbol;Acc:HGNC:253] | | | |
| 8309 | ACOX2 | acyl-CoA oxidase 2 [Source:HGNC Symbol;Acc:HGNC:120] | | | |
| 2571 | GAD1 | glutamate decarboxylase 1 [Source:HGNC Symbol;Acc:HGNC:4092] | | | |
| 51733 | UPB1 | beta-ureidopropionase 1 [Source:HGNC Symbol;Acc:HGNC:16297] | | | |
| 8529 | CYP4F2 | cytochrome P450 family 4 subfamily F member 2 [Source:HGNC Symbol;Acc:HGNC:2645] | | | |
| 866 | SERPINA6 | serpin family A member 6 [Source:HGNC Symbol;Acc:HGNC:1540] | | | |
| 2326 | FMO1 | flavin containing dimethylaniline monoxygenase 1 [Source:HGNC Symbol;Acc:HGNC:3769] | | | |

Bulk RNA-seq DOWN

| | | |
|---|---|---|
| 670 | BPHL | biphenyl hydrolase like [Source:HGNC Symbol;Acc:HGNC:1094] |
| 2203 | FBP1 | fructose-bisphosphatase 1 [Source:HGNC Symbol;Acc:HGNC:3606] |
| 26007 | TKFC | triokinase and FMN cyclase [Source:HGNC Symbol;Acc:HGNC:24552] |
| 2160 | F11 | coagulation factor XI [Source:HGNC Symbol;Acc:HGNC:3529] |
| 2099 | ESR1 | estrogen receptor 1 [Source:HGNC Symbol;Acc:HGNC:3467] |
| 26999 | CYFIP2 | cytoplasmic FMR1 interacting protein 2 [Source:HGNC Symbol;Acc:HGNC:13760] |
| 3273 | HRG | histidine rich glycoprotein [Source:HGNC Symbol;Acc:HGNC:5181] |
| 7448 | VTN | vitronectin [Source:HGNC Symbol;Acc:HGNC:12724] |
| 26 | AOC1 | amine oxidase copper containing 1 [Source:HGNC Symbol;Acc:HGNC:80] |
| 95 | ACY1 | aminoacylase 1 [Source:HGNC Symbol;Acc:HGNC:177] |
| 8802 | SUCLG1 | succinate-CoA ligase GDP/ADP-forming subunit alpha [Source:HGNC Symbol;Acc:HGNC:11449] |
| 23395 | LARS2 | leucyl-tRNA synthetase 2, mitochondrial [Source:HGNC Symbol;Acc:HGNC:17095] |
| 1384 | CRAT | carnitine O-acetyltransferase [Source:HGNC Symbol;Acc:HGNC:2342] |
| 51179 | HAO2 | hydroxyacid oxidase 2 [Source:HGNC Symbol;Acc:HGNC:4810] |
| 2053 | EPHX2 | epoxide hydrolase 2 [Source:HGNC Symbol;Acc:HGNC:3402] |
| 5264 | PHYH | phytanoyl-CoA 2-hydroxylase [Source:HGNC Symbol;Acc:HGNC:8940] |
| 5193 | PEX12 | peroxisomal biogenesis factor 12 [Source:HGNC Symbol;Acc:HGNC:8854] |
| 5189 | PEX1 | peroxisomal biogenesis factor 1 [Source:HGNC Symbol;Acc:HGNC:8850] |
| 10901 | DHRS4 | dehydrogenase/reductase 4 [Source:HGNC Symbol;Acc:HGNC:16985] |
| 11264 | PXMP4 | peroxisomal membrane protein 4 [Source:HGNC Symbol;Acc:HGNC:15920] |
| 83594 | NUDT12 | nudix hydrolase 12 [Source:HGNC Symbol;Acc:HGNC:18826] |
| 84735 | CNDP1 | carnosine dipeptidase 1 [Source:HGNC Symbol;Acc:HGNC:20675] |
| 3248 | HPGD | 15-hydroxyprostaglandin dehydrogenase [Source:HGNC Symbol;Acc:HGNC:5154] |
| 134526 | ACOT12 | acyl-CoA thioesterase 12 [Source:HGNC Symbol;Acc:HGNC:24436] |
| 26027 | ACOT11 | acyl-CoA thioesterase 11 [Source:HGNC Symbol;Acc:HGNC:18156] |
| 80724 | ACAD10 | acyl-CoA dehydrogenase family member 10 [Source:HGNC Symbol;Acc:HGNC:21597] |
| 81616 | ACSBG2 | acyl-CoA synthetase bubblegum family member 2 [Source:HGNC Symbol;Acc:HGNC:24174] |
| 2166 | FAAH | fatty acid amide hydrolase [Source:HGNC Symbol;Acc:HGNC:3553] |
| 84869 | CBR4 | carbonyl reductase 4 [Source:HGNC Symbol;Acc:HGNC:25891] |
| 7069 | THRSP | thyroid hormone responsive [Source:HGNC Symbol;Acc:HGNC:11800] |
| 2819 | GPD1 | glycerol-3-phosphate dehydrogenase 1 [Source:HGNC Symbol;Acc:HGNC:4455] |
| 1576 | CYP3A4 | cytochrome P450 family 3 subfamily A member 4 [Source:HGNC Symbol;Acc:HGNC:2637] |
| 56894 | AGPAT3 | 1-acylglycerol-3-phosphate O-acyltransferase 3 [Source:HGNC Symbol;Acc:HGNC:326] |
| 80339 | PNPLA3 | patatin like phospholipase domain containing 3 [Source:HGNC Symbol;Acc:HGNC:18590] |
| 9468 | PCYT1B | phosphate cytidylyltransferase 1, choline, beta [Source:HGNC Symbol;Acc:HGNC:8755] |
| 56848 | SPHK2 | sphingosine kinase 2 [Source:HGNC Symbol;Acc:HGNC:18859] |
| 5783 | PTPN13 | protein tyrosine phosphatase non-receptor type 13 [Source:HGNC Symbol;Acc:HGNC:9646] |
| 64850 | ETNPPL | ethanolamine-phosphate phospho-lyase [Source:HGNC Symbol;Acc:HGNC:14404] |
| 9362 | CPNE6 | copine 6 [Source:HGNC Symbol;Acc:HGNC:2319] |
| 22908 | SACM1L | SAC1 like phosphatidylinositide phosphatase [Source:HGNC Symbol;Acc:HGNC:17059] |
| 54477 | PLEKHA5 | pleckstrin homology domain containing A5 [Source:HGNC Symbol;Acc:HGNC:30036] |
| 133121 | ENPP6 | ectonucleotide pyrophosphatase/phosphodiesterase 6 [Source:HGNC Symbol;Acc:HGNC:23409] |
| 416 | ARSF | arylsulfatase F [Source:HGNC Symbol;Acc:HGNC:721] |
| 80168 | MOGAT2 | monoacylglycerol O-acyltransferase 2 [Source:HGNC Symbol;Acc:HGNC:23248] |
| 54511 | HMGCLL1 | 3-hydroxymethyl-3-methylglutaryl-CoA lyase like 1 [Source:HGNC Symbol;Acc:HGNC:21359] |
| 6555 | SLC10A2 | solute carrier family 10 member 2 [Source:HGNC Symbol;Acc:HGNC:10906] |
| 114876 | OSBPL1A | oxysterol binding protein like 1A [Source:HGNC Symbol;Acc:HGNC:16398] |
| 23179 | RGL1 | ral guanine nucleotide dissociation stimulator like 1 [Source:HGNC Symbol;Acc:HGNC:30281] |
| 8706 | B3GALNT1 | beta-1,3-N-acetylgalactosaminyltransferase 1 (globoside blood group) [Source:HGNC Symbol;Acc:HGNC:918] |
| 8801 | SUCLG2 | succinate-CoA ligase GDP-forming subunit beta [Source:HGNC Symbol;Acc:HGNC:11450] |
| 210 | ALAD | aminolevulinate dehydratase [Source:HGNC Symbol;Acc:HGNC:395] |
| 55753 | OGDHL | oxoglutarate dehydrogenase L [Source:HGNC Symbol;Acc:HGNC:25590] |
| 197257 | LDHD | lactate dehydrogenase D [Source:HGNC Symbol;Acc:HGNC:19708] |
| 131118 | DNAJC19 | DnaJ heat shock protein family (Hsp40) member C19 [Source:HGNC Symbol;Acc:HGNC:30528] |
| 200895 | DHFR2 | dihydrofolate reductase 2 [Source:HGNC Symbol;Acc:HGNC:27309] |
| 29947 | DNMT3L | DNA methyltransferase 3 like [Source:HGNC Symbol;Acc:HGNC:2980] |
| 1551 | CYP3A7 | cytochrome P450 family 3 subfamily A member 7 [Source:HGNC Symbol;Acc:HGNC:2640] |
| 2938 | GSTA1 | glutathione S-transferase alpha 1 [Source:HGNC Symbol;Acc:HGNC:4626] |
| 2939 | GSTA2 | glutathione S-transferase alpha 2 [Source:HGNC Symbol;Acc:HGNC:4627] |
| 130 | ADH6 | alcohol dehydrogenase 6 (class V) [Source:HGNC Symbol;Acc:HGNC:255] |
| 22977 | AKR7A3 | aldo-keto reductase family 7 member A3 [Source:HGNC Symbol;Acc:HGNC:390] |
| 92292 | GLYATL1 | glycine-N-acyltransferase like 1 [Source:HGNC Symbol;Acc:HGNC:30519] |
| 54996 | MTARC2 | mitochondrial amidoxime reducing component 2 [Source:HGNC Symbol;Acc:HGNC:26064] |
| 124975 | GGT6 | gamma-glutamyltransferase 6 [Source:HGNC Symbol;Acc:HGNC:26891] |
| 9 | NAT1 | N-acetyltransferase 1 [Source:HGNC Symbol;Acc:HGNC:7645] |
| 55347 | ABHD10 | abhydrolase domain containing 10, depalmitoylase [Source:HGNC Symbol;Acc:HGNC:25656] |
| 10243 | GPHN | gephyrin [Source:HGNC Symbol;Acc:HGNC:15465] |
| 9963 | SLC23A1 | solute carrier family 23 member 1 [Source:HGNC Symbol;Acc:HGNC:10974] |
| 683 | BST1 | bone marrow stromal cell antigen 1 [Source:HGNC Symbol;Acc:HGNC:1118] |
| 686 | BTD | biotinidase [Source:HGNC Symbol;Acc:HGNC:1122] |
| 349565 | NMNAT3 | nicotinamide nucleotide adenylyltransferase 3 [Source:HGNC Symbol;Acc:HGNC:20989] |
| 9390 | SLC22A13 | solute carrier family 22 member 13 [Source:HGNC Symbol;Acc:HGNC:8494] |
| 2330 | FMO5 | flavin containing dimethylaniline monoxygenase 5 [Source:HGNC Symbol;Acc:HGNC:3773] |
| 2329 | FMO4 | flavin containing dimethylaniline monoxygenase 4 [Source:HGNC Symbol;Acc:HGNC:3772] |
| 260293 | CYP4X1 | cytochrome P450 family 4 subfamily X member 1 [Source:HGNC Symbol;Acc:HGNC:20244] |
| 9027 | NAT8 | N-acetyltransferase 8 (putative) [Source:HGNC Symbol;Acc:HGNC:18069] |
| 89792 | GAL3ST3 | galactose-3-O-sulfotransferase 3 [Source:HGNC Symbol;Acc:HGNC:24144] |
| 783 | CACNB2 | calcium voltage-gated channel auxiliary subunit beta 2 [Source:HGNC Symbol;Acc:HGNC:1402] |
| 57554 | LRRC7 | leucine rich repeat containing 7 [Source:HGNC Symbol;Acc:HGNC:18531] |
| 116443 | GRIN3A | glutamate ionotropic receptor NMDA type subunit 3A [Source:HGNC Symbol;Acc:HGNC:16767] |
| 23229 | ARHGEF9 | Cdc42 guanine nucleotide exchange factor 9 [Source:HGNC Symbol;Acc:HGNC:14561] |
| 2557 | GABRA4 | gamma-aminobutyric acid type A receptor subunit alpha4 [Source:HGNC Symbol;Acc:HGNC:4078] |
| 4137 | MAPT | microtubule associated protein tau [Source:HGNC Symbol;Acc:HGNC:6893] |
| 1143 | CHRNB4 | cholinergic receptor nicotinic beta 4 subunit [Source:HGNC Symbol;Acc:HGNC:1964] |
| 3766 | KCNJ10 | potassium inwardly rectifying channel subfamily J member 10 [Source:HGNC Symbol;Acc:HGNC:6256] |
| 83988 | NCALD | neurocalcin delta [Source:HGNC Symbol;Acc:HGNC:7655] |
| 785 | CACNB4 | calcium voltage-gated channel auxiliary subunit beta 4 [Source:HGNC Symbol;Acc:HGNC:1404] |
| 54207 | KCNK10 | potassium two pore domain channel subfamily K member 10 [Source:HGNC Symbol;Acc:HGNC:6273] |
| 2911 | GRM1 | glutamate metabotropic receptor 1 [Source:HGNC Symbol;Acc:HGNC:4593] |
| 3781 | KCNN2 | potassium calcium-activated channel subfamily N member 2 [Source:HGNC Symbol;Acc:HGNC:6291] |
| 3782 | KCNN3 | potassium calcium-activated channel subfamily N member 3 [Source:HGNC Symbol;Acc:HGNC:6292] |
| 26045 | LRRTM2 | leucine rich repeat transmembrane neuronal 2 [Source:HGNC Symbol;Acc:HGNC:19409] |
| 5106 | PCK2 | phosphoenolpyruvate carboxykinase 2, mitochondrial [Source:HGNC Symbol;Acc:HGNC:8725] |
| 5313 | PKLR | pyruvate kinase L/R [Source:HGNC Symbol;Acc:HGNC:9020] |
| 5224 | PGAM2 | phosphoglycerate mutase 2 [Source:HGNC Symbol;Acc:HGNC:8889] |
| 3709 | ITPR2 | inositol 1,4,5-trisphosphate receptor type 2 [Source:HGNC Symbol;Acc:HGNC:6181] |
| 10142 | AKAP9 | A-kinase anchoring protein 9 [Source:HGNC Symbol;Acc:HGNC:379] |
| 10008 | KCNE3 | potassium voltage-gated channel subfamily E regulatory subunit 3 [Source:HGNC Symbol;Acc:HGNC:6243] |
| 1607 | DGKB | diacylglycerol kinase beta [Source:HGNC Symbol;Acc:HGNC:2850] |
| 2155 | F7 | coagulation factor VII [Source:HGNC Symbol;Acc:HGNC:3544] |
| 3827 | KNG1 | kininogen 1 [Source:HGNC Symbol;Acc:HGNC:6383] |
| 3818 | KLKB1 | kallikrein B1 [Source:HGNC Symbol;Acc:HGNC:6371] |
| 2811 | GP1BA | glycoprotein Ib platelet subunit alpha [Source:HGNC Symbol;Acc:HGNC:4439] |
| 5104 | SERPINA5 | serpin family A member 5 [Source:HGNC Symbol;Acc:HGNC:8723] |
| 3795 | KHK | ketohexokinase [Source:HGNC Symbol;Acc:HGNC:6315] |
| 84951 | TNS4 | tensin 4 [Source:HGNC Symbol;Acc:HGNC:24352] |
| 3973 | LHCGR | luteinizing hormone/choriogonadotropin receptor [Source:HGNC Symbol;Acc:HGNC:6585] |
| 4923 | NTSR1 | neurotensin receptor 1 [Source:HGNC Symbol;Acc:HGNC:8039] |

| | | |
|---|---|---|
| 63928 | CHP2 | calcineurin like EF-hand protein 2 [Source:HGNC Symbol;Acc:HGNC:24927] |
| 145226 | RDH12 | retinol dehydrogenase 12 [Source:HGNC Symbol;Acc:HGNC:19977] |
| 317749 | DHRS4L2 | dehydrogenase/reductase 4 like 2 [Source:HGNC Symbol;Acc:HGNC:19731] |
| 27294 | DHDH | dihydrodiol dehydrogenase [Source:HGNC Symbol;Acc:HGNC:17887] |
| 1787 | TRDMT1 | tRNA aspartic acid methyltransferase 1 [Source:HGNC Symbol;Acc:HGNC:2977] |
| 64122 | FN3K | fructosamine 3 kinase [Source:HGNC Symbol;Acc:HGNC:24822] |
| 2235 | FECH | ferrochelatase [Source:HGNC Symbol;Acc:HGNC:3647] |
| 7029 | TFDP2 | transcription factor Dp-2 [Source:HGNC Symbol;Acc:HGNC:11751] |
| 8031 | NCOA4 | nuclear receptor coactivator 4 [Source:HGNC Symbol;Acc:HGNC:7671] |
| 10107 | TRIM10 | tripartite motif containing 10 [Source:HGNC Symbol;Acc:HGNC:10072] |
| 8991 | SELENBP1 | selenium binding protein 1 [Source:HGNC Symbol;Acc:HGNC:10719] |
| 23039 | XPO7 | exportin 7 [Source:HGNC Symbol;Acc:HGNC:14108] |
| 4647 | MYO7A | myosin VIIA [Source:HGNC Symbol;Acc:HGNC:7606] |
| 5962 | RDX | radixin [Source:HGNC Symbol;Acc:HGNC:9944] |
| 283310 | OTOGL | otogelin like [Source:HGNC Symbol;Acc:HGNC:26901] |
| 65217 | PCDH15 | protocadherin related 15 [Source:HGNC Symbol;Acc:HGNC:14674] |
| 5581 | PRKCE | protein kinase C epsilon [Source:HGNC Symbol;Acc:HGNC:9401] |
| 6000 | RGS7 | regulator of G protein signaling 7 [Source:HGNC Symbol;Acc:HGNC:10003] |
| 1288 | COL4A6 | collagen type IV alpha 6 chain [Source:HGNC Symbol;Acc:HGNC:2208] |
| 596 | BCL2 | BCL2 apoptosis regulator [Source:HGNC Symbol;Acc:HGNC:990] |
| 793 | CALB1 | calbindin 1 [Source:HGNC Symbol;Acc:HGNC:1434] |
| 186 | AGTR2 | angiotensin II receptor type 2 [Source:HGNC Symbol;Acc:HGNC:338] |
| 10013 | HDAC6 | histone deacetylase 6 [Source:HGNC Symbol;Acc:HGNC:14064] |
| 1896 | EDA | ectodysplasin A [Source:HGNC Symbol;Acc:HGNC:3157] |
| 9687 | GREB1 | growth regulating estrogen receptor binding 1 [Source:HGNC Symbol;Acc:HGNC:24885] |
| 5080 | PAX6 | paired box 6 [Source:HGNC Symbol;Acc:HGNC:8620] |
| 1501 | CTNND2 | catenin delta 2 [Source:HGNC Symbol;Acc:HGNC:2516] |
| 57476 | GRAMD1B | GRAM domain containing 1B [Source:HGNC Symbol;Acc:HGNC:29214] |
| 6387 | CXCL12 | C-X-C motif chemokine ligand 12 [Source:HGNC Symbol;Acc:HGNC:10672] |
| 1524 | CX3CR1 | C-X3-C motif chemokine receptor 1 [Source:HGNC Symbol;Acc:HGNC:2558] |
| 846 | CASR | calcium sensing receptor [Source:HGNC Symbol;Acc:HGNC:1514] |
| 56670 | SUCNR1 | succinate receptor 1 [Source:HGNC Symbol;Acc:HGNC:4542] |
| 165140 | OXER1 | oxoeicosanoid receptor 1 [Source:HGNC Symbol;Acc:HGNC:24884] |
| 10451 | VAV3 | vav guanine nucleotide exchange factor 3 [Source:HGNC Symbol;Acc:HGNC:12659] |
| 3062 | HCRTR2 | hypocretin receptor 2 [Source:HGNC Symbol;Acc:HGNC:4849] |
| 170589 | GPHA2 | glycoprotein hormone subunit alpha 2 [Source:HGNC Symbol;Acc:HGNC:18054] |
| 50649 | ARHGEF4 | Rho guanine nucleotide exchange factor 4 [Source:HGNC Symbol;Acc:HGNC:684] |
| 9890 | PLPPR4 | phospholipid phosphatase related 4 [Source:HGNC Symbol;Acc:HGNC:23496] |
| 54886 | PLPPR1 | phospholipid phosphatase related 1 [Source:HGNC Symbol;Acc:HGNC:25993] |
| 151531 | UPP2 | uridine phosphorylase 2 [Source:HGNC Symbol;Acc:HGNC:23061] |
| 124842 | TMEM132E | transmembrane protein 132E [Source:HGNC Symbol;Acc:HGNC:26991] |
| 140690 | CTCFL | CCCTC-binding factor like [Source:HGNC Symbol;Acc:HGNC:16234] |
| 283629 | TSSK4 | testis specific serine kinase 4 [Source:HGNC Symbol;Acc:HGNC:19825] |
| 340719 | NANOS1 | nanos C2HC-type zinc finger 1 [Source:HGNC Symbol;Acc:HGNC:23044] |
| 8858 | PROZ | protein Z, vitamin K dependent plasma glycoprotein [Source:HGNC Symbol;Acc:HGNC:9460] |
| 1515 | CTSV | cathepsin V [Source:HGNC Symbol;Acc:HGNC:2538] |
| 8416 | ANXA9 | annexin A9 [Source:HGNC Symbol;Acc:HGNC:547] |
| 5775 | PTPN4 | protein tyrosine phosphatase non-receptor type 4 [Source:HGNC Symbol;Acc:HGNC:9656] |
| 9365 | KL | klotho [Source:HGNC Symbol;Acc:HGNC:6344] |
| 4482 | MSRA | methionine sulfoxide reductase A [Source:HGNC Symbol;Acc:HGNC:7377] |
| 55789 | DEPDC1B | DEP domain containing 1B [Source:HGNC Symbol;Acc:HGNC:24902] |
| 57498 | KIDINS220 | kinase D interacting substrate 220 [Source:HGNC Symbol;Acc:HGNC:29508] |
| 64220 | STRA6 | signaling receptor and transporter of retinol STRA6 [Source:HGNC Symbol;Acc:HGNC:30650] |
| 135228 | CD109 | CD109 molecule [Source:HGNC Symbol;Acc:HGNC:21685] |
| 7070 | THY1 | Thy-1 cell surface antigen [Source:HGNC Symbol;Acc:HGNC:11801] |
| 84720 | PIGO | phosphatidylinositol glycan anchor biosynthesis class O [Source:HGNC Symbol;Acc:HGNC:23215] |
| 4978 | OPCML | opioid binding protein/cell adhesion molecule like [Source:HGNC Symbol;Acc:HGNC:8143] |
| 53942 | CNTN5 | contactin 5 [Source:HGNC Symbol;Acc:HGNC:2175] |
| 11181 | TREH | trehalase [Source:HGNC Symbol;Acc:HGNC:12266] |
| 1950 | EGF | epidermal growth factor [Source:HGNC Symbol;Acc:HGNC:3229] |
| 6414 | SELENOP | selenoprotein P [Source:HGNC Symbol;Acc:HGNC:10751] |
| 5087 | PBX1 | PBX homeobox 1 [Source:HGNC Symbol;Acc:HGNC:8632] |
| 6640 | SNTA1 | syntrophin alpha 1 [Source:HGNC Symbol;Acc:HGNC:11167] |
| 7259 | TSPYL1 | TSPY like 1 [Source:HGNC Symbol;Acc:HGNC:12382] |
| 259217 | HSPA12A | heat shock protein family A (Hsp70) member 12A [Source:HGNC Symbol;Acc:HGNC:19022] |
| 5071 | PRKN | parkin RBR E3 ubiquitin protein ligase [Source:HGNC Symbol;Acc:HGNC:8607] |
| 1630 | DCC | DCC netrin 1 receptor [Source:HGNC Symbol;Acc:HGNC:2701] |
| 10686 | CLDN16 | claudin 16 [Source:HGNC Symbol;Acc:HGNC:2037] |
| 2620 | GAS2 | growth arrest specific 2 [Source:HGNC Symbol;Acc:HGNC:4167] |
| 1677 | DFFB | DNA fragmentation factor subunit beta [Source:HGNC Symbol;Acc:HGNC:2773] |
| 8972 | MGAM | maltase-glucoamylase [Source:HGNC Symbol;Acc:HGNC:7043] |
| 2254 | FGF9 | fibroblast growth factor 9 [Source:HGNC Symbol;Acc:HGNC:3687] |
| 23769 | FLRT1 | fibronectin leucine rich transmembrane protein 1 [Source:HGNC Symbol;Acc:HGNC:3760] |
| 1523 | CUX1 | cut like homeobox 1 [Source:HGNC Symbol;Acc:HGNC:2557] |
| 89953 | KLC4 | kinesin light chain 4 [Source:HGNC Symbol;Acc:HGNC:21624] |
| 2734 | GLG1 | golgi glycoprotein 1 [Source:HGNC Symbol;Acc:HGNC:4316] |
| 83953 | FCAMR | Fc fragment of IgA and IgM receptor [Source:HGNC Symbol;Acc:HGNC:24692] |
| 9700 | ESPL1 | extra spindle pole bodies like 1, separase [Source:HGNC Symbol;Acc:HGNC:16856] |
| 63923 | TNN | tenascin N [Source:HGNC Symbol;Acc:HGNC:22942] |
| 1101 | CHAD | chondroadherin [Source:HGNC Symbol;Acc:HGNC:1909] |
| 10788 | IQGAP2 | IQ motif containing GTPase activating protein 2 [Source:HGNC Symbol;Acc:HGNC:6111] |
| 83478 | ARHGAP24 | Rho GTPase activating protein 24 [Source:HGNC Symbol;Acc:HGNC:25361] |
| 9754 | STARD8 | StAR related lipid transfer domain containing 8 [Source:HGNC Symbol;Acc:HGNC:19161] |
| 84144 | SYDE2 | synapse defective Rho GTPase homolog 2 [Source:HGNC Symbol;Acc:HGNC:25841] |
| 6242 | RTKN | rhotekin [Source:HGNC Symbol;Acc:HGNC:10466] |
| 2348 | FOLR1 | folate receptor alpha [Source:HGNC Symbol;Acc:HGNC:3791] |
| 25928 | SOSTDC1 | sclerostin domain containing 1 [Source:HGNC Symbol;Acc:HGNC:21748] |
| 5596 | MAPK4 | mitogen-activated protein kinase 4 [Source:HGNC Symbol;Acc:HGNC:6878] |
| 7429 | VIL1 | villin 1 [Source:HGNC Symbol;Acc:HGNC:12690] |
| 2525 | FUT3 | fucosyltransferase 3 (Lewis blood group) [Source:HGNC Symbol;Acc:HGNC:4014] |
| 2528 | FUT6 | fucosyltransferase 6 [Source:HGNC Symbol;Acc:HGNC:4017] |
| 9942 | XYLB | xylulokinase [Source:HGNC Symbol;Acc:HGNC:12839] |
| 23729 | SHPK | sedoheptulokinase [Source:HGNC Symbol;Acc:HGNC:1492] |
| 23189 | KANK1 | KN motif and ankyrin repeat domains 1 [Source:HGNC Symbol;Acc:HGNC:19309] |
| 255626 | H2BC1 | H2B clustered histone 1 [Source:HGNC Symbol;Acc:HGNC:18730] |
| 956 | ENTPD3 | ectonucleoside triphosphate diphosphohydrolase 3 [Source:HGNC Symbol;Acc:HGNC:3365] |
| 10463 | SLC30A9 | solute carrier family 30 member 9 [Source:HGNC Symbol;Acc:HGNC:1329] |
| 91750 | LIN52 | lin-52 DREAM MuvB core complex component [Source:HGNC Symbol;Acc:HGNC:19856] |
| 203447 | NRK | Nik related kinase [Source:HGNC Symbol;Acc:HGNC:25391] |
| 2690 | GHR | growth hormone receptor [Source:HGNC Symbol;Acc:HGNC:4263] |
| 83660 | TLN2 | talin 2 [Source:HGNC Symbol;Acc:HGNC:15447] |
| 378708 | CENPS | centromere protein S [Source:HGNC Symbol;Acc:HGNC:23163] |
| 2176 | FANCC | FA complementation group C [Source:HGNC Symbol;Acc:HGNC:3584] |
| 22909 | FAN1 | FANCD2 and FANCI associated nuclease 1 [Source:HGNC Symbol;Acc:HGNC:29170] |
| 9620 | CELSR1 | cadherin EGF LAG seven-pass G-type receptor 1 [Source:HGNC Symbol;Acc:HGNC:1850] |
| 161823 | ADAL | adenosine deaminase like [Source:HGNC Symbol;Acc:HGNC:31853] |
| 25961 | NUDT13 | nudix hydrolase 13 [Source:HGNC Symbol;Acc:HGNC:18827] |
| 9321 | TRIP11 | thyroid hormone receptor interactor 11 [Source:HGNC Symbol;Acc:HGNC:12305] |

| | | |
|---|---|---|
| 9053 | MAP7 | microtubule associated protein 7 [Source:HGNC Symbol;Acc:HGNC:6869] |
| 4902 | NRTN | neurturin [Source:HGNC Symbol;Acc:HGNC:8007] |
| 170691 | ADAMTS17 | ADAM metallopeptidase with thrombospondin type 1 motif 17 [Source:HGNC Symbol;Acc:HGNC:17109] |
| 84078 | KBTBD7 | kelch repeat and BTB domain containing 7 [Source:HGNC Symbol;Acc:HGNC:25266] |
| 255631 | COL24A1 | collagen type XXIV alpha 1 chain [Source:HGNC Symbol;Acc:HGNC:20821] |
| 1310 | COL19A1 | collagen type XIX alpha 1 chain [Source:HGNC Symbol;Acc:HGNC:2196] |
| 345456 | PFN3 | profilin 3 [Source:HGNC Symbol;Acc:HGNC:18627] |
| 27109 | DMAC2L | distal membrane arm assembly complex 2 like [Source:HGNC Symbol;Acc:HGNC:18799] |
| 80224 | NUBPL | nucleotide binding protein like [Source:HGNC Symbol;Acc:HGNC:20278] |
| 93058 | COQ10A | coenzyme Q10A [Source:HGNC Symbol;Acc:HGNC:26515] |
| 54829 | ASPN | asporin [Source:HGNC Symbol;Acc:HGNC:14872] |
| 84620 | ST6GAL2 | ST6 beta-galactoside alpha-2,6-sialyltransferase 2 [Source:HGNC Symbol;Acc:HGNC:10861] |
| 84861 | KLHL22 | kelch like family member 22 [Source:HGNC Symbol;Acc:HGNC:25888] |
| 140462 | ASB9 | ankyrin repeat and SOCS box containing 9 [Source:HGNC Symbol;Acc:HGNC:17184] |
| 154881 | KCTD7 | potassium channel tetramerization domain containing 7 [Source:HGNC Symbol;Acc:HGNC:21957] |
| 26249 | KLHL3 | kelch like family member 3 [Source:HGNC Symbol;Acc:HGNC:6354] |
| 285231 | FBXW12 | F-box and WD repeat domain containing 12 [Source:HGNC Symbol;Acc:HGNC:20729] |
| 89890 | KBTBD6 | kelch repeat and BTB domain containing 6 [Source:HGNC Symbol;Acc:HGNC:25340] |
| 147111 | NOTUM | notum, palmitoleoyl-protein carboxylesterase [Source:HGNC Symbol;Acc:HGNC:27106] |
| 63917 | GALNT11 | polypeptide N-acetylgalactosaminyltransferase 11 [Source:HGNC Symbol;Acc:HGNC:19875] |
| 51146 | A4GNT | alpha-1,4-N-acetylglucosaminyltransferase [Source:HGNC Symbol;Acc:HGNC:17968] |
| 282969 | FUOM | fucose mutarotase [Source:HGNC Symbol;Acc:HGNC:24733] |
| 287 | ANK2 | ankyrin 2 [Source:HGNC Symbol;Acc:HGNC:493] |
| 174 | AFP | alpha fetoprotein [Source:HGNC Symbol;Acc:HGNC:317] |
| 79694 | MANEA | mannosidase endo-alpha [Source:HGNC Symbol;Acc:HGNC:21072] |
| 10869 | USP19 | ubiquitin specific peptidase 19 [Source:HGNC Symbol;Acc:HGNC:12617] |
| 9189 | ZBED1 | zinc finger BED-type containing 1 [Source:HGNC Symbol;Acc:HGNC:447] |
| 22981 | NINL | ninein like [Source:HGNC Symbol;Acc:HGNC:29163] |
| 9662 | CEP135 | centrosomal protein 135 [Source:HGNC Symbol;Acc:HGNC:29086] |
| 9727 | RAB11FIP3 | RAB11 family interacting protein 3 [Source:HGNC Symbol;Acc:HGNC:17224] |
| 79738 | BBS10 | Bardet-Biedl syndrome 10 [Source:HGNC Symbol;Acc:HGNC:26291] |
| 55735 | DNAJC11 | DnaJ heat shock protein family (Hsp40) member C11 [Source:HGNC Symbol;Acc:HGNC:25570] |
| 340348 | TSPAN33 | tetraspanin 33 [Source:HGNC Symbol;Acc:HGNC:28743] |
| 9623 | TCL1B | TCL1 family AKT coactivator B [Source:HGNC Symbol;Acc:HGNC:11649] |
| 128977 | C22orf39 | chromosome 22 open reading frame 39 [Source:HGNC Symbol;Acc:HGNC:27012] |
| 8925 | HERC1 | HECT and RLD domain containing E3 ubiquitin protein ligase family member 1 [Source:HGNC Symbol;Acc:HGNC:4867] |
| 11197 | WIF1 | WNT inhibitory factor 1 [Source:HGNC Symbol;Acc:HGNC:18081] |
| 6096 | RORB | RAR related orphan receptor B [Source:HGNC Symbol;Acc:HGNC:10259] |
| 2104 | ESRRG | estrogen related receptor gamma [Source:HGNC Symbol;Acc:HGNC:3474] |
| 80031 | SEMA6D | semaphorin 6D [Source:HGNC Symbol;Acc:HGNC:16770] |
| 10500 | SEMA6C | semaphorin 6C [Source:HGNC Symbol;Acc:HGNC:10740] |
| 3578 | IL9 | interleukin 9 [Source:HGNC Symbol;Acc:HGNC:6029] |
| 9308 | CD83 | CD83 molecule [Source:HGNC Symbol;Acc:HGNC:1703] |
| 84070 | FAM186B | family with sequence similarity 186 member B [Source:HGNC Symbol;Acc:HGNC:25296] |
| 8456 | FOXN1 | forkhead box N1 [Source:HGNC Symbol;Acc:HGNC:12765] |
| 23122 | CLASP2 | cytoplasmic linker associated protein 2 [Source:HGNC Symbol;Acc:HGNC:17078] |
| 84059 | ADGRV1 | adhesion G protein-coupled receptor V1 [Source:HGNC Symbol;Acc:HGNC:17416] |
| 11189 | CELF3 | CUGBP Elav-like family member 3 [Source:HGNC Symbol;Acc:HGNC:11967] |
| 4437 | MSH3 | mutS homolog 3 [Source:HGNC Symbol;Acc:HGNC:7326] |
| 55703 | POLR3B | RNA polymerase III subunit B [Source:HGNC Symbol;Acc:HGNC:30348] |
| 4053 | LTBP2 | latent transforming growth factor beta binding protein 2 [Source:HGNC Symbol;Acc:HGNC:6715] |
| 7783 | ZP2 | zona pellucida glycoprotein 2 [Source:HGNC Symbol;Acc:HGNC:13188] |
| 1473 | CST5 | cystatin D [Source:HGNC Symbol;Acc:HGNC:2477] |
| 3626 | INHBC | inhibin subunit beta C [Source:HGNC Symbol;Acc:HGNC:6068] |
| 55504 | TNFRSF19 | TNF receptor superfamily member 19 [Source:HGNC Symbol;Acc:HGNC:11915] |
| 127069 | OR2T10 | olfactory receptor family 2 subfamily T member 10 [Source:HGNC Symbol;Acc:HGNC:19573] |
| 127077 | OR2T11 | olfactory receptor family 2 subfamily T member 11 [Source:HGNC Symbol;Acc:HGNC:19574] |
| 391211 | OR2G6 | olfactory receptor family 2 subfamily G member 6 [Source:HGNC Symbol;Acc:HGNC:27019] |
| 401992 | OR2T2 | olfactory receptor family 2 subfamily T member 2 [Source:HGNC Symbol;Acc:HGNC:14725] |
| 403244 | OR2T35 | olfactory receptor family 2 subfamily T member 35 [Source:HGNC Symbol;Acc:HGNC:31257] |
| 78989 | COLEC11 | collectin subfamily member 11 [Source:HGNC Symbol;Acc:HGNC:17213] |
| 9315 | NREP | neuronal regeneration related protein [Source:HGNC Symbol;Acc:HGNC:16834] |
| 1272 | CNTN1 | contactin 1 [Source:HGNC Symbol;Acc:HGNC:2171] |
| 4613 | MYCN | MYCN proto-oncogene, bHLH transcription factor [Source:HGNC Symbol;Acc:HGNC:7559] |
| 5178 | PEG3 | paternally expressed 3 [Source:HGNC Symbol;Acc:HGNC:8826] |
| 5334 | PLCL1 | phospholipase C like 1 (inactive) [Source:HGNC Symbol;Acc:HGNC:9063] |
| 64839 | FBXL17 | F-box and leucine rich repeat protein 17 [Source:HGNC Symbol;Acc:HGNC:13615] |
| 9651 | PLCH2 | phospholipase C eta 2 [Source:HGNC Symbol;Acc:HGNC:29037] |
| 26047 | CNTNAP2 | contactin associated protein 2 [Source:HGNC Symbol;Acc:HGNC:13830] |
| 57224 | NHSL1 | NHS like 1 [Source:HGNC Symbol;Acc:HGNC:21021] |
| 55572 | FOXRED1 | FAD dependent oxidoreductase domain containing 1 [Source:HGNC Symbol;Acc:HGNC:26927] |
| 23228 | PLCL2 | phospholipase C like 2 [Source:HGNC Symbol;Acc:HGNC:9064] |
| 5892 | RAD51D | RAD51 paralog D [Source:HGNC Symbol;Acc:HGNC:9823] |
| 9172 | MYOM2 | myomesin 2 [Source:HGNC Symbol;Acc:HGNC:7614] |
| 101928143 | AC005280.2 | novel transcript |
| 113277 | TMEM106A | transmembrane protein 106A [Source:HGNC Symbol;Acc:HGNC:28288] |
| 3434 | IFIT1 | interferon induced protein with tetratricopeptide repeats 1 [Source:HGNC Symbol;Acc:HGNC:5407] |
| 1016 | CDH18 | cadherin 18 [Source:HGNC Symbol;Acc:HGNC:1757] |
| 494551 | WEE2 | WEE2 oocyte meiosis inhibiting kinase [Source:HGNC Symbol;Acc:HGNC:19684] |
| 25840 | METTL7A | methyltransferase like 7A [Source:HGNC Symbol;Acc:HGNC:24550] |
| 84514 | GHDC | GH3 domain containing [Source:HGNC Symbol;Acc:HGNC:24438] |
| 55600 | ITLN1 | intelectin 1 [Source:HGNC Symbol;Acc:HGNC:18259] |
| 26053 | AUTS2 | activator of transcription and developmental regulator AUTS2 [Source:HGNC Symbol;Acc:HGNC:14262] |
| 392255 | GDF6 | growth differentiation factor 6 [Source:HGNC Symbol;Acc:HGNC:4221] |
| 25989 | ULK3 | unc-51 like kinase 3 [Source:HGNC Symbol;Acc:HGNC:19703] |
| 22873 | DZIP1 | DAZ interacting zinc finger protein 1 [Source:HGNC Symbol;Acc:HGNC:20908] |
| 23301 | EHBP1 | EH domain binding protein 1 [Source:HGNC Symbol;Acc:HGNC:29144] |
| 23072 | HECW1 | HECT, C2 and WW domain containing E3 ubiquitin protein ligase 1 [Source:HGNC Symbol;Acc:HGNC:22195] |
| 55840 | EAF2 | ELL associated factor 2 [Source:HGNC Symbol;Acc:HGNC:23115] |
| 121274 | ZNF641 | zinc finger protein 641 [Source:HGNC Symbol;Acc:HGNC:31834] |
| 155054 | ZNF425 | zinc finger protein 425 [Source:HGNC Symbol;Acc:HGNC:20690] |
| 219578 | ZNF804B | zinc finger protein 804B [Source:HGNC Symbol;Acc:HGNC:21958] |
| 284443 | ZNF493 | zinc finger protein 493 [Source:HGNC Symbol;Acc:HGNC:23708] |
| 388536 | ZNF790 | zinc finger protein 790 [Source:HGNC Symbol;Acc:HGNC:33114] |
| 440519 | ZNF724 | zinc finger protein 724 [Source:HGNC Symbol;Acc:HGNC:32460] |
| 619279 | ZNF704 | zinc finger protein 704 [Source:HGNC Symbol;Acc:HGNC:32291] |
| 90075 | ZNF30 | zinc finger protein 30 [Source:HGNC Symbol;Acc:HGNC:13090] |
| 51028 | VPS36 | vacuolar protein sorting 36 homolog [Source:HGNC Symbol;Acc:HGNC:20312] |
| 163486 | DENND1B | DENN domain containing 1B [Source:HGNC Symbol;Acc:HGNC:28404] |
| 54662 | TBC1D13 | TBC1 domain family member 13 [Source:HGNC Symbol;Acc:HGNC:25571] |
| 1842 | ECM2 | extracellular matrix protein 2 [Source:HGNC Symbol;Acc:HGNC:3154] |
| 2312 | FLG | filaggrin [Source:HGNC Symbol;Acc:HGNC:3748] |
| 55364 | IMPACT | impact RWD domain protein [Source:HGNC Symbol;Acc:HGNC:20387] |
| 8663 | EIF3C | eukaryotic translation initiation factor 3 subunit C [Source:HGNC Symbol;Acc:HGNC:3279] |
| 9724 | UTP14C | UTP14C small subunit processome component [Source:HGNC Symbol;Acc:HGNC:20321] |
| 8110 | DPF3 | double PHD fingers 3 [Source:HGNC Symbol;Acc:HGNC:17427] |
| 2135 | EXTL2 | exostosin like glycosyltransferase 2 [Source:HGNC Symbol;Acc:HGNC:3516] |
| 1620 | BRINP1 | BMP/retinoic acid inducible neural specific 1 [Source:HGNC Symbol;Acc:HGNC:2687] |

| | | |
|---|---|---|
| 23742 | NPAP1 | nuclear pore associated protein 1 [Source:HGNC Symbol;Acc:HGNC:1190] |
| 90485 | ZNF835 | zinc finger protein 835 [Source:HGNC Symbol;Acc:HGNC:34332] |
| 51421 | AMOTL2 | angiomotin like 2 [Source:HGNC Symbol;Acc:HGNC:17812] |
| 90624 | LYRM7 | LYR motif containing 7 [Source:HGNC Symbol;Acc:HGNC:28072] |
| 101978719 | LINC00970 | long intergenic non-protein coding RNA 970 [Source:HGNC Symbol;Acc:HGNC:48730] |
| 100128191 | TMPO-AS1 | TMPO antisense RNA 1 [Source:HGNC Symbol;Acc:HGNC:44158] |
| 100128494 | AP002490.1 | novel transcript |
| 100128590 | SLC8A1-AS1 | SLC8A1 antisense RNA 1 [Source:HGNC Symbol;Acc:HGNC:44102] |
| 100128977 | MAPT-AS1 | MAPT antisense RNA 1 [Source:HGNC Symbol;Acc:HGNC:43738] |
| 100129029 | PKP4-AS1 | PKP4 antisense RNA 1 [Source:HGNC Symbol;Acc:HGNC:52580] |
| 100129427 | DDC-AS1 | DDC antisense RNA 1 [Source:HGNC Symbol;Acc:HGNC:40171] |
| 100129654 | TCF24 | transcription factor 24 [Source:HGNC Symbol;Acc:HGNC:32275] |
| 100129792 | CCDC152 | coiled-coil domain containing 152 [Source:HGNC Symbol;Acc:HGNC:34438] |
| 100129969 | FAM205C | family with sequence similarity 205 member C [Source:HGNC Symbol;Acc:HGNC:42673] |
| 100130587 | AL121827.1 | novel transcript, antisense to CHRNA4 |
| 100131193 | CCDC183-AS1 | CCDC183 antisense RNA 1 [Source:HGNC Symbol;Acc:HGNC:44105] |
| 100131244 | ANKRD63 | ankyrin repeat domain 63 [Source:HGNC Symbol;Acc:HGNC:40027] |
| 100132147 | LINC01783 | long intergenic non-protein coding RNA 1783 [Source:HGNC Symbol;Acc:HGNC:52573] |
| 100133985 | LINC01816 | long intergenic non-protein coding RNA 1816 [Source:HGNC Symbol;Acc:HGNC:52621] |
| 100144748 | KLLN | killin, p53 regulated DNA replication inhibitor [Source:HGNC Symbol;Acc:HGNC:37212] |
| 100189589 | DCTN1-AS1 | DCTN1 antisense RNA 1 [Source:HGNC Symbol;Acc:HGNC:44151] |
| 100271702 | LINC00940 | long intergenic non-protein coding RNA 940 [Source:HGNC Symbol;Acc:HGNC:48634] |
| 100287049 | AL139011.1 | novel transcript |
| 100288801 | FRG2C | FSHD region gene 2 family member C [Source:HGNC Symbol;Acc:HGNC:33626] |
| 100288842 | B3GNT10 | UDP-GlcNAc:betaGal beta-1,3-N-acetylglucosaminyltransferase 10 (putative) [Source:HGNC Symbol;Acc:HGNC:53652] |
| 100303491 | ZEB2-AS1 | ZEB2 antisense RNA 1 [Source:HGNC Symbol;Acc:HGNC:37149] |
| 100329135 | TRPC5OS | TRPC5 opposite strand [Source:HGNC Symbol;Acc:HGNC:40593] |
| 100379571 | BACE1-AS | BACE1 antisense RNA [Source:HGNC Symbol;Acc:HGNC:37125] |
| 100505636 | MALINC1 | mitosis associated long intergenic non-coding RNA 1 [Source:HGNC Symbol;Acc:HGNC:49009] |
| 100505676 | RRS1-AS1 | RRS1 antisense RNA 1 (head to head) [Source:HGNC Symbol;Acc:HGNC:50465] |
| 100505678 | STARD4-AS1 | STARD4 antisense RNA 1 [Source:HGNC Symbol;Acc:HGNC:44117] |
| 100505967 | LINC00645 | long intergenic non-protein coding RNA 645 [Source:HGNC Symbol;Acc:HGNC:44299] |
| 100505985 | AL138826.1 | uncharacterized LOC100505985 [Source:NCBI gene (formerly Entrezgene);Acc:100505985] |
| 100506007 | RTCA-AS1 | RTCA antisense RNA 1 [Source:HGNC Symbol;Acc:HGNC:50573] |
| 100506013 | APELA | apelin receptor early endogenous ligand [Source:HGNC Symbol;Acc:HGNC:48925] |
| 100506826 | MYLK-AS1 | MYLK antisense RNA 1 [Source:HGNC Symbol;Acc:HGNC:42440] |
| 100506895 | LUARIS | lncRNA upregulator of antiviral response interferon signaling [Source:HGNC Symbol;Acc:HGNC:40992] |
| 100507027 | MRLN | myoregulin [Source:HGNC Symbol;Acc:HGNC:48649] |
| 100507058 | LINC00845 | long intergenic non-protein coding RNA 845 [Source:HGNC Symbol;Acc:HGNC:45033] |
| 100507203 | SMLR1 | small leucine rich protein 1 [Source:HGNC Symbol;Acc:HGNC:44670] |
| 100507283 | AP000866.1 | novel transcript, antisense to C11orf61 |
| 100507437 | AL928654.1 | novel transcript, antisense to MTA1 |
| 100507466 | DPH6-DT | DPH6 divergent transcript [Source:HGNC Symbol;Acc:HGNC:44147] |
| 100507650 | RNF212B | ring finger protein 212B [Source:HGNC Symbol;Acc:HGNC:20438] |
| 100507657 | LINC02554 | long intergenic non-protein coding RNA 2554 [Source:HGNC Symbol;Acc:HGNC:53594] |
| 100526739 | CENPS-CORT | CENPS-CORT readthrough [Source:HGNC Symbol;Acc:HGNC:38843] |
| 100526760 | ABHD14A-ACY1 | ABHD14A-ACY1 readthrough [Source:HGNC Symbol;Acc:HGNC:38856] |
| 100526771 | SMIM35 | small integral membrane protein 35 [Source:HGNC Symbol;Acc:HGNC:44179] |
| 100526835 | FPGT-TNNI3K | FPGT-TNNI3K readthrough [Source:HGNC Symbol;Acc:HGNC:42952] |
| 100528007 | BORCS7-ASMT | BORCS7-ASMT readthrough (NMD candidate) [Source:HGNC Symbol;Acc:HGNC:49183] |
| 100616530 | C8orf37-AS1 | C8orf37 antisense RNA 1 [Source:HGNC Symbol;Acc:HGNC:50444] |
| 100652999 | AC023509.2 | novel transcript, antisense to ATF7 |
| 100775107 | RBM5-AS1 | RBM5 antisense RNA 1 [Source:HGNC Symbol;Acc:HGNC:48871] |
| 100861507 | FTCD-AS1 | FTCD antisense RNA 1 [Source:HGNC Symbol;Acc:HGNC:40243] |
| 100861519 | GDNF-AS1 | GDNF antisense RNA 1 [Source:HGNC Symbol;Acc:HGNC:43592] |
| 100861540 | CYP3A7-CYP3A51P | CYP3A7-CYP3A51P readthrough [Source:HGNC Symbol;Acc:HGNC:51504] |
| 100873935 | MTOR-AS1 | MTOR antisense RNA 1 [Source:HGNC Symbol;Acc:HGNC:40242] |
| 100873975 | CNTN4-AS1 | CNTN4 antisense RNA 1 [Source:HGNC Symbol;Acc:HGNC:39985] |
| 100873980 | RERG-AS1 | RERG antisense RNA 1 [Source:HGNC Symbol;Acc:HGNC:39990] |
| 100873984 | CHRM3-AS1 | CHRM3 antisense RNA 1 [Source:HGNC Symbol;Acc:HGNC:40150] |
| 100874006 | TTC3-AS1 | TTC3 antisense RNA 1 [Source:HGNC Symbol;Acc:HGNC:40595] |
| 100874024 | TRPC7-AS1 | TRPC7 antisense RNA 1 [Source:HGNC Symbol;Acc:HGNC:40936] |
| 100874238 | ARHGEF7-AS2 | ARHGEF7 antisense RNA 2 [Source:HGNC Symbol;Acc:HGNC:40717] |
| 100874261 | CCDC200 | coiled-coil domain containing 200 [Source:HGNC Symbol;Acc:HGNC:43658] |
| 100885776 | UGDH-AS1 | UGDH antisense RNA 1 [Source:HGNC Symbol;Acc:HGNC:40601] |
| 100886964 | PTCSC3 | papillary thyroid carcinoma susceptibility candidate 3 [Source:HGNC Symbol;Acc:HGNC:43959] |
| 100996669 | LINC01544 | long intergenic non-protein coding RNA 1544 [Source:HGNC Symbol;Acc:HGNC:51312] |
| 10100 | TSPAN2 | tetraspanin 2 [Source:HGNC Symbol;Acc:HGNC:20659] |
| 101926906 | SLC16A12-AS1 | SLC16A12 antisense RNA 1 [Source:HGNC Symbol;Acc:HGNC:51205] |
| 101926967 | AC004596.1 | novel transcript, antisense to ATXN7L3 |
| 101927249 | PDZRN3-AS1 | PDZRN3 antisense RNA 1 [Source:HGNC Symbol;Acc:HGNC:40814] |
| 101927347 | GRM7-AS3 | GRM7 antisense RNA 3 [Source:HGNC Symbol;Acc:HGNC:42444] |
| 101927477 | LINC01507 | long intergenic non-protein coding RNA 1507 [Source:HGNC Symbol;Acc:HGNC:51189] |
| 101927574 | LINC01186 | long intergenic non-protein coding RNA 1186 [Source:HGNC Symbol;Acc:HGNC:49573] |
| 101927702 | AL157756.1 | novel transcript, antisense to PPM1A |
| 101927745 | AJ009632.2 | novel transcript |
| 101927828 | OVOL1-AS1 | OVOL1 antisense RNA 1 [Source:HGNC Symbol;Acc:HGNC:49319] |
| 101927911 | AC015921.1 | novel transcript, antisense to RAP1GAP2 |
| 101927992 | LINC01213 | long intergenic non-protein coding RNA 1213 [Source:HGNC Symbol;Acc:HGNC:49648] |
| 101928099 | PCAT7 | prostate cancer associated transcript 7 [Source:HGNC Symbol;Acc:HGNC:48824] |
| 101928123 | AC004846.1 | novel transcript, antisense to PAPLN |
| 101928154 | LINC01485 | long intergenic non-protein coding RNA 1485 [Source:HGNC Symbol;Acc:HGNC:51138] |
| 101928377 | AF111167.2 | novel transcript, antisense to JDP2 |
| 101928470 | TMEM246-AS1 | TMEM246 antisense RNA 1 [Source:HGNC Symbol;Acc:HGNC:51191] |
| 101928578 | RAP2C-AS1 | RAP2C antisense RNA 1 [Source:HGNC Symbol;Acc:HGNC:40957] |
| 101928596 | AL031726.1 | novel transcript |
| 101928656 | LINC01099 | long intergenic non-protein coding RNA 1099 [Source:HGNC Symbol;Acc:HGNC:49222] |
| 101928781 | TMEM26-AS1 | TMEM26 antisense RNA 1 [Source:HGNC Symbol;Acc:HGNC:51209] |
| 101928844 | AC024592.2 | novel transcript, antisense to FUT3 |
| 101929086 | LINC01447 | long intergenic non-protein coding RNA 1447 [Source:HGNC Symbol;Acc:HGNC:50783] |
| 101929140 | AL354696.2 | novel transcript, antisense to KBTBD7 and MTRF1 |
| 101929265 | LINC01428 | long intergenic non-protein coding RNA 1428 [Source:HGNC Symbol;Acc:HGNC:50738] |
| 101929420 | AL606970.3 | novel transcript |
| 101929469 | TEX52 | testis expressed 52 [Source:HGNC Symbol;Acc:HGNC:53643] |
| 101929625 | LINC01727 | long intergenic non-protein coding RNA 1727 [Source:HGNC Symbol;Acc:HGNC:52515] |
| 101929718 | AL161668.3 | novel transcript, antisense to SLC39A2 |
| 101929733 | LINC01954 | long intergenic non-protein coding RNA 1954 [Source:HGNC Symbol;Acc:HGNC:52779] |
| 101929745 | LINC02119 | long intergenic non-protein coding RNA 2119 [Source:HGNC Symbol;Acc:HGNC:52975] |
| 101929926 | TEX51 | testis expressed 51 [Source:HGNC Symbol;Acc:HGNC:52387] |
| 101930085 | AC018647.2 | novel transcript |
| 102216342 | LINC01108 | long intergenic non-protein coding RNA 1108 [Source:HGNC Symbol;Acc:HGNC:49234] |
| 102467214 | LINC02200 | long intergenic non-protein coding RNA 2200 [Source:HGNC Symbol;Acc:HGNC:53066] |
| 102723701 | AC138356.1 | novel transcript, antisense to ERLIN2 |
| 102724153 | AC007182.1 | novel transcript, antisense to FLVCR2 |
| 102724265 | AP000439.2 | uncharacterized LOC102724265 [Source:NCBI gene (formerly Entrezgene);Acc:102724265] |
| 102724474 | AL137002.2 | WAS/WASL-interacting protein family member 2-like [Source:NCBI gene (formerly Entrezgene);Acc:102724474] |
| 103689918 | LINC01378 | long intergenic non-protein coding RNA 1378 [Source:HGNC Symbol;Acc:HGNC:50645] |
| 103752554 | MYCNUT | MYCN upstream transcript [Source:HGNC Symbol;Acc:HGNC:32684] |

| | | |
|---|---|---|
| 10408 | MYCNOS | MYCN opposite strand [Source:HGNC Symbol;Acc:HGNC:16911] |
| 104355295 | SGMS1-AS1 | SGMS1 antisense RNA 1 [Source:HGNC Symbol;Acc:HGNC:49683] |
| 105369920 | LINC02410 | long intergenic non-protein coding RNA 2410 [Source:HGNC Symbol;Acc:HGNC:53339] |
| 105370854 | AC103740.1 | uncharacterized LOC105370854 [Source:NCBI gene (formerly Entrezgene);Acc:105370854] |
| 105371485 | AC099684.1 | novel transcript, antisense to RTN4RL1 |
| 105371592 | AC005696.1 | novel transcript, antisense to CLUH |
| 105371795 | AC138150.2 | novel transcript, antisense to HEXIM2 |
| 105373536 | LINC01789 | long intergenic non-protein coding RNA 1789 [Source:HGNC Symbol;Acc:HGNC:52578] |
| 105373537 | LINC01886 | long intergenic non-protein coding RNA 1886 [Source:HGNC Symbol;Acc:HGNC:52705] |
| 105374631 | LINC02121 | long intergenic non-protein coding RNA 2121 [Source:HGNC Symbol;Acc:HGNC:52977] |
| 105374714 | LINC02061 | long intergenic non-protein coding RNA 2061 [Source:HGNC Symbol;Acc:HGNC:52906] |
| 105375115 | AC093627.2 | uncharacterized LOC105375115 [Source:NCBI gene (formerly Entrezgene);Acc:105375115] |
| 105378499 | AC016825.1 | novel transcript, antisense to HSPA12A |
| 105378788 | LINC01758 | long intergenic non-protein coding RNA 1758 [Source:HGNC Symbol;Acc:HGNC:52547] |
| 105378934 | LINC01780 | long intergenic non-protein coding RNA 1780 [Source:HGNC Symbol;Acc:HGNC:52570] |
| 105379030 | AC116345.3 | novel transcript |
| 105664404 | WSPAR | WNT signaling pathway activating non-coding RNA [Source:HGNC Symbol;Acc:HGNC:51639] |
| 10669 | CGREF1 | cell growth regulator with EF-hand domain 1 [Source:HGNC Symbol;Acc:HGNC:16962] |
| 10741 | RBBP9 | RB binding protein 9, serine hydrolase [Source:HGNC Symbol;Acc:HGNC:9892] |
| 107984773 | LINC01852 | long intergenic non-protein coding RNA 1852 [Source:HGNC Symbol;Acc:HGNC:52668] |
| 107985343 | LINC01836 | long intergenic non-protein coding RNA 1836 [Source:HGNC Symbol;Acc:HGNC:52652] |
| 107986818 | FAM237B | family with sequence similarity 237 member B [Source:HGNC Symbol;Acc:HGNC:53217] |
| 10826 | FAXDC2 | fatty acid hydroxylase domain containing 2 [Source:HGNC Symbol;Acc:HGNC:1334] |
| 10927 | SPIN1 | spindlin 1 [Source:HGNC Symbol;Acc:HGNC:11243] |
| 11278 | KLF12 | Kruppel like factor 12 [Source:HGNC Symbol;Acc:HGNC:6346] |
| 113419 | TEX261 | testis expressed 261 [Source:HGNC Symbol;Acc:HGNC:30712] |
| 114796 | PSMG3-AS1 | PSMG3 antisense RNA 1 (head to head) [Source:HGNC Symbol;Acc:HGNC:22230] |
| 114905 | C1QTNF7 | C1q and TNF related 7 [Source:HGNC Symbol;Acc:HGNC:14342] |
| 114987 | WDR31 | WD repeat domain 31 [Source:HGNC Symbol;Acc:HGNC:21421] |
| 115752 | DIS3L | DIS3 like exosome 3'-5' exoribonuclease [Source:HGNC Symbol;Acc:HGNC:28698] |
| 118980 | SFXN2 | sideroflexin 2 [Source:HGNC Symbol;Acc:HGNC:16086] |
| 119545625 | AL022069.1 | novel transcript, antisense to BRP44L |
| 122664 | TPPP2 | tubulin polymerization promoting protein family member 2 [Source:HGNC Symbol;Acc:HGNC:19293] |
| 123103 | KLHL33 | kelch like family member 33 [Source:HGNC Symbol;Acc:HGNC:31952] |
| 124751 | KRBA2 | KRAB-A domain containing 2 [Source:HGNC Symbol;Acc:HGNC:26989] |
| 124790 | HEXIM2 | HEXIM P-TEFb complex subunit 2 [Source:HGNC Symbol;Acc:HGNC:28591] |
| 124936 | CYB5D2 | cytochrome b5 domain containing 2 [Source:HGNC Symbol;Acc:HGNC:28471] |
| 127294 | MYOM3 | myomesin 3 [Source:HGNC Symbol;Acc:HGNC:26679] |
| 128497 | SPATA25 | spermatogenesis associated 25 [Source:HGNC Symbol;Acc:HGNC:16158] |
| 129285 | PPP1R21 | protein phosphatase 1 regulatory subunit 21 [Source:HGNC Symbol;Acc:HGNC:30595] |
| 130574 | LYPD6 | LY6/PLAUR domain containing 6 [Source:HGNC Symbol;Acc:HGNC:28751] |
| 131920 | TMEM207 | transmembrane protein 207 [Source:HGNC Symbol;Acc:HGNC:33705] |
| 134553 | C5orf24 | chromosome 5 open reading frame 24 [Source:HGNC Symbol;Acc:HGNC:26746] |
| 137902 | PXDNL | peroxidasin like [Source:HGNC Symbol;Acc:HGNC:26359] |
| 138046 | RALYL | RALY RNA binding protein like [Source:HGNC Symbol;Acc:HGNC:27036] |
| 138948 | DBH-AS1 | DBH antisense RNA 1 [Source:HGNC Symbol;Acc:HGNC:24155] |
| 140689 | CBLN4 | cerebellin 4 precursor [Source:HGNC Symbol;Acc:HGNC:16231] |
| 140902 | R3HDML | R3H domain containing like [Source:HGNC Symbol;Acc:HGNC:16249] |
| 143941 | TTC36 | tetratricopeptide repeat domain 36 [Source:HGNC Symbol;Acc:HGNC:33708] |
| 144100 | PLEKHA7 | pleckstrin homology domain containing A7 [Source:HGNC Symbol;Acc:HGNC:27049] |
| 145837 | DRAIC | downregulated RNA in cancer, inhibitor of cell invasion and migration [Source:HGNC Symbol;Acc:HGNC:27082] |
| 146562 | C16orf71 | chromosome 16 open reading frame 71 [Source:HGNC Symbol;Acc:HGNC:25081] |
| 146691 | TOM1L2 | target of myb1 like 2 membrane trafficking protein [Source:HGNC Symbol;Acc:HGNC:11984] |
| 148808 | MFSD4A | major facilitator superfamily domain containing 4A [Source:HGNC Symbol;Acc:HGNC:25433] |
| 149465 | CFAP57 | cilia and flagella associated protein 57 [Source:HGNC Symbol;Acc:HGNC:26485] |
| 151126 | ZNF385B | zinc finger protein 385B [Source:HGNC Symbol;Acc:HGNC:26332] |
| 151295 | SLC23A3 | solute carrier family 23 member 3 [Source:HGNC Symbol;Acc:HGNC:20601] |
| 151393 | RMDN2 | regulator of microtubule dynamics 2 [Source:HGNC Symbol;Acc:HGNC:26567] |
| 151516 | ASPRV1 | aspartic peptidase retroviral like 1 [Source:HGNC Symbol;Acc:HGNC:26321] |
| 152404 | IGSF11 | immunoglobulin superfamily member 11 [Source:HGNC Symbol;Acc:HGNC:16669] |
| 153364 | MBLAC2 | metallo-beta-lactamase domain containing 2 [Source:HGNC Symbol;Acc:HGNC:33711] |
| 157567 | ANKRD46 | ankyrin repeat domain 46 [Source:HGNC Symbol;Acc:HGNC:27229] |
| 157983 | DOCK8-AS1 | DOCK8 antisense RNA 1 [Source:HGNC Symbol;Acc:HGNC:26436] |
| 161779 | PGBD4 | piggyBac transposable element derived 4 [Source:HGNC Symbol;Acc:HGNC:19401] |
| 163782 | KANK4 | KN motif and ankyrin repeat domains 4 [Source:HGNC Symbol;Acc:HGNC:27263] |
| 165082 | ADGRF3 | adhesion G protein-coupled receptor F3 [Source:HGNC Symbol;Acc:HGNC:18989] |
| 167681 | PRSS35 | serine protease 35 [Source:HGNC Symbol;Acc:HGNC:21387] |
| 1773 | DNASE1 | deoxyribonuclease 1 [Source:HGNC Symbol;Acc:HGNC:2956] |
| 196740 | VSTM4 | V-set and transmembrane domain containing 4 [Source:HGNC Symbol;Acc:HGNC:26470] |
| 200150 | PLD5 | phospholipase D family member 5 [Source:HGNC Symbol;Acc:HGNC:26879] |
| 201305 | SPNS3 | sphingolipid transporter 3 (putative) [Source:HGNC Symbol;Acc:HGNC:28433] |
| 201617 | LINC00870 | long intergenic non-protein coding RNA 870 [Source:HGNC Symbol;Acc:HGNC:27319] |
| 219347 | TMEM254-AS1 | TMEM254 antisense RNA 1 [Source:HGNC Symbol;Acc:HGNC:27340] |
| 219527 | LRRC55 | leucine rich repeat containing 55 [Source:HGNC Symbol;Acc:HGNC:32324] |
| 219623 | TMEM26 | transmembrane protein 26 [Source:HGNC Symbol;Acc:HGNC:28550] |
| 220963 | SLC16A9 | solute carrier family 16 member 9 [Source:HGNC Symbol;Acc:HGNC:23520] |
| 220980 | TMEM72-AS1 | TMEM72 antisense RNA 1 [Source:HGNC Symbol;Acc:HGNC:27349] |
| 221188 | ADGRG5 | adhesion G protein-coupled receptor G5 [Source:HGNC Symbol;Acc:HGNC:19010] |
| 2217 | FCGRT | Fc fragment of IgG receptor and transporter [Source:HGNC Symbol;Acc:HGNC:3621] |
| 222255 | ATXN7L1 | ataxin 7 like 1 [Source:HGNC Symbol;Acc:HGNC:22210] |
| 22844 | FRMPD1 | FERM and PDZ domain containing 1 [Source:HGNC Symbol;Acc:HGNC:29159] |
| 22982 | DIP2C | disco interacting protein 2 homolog C [Source:HGNC Symbol;Acc:HGNC:29150] |
| 23024 | PDZRN3 | PDZ domain containing ring finger 3 [Source:HGNC Symbol;Acc:HGNC:17704] |
| 23078 | VWA8 | von Willebrand factor A domain containing 8 [Source:HGNC Symbol;Acc:HGNC:29071] |
| 23264 | ZC3H7B | zinc finger CCCH-type containing 7B [Source:HGNC Symbol;Acc:HGNC:30869] |
| 23335 | WDR7 | WD repeat domain 7 [Source:HGNC Symbol;Acc:HGNC:13490] |
| 23341 | DNAJC16 | DnaJ heat shock protein family (Hsp40) member C16 [Source:HGNC Symbol;Acc:HGNC:29157] |
| 23506 | BICRAL | BRD4 interacting chromatin remodeling complex associated protein like [Source:HGNC Symbol;Acc:HGNC:21111] |
| 252995 | FNDC5 | fibronectin type III domain containing 5 [Source:HGNC Symbol;Acc:HGNC:20240] |
| 254778 | VXN | vexin [Source:HGNC Symbol;Acc:HGNC:28498] |
| 255167 | LINC01018 | long intergenic non-protein coding RNA 1018 [Source:HGNC Symbol;Acc:HGNC:27394] |
| 257019 | FRMD3 | FERM domain containing 3 [Source:HGNC Symbol;Acc:HGNC:24125] |
| 257396 | AC008966.1 | novel transcript, antisense to MOCS2 |
| 25849 | PARM1 | prostate androgen-regulated mucin-like protein 1 [Source:HGNC Symbol;Acc:HGNC:24536] |
| 25864 | ABHD14A | abhydrolase domain containing 14A [Source:HGNC Symbol;Acc:HGNC:24538] |
| 25932 | CLIC4 | chloride intracellular channel 4 [Source:HGNC Symbol;Acc:HGNC:13518] |
| 25975 | EGFL6 | EGF like domain multiple 6 [Source:HGNC Symbol;Acc:HGNC:3235] |
| 26080 | FAM230C | family with sequence similarity 230 member C [Source:HGNC Symbol;Acc:HGNC:24482] |
| 26273 | FBXO3 | F-box protein 3 [Source:HGNC Symbol;Acc:HGNC:13582] |
| 283129 | MAJIN | membrane anchored junction protein [Source:HGNC Symbol;Acc:HGNC:27441] |
| 283514 | SIAH3 | siah E3 ubiquitin protein ligase family member 3 [Source:HGNC Symbol;Acc:HGNC:30553] |
| 283518 | KCNRG | potassium channel regulator [Source:HGNC Symbol;Acc:HGNC:18893] |
| 284948 | SH2D6 | SH2 domain containing 6 [Source:HGNC Symbol;Acc:HGNC:30439] |
| 284992 | CCDC150 | coiled-coil domain containing 150 [Source:HGNC Symbol;Acc:HGNC:26834] |
| 285154 | CYP1B1-AS1 | CYP1B1 antisense RNA 1 [Source:HGNC Symbol;Acc:HGNC:28543] |
| 285441 | F11-AS1 | F11 antisense RNA 1 [Source:HGNC Symbol;Acc:HGNC:27725] |
| 285605 | DTWD2 | DTW domain containing 2 [Source:HGNC Symbol;Acc:HGNC:19334] |
| 285848 | PNPLA1 | patatin like phospholipase domain containing 1 [Source:HGNC Symbol;Acc:HGNC:21246] |

| | | |
|---|---|---|
| 286002 | SLC26A4-AS1 | SLC26A4 antisense RNA 1 [Source:HGNC Symbol;Acc:HGNC:22385] |
| 28992 | MACROD1 | mono-ADP ribosylhydrolase 1 [Source:HGNC Symbol;Acc:HGNC:29598] |
| 29114 | TAGLN3 | transgelin 3 [Source:HGNC Symbol;Acc:HGNC:29868] |
| 29953 | TRHDE | thyrotropin releasing hormone degrading enzyme [Source:HGNC Symbol;Acc:HGNC:30748] |
| 338094 | FAM151A | family with sequence similarity 151 member A [Source:HGNC Symbol;Acc:HGNC:25032] |
| 339318 | ZNF181 | zinc finger protein 181 [Source:HGNC Symbol;Acc:HGNC:12971] |
| 339855 | KY | kyphoscoliosis peptidase [Source:HGNC Symbol;Acc:HGNC:26576] |
| 340094 | LINC01020 | long intergenic non-protein coding RNA 1020 [Source:HGNC Symbol;Acc:HGNC:27968] |
| 340595 | RTL4 | retrotransposon Gag like 4 [Source:HGNC Symbol;Acc:HGNC:25214] |
| 341116 | MS4A10 | membrane spanning 4-domains A10 [Source:HGNC Symbol;Acc:HGNC:13368] |
| 342346 | C16orf96 | chromosome 16 open reading frame 96 [Source:HGNC Symbol;Acc:HGNC:40031] |
| 347051 | SLC10A5 | solute carrier family 10 member 5 [Source:HGNC Symbol;Acc:HGNC:22981] |
| 348801 | LNP1 | leukemia NUP98 fusion partner 1 [Source:HGNC Symbol;Acc:HGNC:28014] |
| 375791 | CYSRT1 | cysteine rich tail 1 [Source:HGNC Symbol;Acc:HGNC:30529] |
| 3769 | KCNJ13 | potassium inwardly rectifying channel subfamily J member 13 [Source:HGNC Symbol;Acc:HGNC:6259] |
| 378832 | COL18A1-AS1 | COL18A1 antisense RNA 1 [Source:HGNC Symbol;Acc:HGNC:23132] |
| 387640 | SKIDA1 | SKI/DACH domain containing 1 [Source:HGNC Symbol;Acc:HGNC:32697] |
| 387700 | SLC16A12 | solute carrier family 16 member 12 [Source:HGNC Symbol;Acc:HGNC:23094] |
| 387804 | VSTM5 | V-set and transmembrane domain containing 5 [Source:HGNC Symbol;Acc:HGNC:34443] |
| 388011 | LINC01550 | long intergenic non-protein coding RNA 1550 [Source:HGNC Symbol;Acc:HGNC:20111] |
| 388324 | INCA1 | inhibitor of CDK, cyclin A1 interacting protein 1 [Source:HGNC Symbol;Acc:HGNC:32224] |
| 388595 | TMEM82 | transmembrane protein 82 [Source:HGNC Symbol;Acc:HGNC:32350] |
| 389643 | NUGGC | nuclear GTPase, germinal center associated [Source:HGNC Symbol;Acc:HGNC:33550] |
| 389827 | MYMK | myomaker, myoblast fusion factor [Source:HGNC Symbol;Acc:HGNC:33778] |
| 389941 | C1QL3 | complement C1q like 3 [Source:HGNC Symbol;Acc:HGNC:19359] |
| 399948 | COLCA1 | colorectal cancer associated 1 [Source:HGNC Symbol;Acc:HGNC:33789] |
| 400165 | ATP11AUN | ATP11A upstream neighbor [Source:HGNC Symbol;Acc:HGNC:33793] |
| 400548 | LINC02139 | long intergenic non-protein coding RNA 2139 [Source:HGNC Symbol;Acc:HGNC:52999] |
| 400673 | VMAC | vimentin type intermediate filament associated coiled-coil protein [Source:HGNC Symbol;Acc:HGNC:33803] |
| 401014 | TEX41 | testis expressed 41 [Source:HGNC Symbol;Acc:HGNC:48667] |
| 401081 | C3orf85 | chromosome 3 open reading frame 85 [Source:HGNC Symbol;Acc:HGNC:53432] |
| 401145 | CCSER1 | coiled-coil serine rich protein 1 [Source:HGNC Symbol;Acc:HGNC:29349] |
| 401264 | TRAM2-AS1 | TRAM2 antisense RNA 1 (head to head) [Source:HGNC Symbol;Acc:HGNC:48663] |
| 401548 | SNX30 | sorting nexin family member 30 [Source:HGNC Symbol;Acc:HGNC:23685] |
| 404216 | LINC01561 | long intergenic non-protein coding RNA 1561 [Source:HGNC Symbol;Acc:HGNC:31365] |
| 414328 | IDNK | IDNK gluconokinase [Source:HGNC Symbol;Acc:HGNC:31367] |
| 440556 | PRDM16-DT | PRDM16 divergent transcript [Source:HGNC Symbol;Acc:HGNC:48664] |
| 440590 | ZYG11A | zyg-11 family member A, cell cycle regulator [Source:HGNC Symbol;Acc:HGNC:32058] |
| 441376 | AARD | alanine and arginine rich domain containing protein [Source:HGNC Symbol;Acc:HGNC:33842] |
| 4675 | NAP1L3 | nucleosome assembly protein 1 like 3 [Source:HGNC Symbol;Acc:HGNC:7639] |
| 4745 | NELL1 | neural EGFL like 1 [Source:HGNC Symbol;Acc:HGNC:7750] |
| 4883 | NPR3 | natriuretic peptide receptor 3 [Source:HGNC Symbol;Acc:HGNC:7945] |
| 51062 | ATL1 | atlastin GTPase 1 [Source:HGNC Symbol;Acc:HGNC:11231] |
| 51066 | SSUH2 | ssu-2 homolog [Source:HGNC Symbol;Acc:HGNC:24809] |
| 51086 | TNNI3K | TNNI3 interacting kinase [Source:HGNC Symbol;Acc:HGNC:19661] |
| 51337 | THEM6 | thioesterase superfamily member 6 [Source:HGNC Symbol;Acc:HGNC:29656] |
| 51463 | GPR89B | G protein-coupled receptor 89B [Source:HGNC Symbol;Acc:HGNC:13840] |
| 51557 | LGSN | lengsin, lens protein with glutamine synthetase domain [Source:HGNC Symbol;Acc:HGNC:21016] |
| 5184 | PEPD | peptidase D [Source:HGNC Symbol;Acc:HGNC:8840] |
| 54088 | LINC00113 | long intergenic non-protein coding RNA 113 [Source:HGNC Symbol;Acc:HGNC:1264] |
| 54346 | UNC93A | unc-93 homolog A [Source:HGNC Symbol;Acc:HGNC:12570] |
| 54550 | NECAB2 | N-terminal EF-hand calcium binding protein 2 [Source:HGNC Symbol;Acc:HGNC:23746] |
| 54923 | LIME1 | Lck interacting transmembrane adaptor 1 [Source:HGNC Symbol;Acc:HGNC:26016] |
| 54943 | DNAJC28 | DnaJ heat shock protein family (Hsp40) member C28 [Source:HGNC Symbol;Acc:HGNC:1297] |
| 54993 | ZSCAN2 | zinc finger and SCAN domain containing 2 [Source:HGNC Symbol;Acc:HGNC:20994] |
| 55020 | TTC38 | tetratricopeptide repeat domain 38 [Source:HGNC Symbol;Acc:HGNC:26082] |
| 55068 | ENOX1 | ecto-NOX disulfide-thiol exchanger 1 [Source:HGNC Symbol;Acc:HGNC:25474] |
| 55266 | TMEM19 | transmembrane protein 19 [Source:HGNC Symbol;Acc:HGNC:25605] |
| 55316 | RSAD1 | radical S-adenosyl methionine domain containing 1 [Source:HGNC Symbol;Acc:HGNC:25634] |
| 55333 | SYNJ2BP | synaptojanin 2 binding protein [Source:HGNC Symbol;Acc:HGNC:18955] |
| 554235 | ASPDH | aspartate dehydrogenase domain containing [Source:HGNC Symbol;Acc:HGNC:33856] |
| 55449 | DHRS4-AS1 | DHRS4 antisense RNA 1 [Source:HGNC Symbol;Acc:HGNC:23175] |
| 55530 | SVOP | SV2 related protein [Source:HGNC Symbol;Acc:HGNC:25417] |
| 55612 | FERMT1 | fermitin family member 1 [Source:HGNC Symbol;Acc:HGNC:15889] |
| 56062 | KLHL4 | kelch like family member 4 [Source:HGNC Symbol;Acc:HGNC:6355] |
| 56181 | MTFR1L | mitochondrial fission regulator 1 like [Source:HGNC Symbol;Acc:HGNC:28836] |
| 56241 | SUSD2 | sushi domain containing 2 [Source:HGNC Symbol;Acc:HGNC:30667] |
| 56890 | MDM1 | Mdm1 nuclear protein [Source:HGNC Symbol;Acc:HGNC:29917] |
| 57626 | KLHL1 | kelch like family member 1 [Source:HGNC Symbol;Acc:HGNC:6352] |
| 57685 | CACHD1 | cache domain containing 1 [Source:HGNC Symbol;Acc:HGNC:29314] |
| 613212 | CTXN3 | cortexin 3 [Source:HGNC Symbol;Acc:HGNC:31110] |
| 64081 | PBLD | phenazine biosynthesis like protein domain containing [Source:HGNC Symbol;Acc:HGNC:23301] |
| 642484 | AC006305.1 | novel transcript |
| 643236 | TMEM72 | transmembrane protein 72 [Source:HGNC Symbol;Acc:HGNC:31658] |
| 643596 | RNF224 | ring finger protein 224 [Source:HGNC Symbol;Acc:HGNC:41912] |
| 64478 | CSMD1 | CUB and Sushi multiple domains 1 [Source:HGNC Symbol;Acc:HGNC:14026] |
| 645027 | EVPLL | envoplakin like [Source:HGNC Symbol;Acc:HGNC:35236] |
| 645434 | LINC01625 | long intergenic non-protein coding RNA 1625 [Source:HGNC Symbol;Acc:HGNC:52052] |
| 646019 | CBY3 | chibby family member 3 [Source:HGNC Symbol;Acc:HGNC:33278] |
| 64755 | RUSF1 | RUS family member 1 [Source:HGNC Symbol;Acc:HGNC:25848] |
| 654463 | FER1L6 | fer-1 like family member 6 [Source:HGNC Symbol;Acc:HGNC:28065] |
| 6744 | ITPRID2 | ITPR interacting domain containing 2 [Source:HGNC Symbol;Acc:HGNC:11319] |
| 6887 | TAL2 | TAL bHLH transcription factor 2 [Source:HGNC Symbol;Acc:HGNC:11557] |
| 6920 | TCEA3 | transcription elongation factor A3 [Source:HGNC Symbol;Acc:HGNC:11615] |
| 727944 | LINC01874 | long intergenic non-protein coding RNA 1874 [Source:HGNC Symbol;Acc:HGNC:52693] |
| 728040 | LINC02499 | long intergenic non-protein coding RNA 2499 [Source:HGNC Symbol;Acc:HGNC:53484] |
| 730091 | LINC00886 | long intergenic non-protein coding RNA 886 [Source:HGNC Symbol;Acc:HGNC:48572] |
| 7447 | VSNL1 | visinin like 1 [Source:HGNC Symbol;Acc:HGNC:12722] |
| 7691 | ZNF132 | zinc finger protein 132 [Source:HGNC Symbol;Acc:HGNC:12916] |
| 79000 | AUNIP | aurora kinase A and ninein interacting protein [Source:HGNC Symbol;Acc:HGNC:28363] |
| 79154 | DHRS11 | dehydrogenase/reductase 11 [Source:HGNC Symbol;Acc:HGNC:28639] |
| 79763 | ISOC2 | isochorismatase domain containing 2 [Source:HGNC Symbol;Acc:HGNC:26278] |
| 79783 | SUGCT | succinyl-CoA:glutarate-CoA transferase [Source:HGNC Symbol;Acc:HGNC:16001] |
| 79831 | KDM8 | lysine demethylase 8 [Source:HGNC Symbol;Acc:HGNC:25840] |
| 79896 | THNSL1 | threonine synthase like 1 [Source:HGNC Symbol;Acc:HGNC:26160] |
| 79981 | FRMD1 | FERM domain containing 1 [Source:HGNC Symbol;Acc:HGNC:21240] |
| 80014 | WWC2 | WW and C2 domain containing 2 [Source:HGNC Symbol;Acc:HGNC:24148] |
| 80069 | LINC00574 | long intergenic non-protein coding RNA 574 [Source:HGNC Symbol;Acc:HGNC:21598] |
| 80125 | CCDC33 | coiled-coil domain containing 33 [Source:HGNC Symbol;Acc:HGNC:26552] |
| 80127 | BBOF1 | basal body orientation factor 1 [Source:HGNC Symbol;Acc:HGNC:19855] |
| 80157 | CWH43 | cell wall biogenesis 43 C-terminal homolog [Source:HGNC Symbol;Acc:HGNC:26133] |
| 80206 | FHOD3 | formin homology 2 domain containing 3 [Source:HGNC Symbol;Acc:HGNC:26178] |
| 83876 | MRO | maestro [Source:HGNC Symbol;Acc:HGNC:24121] |
| 84417 | ECRG4 | ECRG4 augurin precursor [Source:HGNC Symbol;Acc:HGNC:24642] |
| 84461 | NEURL4 | neuralized E3 ubiquitin protein ligase 4 [Source:HGNC Symbol;Acc:HGNC:34410] |
| 84530 | SRRM4 | serine/arginine repetitive matrix 4 [Source:HGNC Symbol;Acc:HGNC:29389] |
| 84656 | GLYR1 | glyoxylate reductase 1 homolog [Source:HGNC Symbol;Acc:HGNC:24434] |
| 84898 | PLXDC2 | plexin domain containing 2 [Source:HGNC Symbol;Acc:HGNC:21013] |

| | | |
|---|---|---|
| 84960 | CCDC183 | coiled-coil domain containing 183 [Source:HGNC Symbol;Acc:HGNC:28236] |
| 8521 | GCM1 | glial cells missing transcription factor 1 [Source:HGNC Symbol;Acc:HGNC:4197] |
| 85314 | ERVE-1 | endogenous retrovirus group E member 1 [Source:HGNC Symbol;Acc:HGNC:14602] |
| 8715 | NOL4 | nucleolar protein 4 [Source:HGNC Symbol;Acc:HGNC:7870] |
| 87769 | GGACT | gamma-glutamylamine cyclotransferase [Source:HGNC Symbol;Acc:HGNC:25100] |
| 8814 | CDKL1 | cyclin dependent kinase like 1 [Source:HGNC Symbol;Acc:HGNC:1781] |
| 89758 | LINC01621 | long intergenic non-protein coding RNA 1621 [Source:HGNC Symbol;Acc:HGNC:14109] |
| 93432 | MGAM2 | maltase-glucoamylase 2 (putative) [Source:HGNC Symbol;Acc:HGNC:28101] |
| 9389 | SLC22A14 | solute carrier family 22 member 14 [Source:HGNC Symbol;Acc:HGNC:8495] |
| 9543 | IGDCC3 | immunoglobulin superfamily DCC subclass member 3 [Source:HGNC Symbol;Acc:HGNC:9700] |
| 9717 | SEC14L5 | SEC14 like lipid binding 5 [Source:HGNC Symbol;Acc:HGNC:29032] |
| 9772 | TMEM94 | transmembrane protein 94 [Source:HGNC Symbol;Acc:HGNC:28983] |
| 9865 | TRIL | TLR4 interactor with leucine rich repeats [Source:HGNC Symbol;Acc:HGNC:22200] |
| 987 | LRBA | LPS responsive beige-like anchor protein [Source:HGNC Symbol;Acc:HGNC:1742] |

Bulk RNA-seq DOWN

| FDR q-value |
|---|
| 1.83E-13 |
| 6.82E-13 |
| 4.50E-11 |
| 2.17E-09 |
| 1.04E-08 |
| 7.12E-08 |
| 7.12E-08 |
| 1.04E-07 |
| 1.04E-07 |
| 5.50E-07 |
| 6.85E-07 |
| 9.58E-07 |
| 1.11E-06 |
| 1.57E-06 |
| 1.78E-06 |
| 1.83E-06 |
| 8.13E-06 |
| 1.24E-05 |
| 1.60E-05 |
| 1.60E-05 |
| 3.90E-05 |
| 3.90E-05 |
| 1.72E-04 |
| 1.78E-04 |
| 2.08E-04 |
| 3.21E-04 |
| 4.41E-04 |
| 7.71E-04 |
| 8.63E-04 |
| 1.22E-03 |
| 1.23E-03 |
| 1.31E-03 |
| 1.32E-03 |
| 1.51E-03 |
| 1.74E-03 |
| 1.74E-03 |
| 2.05E-03 |
| 2.34E-03 |
| 2.34E-03 |
| 2.59E-03 |
| 2.60E-03 |
| 2.60E-03 |
| 3.44E-03 |
| 3.46E-03 |
| 3.46E-03 |
| 3.46E-03 |
| 3.57E-03 |
| 3.57E-03 |
| 3.59E-03 |
| 3.59E-03 |
| 3.59E-03 |
| 3.71E-03 |
| 3.71E-03 |
| 3.81E-03 |
| 4.44E-03 |
| 5.05E-03 |
| 6.56E-03 |
| 7.85E-03 |
| 7.87E-03 |
| 8.12E-03 |
| 8.72E-03 |
| 8.72E-03 |
| 8.72E-03 |
| 9.14E-03 |
| 9.21E-03 |
| 9.21E-03 |
| 9.21E-03 |
| 9.21E-03 |
| 1.28E-02 |
| 1.46E-02 |
| 1.55E-02 |
| 1.65E-02 |
| 1.79E-02 |
| 1.79E-02 |
| 1.79E-02 |
| 1.79E-02 |
| 1.89E-02 |
| 2.05E-02 |
| 2.05E-02 |
| 2.05E-02 |
| 2.05E-02 |
| 2.05E-02 |
| 2.21E-02 |
| 2.29E-02 |
| 2.31E-02 |
| 2.31E-02 |
| 2.35E-02 |
| 2.66E-02 |
| 2.66E-02 |
| 2.85E-02 |
| 3.16E-02 |
| 3.96E-02 |
| 4.04E-02 |
| 4.18E-02 |
| 4.18E-02 |

Bulk RNA-seq DOWN

| | | | | | | | | |
|---|---|---|---|---|---|---|---|---|
| REACTOME_SLC_MEDIATED_TRANSMEMBRANE_TRANSPORT | REACTOME_TRANSPORT_OF_INORGANIC_CATIONS_ANIONS_AND_AMINO_ACIDS_OLIGOPEPTIDES | KEGG_GLYCINE_SERINE_AND_THREONINE_METABOLISM | REACTOME_GLYOXYLATE_METABOLISM_AND_GLYCINE_DEGRADATION | HALLMARK_XENOBIOTIC_METABOLISM | KEGG_ARGININE_AND_PROLINE_METABOLISM | WP_AMINO_ACID_METABOLISM | KEGG_PEROXISOME | KEGG_BETA_ALANINE_METABOLISM |
| | | | REACTOME_GLYOXYLATE_METABOLISM_AND_GLYCINE_DEGRADATION | | | | | |
| | | | REACTOME_GLYOXYLATE_METABOLISM_AND_GLYCINE_DEGRADATION | | | | | |
| | | | | HALLMARK_XENOBIOTIC_METABOLISM | | | | |
| | | | | | KEGG_ARGININE_AND_PROLINE_METABOLISM | WP_AMINO_ACID_METABOLISM | | KEGG_BETA_ALANINE_METABOLISM |
| | | | | | | WP_AMINO_ACID_METABOLISM | | KEGG_BETA_ALANINE_METABOLISM |
| | | KEGG_GLYCINE_SERINE_AND_THREONINE_METABOLISM | REACTOME_GLYOXYLATE_METABOLISM_AND_GLYCINE_DEGRADATION | | KEGG_ARGININE_AND_PROLINE_METABOLISM | | KEGG_PEROXISOME | |
| | | KEGG_GLYCINE_SERINE_AND_THREONINE_METABOLISM | REACTOME_GLYOXYLATE_METABOLISM_AND_GLYCINE_DEGRADATION | | | | | |
| | | KEGG_GLYCINE_SERINE_AND_THREONINE_METABOLISM | | | KEGG_ARGININE_AND_PROLINE_METABOLISM | | | |
| | | KEGG_GLYCINE_SERINE_AND_THREONINE_METABOLISM | | | | WP_AMINO_ACID_METABOLISM | | |
| | | KEGG_GLYCINE_SERINE_AND_THREONINE_METABOLISM | | | | | KEGG_PEROXISOME | |
| | | KEGG_GLYCINE_SERINE_AND_THREONINE_METABOLISM | | | | | | |
| | | KEGG_GLYCINE_SERINE_AND_THREONINE_METABOLISM | | | | | | |
| | | KEGG_GLYCINE_SERINE_AND_THREONINE_METABOLISM | | | | | | |
| | | KEGG_GLYCINE_SERINE_AND_THREONINE_METABOLISM | | | | | | |
| | | | REACTOME_GLYOXYLATE_METABOLISM_AND_GLYCINE_DEGRADATION | | KEGG_ARGININE_AND_PROLINE_METABOLISM | | | |
| | | | REACTOME_GLYOXYLATE_METABOLISM_AND_GLYCINE_DEGRADATION | | KEGG_ARGININE_AND_PROLINE_METABOLISM | | | |
| | | | REACTOME_GLYOXYLATE_METABOLISM_AND_GLYCINE_DEGRADATION | | | WP_AMINO_ACID_METABOLISM | | |
| | | | REACTOME_GLYOXYLATE_METABOLISM_AND_GLYCINE_DEGRADATION | | | | KEGG_PEROXISOME | |
| | | | REACTOME_GLYOXYLATE_METABOLISM_AND_GLYCINE_DEGRADATION | | | | | |
| | | | REACTOME_GLYOXYLATE_METABOLISM_AND_GLYCINE_DEGRADATION | | | | | |
| | | | | HALLMARK_XENOBIOTIC_METABOLISM | | WP_AMINO_ACID_METABOLISM | | |
| | | | | HALLMARK_XENOBIOTIC_METABOLISM | | | | |
| | | | | | KEGG_ARGININE_AND_PROLINE_METABOLISM | WP_AMINO_ACID_METABOLISM | | |
| | | | | | KEGG_ARGININE_AND_PROLINE_METABOLISM | | | |
| | | | | | KEGG_ARGININE_AND_PROLINE_METABOLISM | | | |
| | | | | | KEGG_ARGININE_AND_PROLINE_METABOLISM | | | |
| | | | | | | WP_AMINO_ACID_METABOLISM | | |
| | | | | | | WP_AMINO_ACID_METABOLISM | | |
| REACTOME_SLC_MEDIATED_TRANSMEMBRANE_TRANSPORT | REACTOME_TRANSPORT_OF_INORGANIC_CATIONS_ANIONS_AND_AMINO_ACIDS_OLIGOPEPTIDES | | | | | | | |
| REACTOME_SLC_MEDIATED_TRANSMEMBRANE_TRANSPORT | REACTOME_TRANSPORT_OF_INORGANIC_CATIONS_ANIONS_AND_AMINO_ACIDS_OLIGOPEPTIDES | | | | | | | |
| REACTOME_SLC_MEDIATED_TRANSMEMBRANE_TRANSPORT | REACTOME_TRANSPORT_OF_INORGANIC_CATIONS_ANIONS_AND_AMINO_ACIDS_OLIGOPEPTIDES | | | | | | | |
| REACTOME_SLC_MEDIATED_TRANSMEMBRANE_TRANSPORT | REACTOME_TRANSPORT_OF_INORGANIC_CATIONS_ANIONS_AND_AMINO_ACIDS_OLIGOPEPTIDES | | | | | | | |
| REACTOME_SLC_MEDIATED_TRANSMEMBRANE_TRANSPORT | REACTOME_TRANSPORT_OF_INORGANIC_CATIONS_ANIONS_AND_AMINO_ACIDS_OLIGOPEPTIDES | | | | | | | |
| REACTOME_SLC_MEDIATED_TRANSMEMBRANE_TRANSPORT | REACTOME_TRANSPORT_OF_INORGANIC_CATIONS_ANIONS_AND_AMINO_ACIDS_OLIGOPEPTIDES | | | | | | | |
| REACTOME_SLC_MEDIATED_TRANSMEMBRANE_TRANSPORT | REACTOME_TRANSPORT_OF_INORGANIC_CATIONS_ANIONS_AND_AMINO_ACIDS_OLIGOPEPTIDES | | | | | | | |
| REACTOME_SLC_MEDIATED_TRANSMEMBRANE_TRANSPORT | REACTOME_TRANSPORT_OF_INORGANIC_CATIONS_ANIONS_AND_AMINO_ACIDS_OLIGOPEPTIDES | | | | | | | |
| REACTOME_SLC_MEDIATED_TRANSMEMBRANE_TRANSPORT | REACTOME_TRANSPORT_OF_INORGANIC_CATIONS_ANIONS_AND_AMINO_ACIDS_OLIGOPEPTIDES | | | | | | | |
| REACTOME_SLC_MEDIATED_TRANSMEMBRANE_TRANSPORT | REACTOME_TRANSPORT_OF_INORGANIC_CATIONS_ANIONS_AND_AMINO_ACIDS_OLIGOPEPTIDES | | | | | | | |
| REACTOME_SLC_MEDIATED_TRANSMEMBRANE_TRANSPORT | REACTOME_TRANSPORT_OF_INORGANIC_CATIONS_ANIONS_AND_AMINO_ACIDS_OLIGOPEPTIDES | | | | | | | |
| REACTOME_SLC_MEDIATED_TRANSMEMBRANE_TRANSPORT | REACTOME_TRANSPORT_OF_INORGANIC_CATIONS_ANIONS_AND_AMINO_ACIDS_OLIGOPEPTIDES | | | | | | | |
| REACTOME_SLC_MEDIATED_TRANSMEMBRANE_TRANSPORT | REACTOME_TRANSPORT_OF_INORGANIC_CATIONS_ANIONS_AND_AMINO_ACIDS_OLIGOPEPTIDES | | | | | | | |
| REACTOME_SLC_MEDIATED_TRANSMEMBRANE_TRANSPORT | REACTOME_TRANSPORT_OF_INORGANIC_CATIONS_ANIONS_AND_AMINO_ACIDS_OLIGOPEPTIDES | | | | | | | |
| REACTOME_SLC_MEDIATED_TRANSMEMBRANE_TRANSPORT | REACTOME_TRANSPORT_OF_INORGANIC_CATIONS_ANIONS_AND_AMINO_ACIDS_OLIGOPEPTIDES | | | | | | | |
| REACTOME_SLC_MEDIATED_TRANSMEMBRANE_TRANSPORT | REACTOME_TRANSPORT_OF_INORGANIC_CATIONS_ANIONS_AND_AMINO_ACIDS_OLIGOPEPTIDES | | | | | | | |
| REACTOME_SLC_MEDIATED_TRANSMEMBRANE_TRANSPORT | REACTOME_TRANSPORT_OF_INORGANIC_CATIONS_ANIONS_AND_AMINO_ACIDS_OLIGOPEPTIDES | | | | | | | |
| REACTOME_SLC_MEDIATED_TRANSMEMBRANE_TRANSPORT | | | | | | | | |
| REACTOME_SLC_MEDIATED_TRANSMEMBRANE_TRANSPORT | | | | | | | | |
| REACTOME_SLC_MEDIATED_TRANSMEMBRANE_TRANSPORT | | | | | | | | |
| REACTOME_SLC_MEDIATED_TRANSMEMBRANE_TRANSPORT | | | | | | | | |
| REACTOME_SLC_MEDIATED_TRANSMEMBRANE_TRANSPORT | | | | | | | | |
| REACTOME_SLC_MEDIATED_TRANSMEMBRANE_TRANSPORT | | | | | | | | |
| REACTOME_SLC_MEDIATED_TRANSMEMBRANE_TRANSPORT | | | | | | | | |
| REACTOME_SLC_MEDIATED_TRANSMEMBRANE_TRANSPORT | | | | | | | | |
| REACTOME_SLC_MEDIATED_TRANSMEMBRANE_TRANSPORT | | | | | | | | |
| | | | | HALLMARK_XENOBIOTIC_METABOLISM | | | | |
| | | | | HALLMARK_XENOBIOTIC_METABOLISM | | | | |
| | | | | | KEGG_ARGININE_AND_PROLINE_METABOLISM | | | KEGG_BETA_ALANINE_METABOLISM |
| | | | | | | WP_AMINO_ACID_METABOLISM | | KEGG_BETA_ALANINE_METABOLISM |
| | | | | | | WP_AMINO_ACID_METABOLISM | | |
| | | | | | | | | KEGG_BETA_ALANINE_METABOLISM |
| | | KEGG_GLYCINE_SERINE_AND_THREONINE_METABOLISM | | | | | | |
| | | | | HALLMARK_XENOBIOTIC_METABOLISM | | WP_AMINO_ACID_METABOLISM | | |
| | | | | HALLMARK_XENOBIOTIC_METABOLISM | | WP_AMINO_ACID_METABOLISM | | |
| | | | | HALLMARK_XENOBIOTIC_METABOLISM | | | KEGG_PEROXISOME | |
| | | | | HALLMARK_XENOBIOTIC_METABOLISM | | | | KEGG_BETA_ALANINE_METABOLISM |
| | | | | HALLMARK_XENOBIOTIC_METABOLISM | | | | KEGG_BETA_ALANINE_METABOLISM |
| | | | | HALLMARK_XENOBIOTIC_METABOLISM | | | | |
| | | | | HALLMARK_XENOBIOTIC_METABOLISM | | | | |
| | | | | HALLMARK_XENOBIOTIC_METABOLISM | | | | |

Bulk RNA-seq DOWN

| | | | | |
|---|---|---|---|---|
| HALLMARK_XENOBIOTIC_METABOLISM | | | | |
| HALLMARK_XENOBIOTIC_METABOLISM | | | | |
| HALLMARK_XENOBIOTIC_METABOLISM | | | | |
| HALLMARK_XENOBIOTIC_METABOLISM | | | | |
| HALLMARK_XENOBIOTIC_METABOLISM | | | | |
| HALLMARK_XENOBIOTIC_METABOLISM | | | | |
| HALLMARK_XENOBIOTIC_METABOLISM | | | | |
| HALLMARK_XENOBIOTIC_METABOLISM | | | | |
| | KEGG_ARGININE_AND_PROLINE_METABOLISM | | | |
| | KEGG_ARGININE_AND_PROLINE_METABOLISM | | | |
| | | WP_AMINO_ACID_METABOLISM | | |
| | | WP_AMINO_ACID_METABOLISM | | |
| | | | KEGG_PEROXISOME | |
| | | | KEGG_PEROXISOME | |
| | | | KEGG_PEROXISOME | |
| | | | KEGG_PEROXISOME | |
| | | | KEGG_PEROXISOME | |
| | | | KEGG_PEROXISOME | |
| | | | KEGG_PEROXISOME | |
| | | | KEGG_PEROXISOME | |
| | | | KEGG_PEROXISOME | |
| | | | | KEGG_BETA_ALANINE_METABOLISM |

Bulk RNA-seq DOWN

Bulk RNA-seq DOWN

Bulk RNA-seq DOWN

Bulk RNA-seq DOWN

Bulk RNA-seq DOWN

Bulk RNA-seq DOWN

Bulk RNA-seq DOWN

Bulk RNA-seq DOWN

| | | | | | | | | | | |
|---|---|---|---|---|---|---|---|---|---|---|
| REACTOME_METABOLISM_OF_LIPIDS | KEGG_PROPANOATE_METABOLISM | REACTOME_FATTY_ACID_METABOLISM | HALLMARK_FATTY_ACID_METABOLISM | KEGG_TRYPTOPHAN_METABOLISM | REACTOME_BRANCHED_CHAIN_AMINO_ACID_CATABOLISM | REACTOME_PROTEIN_LOCALIZATION | WP_TRANSSULFURATION_AND_ONE_CARBON_METABOLISM | REACTOME_PEROXISOMAL_PROTEIN_IMPORT | REACTOME_BIOLOGICAL_OXIDATIONS | WP_PROXIMAL_TUBULE_TRANSPORT |
| | | | HALLMARK_FATTY_ACID_METABOLISM | | REACTOME_BRANCHED_CHAIN_AMINO_ACID_CATABOLISM | | | | | |
| | | | | | REACTOME_BRANCHED_CHAIN_AMINO_ACID_CATABOLISM | | | | | |
| | | | | | REACTOME_BRANCHED_CHAIN_AMINO_ACID_CATABOLISM | | | | | |
| | KEGG_PROPANOATE_METABOLISM | | | KEGG_TRYPTOPHAN_METABOLISM | | | | | | |
| | KEGG_PROPANOATE_METABOLISM | | HALLMARK_FATTY_ACID_METABOLISM | | REACTOME_BRANCHED_CHAIN_AMINO_ACID_CATABOLISM | | | | | |
| | KEGG_PROPANOATE_METABOLISM | | | | REACTOME_BRANCHED_CHAIN_AMINO_ACID_CATABOLISM | | | | | |
| | | | | | REACTOME_BRANCHED_CHAIN_AMINO_ACID_CATABOLISM | | | | | |
| | | | | | REACTOME_BRANCHED_CHAIN_AMINO_ACID_CATABOLISM | | | | | |
| | | | | | | REACTOME_PROTEIN_LOCALIZATION | | REACTOME_PEROXISOMAL_PROTEIN_IMPORT | | |
| | | | | | | | WP_TRANSSULFURATION_AND_ONE_CARBON_METABOLISM | | | |
| | | | | | | | WP_TRANSSULFURATION_AND_ONE_CARBON_METABOLISM | | | |
| | | | | | | REACTOME_PROTEIN_LOCALIZATION | | REACTOME_PEROXISOMAL_PROTEIN_IMPORT | | |
| | | | | | | | WP_TRANSSULFURATION_AND_ONE_CARBON_METABOLISM | | | |
| | | | | | | | WP_TRANSSULFURATION_AND_ONE_CARBON_METABOLISM | | | |
| | | | | KEGG_TRYPTOPHAN_METABOLISM | | | | | | |
| | | | | | | REACTOME_PROTEIN_LOCALIZATION | | | | |
| | | | | KEGG_TRYPTOPHAN_METABOLISM | | | | | | |
| | | | | | | | WP_TRANSSULFURATION_AND_ONE_CARBON_METABOLISM | | REACTOME_BIOLOGICAL_OXIDATIONS | |
| | | | | KEGG_TRYPTOPHAN_METABOLISM | | | | | | |
| | | | | KEGG_TRYPTOPHAN_METABOLISM | | | | | | |
| | | | | | | | WP_TRANSSULFURATION_AND_ONE_CARBON_METABOLISM | | | |
| | | | | | | | | | | WP_PROXIMAL_TUBULE_TRANSPORT |
| | | | | | | | | | | WP_PROXIMAL_TUBULE_TRANSPORT |
| | | | | | | | | | | WP_PROXIMAL_TUBULE_TRANSPORT |
| | | | | | | | | | | WP_PROXIMAL_TUBULE_TRANSPORT |
| | | | | | | | | | | WP_PROXIMAL_TUBULE_TRANSPORT |
| | | | | | | | | | | WP_PROXIMAL_TUBULE_TRANSPORT |
| | | | | | | | | | | WP_PROXIMAL_TUBULE_TRANSPORT |
| | | | HALLMARK_FATTY_ACID_METABOLISM | | | | | | | WP_PROXIMAL_TUBULE_TRANSPORT |
| REACTOME_METABOLISM_OF_LIPIDS | | | | | | | | | | |
| | | | | | | | | | | WP_PROXIMAL_TUBULE_TRANSPORT |
| REACTOME_METABOLISM_OF_LIPIDS | KEGG_PROPANOATE_METABOLISM | REACTOME_FATTY_ACID_METABOLISM | HALLMARK_FATTY_ACID_METABOLISM | KEGG_TRYPTOPHAN_METABOLISM | | REACTOME_PROTEIN_LOCALIZATION | | | | |
| REACTOME_METABOLISM_OF_LIPIDS | KEGG_PROPANOATE_METABOLISM | REACTOME_FATTY_ACID_METABOLISM | HALLMARK_FATTY_ACID_METABOLISM | | | | | | | |
| REACTOME_METABOLISM_OF_LIPIDS | | REACTOME_FATTY_ACID_METABOLISM | HALLMARK_FATTY_ACID_METABOLISM | KEGG_TRYPTOPHAN_METABOLISM | | | | | | |
| | KEGG_PROPANOATE_METABOLISM | | | | | | | | | |
| REACTOME_METABOLISM_OF_LIPIDS | KEGG_PROPANOATE_METABOLISM | REACTOME_FATTY_ACID_METABOLISM | HALLMARK_FATTY_ACID_METABOLISM | | | | | | | |
| REACTOME_METABOLISM_OF_LIPIDS | | | | | | | | | | |
| | | | HALLMARK_FATTY_ACID_METABOLISM | | | REACTOME_PROTEIN_LOCALIZATION | | | | |
| | | | | | | | | | REACTOME_BIOLOGICAL_OXIDATIONS | |
| REACTOME_METABOLISM_OF_LIPIDS | | REACTOME_FATTY_ACID_METABOLISM | | | | REACTOME_PROTEIN_LOCALIZATION | | REACTOME_PEROXISOMAL_PROTEIN_IMPORT | | |
| REACTOME_METABOLISM_OF_LIPIDS | | REACTOME_FATTY_ACID_METABOLISM | | | | | | | REACTOME_BIOLOGICAL_OXIDATIONS | |
| REACTOME_METABOLISM_OF_LIPIDS | | | | | | | | | | |
| | | | HALLMARK_FATTY_ACID_METABOLISM | | | | | | REACTOME_BIOLOGICAL_OXIDATIONS | |

Bulk RNA-seq DOWN

| | | | | | | | | |
|---|---|---|---|---|---|---|---|---|
| | | | HALLMARK_FATTY_ACID_METABOLISM | | | | | REACTOME_BIOLOGICAL_OXIDATIONS |
| | | | | KEGG_TRYPTOPHAN_METABOLISM | | | | REACTOME_BIOLOGICAL_OXIDATIONS |
| | | | | | | | | REACTOME_BIOLOGICAL_OXIDATIONS |
| | KEGG_PROPANOATE_METABOLISM | | HALLMARK_FATTY_ACID_METABOLISM | | | | | |
| REACTOME_METABOLISM_OF_LIPIDS | | REACTOME_FATTY_ACID_METABOLISM | HALLMARK_FATTY_ACID_METABOLISM | | REACTOME_PROTEIN_LOCALIZATION | | REACTOME_PEROXISOMAL_PROTEIN_IMPORT | |
| REACTOME_METABOLISM_OF_LIPIDS | | REACTOME_FATTY_ACID_METABOLISM | HALLMARK_FATTY_ACID_METABOLISM | | REACTOME_PROTEIN_LOCALIZATION | | REACTOME_PEROXISOMAL_PROTEIN_IMPORT | |
| REACTOME_METABOLISM_OF_LIPIDS | | REACTOME_FATTY_ACID_METABOLISM | | | REACTOME_PROTEIN_LOCALIZATION | | REACTOME_PEROXISOMAL_PROTEIN_IMPORT | |
| REACTOME_METABOLISM_OF_LIPIDS | | REACTOME_FATTY_ACID_METABOLISM | | | REACTOME_PROTEIN_LOCALIZATION | | REACTOME_PEROXISOMAL_PROTEIN_IMPORT | |
| | | | | | REACTOME_PROTEIN_LOCALIZATION | | REACTOME_PEROXISOMAL_PROTEIN_IMPORT | |
| | | | | | REACTOME_PROTEIN_LOCALIZATION | | REACTOME_PEROXISOMAL_PROTEIN_IMPORT | |
| | | | | | REACTOME_PROTEIN_LOCALIZATION | | REACTOME_PEROXISOMAL_PROTEIN_IMPORT | |
| | | | | | REACTOME_PROTEIN_LOCALIZATION | | | |
| REACTOME_METABOLISM_OF_LIPIDS | | REACTOME_FATTY_ACID_METABOLISM | HALLMARK_FATTY_ACID_METABOLISM | | | | | |
| REACTOME_METABOLISM_OF_LIPIDS | | REACTOME_FATTY_ACID_METABOLISM | | | | | | |
| REACTOME_METABOLISM_OF_LIPIDS | | REACTOME_FATTY_ACID_METABOLISM | | | | | | |
| REACTOME_METABOLISM_OF_LIPIDS | | REACTOME_FATTY_ACID_METABOLISM | | | | | | |
| REACTOME_METABOLISM_OF_LIPIDS | | REACTOME_FATTY_ACID_METABOLISM | | | | | | |
| REACTOME_METABOLISM_OF_LIPIDS | | REACTOME_FATTY_ACID_METABOLISM | | | | | | |
| REACTOME_METABOLISM_OF_LIPIDS | | REACTOME_FATTY_ACID_METABOLISM | | | | | | |
| REACTOME_METABOLISM_OF_LIPIDS | | REACTOME_FATTY_ACID_METABOLISM | | | | | | |
| REACTOME_METABOLISM_OF_LIPIDS | | | HALLMARK_FATTY_ACID_METABOLISM | | | | | |
| REACTOME_METABOLISM_OF_LIPIDS | | | | | | | | REACTOME_BIOLOGICAL_OXIDATIONS |
| REACTOME_METABOLISM_OF_LIPIDS | | | | | | | | |
| REACTOME_METABOLISM_OF_LIPIDS | | | | | | | | |
| REACTOME_METABOLISM_OF_LIPIDS | | | | | | | | |
| REACTOME_METABOLISM_OF_LIPIDS | | | | | | | | |
| REACTOME_METABOLISM_OF_LIPIDS | | | | | | | | |
| REACTOME_METABOLISM_OF_LIPIDS | | | | | | | | |
| REACTOME_METABOLISM_OF_LIPIDS | | | | | | | | |
| REACTOME_METABOLISM_OF_LIPIDS | | | | | | | | |
| REACTOME_METABOLISM_OF_LIPIDS | | | | | | | | |
| REACTOME_METABOLISM_OF_LIPIDS | | | | | | | | |
| REACTOME_METABOLISM_OF_LIPIDS | | | | | | | | |
| REACTOME_METABOLISM_OF_LIPIDS | | | | | | | | |
| REACTOME_METABOLISM_OF_LIPIDS | | | | | | | | |
| REACTOME_METABOLISM_OF_LIPIDS | | | | | | | | |
| REACTOME_METABOLISM_OF_LIPIDS | | | | | | | | |
| REACTOME_METABOLISM_OF_LIPIDS | | | | | | | | |
| REACTOME_METABOLISM_OF_LIPIDS | | | | | | | | |
| | KEGG_PROPANOATE_METABOLISM | | HALLMARK_FATTY_ACID_METABOLISM | | | | | |
| | | | HALLMARK_FATTY_ACID_METABOLISM | | | | | |
| | | | | KEGG_TRYPTOPHAN_METABOLISM | | | | |
| | | | | | REACTOME_PROTEIN_LOCALIZATION | | | |
| | | | | | REACTOME_PROTEIN_LOCALIZATION | | | |
| | | | | | | WP_TRANSSULFURATION_AND_ONE_CARBON_METABOLISM | | |
| | | | | | | WP_TRANSSULFURATION_AND_ONE_CARBON_METABOLISM | | |
| | | | | | | | | REACTOME_BIOLOGICAL_OXIDATIONS |
| | | | | | | | | REACTOME_BIOLOGICAL_OXIDATIONS |
| | | | | | | | | REACTOME_BIOLOGICAL_OXIDATIONS |
| | | | | | | | | REACTOME_BIOLOGICAL_OXIDATIONS |
| | | | | | | | | REACTOME_BIOLOGICAL_OXIDATIONS |
| | | | | | | | | REACTOME_BIOLOGICAL_OXIDATIONS |
| | | | | | | | | REACTOME_BIOLOGICAL_OXIDATIONS |
| | | | | | | | | REACTOME_BIOLOGICAL_OXIDATIONS |
| | | | | | | | | REACTOME_BIOLOGICAL_OXIDATIONS |
| | | | | | | | | REACTOME_BIOLOGICAL_OXIDATIONS |

Bulk RNA-seq DOWN

Bulk RNA-seq DOWN

Bulk RNA-seq DOWN

Bulk RNA-seq DOWN

Bulk RNA-seq DOWN

Bulk RNA-seq DOWN

Bulk RNA-seq DOWN

Bulk RNA-seq DOWN

| WP_ONE_CARBON_METABOLISM | REACTOME_CHOLINE_CATABOLISM | REACTOME_SLC_TRANSPORTER_DISORDERS | REACTOME_METABOLISM_OF_WATER_SOLUBLE_VITAMINS_AND_COFACTORS | REACTOME_DISORDERS_OF_TRANSMEMBRANE_TRANSPORTERS | KEGG_DRUG_METABOLISM_CYTOCHROME_P450 | WP_TRYPTOPHAN_METABOLISM | WP_METAPATHWAY_BIOTRANSFORMATION_PHASE_I_AND_II | KEGG_HISTIDINE_METABOLISM | REACTOME_NEURONAL_SYSTEM |
|---|---|---|---|---|---|---|---|---|---|
| | | | REACTOME_METABOLISM_OF_WATER_SOLUBLE_VITAMINS_AND_COFACTORS | | | | | | |
| | REACTOME_CHOLINE_CATABOLISM | | | | | | | KEGG_HISTIDINE_METABOLISM | |
| WP_ONE_CARBON_METABOLISM | | | | | | | | | |
| WP_ONE_CARBON_METABOLISM | REACTOME_CHOLINE_CATABOLISM | | | | | | | | |
| WP_ONE_CARBON_METABOLISM | | | REACTOME_METABOLISM_OF_WATER_SOLUBLE_VITAMINS_AND_COFACTORS | | | | | | |
| | REACTOME_CHOLINE_CATABOLISM | | | | | | | | |
| | REACTOME_CHOLINE_CATABOLISM | | | | | | | | |
| | | | | | | WP_TRYPTOPHAN_METABOLISM | | | |
| | | | | | | WP_TRYPTOPHAN_METABOLISM | | KEGG_HISTIDINE_METABOLISM | |
| WP_ONE_CARBON_METABOLISM | | | | | | | | | |
| WP_ONE_CARBON_METABOLISM | | | | | | | | KEGG_HISTIDINE_METABOLISM | |
| | | | | | | WP_TRYPTOPHAN_METABOLISM | | | |
| WP_ONE_CARBON_METABOLISM | | | | | | | | | |
| WP_ONE_CARBON_METABOLISM | | | | | | | | | |
| | | REACTOME_SLC_TRANSPORTER_DISORDERS | | REACTOME_DISORDERS_OF_TRANSMEMBRANE_TRANSPORTERS | | | | | |
| | | REACTOME_SLC_TRANSPORTER_DISORDERS | | REACTOME_DISORDERS_OF_TRANSMEMBRANE_TRANSPORTERS | | | | | |
| | | REACTOME_SLC_TRANSPORTER_DISORDERS | | REACTOME_DISORDERS_OF_TRANSMEMBRANE_TRANSPORTERS | | | | | |
| | | REACTOME_SLC_TRANSPORTER_DISORDERS | | REACTOME_DISORDERS_OF_TRANSMEMBRANE_TRANSPORTERS | | | | | |
| | | REACTOME_SLC_TRANSPORTER_DISORDERS | | REACTOME_DISORDERS_OF_TRANSMEMBRANE_TRANSPORTERS | | | | | |
| | | REACTOME_SLC_TRANSPORTER_DISORDERS | | REACTOME_DISORDERS_OF_TRANSMEMBRANE_TRANSPORTERS | | | | | |
| | | REACTOME_SLC_TRANSPORTER_DISORDERS | | REACTOME_DISORDERS_OF_TRANSMEMBRANE_TRANSPORTERS | | | | | |
| | | REACTOME_SLC_TRANSPORTER_DISORDERS | | REACTOME_DISORDERS_OF_TRANSMEMBRANE_TRANSPORTERS | | | | | |
| | | REACTOME_SLC_TRANSPORTER_DISORDERS | | REACTOME_DISORDERS_OF_TRANSMEMBRANE_TRANSPORTERS | | | | | |
| | | REACTOME_SLC_TRANSPORTER_DISORDERS | | REACTOME_DISORDERS_OF_TRANSMEMBRANE_TRANSPORTERS | | | | | |
| | | REACTOME_SLC_TRANSPORTER_DISORDERS | | REACTOME_DISORDERS_OF_TRANSMEMBRANE_TRANSPORTERS | | | | | |
| | | | | | | | | | REACTOME_NEURONAL_SYSTEM |
| | | | REACTOME_METABOLISM_OF_WATER_SOLUBLE_VITAMINS_AND_COFACTORS | | | | | | |
| | | | REACTOME_METABOLISM_OF_WATER_SOLUBLE_VITAMINS_AND_COFACTORS | | | | | | |
| | | | | REACTOME_DISORDERS_OF_TRANSMEMBRANE_TRANSPORTERS | | | | | |
| | | | | REACTOME_DISORDERS_OF_TRANSMEMBRANE_TRANSPORTERS | | | | | |
| | | | | REACTOME_DISORDERS_OF_TRANSMEMBRANE_TRANSPORTERS | | | | | |
| | | | | | | | | | REACTOME_NEURONAL_SYSTEM |
| | | | | | | | | | REACTOME_NEURONAL_SYSTEM |
| | | | | | | | | | REACTOME_NEURONAL_SYSTEM |
| | | | | | | | | | REACTOME_NEURONAL_SYSTEM |
| | | | | | | WP_TRYPTOPHAN_METABOLISM | | KEGG_HISTIDINE_METABOLISM | |
| | | | | | | WP_TRYPTOPHAN_METABOLISM | | | |
| | | | | | | | | | REACTOME_NEURONAL_SYSTEM |
| | | | | | KEGG_DRUG_METABOLISM_CYTOCHROME_P450 | | | | |
| | | | | | | | | | REACTOME_NEURONAL_SYSTEM |
| | | | | | | | WP_METAPATHWAY_BIOTRANSFORMATION_PHASE_I_AND_II | | |
| | | | | | KEGG_DRUG_METABOLISM_CYTOCHROME_P450 | | WP_METAPATHWAY_BIOTRANSFORMATION_PHASE_I_AND_II | | |

Bulk RNA-seq DOWN

| | | | | | |
|---|---|---|---|---|---|
| | | WP_TRYPTOPHAN_METABOLISM | | KEGG_HISTIDINE_METABOLISM | |
| | | | WP_METAPATHWAY_BIOTRANSFORMATION_PHASE_I_AND_II | | |
| REACTOME_METABOLISM_OF_WATER_SOLUBLE_VITAMINS_AND_COFACTORS | | | | | |
| | | | | KEGG_HISTIDINE_METABOLISM | |
| | KEGG_DRUG_METABOLISM_CYTOCHROME_P450 | WP_TRYPTOPHAN_METABOLISM | WP_METAPATHWAY_BIOTRANSFORMATION_PHASE_I_AND_II | | |
| REACTOME_METABOLISM_OF_WATER_SOLUBLE_VITAMINS_AND_COFACTORS | | | | | |
| | KEGG_DRUG_METABOLISM_CYTOCHROME_P450 | | WP_METAPATHWAY_BIOTRANSFORMATION_PHASE_I_AND_II | | |
| | KEGG_DRUG_METABOLISM_CYTOCHROME_P450 | | WP_METAPATHWAY_BIOTRANSFORMATION_PHASE_I_AND_II | | |
| | KEGG_DRUG_METABOLISM_CYTOCHROME_P450 | | WP_METAPATHWAY_BIOTRANSFORMATION_PHASE_I_AND_II | | |
| | KEGG_DRUG_METABOLISM_CYTOCHROME_P450 | | | | |
| | | | WP_METAPATHWAY_BIOTRANSFORMATION_PHASE_I_AND_II | | |
| | | | WP_METAPATHWAY_BIOTRANSFORMATION_PHASE_I_AND_II | | |
| REACTOME_METABOLISM_OF_WATER_SOLUBLE_VITAMINS_AND_COFACTORS | | | | | |
| REACTOME_METABOLISM_OF_WATER_SOLUBLE_VITAMINS_AND_COFACTORS | | | | | |
| REACTOME_METABOLISM_OF_WATER_SOLUBLE_VITAMINS_AND_COFACTORS | | | | | |
| REACTOME_METABOLISM_OF_WATER_SOLUBLE_VITAMINS_AND_COFACTORS | | | | | |
| REACTOME_METABOLISM_OF_WATER_SOLUBLE_VITAMINS_AND_COFACTORS | | | | | |
| REACTOME_METABOLISM_OF_WATER_SOLUBLE_VITAMINS_AND_COFACTORS | | | | | |
| | KEGG_DRUG_METABOLISM_CYTOCHROME_P450 | | WP_METAPATHWAY_BIOTRANSFORMATION_PHASE_I_AND_II | | |
| | KEGG_DRUG_METABOLISM_CYTOCHROME_P450 | | WP_METAPATHWAY_BIOTRANSFORMATION_PHASE_I_AND_II | | |
| | | | WP_METAPATHWAY_BIOTRANSFORMATION_PHASE_I_AND_II | | |
| | | | WP_METAPATHWAY_BIOTRANSFORMATION_PHASE_I_AND_II | | |
| | | | WP_METAPATHWAY_BIOTRANSFORMATION_PHASE_I_AND_II | | |
| | | | | | REACTOME_NEURONAL_SYSTEM |
| | | | | | REACTOME_NEURONAL_SYSTEM |
| | | | | | REACTOME_NEURONAL_SYSTEM |
| | | | | | REACTOME_NEURONAL_SYSTEM |
| | | | | | REACTOME_NEURONAL_SYSTEM |
| | | | | | REACTOME_NEURONAL_SYSTEM |
| | | | | | REACTOME_NEURONAL_SYSTEM |
| | | | | | REACTOME_NEURONAL_SYSTEM |
| | | | | | REACTOME_NEURONAL_SYSTEM |
| | | | | | REACTOME_NEURONAL_SYSTEM |
| | | | | | REACTOME_NEURONAL_SYSTEM |
| | | | | | REACTOME_NEURONAL_SYSTEM |
| | | | | | REACTOME_NEURONAL_SYSTEM |
| | | | | | REACTOME_NEURONAL_SYSTEM |
| | | | | | REACTOME_NEURONAL_SYSTEM |

Bulk RNA-seq DOWN

Bulk RNA-seq DOWN

Bulk RNA-seq DOWN

Bulk RNA-seq DOWN

Bulk RNA-seq DOWN

Bulk RNA-seq DOWN

Bulk RNA-seq DOWN

Bulk RNA-seq DOWN

| | | | | | | | | | |
|---|---|---|---|---|---|---|---|---|---|
| REACTOME_TRANSMISSION_ACROSS_CHEMICAL_SYNAPSES | KEGG_GLYCOLYSIS_GLUCONEOGENESIS | KEGG_CITRATE_CYCLE_TCA_CYCLE | KEGG_ALANINE_ASPARTATE_AND_GLUTAMATE_METABOLISM | HALLMARK_ADIPOGENESIS | REACTOME_CARDIAC_CONDUCTION | KEGG_GLYCEROLIPID_METABOLISM | WP_UREA_CYCLE_AND_METABOLISM_OF_AMINO_GROUPS | KEGG_BUTANOATE_METABOLISM | REACTOME_ION_CHANNEL_TRANSPORT |
| | | | | HALLMARK_ADIPOGENESIS | | | | | |
| | KEGG_GLYCOLYSIS_GLUCONEOGENESIS | | | | | KEGG_GLYCEROLIPID_METABOLISM | | KEGG_BUTANOATE_METABOLISM | |
| | | | | HALLMARK_ADIPOGENESIS | | | | | |
| | | | | | | | WP_UREA_CYCLE_AND_METABOLISM_OF_AMINO_GROUPS | | |
| | | | | | | | WP_UREA_CYCLE_AND_METABOLISM_OF_AMINO_GROUPS | | |
| | | | KEGG_ALANINE_ASPARTATE_AND_GLUTAMATE_METABOLISM | | | | | | |
| | | KEGG_CITRATE_CYCLE_TCA_CYCLE | | | | | | | |
| | | | KEGG_ALANINE_ASPARTATE_AND_GLUTAMATE_METABOLISM | | | | WP_UREA_CYCLE_AND_METABOLISM_OF_AMINO_GROUPS | | |
| | | | KEGG_ALANINE_ASPARTATE_AND_GLUTAMATE_METABOLISM | | | | | | |
| | | | | | | | WP_UREA_CYCLE_AND_METABOLISM_OF_AMINO_GROUPS | | |
| | | | KEGG_ALANINE_ASPARTATE_AND_GLUTAMATE_METABOLISM | | | | | | |
| | | | | HALLMARK_ADIPOGENESIS | | | | | |
| | | | | | REACTOME_CARDIAC_CONDUCTION | | | | |
| REACTOME_TRANSMISSION_ACROSS_CHEMICAL_SYNAPSES | | | | | | | | | |
| | | | | | REACTOME_CARDIAC_CONDUCTION | | | | REACTOME_ION_CHANNEL_TRANSPORT |
| REACTOME_TRANSMISSION_ACROSS_CHEMICAL_SYNAPSES | | | | | REACTOME_CARDIAC_CONDUCTION | | | | REACTOME_ION_CHANNEL_TRANSPORT |
| REACTOME_TRANSMISSION_ACROSS_CHEMICAL_SYNAPSES | | | | | REACTOME_CARDIAC_CONDUCTION | | | | REACTOME_ION_CHANNEL_TRANSPORT |
| REACTOME_TRANSMISSION_ACROSS_CHEMICAL_SYNAPSES | | | | | REACTOME_CARDIAC_CONDUCTION | | | | REACTOME_ION_CHANNEL_TRANSPORT |
| REACTOME_TRANSMISSION_ACROSS_CHEMICAL_SYNAPSES | | | | | | | | | |
| | | | | HALLMARK_ADIPOGENESIS | | | | | |
| | | | | | REACTOME_CARDIAC_CONDUCTION | | | | REACTOME_ION_CHANNEL_TRANSPORT |
| | | | | | | | | | REACTOME_ION_CHANNEL_TRANSPORT |
| | | | | | | | | | REACTOME_ION_CHANNEL_TRANSPORT |
| | | | | | | | | | REACTOME_ION_CHANNEL_TRANSPORT |
| | | | | | | | | | REACTOME_ION_CHANNEL_TRANSPORT |
| | | | | | | | | | REACTOME_ION_CHANNEL_TRANSPORT |
| | | | | | | | | | REACTOME_ION_CHANNEL_TRANSPORT |
| | | | | | | | | | REACTOME_ION_CHANNEL_TRANSPORT |
| | | | | | | | | | REACTOME_ION_CHANNEL_TRANSPORT |
| | KEGG_GLYCOLYSIS_GLUCONEOGENESIS | | | | | KEGG_GLYCEROLIPID_METABOLISM | | KEGG_BUTANOATE_METABOLISM | |
| | | | | HALLMARK_ADIPOGENESIS | | | | | |
| | | | | HALLMARK_ADIPOGENESIS | | | | KEGG_BUTANOATE_METABOLISM | |
| REACTOME_TRANSMISSION_ACROSS_CHEMICAL_SYNAPSES | | | KEGG_ALANINE_ASPARTATE_AND_GLUTAMATE_METABOLISM | | | | | KEGG_BUTANOATE_METABOLISM | |
| | | | | | | | | KEGG_BUTANOATE_METABOLISM | |
| | | | | | | KEGG_GLYCEROLIPID_METABOLISM | | | |
| | | KEGG_CITRATE_CYCLE_TCA_CYCLE | | HALLMARK_ADIPOGENESIS | | | | | |
| | KEGG_GLYCOLYSIS_GLUCONEOGENESIS | | | | | | | | |
| REACTOME_TRANSMISSION_ACROSS_CHEMICAL_SYNAPSES | | | KEGG_ALANINE_ASPARTATE_AND_GLUTAMATE_METABOLISM | | | | | KEGG_BUTANOATE_METABOLISM | |

Bulk RNA-seq DOWN

KEGG_GLYCOLYSIS_GLUCONEOGENESIS

KEGG_GLYCEROLIPID_METABOLISM

WP_UREA_CYCLE_AND_METABOLISM_OF_AMINO_GROUPS

KEGG_CITRATE_CYCLE_TCA_CYCLE
HALLMARK_ADIPOGENESIS

HALLMARK_ADIPOGENESIS

HALLMARK_ADIPOGENESIS
HALLMARK_ADIPOGENESIS

HALLMARK_ADIPOGENESIS
KEGG_GLYCEROLIPID_METABOLISM
KEGG_GLYCEROLIPID_METABOLISM

KEGG_CITRATE_CYCLE_TCA_CYCLE

KEGG_CITRATE_CYCLE_TCA_CYCLE

KEGG_GLYCOLYSIS_GLUCONEOGENESIS

HALLMARK_ADIPOGENESIS

HALLMARK_ADIPOGENESIS

REACTOME_TRANSMISSION_ACROSS_CHEMICAL_SYNAPSES
REACTOME_CARDIAC_CONDUCTION
REACTOME_TRANSMISSION_ACROSS_CHEMICAL_SYNAPSES
REACTOME_TRANSMISSION_ACROSS_CHEMICAL_SYNAPSES
REACTOME_TRANSMISSION_ACROSS_CHEMICAL_SYNAPSES
REACTOME_TRANSMISSION_ACROSS_CHEMICAL_SYNAPSES
REACTOME_TRANSMISSION_ACROSS_CHEMICAL_SYNAPSES
REACTOME_TRANSMISSION_ACROSS_CHEMICAL_SYNAPSES
REACTOME_TRANSMISSION_ACROSS_CHEMICAL_SYNAPSES
REACTOME_TRANSMISSION_ACROSS_CHEMICAL_SYNAPSES
REACTOME_TRANSMISSION_ACROSS_CHEMICAL_SYNAPSES

REACTOME_CARDIAC_CONDUCTION

KEGG_GLYCOLYSIS_GLUCONEOGENESIS
KEGG_CITRATE_CYCLE_TCA_CYCLE
KEGG_GLYCOLYSIS_GLUCONEOGENESIS
KEGG_GLYCOLYSIS_GLUCONEOGENESIS

REACTOME_CARDIAC_CONDUCTION
REACTOME_CARDIAC_CONDUCTION
REACTOME_CARDIAC_CONDUCTION
KEGG_GLYCEROLIPID_METABOLISM

Bulk RNA-seq DOWN

Bulk RNA-seq DOWN

Bulk RNA-seq DOWN

Bulk RNA-seq DOWN

Bulk RNA-seq DOWN

Bulk RNA-seq DOWN

Bulk RNA-seq DOWN

Bulk RNA-seq DOWN

| WP_ALANINE_AND_ASPARTATE_METABOLISM | WP_BIOMARKERS_FOR_UREA_CYCLE_DISORDERS | HALLMARK_BILE_ACID_METABOLISM | KEGG_PROXIMAL_TUBULE_BICARBONATE_RECLAMATION | REACTOME_INTRINSIC_PATHWAY_OF_FIBRIN_CLOT_FORMATION | REACTOME_MITOCHONDRIAL_FATTY_ACID_BETA_OXIDATION | REACTOME_FRUCTOSE_CATABOLISM | WP_CATALYTIC_CYCLE_OF_MAMMALIAN_FLAVINCONTAINING_MONOOXYGENASES_FMOS |
|---|---|---|---|---|---|---|---|
| | WP_BIOMARKERS_FOR_UREA_CYCLE_DISORDERS | | | | | | |
| | | HALLMARK_BILE_ACID_METABOLISM | | | | | |
| | | HALLMARK_BILE_ACID_METABOLISM | | | | | |
| WP_ALANINE_AND_ASPARTATE_METABOLISM | WP_BIOMARKERS_FOR_UREA_CYCLE_DISORDERS | | | | | | |
| | | | KEGG_PROXIMAL_TUBULE_BICARBONATE_RECLAMATION | | | | |
| WP_ALANINE_AND_ASPARTATE_METABOLISM | WP_BIOMARKERS_FOR_UREA_CYCLE_DISORDERS | | | | | | |
| | | | KEGG_PROXIMAL_TUBULE_BICARBONATE_RECLAMATION | | | | |
| | | | KEGG_PROXIMAL_TUBULE_BICARBONATE_RECLAMATION | | | | |
| | | | KEGG_PROXIMAL_TUBULE_BICARBONATE_RECLAMATION | | | | |
| | | HALLMARK_BILE_ACID_METABOLISM | | | | | |
| | | | | | REACTOME_MITOCHONDRIAL_FATTY_ACID_BETA_OXIDATION | | |
| | | | | | REACTOME_MITOCHONDRIAL_FATTY_ACID_BETA_OXIDATION | | |
| WP_ALANINE_AND_ASPARTATE_METABOLISM | | | | | | | |
| | | | | | REACTOME_MITOCHONDRIAL_FATTY_ACID_BETA_OXIDATION | | |
| | | | | | | REACTOME_FRUCTOSE_CATABOLISM | |
| WP_ALANINE_AND_ASPARTATE_METABOLISM | | | | | | | |
| | | HALLMARK_BILE_ACID_METABOLISM | | | | | |
| | | | | | | | WP_CATALYTIC_CYCLE_OF_MAMMALIAN_FLAVINCONTAINING_MONOOXYGENASES_FMOS |

Bulk RNA-seq DOWN

REACTOME_FRUCTOSE_CATABOLISM
REACTOME_INTRINSIC_PATHWAY_OF_FIBRIN_CLOT_FORMATION

HALLMARK_BILE_ACID_METABOLISM
HALLMARK_BILE_ACID_METABOLISM
HALLMARK_BILE_ACID_METABOLISM
HALLMARK_BILE_ACID_METABOLISM

HALLMARK_BILE_ACID_METABOLISM

REACTOME_MITOCHONDRIAL_FATTY_ACID_BETA_OXIDATION
REACTOME_MITOCHONDRIAL_FATTY_ACID_BETA_OXIDATION
REACTOME_MITOCHONDRIAL_FATTY_ACID_BETA_OXIDATION

HALLMARK_BILE_ACID_METABOLISM

WP_CATALYTIC_CYCLE_OF_MAMMALIAN_FLAVINCONTAINING_MONOOXYGENASES_FMOS
WP_CATALYTIC_CYCLE_OF_MAMMALIAN_FLAVINCONTAINING_MONOOXYGENASES_FMOS

KEGG_PROXIMAL_TUBULE_BICARBONATE_RECLAMATION

WP_BIOMARKERS_FOR_UREA_CYCLE_DISORDERS

REACTOME_INTRINSIC_PATHWAY_OF_FIBRIN_CLOT_FORMATION
REACTOME_INTRINSIC_PATHWAY_OF_FIBRIN_CLOT_FORMATION
REACTOME_INTRINSIC_PATHWAY_OF_FIBRIN_CLOT_FORMATION
REACTOME_INTRINSIC_PATHWAY_OF_FIBRIN_CLOT_FORMATION

REACTOME_FRUCTOSE_CATABOLISM

Bulk RNA-seq DOWN

Bulk RNA-seq DOWN

Bulk RNA-seq DOWN

Bulk RNA-seq DOWN

Bulk RNA-seq DOWN

Bulk RNA-seq DOWN

Bulk RNA-seq DOWN

Bulk RNA-seq DOWN

| REACTOME_ION_HOMEOSTASIS | WP_ONE_CARBON_METABOLISM_AND_RELATED_PATHWAYS | REACTOME_METABOLISM_OF_VITAMINS_AND_COFACTORS | REACTOME_FORMATION_OF_FIBRIN_CLOT_CLOTTING_CASCADE | KEGG_PYRUVATE_METABOLISM | KEGG_FATTY_ACID_METABOLISM | WP_NUCLEAR_RECEPTORS_METAPATHWAY | KEGG_CALCIUM_SIGNALING_PATHWAY | KEGG_LYSINE_DEGRADATION | WP_BIOGENIC_AMINE_SYNTHESIS |
|---|---|---|---|---|---|---|---|---|---|
| | | REACTOME_METABOLISM_OF_VITAMINS_AND_COFACTORS | | | | | | | |
| | | | | KEGG_PYRUVATE_METABOLISM | KEGG_FATTY_ACID_METABOLISM | | | KEGG_LYSINE_DEGRADATION | |
| | WP_ONE_CARBON_METABOLISM_AND_RELATED_PATHWAYS | | | | | | | | |
| | | | | | | | | KEGG_LYSINE_DEGRADATION | |
| | WP_ONE_CARBON_METABOLISM_AND_RELATED_PATHWAYS | REACTOME_METABOLISM_OF_VITAMINS_AND_COFACTORS | | | | | | | |
| | WP_ONE_CARBON_METABOLISM_AND_RELATED_PATHWAYS | | | | | | | | |
| | WP_ONE_CARBON_METABOLISM_AND_RELATED_PATHWAYS | | | | | | | | |
| | | | | | | | | KEGG_LYSINE_DEGRADATION | |
| | | | | | | | | | WP_BIOGENIC_AMINE_SYNTHESIS |
| | | | | | | | | | WP_BIOGENIC_AMINE_SYNTHESIS |
| | | | | | | | | | WP_BIOGENIC_AMINE_SYNTHESIS |
| | | | | | | WP_NUCLEAR_RECEPTORS_METAPATHWAY | | | |
| | | | | | | WP_NUCLEAR_RECEPTORS_METAPATHWAY | | | |
| REACTOME_ION_HOMEOSTASIS | | | | | | | KEGG_CALCIUM_SIGNALING_PATHWAY | | |
| | | | | | | WP_NUCLEAR_RECEPTORS_METAPATHWAY | | | |
| | | | | | | WP_NUCLEAR_RECEPTORS_METAPATHWAY | | | |
| | | | | | | WP_NUCLEAR_RECEPTORS_METAPATHWAY | | | |
| | | | | | | WP_NUCLEAR_RECEPTORS_METAPATHWAY | | | |
| | | | | | | WP_NUCLEAR_RECEPTORS_METAPATHWAY | | | |
| | | | | | | WP_NUCLEAR_RECEPTORS_METAPATHWAY | | | |
| | | REACTOME_METABOLISM_OF_VITAMINS_AND_COFACTORS | | | | | | | |
| REACTOME_ION_HOMEOSTASIS | | | | | | | | | |
| | | REACTOME_METABOLISM_OF_VITAMINS_AND_COFACTORS | | | | | | | |
| | | | | | | WP_NUCLEAR_RECEPTORS_METAPATHWAY | | | |
| REACTOME_ION_HOMEOSTASIS | | | | | | | KEGG_CALCIUM_SIGNALING_PATHWAY | | |
| REACTOME_ION_HOMEOSTASIS | | | | | | | KEGG_CALCIUM_SIGNALING_PATHWAY | | |
| REACTOME_ION_HOMEOSTASIS | | | | | | | KEGG_CALCIUM_SIGNALING_PATHWAY | | |
| REACTOME_ION_HOMEOSTASIS | | | | | | | KEGG_CALCIUM_SIGNALING_PATHWAY | | |
| | | REACTOME_METABOLISM_OF_VITAMINS_AND_COFACTORS | | | | WP_NUCLEAR_RECEPTORS_METAPATHWAY | | | |
| | | | | KEGG_PYRUVATE_METABOLISM | KEGG_FATTY_ACID_METABOLISM | | | KEGG_LYSINE_DEGRADATION | |
| | | | | | KEGG_FATTY_ACID_METABOLISM | WP_NUCLEAR_RECEPTORS_METAPATHWAY | | | |
| | | | | | KEGG_FATTY_ACID_METABOLISM | | | KEGG_LYSINE_DEGRADATION | |
| | | | | | KEGG_FATTY_ACID_METABOLISM | | | | |
| | WP_ONE_CARBON_METABOLISM_AND_RELATED_PATHWAYS | | | | | | | | WP_BIOGENIC_AMINE_SYNTHESIS |

Bulk RNA-seq DOWN

REACTOME_FORMATION_OF_FIBRIN_CLOT_CLOTTING_CASCADE

WP_NUCLEAR_RECEPTORS_METAPATHWAY

REACTOME_METABOLISM_OF_VITAMINS_AND_COFACTORS

KEGG_PYRUVATE_METABOLISM

WP_NUCLEAR_RECEPTORS_METAPATHWAY

WP_ONE_CARBON_METABOLISM_AND_RELATED_PATHWAYS

KEGG_CALCIUM_SIGNALING_PATHWAY

KEGG_LYSINE_DEGRADATION

KEGG_PYRUVATE_METABOLISM

WP_ONE_CARBON_METABOLISM_AND_RELATED_PATHWAYS REACTOME_METABOLISM_OF_VITAMINS_AND_COFACTORS

WP_NUCLEAR_RECEPTORS_METAPATHWAY
WP_NUCLEAR_RECEPTORS_METAPATHWAY
WP_NUCLEAR_RECEPTORS_METAPATHWAY

KEGG_FATTY_ACID_METABOLISM

REACTOME_METABOLISM_OF_VITAMINS_AND_COFACTORS
REACTOME_METABOLISM_OF_VITAMINS_AND_COFACTORS
REACTOME_METABOLISM_OF_VITAMINS_AND_COFACTORS
REACTOME_METABOLISM_OF_VITAMINS_AND_COFACTORS
REACTOME_METABOLISM_OF_VITAMINS_AND_COFACTORS
REACTOME_METABOLISM_OF_VITAMINS_AND_COFACTORS

KEGG_CALCIUM_SIGNALING_PATHWAY

KEGG_CALCIUM_SIGNALING_PATHWAY

KEGG_PYRUVATE_METABOLISM
KEGG_PYRUVATE_METABOLISM

REACTOME_ION_HOMEOSTASIS

KEGG_CALCIUM_SIGNALING_PATHWAY

REACTOME_FORMATION_OF_FIBRIN_CLOT_CLOTTING_CASCADE
REACTOME_FORMATION_OF_FIBRIN_CLOT_CLOTTING_CASCADE
REACTOME_FORMATION_OF_FIBRIN_CLOT_CLOTTING_CASCADE
REACTOME_FORMATION_OF_FIBRIN_CLOT_CLOTTING_CASCADE
REACTOME_FORMATION_OF_FIBRIN_CLOT_CLOTTING_CASCADE

WP_NUCLEAR_RECEPTORS_METAPATHWAY

KEGG_CALCIUM_SIGNALING_PATHWAY
KEGG_CALCIUM_SIGNALING_PATHWAY

Bulk RNA-seq DOWN

KEGG_CALCIUM_SIGNALING_PATHWAY

Bulk RNA-seq DOWN

Bulk RNA-seq DOWN

Bulk RNA-seq DOWN

Bulk RNA-seq DOWN

Bulk RNA-seq DOWN

Bulk RNA-seq DOWN

Bulk RNA-seq DOWN

REACTOME_PHASE_I_FUNCTIONALIZATION_OF_COMPOUNDS REACTOME_PEROXISOMAL_LIPID_METABOLISM KEGG_RETINOL_METABOLISM REACTOME_CATION_COUPLED_CHLORIDE_COTRANSPORTERS REACTOME_FRUCTOSE_METABOLISM WP_DISORDERS_OF_THE_KREBS_CYCLE WP_OXYSTEROLS_DERIVED_FROM_CHOLESTEROL WP_TCA_CYCLE_AKA_KREBS_OR_CITRIC_ACID_CYCLE KEGG_METABOLISM_OF_XENOBIOTICS_BY_CYTOCHROME_P450

WP_TCA_CYCLE_AKA_KREBS_OR_CITRIC_ACID_CYCLE

WP_DISORDERS_OF_THE_KREBS_CYCLE

REACTOME_CATION_COUPLED_CHLORIDE_COTRANSPORTERS
REACTOME_CATION_COUPLED_CHLORIDE_COTRANSPORTERS
REACTOME_CATION_COUPLED_CHLORIDE_COTRANSPORTERS

REACTOME_PEROXISOMAL_LIPID_METABOLISM

REACTOME_FRUCTOSE_METABOLISM

WP_TCA_CYCLE_AKA_KREBS_OR_CITRIC_ACID_CYCLE

REACTOME_PHASE_I_FUNCTIONALIZATION_OF_COMPOUNDS KEGG_RETINOL_METABOLISM KEGG_METABOLISM_OF_XENOBIOTICS_BY_CYTOCHROME_P450

REACTOME_PEROXISOMAL_LIPID_METABOLISM WP_OXYSTEROLS_DERIVED_FROM_CHOLESTEROL

REACTOME_PHASE_I_FUNCTIONALIZATION_OF_COMPOUNDS

REACTOME_PHASE_I_FUNCTIONALIZATION_OF_COMPOUNDS

Bulk RNA-seq DOWN

| | | | | | | | |
|---|---|---|---|---|---|---|---|
| REACTOME_PHASE_I_FUNCTIONALIZATION_OF_COMPOUNDS | | | | | | | |
| | | | REACTOME_FRUCTOSE_METABOLISM | | | | |
| | | | | | WP_OXYSTEROLS_DERIVED_FROM_CHOLESTEROL | | |
| REACTOME_PHASE_I_FUNCTIONALIZATION_OF_COMPOUNDS | | | | | | | |
| | | | | WP_DISORDERS_OF_THE_KREBS_CYCLE | | WP_TCA_CYCLE_AKA_KREBS_OR_CITRIC_ACID_CYCLE | |
| | REACTOME_PEROXISOMAL_LIPID_METABOLISM | | | | | | |
| | REACTOME_PEROXISOMAL_LIPID_METABOLISM | | | | | | |
| | | | | | WP_OXYSTEROLS_DERIVED_FROM_CHOLESTEROL | | |
| | REACTOME_PEROXISOMAL_LIPID_METABOLISM | | | | | | |
| | | KEGG_RETINOL_METABOLISM | | | | | |
| | | | | | WP_OXYSTEROLS_DERIVED_FROM_CHOLESTEROL | | |
| | | | | | WP_OXYSTEROLS_DERIVED_FROM_CHOLESTEROL | | |
| REACTOME_PHASE_I_FUNCTIONALIZATION_OF_COMPOUNDS | | KEGG_RETINOL_METABOLISM | | | WP_OXYSTEROLS_DERIVED_FROM_CHOLESTEROL | | KEGG_METABOLISM_OF_XENOBIOTICS_BY_CYTOCHROME_P450 |
| | | | | WP_DISORDERS_OF_THE_KREBS_CYCLE | | WP_TCA_CYCLE_AKA_KREBS_OR_CITRIC_ACID_CYCLE | |
| REACTOME_PHASE_I_FUNCTIONALIZATION_OF_COMPOUNDS | | KEGG_RETINOL_METABOLISM | | | | | KEGG_METABOLISM_OF_XENOBIOTICS_BY_CYTOCHROME_P450 |
| | | | | | | | KEGG_METABOLISM_OF_XENOBIOTICS_BY_CYTOCHROME_P450 |
| | | | | | | | KEGG_METABOLISM_OF_XENOBIOTICS_BY_CYTOCHROME_P450 |
| REACTOME_PHASE_I_FUNCTIONALIZATION_OF_COMPOUNDS | | KEGG_RETINOL_METABOLISM | | | | | KEGG_METABOLISM_OF_XENOBIOTICS_BY_CYTOCHROME_P450 |
| REACTOME_PHASE_I_FUNCTIONALIZATION_OF_COMPOUNDS | | | | | | | |
| | | | REACTOME_FRUCTOSE_METABOLISM | | | | |

Bulk RNA-seq DOWN

KEGG_RETINOL_METABOLISM
KEGG_RETINOL_METABOLISM

KEGG_METABOLISM_OF_XENOBIOTICS_BY_CYTOCHROME_P450

Bulk RNA-seq DOWN

Bulk RNA-seq DOWN

Bulk RNA-seq DOWN

Bulk RNA-seq DOWN

Bulk RNA-seq DOWN

Bulk RNA-seq DOWN

Bulk RNA-seq DOWN

| REACTOME_AFLATOXIN_ACTIVATION_AND_DETOXIFICATION | KEGG_CYSTEINE_AND_METHIONINE_METABOLISM | WP_NRF2_PATHWAY | HALLMARK_HEME_METABOLISM | REACTOME_CLASS_I_PEROXISOMAL_MEMBRANE_PROTEIN_IMPORT | REACTOME_RAS_ACTIVATION_UPON_CA2_INFLUX_THROUGH_NMDA_RECEPTOR | REACTOME_MULTIFUNCTIONAL_ANION_EXCHANGERS | REACTOME_NEGATIVE_REGULATION_OF_NMDA_RECEPTOR_MEDIATED_NEURONAL_TRANSMISSION |
|---|---|---|---|---|---|---|---|
| | | | HALLMARK_HEME_METABOLISM | | | | |
| | KEGG_CYSTEINE_AND_METHIONINE_METABOLISM | | | | | | |
| | | | | REACTOME_CLASS_I_PEROXISOMAL_MEMBRANE_PROTEIN_IMPORT | | | |
| | KEGG_CYSTEINE_AND_METHIONINE_METABOLISM | | | | | | |
| | | | HALLMARK_HEME_METABOLISM | | | | |
| | KEGG_CYSTEINE_AND_METHIONINE_METABOLISM | | | | | | |
| | | WP_NRF2_PATHWAY | | | | | |
| | | WP_NRF2_PATHWAY | | | | | |
| | | | HALLMARK_HEME_METABOLISM | | | | |
| | | | | | | REACTOME_MULTIFUNCTIONAL_ANION_EXCHANGERS | |
| | | WP_NRF2_PATHWAY | | | | REACTOME_MULTIFUNCTIONAL_ANION_EXCHANGERS | |
| | | | | | | REACTOME_MULTIFUNCTIONAL_ANION_EXCHANGERS | |
| | | WP_NRF2_PATHWAY | | | | | |
| | | WP_NRF2_PATHWAY | | | | | |
| | | WP_NRF2_PATHWAY | | | | | |
| | | WP_NRF2_PATHWAY | | | | | |
| | | WP_NRF2_PATHWAY | | | | | |
| | | | HALLMARK_HEME_METABOLISM | | | | |
| | | | | | REACTOME_RAS_ACTIVATION_UPON_CA2_INFLUX_THROUGH_NMDA_RECEPTOR | | REACTOME_NEGATIVE_REGULATION_OF_NMDA_RECEPTOR_MEDIATED_NEURONAL_TRANSMISSION |
| | | | | | REACTOME_RAS_ACTIVATION_UPON_CA2_INFLUX_THROUGH_NMDA_RECEPTOR | | REACTOME_NEGATIVE_REGULATION_OF_NMDA_RECEPTOR_MEDIATED_NEURONAL_TRANSMISSION |
| | | | | | REACTOME_RAS_ACTIVATION_UPON_CA2_INFLUX_THROUGH_NMDA_RECEPTOR | | REACTOME_NEGATIVE_REGULATION_OF_NMDA_RECEPTOR_MEDIATED_NEURONAL_TRANSMISSION |
| | | | | REACTOME_CLASS_I_PEROXISOMAL_MEMBRANE_PROTEIN_IMPORT | | | |

Bulk RNA-seq DOWN

REACTOME_AFLATOXIN_ACTIVATION_AND_DETOXIFICATION

REACTOME_CLASS_I_PEROXISOMAL_MEMBRANE_PROTEIN_IMPORT

REACTOME_CLASS_I_PEROXISOMAL_MEMBRANE_PROTEIN_IMPORT

REACTOME_AFLATOXIN_ACTIVATION_AND_DETOXIFICATION

HALLMARK_HEME_METABOLISM

KEGG_CYSTEINE_AND_METHIONINE_METABOLISM

WP_NRF2_PATHWAY
WP_NRF2_PATHWAY

REACTOME_AFLATOXIN_ACTIVATION_AND_DETOXIFICATION

REACTOME_AFLATOXIN_ACTIVATION_AND_DETOXIFICATION

REACTOME_RAS_ACTIVATION_UPON_CA2_INFLUX_THROUGH_NMDA_RECEPTOR

REACTOME_NEGATIVE_REGULATION_OF_NMDA_RECEPTOR_MEDIATED_NEURONAL_TRANSMISSION

Bulk RNA-seq DOWN

| | |
|---|---|
| KEGG_CYSTEINE_AND_METHIONINE_METABOLISM | |
| | HALLMARK_HEME_METABOLISM |
| | HALLMARK_HEME_METABOLISM |
| | HALLMARK_HEME_METABOLISM |
| | HALLMARK_HEME_METABOLISM |
| | HALLMARK_HEME_METABOLISM |
| | HALLMARK_HEME_METABOLISM |
| | HALLMARK_HEME_METABOLISM |

Bulk RNA-seq DOWN

Bulk RNA-seq DOWN

Bulk RNA-seq DOWN

Bulk RNA-seq DOWN

Bulk RNA-seq DOWN

Bulk RNA-seq DOWN

Bulk RNA-seq DOWN

| REACTOME_UNBLOCKING_OF_NMDA_RECEPTORS_GLUTAMATE_BINDING_AND_ACTIVATION | REACTOME_NEUROTRANSMITTER_RECEPTORS_AND_POSTSYNAPTIC_SIGNAL_TRANSMISSION | REACTOME_ION_TRANSPORT_BY_P_TYPE_ATPASES | REACTOME_SENSORY_PROCESSING_OF_SOUND_BY_OUTER_HAIR_CELLS_OF_THE_COCHLEA | REACTOME_SENSORY_PROCESSING_OF_SOUND | WP_CALCIUM_REGULATION_IN_THE_CARDIAC_CELL |
|---|---|---|---|---|---|
| | | | | | WP_CALCIUM_REGULATION_IN_THE_CARDIAC_CELL |
| | | REACTOME_ION_TRANSPORT_BY_P_TYPE_ATPASES | | | WP_CALCIUM_REGULATION_IN_THE_CARDIAC_CELL |
| REACTOME_UNBLOCKING_OF_NMDA_RECEPTORS_GLUTAMATE_BINDING_AND_ACTIVATION | REACTOME_NEUROTRANSMITTER_RECEPTORS_AND_POSTSYNAPTIC_SIGNAL_TRANSMISSION | REACTOME_ION_TRANSPORT_BY_P_TYPE_ATPASES | | | WP_CALCIUM_REGULATION_IN_THE_CARDIAC_CELL |
| REACTOME_UNBLOCKING_OF_NMDA_RECEPTORS_GLUTAMATE_BINDING_AND_ACTIVATION | REACTOME_NEUROTRANSMITTER_RECEPTORS_AND_POSTSYNAPTIC_SIGNAL_TRANSMISSION | REACTOME_ION_TRANSPORT_BY_P_TYPE_ATPASES | | | WP_CALCIUM_REGULATION_IN_THE_CARDIAC_CELL |
| REACTOME_UNBLOCKING_OF_NMDA_RECEPTORS_GLUTAMATE_BINDING_AND_ACTIVATION | REACTOME_NEUROTRANSMITTER_RECEPTORS_AND_POSTSYNAPTIC_SIGNAL_TRANSMISSION | REACTOME_ION_TRANSPORT_BY_P_TYPE_ATPASES | | | WP_CALCIUM_REGULATION_IN_THE_CARDIAC_CELL |
| | REACTOME_NEUROTRANSMITTER_RECEPTORS_AND_POSTSYNAPTIC_SIGNAL_TRANSMISSION | | | | WP_CALCIUM_REGULATION_IN_THE_CARDIAC_CELL |
| | | REACTOME_ION_TRANSPORT_BY_P_TYPE_ATPASES | REACTOME_SENSORY_PROCESSING_OF_SOUND_BY_OUTER_HAIR_CELLS_OF_THE_COCHLEA | REACTOME_SENSORY_PROCESSING_OF_SOUND | WP_CALCIUM_REGULATION_IN_THE_CARDIAC_CELL |
| | | REACTOME_ION_TRANSPORT_BY_P_TYPE_ATPASES | | | |

Bulk RNA-seq DOWN

REACTOME_SENSORY_PROCESSING_OF_SOUND

REACTOME_UNBLOCKING_OF_NMDA_RECEPTORS_GLUTAMATE_BINDING_AND_ACTIVATION
REACTOME_NEUROTRANSMITTER_RECEPTORS_AND_POSTSYNAPTIC_SIGNAL_TRANSMISSION
REACTOME_NEUROTRANSMITTER_RECEPTORS_AND_POSTSYNAPTIC_SIGNAL_TRANSMISSION
REACTOME_NEUROTRANSMITTER_RECEPTORS_AND_POSTSYNAPTIC_SIGNAL_TRANSMISSION
REACTOME_NEUROTRANSMITTER_RECEPTORS_AND_POSTSYNAPTIC_SIGNAL_TRANSMISSION
REACTOME_NEUROTRANSMITTER_RECEPTORS_AND_POSTSYNAPTIC_SIGNAL_TRANSMISSION
REACTOME_NEUROTRANSMITTER_RECEPTORS_AND_POSTSYNAPTIC_SIGNAL_TRANSMISSION
REACTOME_NEUROTRANSMITTER_RECEPTORS_AND_POSTSYNAPTIC_SIGNAL_TRANSMISSION
REACTOME_NEUROTRANSMITTER_RECEPTORS_AND_POSTSYNAPTIC_SIGNAL_TRANSMISSION

REACTOME_SENSORY_PROCESSING_OF_SOUND_BY_OUTER_HAIR_CELLS_OF_THE_COCHLEA
REACTOME_SENSORY_PROCESSING_OF_SOUND

WP_CALCIUM_REGULATION_IN_THE_CARDIAC_CELL

Bulk RNA-seq DOWN

| | | |
|---|---|---|
| REACTOME_SENSORY_PROCESSING_OF_SOUND_BY_OUTER_HAIR_CELLS_OF_THE_COCHLEA | REACTOME_SENSORY_PROCESSING_OF_SOUND | |
| REACTOME_SENSORY_PROCESSING_OF_SOUND_BY_OUTER_HAIR_CELLS_OF_THE_COCHLEA | REACTOME_SENSORY_PROCESSING_OF_SOUND | |
| REACTOME_SENSORY_PROCESSING_OF_SOUND_BY_OUTER_HAIR_CELLS_OF_THE_COCHLEA | REACTOME_SENSORY_PROCESSING_OF_SOUND | |
| REACTOME_SENSORY_PROCESSING_OF_SOUND_BY_OUTER_HAIR_CELLS_OF_THE_COCHLEA | REACTOME_SENSORY_PROCESSING_OF_SOUND | |
| | | WP_CALCIUM_REGULATION_IN_THE_CARDIAC_CELL |
| | | WP_CALCIUM_REGULATION_IN_THE_CARDIAC_CELL |

Bulk RNA-seq DOWN

Bulk RNA-seq DOWN

Bulk RNA-seq DOWN

Bulk RNA-seq DOWN

Bulk RNA-seq DOWN

Bulk RNA-seq DOWN

Bulk RNA-seq DOWN

| REACTOME_CITRIC_ACID_CYCLE_TCA_CYCLE | REACTOME_DEFECTS_IN_VITAMIN_AND_COFACTOR_METABOLISM | REACTOME_CAMK_IV_MEDIATED_PHOSPHORYLATION_OF_CREB | BIOCARTA_INTRINSIC_PATHWAY | REACTOME_LONG_TERM_POTENTIATION | WP_VITAMIN_DSENSITIVE_CALCIUM_SIGNALING_IN_DEPRESSION | REACTOME_DAG_AND_IP3_SIGNALING | REACTOME_LYSINE_CATABOLISM | REACTOME_MUSCLE_CONTRACTION |
|---|---|---|---|---|---|---|---|---|
| | REACTOME_DEFECTS_IN_VITAMIN_AND_COFACTOR_METABOLISM | | | | | | | |
| | | | | | | | REACTOME_LYSINE_CATABOLISM | |
| | | | | | | | REACTOME_LYSINE_CATABOLISM | |
| REACTOME_CITRIC_ACID_CYCLE_TCA_CYCLE | | | | | | | REACTOME_LYSINE_CATABOLISM | |
| | | | | | WP_VITAMIN_DSENSITIVE_CALCIUM_SIGNALING_IN_DEPRESSION | | | REACTOME_MUSCLE_CONTRACTION |
| | REACTOME_DEFECTS_IN_VITAMIN_AND_COFACTOR_METABOLISM | | | | | | | |
| | | | | | | | | REACTOME_MUSCLE_CONTRACTION |
| | REACTOME_DEFECTS_IN_VITAMIN_AND_COFACTOR_METABOLISM | | | | | | | |
| | | REACTOME_CAMK_IV_MEDIATED_PHOSPHORYLATION_OF_CREB | | REACTOME_LONG_TERM_POTENTIATION | | REACTOME_DAG_AND_IP3_SIGNALING | | REACTOME_MUSCLE_CONTRACTION |
| | | REACTOME_CAMK_IV_MEDIATED_PHOSPHORYLATION_OF_CREB | | REACTOME_LONG_TERM_POTENTIATION | | REACTOME_DAG_AND_IP3_SIGNALING | | REACTOME_MUSCLE_CONTRACTION |
| | | REACTOME_CAMK_IV_MEDIATED_PHOSPHORYLATION_OF_CREB | | REACTOME_LONG_TERM_POTENTIATION | | REACTOME_DAG_AND_IP3_SIGNALING | | REACTOME_MUSCLE_CONTRACTION |
| | | | | | WP_VITAMIN_DSENSITIVE_CALCIUM_SIGNALING_IN_DEPRESSION | | | REACTOME_MUSCLE_CONTRACTION |
| REACTOME_CITRIC_ACID_CYCLE_TCA_CYCLE | | | | | | | | |

Bulk RNA-seq DOWN

BIOCARTA_INTRINSIC_PATHWAY

REACTOME_CITRIC_ACID_CYCLE_TCA_CYCLE

REACTOME_CITRIC_ACID_CYCLE_TCA_CYCLE

REACTOME_DEFECTS_IN_VITAMIN_AND_COFACTOR_METABOLISM

REACTOME_MUSCLE_CONTRACTION

REACTOME_LONG_TERM_POTENTIATION

REACTOME_MUSCLE_CONTRACTION

WP_VITAMIN_DSENSITIVE_CALCIUM_SIGNALING_IN_DEPRESSION REACTOME_DAG_AND_IP3_SIGNALING REACTOME_MUSCLE_CONTRACTION

REACTOME_MUSCLE_CONTRACTION

REACTOME_MUSCLE_CONTRACTION

BIOCARTA_INTRINSIC_PATHWAY

BIOCARTA_INTRINSIC_PATHWAY

Bulk RNA-seq DOWN

REACTOME_DAG_AND_IP3_SIGNALING

BIOCARTA_INTRINSIC_PATHWAY

WP_VITAMIN_DSENSITIVE_CALCIUM_SIGNALING_IN_DEPRESSION

WP_VITAMIN_DSENSITIVE_CALCIUM_SIGNALING_IN_DEPRESSION

Bulk RNA-seq DOWN

Bulk RNA-seq DOWN

Bulk RNA-seq DOWN

Bulk RNA-seq DOWN

Bulk RNA-seq DOWN

Bulk RNA-seq DOWN

Bulk RNA-seq DOWN

| | |
|---|---|
| REACTOME_ASSEMBLY_AND_CELL_SURFACE_PRESENTATION_OF_NMDA_RECEPTORS | WP_RENIN_ANGIOTENSIN_ALDOSTERONE_SYSTEM_RAAS |

| | |
|---|---|
| REACTOME_ASSEMBLY_AND_CELL_SURFACE_PRESENTATION_OF_NMDA_RECEPTORS | WP_RENIN_ANGIOTENSIN_ALDOSTERONE_SYSTEM_RAAS |
| REACTOME_ASSEMBLY_AND_CELL_SURFACE_PRESENTATION_OF_NMDA_RECEPTORS | WP_RENIN_ANGIOTENSIN_ALDOSTERONE_SYSTEM_RAAS |
| REACTOME_ASSEMBLY_AND_CELL_SURFACE_PRESENTATION_OF_NMDA_RECEPTORS | WP_RENIN_ANGIOTENSIN_ALDOSTERONE_SYSTEM_RAAS |

Bulk RNA-seq DOWN

REACTOME_ASSEMBLY_AND_CELL_SURFACE_PRESENTATION_OF_NMDA_RECEPTORS
REACTOME_ASSEMBLY_AND_CELL_SURFACE_PRESENTATION_OF_NMDA_RECEPTORS

WP_RENIN_ANGIOTENSIN_ALDOSTERONE_SYSTEM_RAAS

Bulk RNA-seq DOWN

WP_RENIN_ANGIOTENSIN_ALDOSTERONE_SYSTEM_RAAS
