## Supplemental Table S5 for "Transcriptomic responses of the human kidney to acute injury at single cell resolution"

**Marker gene definition:** A gene is considered to be a marker gene of the respective cluster if it is significantly upregulated when compared to cells from healthy tissue of the respective cell type (as defined by an adjusted p-value<0.05 provided by the FindAllMarkers Seurat function) and if its average expression is higher than in all other AKI-associated cell states o the respective cell type.

Oxidative stress
Source:
https://www.gsea-msigdb.org/gsea/msigdb/cards/HALLMARK_REACTIVE_OXYGEN_SPECIES_PATHWAY.html
https://www.gsea-msigdb.org/gsea/msigdb/cards/WP_NRF2_PATHWAY.html
https://www.genenames.org/data/genegroup/#!/group/638
1 = marker gene of the respective cluster, 0 = no marker gene of the respective cluster (see methods)

| | PT-New1 | TAL-New1 | DCT-New1 |
|---|---|---|---|
| FTL | 1 | 1 | 1 |
| GCLM | 1 | 1 | 1 |
| GPX3 | 1 | 1 | 1 |
| MT1A | 1 | 1 | 1 |
| MT1F | 1 | 1 | 1 |
| MT1G | 1 | 1 | 1 |
| MT1H | 1 | 1 | 1 |
| MT1HL1 | 1 | 1 | 1 |
| MT1M | 1 | 1 | 1 |
| MT1X | 1 | 1 | 1 |
| SLC39A5 | 1 | 1 | 1 |
| CES2 | 1 | 1 | 1 |
| SLC2A8 | 1 | 1 | 1 |
| ABCC2 | 0 | 1 | 1 |
| CYP4A11 | 0 | 1 | 1 |
| DNAJB1 | 1 | 0 | 1 |
| FES | 0 | 1 | 1 |
| GGT1 | 0 | 1 | 1 |
| GPX2 | 1 | 1 | 0 |
| GSTA2 | 1 | 0 | 1 |
| HMOX1 | 1 | 0 | 1 |
| MT1E | 1 | 0 | 1 |
| MT2A | 1 | 0 | 1 |
| SLC2A2 | 0 | 1 | 1 |
| SLC5A10 | 0 | 1 | 1 |
| SLC5A12 | 0 | 1 | 1 |
| SLC5A2 | 0 | 1 | 1 |
| SLC5A4 | 1 | 1 | 0 |
| SLC5A5 | 0 | 1 | 1 |
| SLC6A13 | 0 | 1 | 1 |
| SLC6A19 | 0 | 1 | 1 |
| ATOX1 | 1 | 0 | 1 |
| CDKN2D | 1 | 0 | 1 |
| FTH1 | 1 | 0 | 1 |
| GLRX | 1 | 0 | 1 |
| HMOX2 | 1 | 0 | 1 |
| LAMTOR5 | 1 | 0 | 1 |
| NQO1 | 1 | 0 | 1 |
| SLC39A14 | 1 | 0 | 1 |
| SLC39A4 | 1 | 0 | 1 |
| SOD1 | 1 | 0 | 1 |
| TGFB2 | 0 | 1 | 1 |
| TXN | 1 | 0 | 1 |
| UGT2B7 | 0 | 1 | 1 |
| CES3 | 0 | 0 | 1 |
| G6PD | 1 | 0 | 0 |
| GSTA1 | 1 | 0 | 0 |
| HHEX | 0 | 1 | 0 |
| KEAP1 | 1 | 0 | 0 |
| ME1 | 0 | 0 | 1 |
| MGST1 | 0 | 0 | 1 |
| MGST2 | 0 | 0 | 1 |
| MPO | 1 | 0 | 0 |
| SERPINA1 | 0 | 0 | 1 |
| SLC2A10 | 0 | 1 | 0 |
| SLC2A3 | 0 | 1 | 0 |
| SLC2A9 | 0 | 0 | 1 |
| SLC5A11 | 0 | 0 | 1 |
| SLC5A8 | 0 | 0 | 1 |
| SLC5A9 | 0 | 0 | 1 |
| SLC6A2 | 1 | 0 | 0 |
| SLC6A8 | 1 | 0 | 0 |
| SOD3 | 0 | 0 | 1 |
| ABCC3 | 0 | 0 | 1 |
| BLVRB | 1 | 0 | 0 |
| CAT | 1 | 0 | 0 |
| CBR1 | 1 | 0 | 0 |
| CYP2A6 | 1 | 0 | 0 |
| EGR1 | 0 | 0 | 1 |
| EPHA3 | 0 | 0 | 1 |
| ERCC2 | 0 | 0 | 1 |
| GLRX2 | 0 | 0 | 1 |
| GPX4 | 0 | 0 | 1 |
| HSP90AA1 | 1 | 0 | 0 |
| HSP90AB1 | 1 | 0 | 0 |
| HSPA1A | 1 | 0 | 0 |
| LSP1 | 0 | 1 | 0 |
| MAFG | 1 | 0 | 0 |
| MGST3 | 1 | 0 | 0 |
| MT3 | 1 | 0 | 0 |

| | | | |
|---|---|---|---|
| NDUFB4 | 1 | 0 | 0 |
| NDUFS2 | 1 | 0 | 0 |
| OXSR1 | 0 | 0 | 1 |
| PDGFB | 0 | 0 | 1 |
| PDLIM1 | 0 | 0 | 1 |
| PGD | 1 | 0 | 0 |
| PRDX1 | 1 | 0 | 0 |
| PRDX6 | 1 | 0 | 0 |
| PRNP | 0 | 0 | 1 |
| RXRA | 1 | 0 | 0 |
| SLC2A4 | 0 | 0 | 1 |
| SLC2A6 | 0 | 0 | 1 |
| SLC39A7 | 0 | 0 | 1 |
| SLC5A1 | 0 | 0 | 1 |
| SLC5A6 | 0 | 1 | 0 |
| SLC6A1 | 0 | 1 | 0 |
| SLC6A11 | 0 | 1 | 0 |
| SLC6A18 | 0 | 1 | 0 |
| SLC6A6 | 0 | 0 | 1 |
| SLC6A9 | 0 | 0 | 1 |
| SLC7A11 | 0 | 0 | 1 |
| SRXN1 | 1 | 0 | 0 |
| STK25 | 0 | 0 | 1 |
| TXNRD1 | 0 | 0 | 1 |
| TXNRD3 | 0 | 1 | 0 |
| ABCC4 | 0 | 0 | 0 |
| CBR3 | 0 | 0 | 0 |
| CES5A | 0 | 0 | 0 |
| FGF13 | 0 | 0 | 0 |
| GCLC | 0 | 0 | 0 |
| GGT2 | 0 | 0 | 0 |
| GSR | 0 | 0 | 0 |
| GSTM2 | 0 | 0 | 0 |
| HGF | 0 | 0 | 0 |
| MSRA | 0 | 0 | 0 |
| PPARD | 0 | 0 | 0 |
| PTPA | 0 | 0 | 0 |
| SLC2A12 | 0 | 0 | 0 |
| SLC2A5 | 0 | 0 | 0 |
| SLC6A16 | 0 | 0 | 0 |
| TXNRD2 | 0 | 0 | 0 |

Hypoxia response
Source:
https://www.gsea-msigdb.org/gsea/msigdb/cards/HALLMARK_HYPOXIA.html
1 = marker gene of the respective cluster, 0 = no marker gene of the respective cluster (see methods)

| | PT-New2 | TAL-New2 | DCT-New2 |
|---|---|---|---|
| ERO1A | 1 | 1 | 1 |
| GBE1 | 1 | 1 | 1 |
| GPI | 1 | 1 | 1 |
| MAFF | 1 | 1 | 1 |
| MYH9 | 1 | 1 | 1 |
| P4HA1 | 1 | 1 | 1 |
| PDK1 | 1 | 1 | 1 |
| PFKP | 1 | 1 | 1 |
| PPFIA4 | 1 | 1 | 1 |
| WSB1 | 1 | 1 | 1 |
| ZNF292 | 1 | 1 | 1 |
| ADM | 0 | 1 | 1 |
| AMPD3 | 0 | 1 | 1 |
| ATF3 | 1 | 0 | 1 |
| BHLHE40 | 0 | 1 | 1 |
| BNIP3L | 0 | 1 | 1 |
| CDKN1A | 0 | 1 | 1 |
| DDIT3 | 0 | 1 | 1 |
| ENO2 | 0 | 1 | 1 |
| ETS1 | 1 | 0 | 1 |
| HK1 | 0 | 1 | 1 |
| HK2 | 0 | 1 | 1 |
| KDM3A | 1 | 1 | 0 |
| KLF6 | 1 | 0 | 1 |
| NDRG1 | 0 | 1 | 1 |
| PAM | 1 | 1 | 0 |
| PDK3 | 0 | 1 | 1 |
| PFKFB3 | 0 | 1 | 1 |
| PGK1 | 0 | 1 | 1 |
| PPARGC1A | 0 | 1 | 1 |
| PPP1R15A | 0 | 1 | 1 |
| PPP1R3C | 0 | 1 | 1 |
| PYGM | 1 | 0 | 1 |
| RRAGD | 0 | 1 | 1 |
| SIAH2 | 1 | 1 | 0 |
| SLC2A1 | 0 | 1 | 1 |
| TES | 1 | 0 | 1 |
| VEGFA | 0 | 1 | 1 |
| XPNPEP1 | 1 | 1 | 0 |
| DTNA | 1 | 0 | 1 |
| EFNA3 | 0 | 1 | 1 |
| EGFR | 1 | 1 | 0 |
| FOXO3 | 1 | 0 | 1 |
| LARGE1 | 1 | 0 | 1 |
| NEDD4L | 0 | 1 | 1 |
| NOCT | 1 | 1 | 0 |
| SCARB1 | 0 | 1 | 1 |
| TNFAIP3 | 1 | 0 | 1 |

| | | | |
|---|---|---|---|
| TPST2 | 0 | 1 | 1 |
| ADORA2B | 0 | 1 | 0 |
| ANGPTL4 | 0 | 1 | 0 |
| ANKZF1 | 0 | 1 | 0 |
| CDKN1C | 0 | 1 | 0 |
| CXCR4 | 0 | 1 | 0 |
| DDIT4 | 0 | 1 | 0 |
| EDN2 | 0 | 1 | 0 |
| FAM162A | 0 | 1 | 0 |
| GCK | 1 | 0 | 0 |
| GYS1 | 0 | 1 | 0 |
| IGFBP3 | 0 | 1 | 0 |
| IRS2 | 0 | 0 | 1 |
| JMJD6 | 0 | 1 | 0 |
| KLHL24 | 0 | 0 | 1 |
| MXI1 | 0 | 1 | 0 |
| P4HA2 | 0 | 1 | 0 |
| PIM1 | 0 | 0 | 1 |
| PLAUR | 1 | 0 | 0 |
| PLIN2 | 0 | 1 | 0 |
| RBPJ | 0 | 1 | 0 |
| SAP30 | 0 | 1 | 0 |
| SERPINE1 | 0 | 0 | 1 |
| SLC6A6 | 0 | 1 | 0 |
| TGFBI | 0 | 0 | 1 |
| TIPARP | 0 | 1 | 0 |
| TPD52 | 0 | 1 | 0 |
| VLDLR | 0 | 1 | 0 |
| AK4 | 0 | 1 | 0 |
| AKAP12 | 1 | 0 | 0 |
| ANXA2 | 1 | 0 | 0 |
| BCL2 | 0 | 0 | 1 |
| COL5A1 | 0 | 0 | 1 |
| CSRP2 | 0 | 1 | 0 |
| EXT1 | 1 | 0 | 0 |
| F3 | 0 | 1 | 0 |
| HDLBP | 0 | 0 | 1 |
| HEXA | 0 | 1 | 0 |
| HMOX1 | 0 | 1 | 0 |
| INHA | 0 | 1 | 0 |
| KLF7 | 1 | 0 | 0 |
| LOX | 0 | 1 | 0 |
| PDGFB | 1 | 0 | 0 |
| PGAM2 | 0 | 1 | 0 |
| PGF | 0 | 0 | 1 |
| PHKG1 | 0 | 0 | 1 |
| PRKCA | 1 | 0 | 0 |
| SELENBP1 | 0 | 0 | 1 |
| SLC2A5 | 0 | 1 | 0 |
| STC1 | 0 | 1 | 0 |
| STC2 | 0 | 1 | 0 |
| UGP2 | 0 | 1 | 0 |
| VHL | 0 | 0 | 1 |

| | | | |
|---|---|---|---|
| CCNG2 | 0 | 0 | 0 |
| CHST2 | 0 | 0 | 0 |
| DUSP1 | 0 | 0 | 0 |
| ENO1 | 0 | 0 | 0 |
| ERRFI1 | 0 | 0 | 0 |
| FOS | 0 | 0 | 0 |
| FOSL2 | 0 | 0 | 0 |
| GAPDHS | 0 | 0 | 0 |
| HAS1 | 0 | 0 | 0 |
| HSPA5 | 0 | 0 | 0 |
| IER3 | 0 | 0 | 0 |
| ILVBL | 0 | 0 | 0 |
| JUN | 0 | 0 | 0 |
| LDHA | 0 | 0 | 0 |
| NCAN | 0 | 0 | 0 |
| NDST2 | 0 | 0 | 0 |
| NFIL3 | 0 | 0 | 0 |
| PFKL | 0 | 0 | 0 |
| PGM1 | 0 | 0 | 0 |
| TMEM45A | 0 | 0 | 0 |
| ZFP36 | 0 | 0 | 0 |

Interferon gamma signaling and ribosomal protein-coding genes
Source:
https://www.gsea-msigdb.org/gsea/msigdb/cards/HALLMARK_INTERFERON_GAMMA_RESPONSE.html
http://ribosome.med.miyazaki-u.ac.jp/rpg.cgi?mode=orglist&org=Homo%20sapiens
1 = marker gene of the respective cluster, 0 = no marker gene of the respective cluster (see methods)

| | PT-New3 | TAL-New3 | DCT-New3 |
|---|---|---|---|
| BTG1 | 1 | 1 | 1 |
| RPL10 | 1 | 1 | 1 |
| RPL10A | 1 | 1 | 1 |
| RPL11 | 1 | 1 | 1 |
| RPL12 | 1 | 1 | 1 |
| RPL13 | 1 | 1 | 1 |
| RPL13A | 1 | 1 | 1 |
| RPL14 | 1 | 1 | 1 |
| RPL15 | 1 | 1 | 1 |
| RPL18 | 1 | 1 | 1 |
| RPL19 | 1 | 1 | 1 |
| RPL21 | 1 | 1 | 1 |
| RPL22 | 1 | 1 | 1 |
| RPL23 | 1 | 1 | 1 |
| RPL23A | 1 | 1 | 1 |
| RPL24 | 1 | 1 | 1 |
| RPL27 | 1 | 1 | 1 |
| RPL28 | 1 | 1 | 1 |
| RPL29 | 1 | 1 | 1 |
| RPL3 | 1 | 1 | 1 |
| RPL32 | 1 | 1 | 1 |
| RPL34 | 1 | 1 | 1 |
| RPL35 | 1 | 1 | 1 |
| RPL37A | 1 | 1 | 1 |
| RPL38 | 1 | 1 | 1 |
| RPL4 | 1 | 1 | 1 |
| RPL41 | 1 | 1 | 1 |
| RPL5 | 1 | 1 | 1 |
| RPL6 | 1 | 1 | 1 |
| RPL7 | 1 | 1 | 1 |
| RPL7A | 1 | 1 | 1 |
| RPL8 | 1 | 1 | 1 |
| RPLP0 | 1 | 1 | 1 |
| RPLP1 | 1 | 1 | 1 |
| RPS11 | 1 | 1 | 1 |
| RPS12 | 1 | 1 | 1 |
| RPS13 | 1 | 1 | 1 |
| RPS14 | 1 | 1 | 1 |
| RPS15 | 1 | 1 | 1 |
| RPS15A | 1 | 1 | 1 |
| RPS17 | 1 | 1 | 1 |
| RPS18 | 1 | 1 | 1 |
| RPS2 | 1 | 1 | 1 |
| RPS23 | 1 | 1 | 1 |
| RPS24 | 1 | 1 | 1 |
| RPS27 | 1 | 1 | 1 |
| RPS27A | 1 | 1 | 1 |

| | | | |
|---|---|---|---|
| RPS28 | 1 | 1 | 1 |
| RPS3 | 1 | 1 | 1 |
| RPS4X | 1 | 1 | 1 |
| RPS5 | 1 | 1 | 1 |
| RPS6 | 1 | 1 | 1 |
| RPS8 | 1 | 1 | 1 |
| RPS9 | 1 | 1 | 1 |
| VAMP8 | 1 | 1 | 1 |
| GPR18 | 1 | 1 | 1 |
| MVP | 1 | 1 | 1 |
| RPLP2 | 1 | 1 | 1 |
| SECTM1 | 1 | 1 | 1 |
| IFITM2 | 0 | 1 | 1 |
| LGALS3BP | 0 | 1 | 1 |
| PNP | 0 | 1 | 1 |
| RPL27A | 1 | 1 | 0 |
| RPL30 | 1 | 1 | 0 |
| RPL31 | 1 | 1 | 0 |
| RPL35A | 1 | 1 | 0 |
| RPL36 | 0 | 1 | 1 |
| RPL37 | 1 | 1 | 0 |
| RPL39 | 1 | 1 | 0 |
| RPL9 | 1 | 0 | 1 |
| RPS16 | 1 | 1 | 0 |
| RPS19 | 0 | 1 | 1 |
| RPS20 | 1 | 1 | 0 |
| RPS25 | 1 | 1 | 0 |
| RPS29 | 1 | 1 | 0 |
| RPS3A | 1 | 1 | 0 |
| RPS7 | 1 | 1 | 0 |
| RPSA | 0 | 1 | 1 |
| SERPING1 | 0 | 1 | 1 |
| SOCS1 | 0 | 1 | 1 |
| SOCS3 | 1 | 1 | 0 |
| SRI | 0 | 1 | 1 |
| B2M | 0 | 1 | 1 |
| BPGM | 0 | 1 | 1 |
| BST2 | 0 | 1 | 1 |
| CXCL11 | 0 | 1 | 1 |
| HIF1A | 1 | 1 | 0 |
| HLA-A | 0 | 1 | 1 |
| HLA-B | 0 | 1 | 1 |
| HLA-DRB1 | 0 | 1 | 1 |
| IFITM3 | 0 | 1 | 1 |
| IRF5 | 0 | 1 | 1 |
| ISOC1 | 1 | 0 | 1 |
| MYD88 | 1 | 1 | 0 |
| PSMB8 | 0 | 1 | 1 |
| PSME1 | 0 | 1 | 1 |
| PSME2 | 0 | 1 | 1 |
| RPL18A | 1 | 1 | 0 |
| RPS21 | 0 | 1 | 1 |
| TAP1 | 0 | 1 | 1 |

| | | | |
|---|---|---|---|
| TNFSF10 | 1 | 1 | 0 |
| TXNIP | 0 | 1 | 1 |
| UBE2L6 | 0 | 1 | 1 |
| VAMP5 | 1 | 0 | 1 |
| EIF4E3 | 0 | 0 | 1 |
| FGL2 | 0 | 0 | 1 |
| IL7 | 0 | 0 | 1 |
| LYSMD2 | 0 | 1 | 0 |
| PSMB2 | 0 | 1 | 0 |
| RBCK1 | 0 | 0 | 1 |
| RPS26 | 0 | 1 | 0 |
| SSPN | 0 | 1 | 0 |
| XCL1 | 1 | 0 | 0 |
| ADAR | 0 | 0 | 1 |
| ARL4A | 0 | 1 | 0 |
| BATF2 | 0 | 1 | 0 |
| C1R | 0 | 1 | 0 |
| CASP7 | 0 | 1 | 0 |
| CCL2 | 0 | 0 | 1 |
| CD274 | 0 | 1 | 0 |
| CD40 | 0 | 0 | 1 |
| CD69 | 0 | 0 | 1 |
| CD74 | 0 | 1 | 0 |
| CDKN1A | 1 | 0 | 0 |
| EIF2AK2 | 0 | 1 | 0 |
| FAS | 0 | 0 | 1 |
| FCGR1A | 0 | 1 | 0 |
| HELZ2 | 0 | 0 | 1 |
| IFI35 | 0 | 1 | 0 |
| IFIT2 | 0 | 0 | 1 |
| IFIT3 | 0 | 0 | 1 |
| IRF7 | 0 | 0 | 1 |
| LAP3 | 0 | 1 | 0 |
| LCP2 | 0 | 1 | 0 |
| LY6E | 0 | 1 | 0 |
| METTL7B | 0 | 0 | 1 |
| MT2A | 0 | 1 | 0 |
| MTHFD2 | 1 | 0 | 0 |
| NMI | 0 | 1 | 0 |
| OAS2 | 0 | 0 | 1 |
| OAS3 | 0 | 1 | 0 |
| OASL | 0 | 0 | 1 |
| PLA2G4A | 0 | 0 | 1 |
| PLSCR1 | 0 | 1 | 0 |
| PML | 0 | 0 | 1 |
| PSMB10 | 1 | 0 | 0 |
| PSMB9 | 0 | 1 | 0 |
| PTGS2 | 0 | 0 | 1 |
| RAPGEF6 | 0 | 1 | 0 |
| RNF31 | 1 | 0 | 0 |
| RSAD2 | 0 | 0 | 1 |
| RTP4 | 0 | 1 | 0 |
| SAMHD1 | 0 | 1 | 0 |

| | | | |
|---|---|---|---|
| SLAMF7 | 0 | 1 | 0 |
| SLC25A28 | 0 | 1 | 0 |
| SOD2 | 1 | 0 | 0 |
| ST3GAL5 | 1 | 0 | 0 |
| TRIM21 | 0 | 1 | 0 |
| ZBP1 | 0 | 0 | 1 |
| ITGB7 | 0 | 0 | 0 |
| PSMA3 | 0 | 0 | 0 |

Epithelial-to-mesenchymal transition-associated genes
Source:
https://www.gsea-msigdb.org/gsea/msigdb/cards/HALLMARK_EPITHELIAL_MESENCHYMAL_TRANSITION.html
1 = marker gene of the respective cluster, 0 = no marker gene of the respective cluster (see methods)

| | PT-New4 | TAL-New4 | DCT-New4 |
|---|---|---|---|
| CD44 | 1 | 1 | 1 |
| FSTL1 | 1 | 1 | 1 |
| ITGA2 | 1 | 1 | 1 |
| ITGA5 | 1 | 1 | 1 |
| LAMA3 | 1 | 1 | 1 |
| OXTR | 1 | 1 | 1 |
| TNC | 1 | 1 | 1 |
| CXCL1 | 1 | 1 | 0 |
| CXCL6 | 1 | 1 | 0 |
| GJA1 | 1 | 1 | 0 |
| ITGB1 | 1 | 1 | 0 |
| LAMC1 | 0 | 1 | 1 |
| LAMC2 | 1 | 1 | 0 |
| SAT1 | 0 | 1 | 1 |
| TFPI2 | 1 | 1 | 0 |
| TGM2 | 0 | 1 | 1 |
| THBS1 | 1 | 1 | 0 |
| TNFRSF11B | 1 | 0 | 1 |
| TPM1 | 1 | 1 | 0 |
| TPM4 | 1 | 1 | 0 |
| WNT5A | 1 | 1 | 0 |
| ABI3BP | 1 | 1 | 0 |
| ADAM12 | 1 | 0 | 1 |
| BMP1 | 1 | 1 | 0 |
| COL6A2 | 1 | 1 | 0 |
| COL8A2 | 1 | 0 | 1 |
| EMP3 | 1 | 1 | 0 |
| FBLN2 | 0 | 1 | 1 |
| FBN2 | 1 | 1 | 0 |
| FLNA | 1 | 1 | 0 |
| FSTL3 | 1 | 1 | 0 |
| LOXL2 | 1 | 1 | 0 |
| MCM7 | 1 | 1 | 0 |
| MMP3 | 1 | 0 | 1 |
| MYLK | 1 | 0 | 1 |
| NOTCH2 | 1 | 1 | 0 |
| PLAUR | 0 | 1 | 1 |
| QSOX1 | 1 | 1 | 0 |
| SERPINE1 | 1 | 1 | 0 |
| VCAN | 1 | 1 | 0 |
| COL16A1 | 1 | 0 | 0 |
| CXCL8 | 0 | 0 | 1 |
| DST | 0 | 1 | 0 |
| GPC1 | 1 | 0 | 0 |
| IL6 | 0 | 1 | 0 |
| ITGAV | 0 | 1 | 0 |
| LGALS1 | 1 | 0 | 0 |
| MATN2 | 0 | 1 | 0 |
| MMP14 | 1 | 0 | 0 |
| NID2 | 1 | 0 | 0 |
| NTM | 0 | 1 | 0 |
| PDLIM4 | 0 | 1 | 0 |
| PMEPA1 | 0 | 1 | 0 |
| PTHLH | 0 | 1 | 0 |
| SGCG | 0 | 1 | 0 |
| TIMP1 | 1 | 0 | 0 |
| TNFAIP3 | 0 | 1 | 0 |
| VIM | 1 | 0 | 0 |
| ACTA2 | 1 | 0 | 0 |
| AREG | 0 | 1 | 0 |

| | | | |
|---|---|---|---|
| BGN | 1 | 0 | 0 |
| CDH6 | 1 | 0 | 0 |
| COL12A1 | 1 | 0 | 0 |
| COL1A1 | 1 | 0 | 0 |
| COL1A2 | 1 | 0 | 0 |
| COL3A1 | 1 | 0 | 0 |
| COL5A1 | 0 | 1 | 0 |
| COL5A3 | 1 | 0 | 0 |
| COL6A3 | 1 | 0 | 0 |
| COL7A1 | 0 | 1 | 0 |
| COLGALT1 | 1 | 0 | 0 |
| CXCL12 | 1 | 0 | 0 |
| DPYSL3 | 1 | 0 | 0 |
| ECM1 | 0 | 0 | 1 |
| EDIL3 | 0 | 1 | 0 |
| EFEMP2 | 1 | 0 | 0 |
| ENO2 | 1 | 0 | 0 |
| FAS | 1 | 0 | 0 |
| FGF2 | 1 | 0 | 0 |
| FZD8 | 1 | 0 | 0 |
| GLIPR1 | 0 | 1 | 0 |
| IGFBP2 | 1 | 0 | 0 |
| IGFBP3 | 1 | 0 | 0 |
| JUN | 0 | 1 | 0 |
| LOX | 1 | 0 | 0 |
| LOXL1 | 1 | 0 | 0 |
| MAGEE1 | 1 | 0 | 0 |
| MATN3 | 1 | 0 | 0 |
| MMP1 | 1 | 0 | 0 |
| MMP2 | 1 | 0 | 0 |
| MXRA5 | 1 | 0 | 0 |
| NNMT | 1 | 0 | 0 |
| NT5E | 0 | 1 | 0 |
| PCOLCE | 1 | 0 | 0 |
| PCOLCE2 | 1 | 0 | 0 |
| PDGFRB | 0 | 1 | 0 |
| PFN2 | 0 | 1 | 0 |
| PLOD2 | 0 | 0 | 1 |
| PRRX1 | 0 | 0 | 1 |
| PVR | 0 | 1 | 0 |
| RGS4 | 1 | 0 | 0 |
| RHOB | 0 | 1 | 0 |
| SFRP4 | 0 | 0 | 1 |
| SNTB1 | 0 | 0 | 1 |
| SPARC | 1 | 0 | 0 |
| SPOCK1 | 0 | 1 | 0 |
| TGFB1 | 1 | 0 | 0 |
| THY1 | 1 | 0 | 0 |
| TIMP3 | 1 | 0 | 0 |
| TNFRSF12A | 0 | 1 | 0 |
| TPM2 | 1 | 0 | 0 |
| VCAM1 | 1 | 0 | 0 |
| VEGFC | 1 | 0 | 0 |
| BDNF | 0 | 0 | 0 |
| CADM1 | 0 | 0 | 0 |
| CAPG | 0 | 0 | 0 |
| COPA | 0 | 0 | 0 |
| CTHRC1 | 0 | 0 | 0 |
| ECM2 | 0 | 0 | 0 |
| FAP | 0 | 0 | 0 |
| IL15 | 0 | 0 | 0 |
| MGP | 0 | 0 | 0 |
| PPIB | 0 | 0 | 0 |
| PTX3 | 0 | 0 | 0 |
| SDC4 | 0 | 0 | 0 |

| | | | |
|---|---|---|---|
| SERPINH1 | 0 | 0 | 0 |
| TGFBR3 | 0 | 0 | 0 |
